# Coordinated Transcriptional Networks Program Organelle Expansion and Metabolic Flows for High Endothelial Morphology and Function

**DOI:** 10.1101/2025.06.19.660616

**Authors:** Yuhan Bi, Kevin Brulois, Aiman Ayesha, Menglan Xiang, Romain Ballet, Borja Ocón, Theresa Dinh, Nicole Lazarus, Manali Kun, Griselda Ramos, Hiroto Kawashima, Florea Lupu, Jonathan H Lin, Eugene C. Butcher, Junliang Pan

## Abstract

High endothelial cells (HECs) are specialized vascular gatekeepers that control lymphocyte entry into lymph nodes, a process essential for immune surveillance and adaptive responses. However, how HECs coordinate their high biosynthetic demands with the secretory apparatus to support immune function remains unclear. Using multi-omic approaches, we identify IRE1α–XBP1–centered transcriptional networks that, together with CREB3L2–associated gene programs, coordinate inter-organelle metabolic and secretory pathways required for glycosylation and assembly of peripheral node addressin (PNAd), a sulfated sialomucin critical for lymphocyte homing. Genetic or pharmacological perturbation of these pathways disrupts HEC morphology and lymphocyte recruitment during homeostasis and impairs HEC induction during inflammation. Parallel transcriptional programs operate in mucin-producing intestinal goblet cells, suggesting common regulatory pathways linking metabolism and sulfated mucin specialization with the associated expansion of secretory organelles across immune and barrier tissues. Thus, our findings identify transcriptional programs that coordinately scale metabolic pathways and secretory organelles to support the biosynthetic infrastructure underlying the morphology and immune trafficking functions of HEVs.

**Key points:**

XBP1-centered transcriptional networks coordinate metabolic and secretory programs for PNAd biosynthesis.
IRE1α or S1P inhibition flattens HEVs, prevents ectopic HEV formation and, reduces lymphocyte recruitment
Endothelial XBP1 deletion disrupts HEV morphology and lymphocyte trafficking.
Goblet cells share secretory transcriptional programs and regulatory logic with HEV.

## Introduction

All immune, vascular, and epithelial cells must dynamically integrate metabolic fluxes and organelle architecture to meet the biosynthetic demands of their specialized functions^1^. In the immune system, this integration is particularly critical, as cells must rapidly adapt to environmental cues during homeostasis, inflammation, and defense against pathogens. However, how metabolic pathways and organelle remodeling are logistically and transcriptionally coordinated to sustain specialized immune cell states remains incompletely understood^2–4^.

High endothelial cells (HECs) exemplify this challenge. As specialized vascular gatekeepers lining high endothelial venules (HEVs) in lymph nodes (LNs), HECs control lymphocyte entry, a process essential for immune surveillance and adaptive immune responses^5,6^. Beyond lymph nodes, HEC-like cells can emerge *de novo* in chronically inflamed tissues and tumors, where they facilitate lymphocyte infiltration and tertiary lymphoid structure formation. This trafficking function is mediated by peripheral node addressin (PNAd), a set of sialomucins densely decorated with O-glycans bearing 6-sulfo sialyl Lewis X (6-sulfo sLe^X^), the high-affinity ligand for L-selectin^7,8^. PNAd synthesis is metabolically intensive and depends on substantial secretory and glycosylation capacity, supported by expanded endoplasmic reticulum (ER), Golgi, and associated biosynthetic machinery that underlie the characteristic “high” morphology of HEVs and sustain the high-level, stepwise glycosylation and sulfation required for functional PNAd ligands^7,9–11^.These biosynthetic demands parallel those required for the production of sulfated mucins in intestinal goblet cells^12–14^, suggesting that conserved principles link metabolism, glycosylation, secretory organelle architecture, and specialized secretory functions across immune and barrier tissues.

PNAd⁺ HECs are induced in diverse inflammatory settings, including persistent infection, autoimmunity, graft rejection, and cancer, where their presence correlates with lymphocyte infiltration and local immune activation^15–17^. While lymphotoxin signaling and noncanonical NF-κB activation are required for HEV induction and maintenance^15,18–20^, these pathways do not explain how HECs acquire their distinct cuboidal morphology and biosynthetic capacity for PNAd assembly. Thus, despite their central role in immune surveillance and inflammation, the transcriptional programs that integrate organelle and metabolic remodeling to enable HEC function remain poorly defined.

Unlike template-driven processes such as transcription or translation, PNAd biosynthesis depends on coordinated metabolic fluxes and spatial organization^7,21^. This includes sustained production of nucleotide-activated sugar donors and 3′-phosphoadenosine 5′-phosphosulfate (PAPS), their transport into the Golgi lumen, and precise deployment of glycosyltransferases and sulfotransferases across Golgi cisternae^22^. While individual enzymatic steps have been characterized, how these metabolic pathways are transcriptionally programmed and coupled to organelle expansion *in vivo* remains unknown.

These biosynthetic demands suggest that specialized transcriptional programs must coordinate metabolic pathways with secretory capacity to sustain PNAd production and the distinctive morphology of HEVs. Here, we identify conserved IRE1α–XBP1–centered gene regulatory networks (GRNs) as key components of the biosynthetic programs that sustain HEV identity and immune trafficking. Computational and biochemical analyses further implicate the bZIP transcription factor CREB3L2 as a component of the regulatory architecture associated with these programs. Using single-cell transcriptomics, genetic perturbation, and phylogenomic analyses, we show that these GRNs upregulate nutrient transporters and enzymes required for nucleotide sugar and PAPS synthesis, promote endomembrane transport of precursors into the Golgi, coordinate spatial deployment of glycosyltransferases, and support the secretory capacity underlying PNAd production. Disruption of these pathways impairs PNAd expression, compromises lymphocyte homing, and prevents the ectopic emergence of functional HECs during inflammation. Parallel transcriptional programs in goblet cells support sulfated mucin biosynthesis and secretory specialization in barrier tissues^12^ ^14,23–26^. Together, these findings identify IRE1α–XBP1–centered transcriptional networks and implicate CREB3L2-associated pathways as key components of the biosynthetic programs that sustain HEV identity and immune trafficking, revealing a conserved regulatory logic underlying specialized immune and epithelial cell states.

## Results

### HECs Coordinately Express PNAd Biosynthesis, Secretory Organelle Machinery, and UPR Programs to Support HEC Specialization

The biosynthesis of PNAd-bearing mucins, a defining feature of HECs, requires sustained metabolic input and specialized organelle machinery, the expansion of which underlies the characteristic ’high’ endothelial morphology^27–29^. To define the metabolic programs supporting this process, we analyzed single-cell transcriptomes of murine lymph node (LN) endothelial subsets^28^ (Suppl. Fig. 1A). Differential expression analysis revealed that HECs uniquely upregulate a suite of genes required for PNAd biosynthesis, including nutrient transporters, metabolic enzymes for precursor synthesis, Golgi-resident transporters, glycosyltransferases and sulfotransferases that mediate stepwise glycan assembly (Fig. 1A, Suppl. Fig. 1B). Among the top 2,000 differentially expressed genes (DEGs) enriched in HECs relative to other endothelial subsets, nearly all key regulators of the inter-organelle metabolic pathways required for PNAd synthesis were represented (Fig. 1A–C, DEG list provided in Table S1). These findings indicate that HECs transcriptionally scale metabolic pathways to meet the biosynthetic demands of PNAd production.

**Fig. 1.**
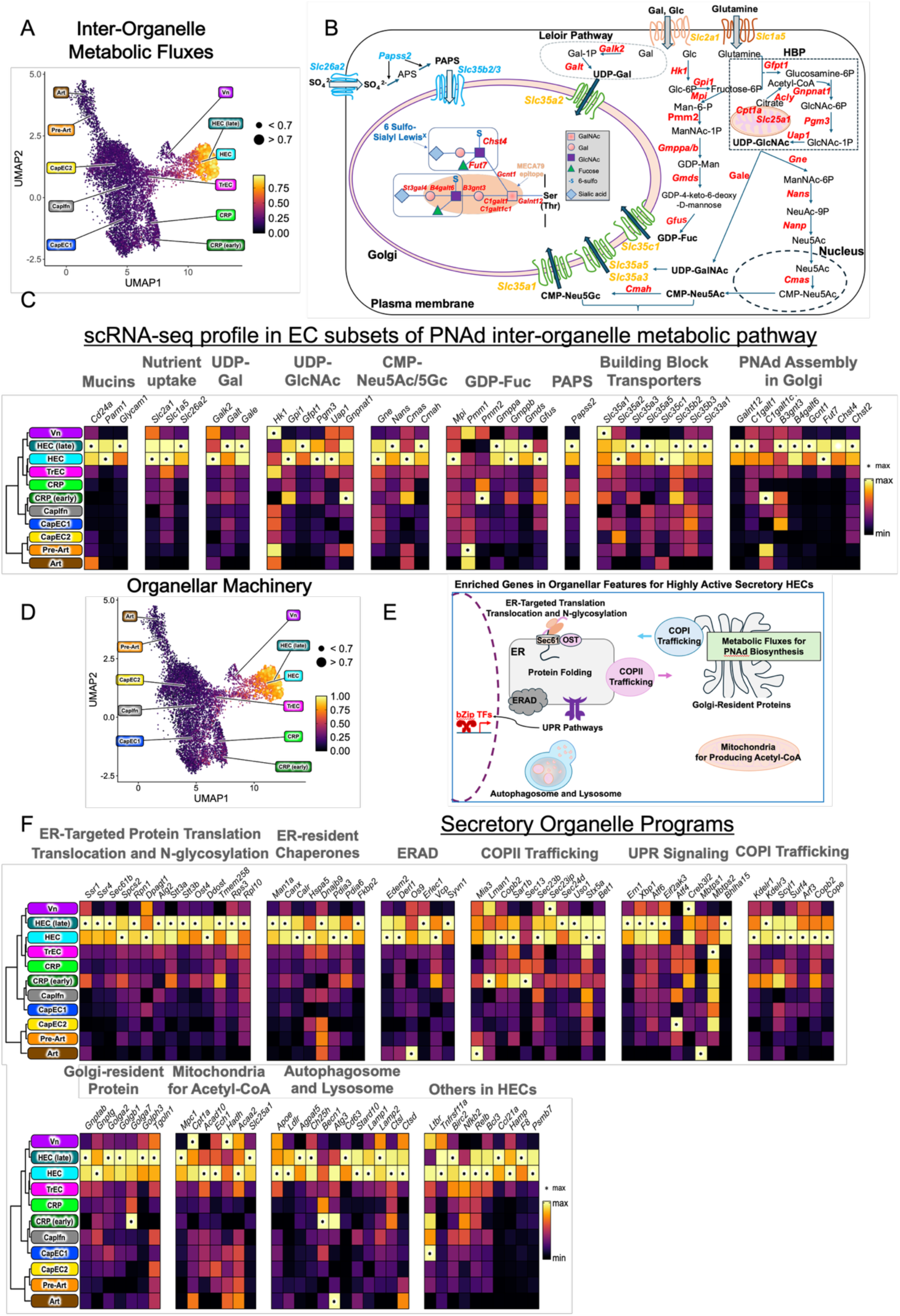
HECs Coordinately Express PNAd Biosynthesis, Secretory Organelle Machinery, and UPR Programs to Support HEC Specialization. **A.** UMAP projection of scRNA-seq data from murine peripheral LN ECs^28^, showing module scores that summarize the expression levels of genes controlling metabolic fluxes for PNAd synthesis in HECs compared to other EC subsets. Genes used for these modules are provided with expression values in C. **B.** A schematic chart depicts the genes and flow of metabolic fluxes, starting with nutrient uptake and building block production, followed by their transport into the Golgi lumen, and culminating in the stepwise assembly of PNAd^7^ on mucins within Golgi stacks. Transporters and enzymes involved in each step are color-coded in yellow and red, respectively, with those specifically involved in sulfation shown in blue. **C.** Heat maps showing the enriched expression of genes in HECs that coordinate metabolic fluxes essential for PNAd biosynthesis. **D.** UMAP projection of scRNA-seq data from murine peripheral LN ECs, showing module scores that summarize the expression levels of genes controlling organellar features. Mean expression profiles of the genes used for these module scores are provided in F. **E.** Schematic chart illustrates the organellar protein machineries and their functions that shape the distinctive “high” cuboidal morphology of HECs. **F.** Heat maps illustrate the HEC-enriched expression of genes associated with organellar structures and their functions. Values in heatmaps **C** and **F** are mean expression from three biological replicates and are shown on a color scale from min to max, with an asterisk (*) indicating the maximum value.

These upregulated metabolic gene programs encompass every critical step of PNAd biosynthesis, from precursor generation to glycan assembly (Fig. 1B). PNAd glycans, including the 6-sulfo sialyl Lewis X (6-sulfo SLe^X) carbohydrate ligands for the lymphocyte homing receptor L-selectin, are assembled within the Golgi apparatus from nucleotide sugar and sulfate donors. HEC-enriched genes support the generation of these donors via multiple cytosolic pathways. The hexosamine biosynthetic pathway (HBP) generates UDP-GlcNAc and UDP-GalNAc from glucose, galactose, and glutamine through the actions of Gfpt1, Pgm3, Uap1, and Gale, relying on nutrient uptake (via SLC2A1, SLC1A5) and acetyl-CoA derived from mitochondrial fatty acid β-oxidation and the citrate-malate-pyruvate shuttle^21,30,31^. Additional donors—including UDP-Gal, CMP-Neu5Ac, GDP-Fuc, and PAPS—are synthesized via the Leloir, sialic acid, fucose and sulfate donor pathways^21,32,33^, with specialized Golgi-resident transporters (e.g., Slc35a1, Slc35b2/3, Slc35c1) directing these precursors into the Golgi lumen for glycan assembly^34^. PNAd biosynthesis initiates at cis-Golgi with Galnt12-mediated transfer of GalNAc to mucin core proteins such as Glycam1, CD34, Parm1, Podocalyxin, and CD24, followed by the stepwise action of glycosyltransferases and sulfotransferases including C1galt1, Gcnt1, B3gnt3, Fut7, Chst4, and St3gal4^7,27^, along the Golgi stacks.

The ’high’ endothelial morphology of HEC derives from the expansion of polyribosomes, ER, Golgi, mitochondria, and autophagolysosomal systems, providing the architectural framework for glyco-mucin synthesis and precursor transport between organelles. Alongside metabolic programs, HECs upregulate gene programs required for the generation and function of organelles that support high-output glycoprotein production (Fig. 1D–F). HEC-enriched genes include those for ER-targeted protein translation, translocation and cotranslational N-glycosylation (Ssr1, Sec61b, Spcs2, Rpn1, Dpagt1, Alg2, Stt3a, Stt3b, Ost4, Tmem258, Ddost)^35^, ER chaperones (Hspa5, Calr, Canx, Dnajb9, Pdia3, Pdia6)^36^, and ER-associated degradation (Edem2, Derl1, Os9, Erlec1, Vcp, Syvn1)^37^. Cargo trafficking is coordinated by COPII components (Mia3, Lman1, Sar1b, Sec23b, Sec24d)^38^ and COPI machinery (Kdelr1, Arf3, Copb2)^39^, while Golgi organization relies on scaffold proteins such as Golga2, Golgb1, and Golph3^40^. Additional HEC-enriched programs support mitochondrial fatty acid oxidation and citrate shuttling (Mpc1, Cpt1a, Acad10, Slc25a1, Acly)^41^ as well as membrane maintenance and autophagy (Apoe, Agpat5, Lamp1, Becn1)^42,43^. The inclusion of Slc35b1, an ATP/ADP exchanger that imports cytosolic ATP into the ER lumen in exchange for ADP to support protein folding, further illustrates the tight coupling of metabolic state with organelle function. Immunohistochemical datasets (Human Protein Atlas) reveal protein-level enrichment in HECs of multiple proteins predicted by gene expression, including proteins involved in metabolic flux (PASS2, PGM3, PMM2, CMAS), endomembrane transport (SLC35B2, SLC35C1), PNAd biosynthesis (PARM1, GALNT1, CHST4), organelle homeostasis (EDEM2, STX5A, KDELR2, GOLPH3), and the unfolded protein response (SLC35B1, HSPA5) (Data S1). Together with prior studies^10,44–47^, both transcriptional and proteomic studies support a coordinated upscaling of metabolic and organellar machinery in HEC.

The expansive upregulation of these biosynthetic, metabolic, and organelle programs is accompanied by selective activation of transcriptional regulators associated with the UPR. HECs show marked enrichment of the IRE1α–XBP1 branch, including elevated expression of Hspa5, Ern1 (IRE1α), spliced Xbp1(Fig. F). Expression of Atf6, but not Atf6b, is enriched. Notably, the noncanonical UPR transcription factor Creb3l2, but not other Creb3 family members, is selectively expressed in HECs, together with the S1P/S2P proteases (Mbtps1 and Mbtps2) required for CREB3L2 activation. In contrast, the PERK-ATF4 branch is selectively suppressed in HECs, as indicated by low Atf4 expression and elevated Dnajc3, an inhibitor of PERK signaling^48^. While UPR signaling is classically associated with stress mitigation, these branches seem to exert distinct metabolic outputs. IRE1α–XBP1 and CREB3L2 have been reported to drive adaptive anabolic programs that couple nutrient sensing to ER vesicle trafficking^49,50^, glycosylation^51^ and protein translation^52^ while expanding ER and secretory capacity^53^. ATF6 promotes ER proteostasis through induction of ER-associated degradation components^54,55^, whereas PERK broadly represses global translation to reduce ER load^56^. Thus, the engagement of IRE1α-XBP1 and CREB3L2 pathways in HEC, together with ATF6 activation and suppression of PERK signaling, may establish a specialized UPR configuration for PNAd synthesis and lymphocyte recruitment.

Taken together, these single-cell analyses reveal that PNAd biosynthesis is embedded within a coordinated metabolic and regulatory network spanning multiple organelles in HECs. This integrated program is predicted to coordinately scale metabolic pathways, secretory organelles, and UPR signaling, thereby linking glycan biosynthesis to the distinctive morphology and lymphocyte-recruiting function of the specialized “high” endothelial state.

### Goblet Cells Share Gene Programs for Sulfated Mucin Production and Organelle Expansion with HECs

Intestinal goblet cells (GCs) produce sulfated, complex carbohydrate–coated mucins that constitute a central component of the mucosal barrier, and defects in goblet cells or mucus integrity are associated with colitis^26^. Notably, Galnt12 and Chst4, highly enriched in HECs, are also selectively abundant in colonic GCs at both transcript and protein levels^26^. GALNT12 initiates mucin-type O-glycosylation and, through its lectin domain, preferentially produces heavily O-glycosylated structures like PNAd and mucus^57^. Sulfation of GC-derived mucins is catalyzed by CHST4^58,24^, a sulfotransferase that is also required for PNAd biosynthesis in HEC^47^. We asked whether GCs employ similar molecular pathways to couple sulfated mucin synthesis with organelle function, by analyzing single-cell RNA sequencing data from mouse intestinal epithelial cells in the Tabula Muris atlas^59^. GCs exhibit a strikingly similar molecular profile, sharing nearly all gene sets involved in metabolic and secretory pathways supporting sulfated O-glycan production on mucins (Suppl. Fig. 2A and B). This convergence is particularly evident within precursor biosynthetic pathways and transporters. GCs also display enhanced expression of genes for ER- and Golgi-resident protein machineries that coordinate and streamline cargo trafficking along the secretory pathway (Suppl. Fig. 2C). Finally, GCs appear to engage analogous regulatory mechanisms, characterized by enriched expression of the Ern2-Xbp1 signaling axis^25,60,61^, Creb3l1^24^, and the intramembrane proteases Mbtps1^62^. Together, these findings suggest that GCs and HECs share conserved metabolic and secretory gene programs, secretory organelle machinery deployment, and regulatory logic to support high-output glycosylation.

### hXBP1 and CREB3L2 Are Predicted to Coordinate Conserved Transcriptional Networks that Couple PNAd Biosynthesis to Organelle Scaling

Having identified a shared metabolic and organellar gene program in HECs and GCs, we next sought to define the upstream regulatory networks that drive this convergence. Gene set overlap analysis across endothelial and intestinal epithelial subsets revealed that HECs and GCs share a highly correlated transcriptional signature (Fig. 2A). Among the top 2,000 DEGs enriched in HECs, 346 were co-upregulated in GCs, the majority of which encode components of metabolic pathways and organellar protein machineries (Suppl. 2B and C). This overlap suggests that a common set of *cis*- and *trans*-acting factors underlies their shared transcriptional architecture.

**Fig. 2.**
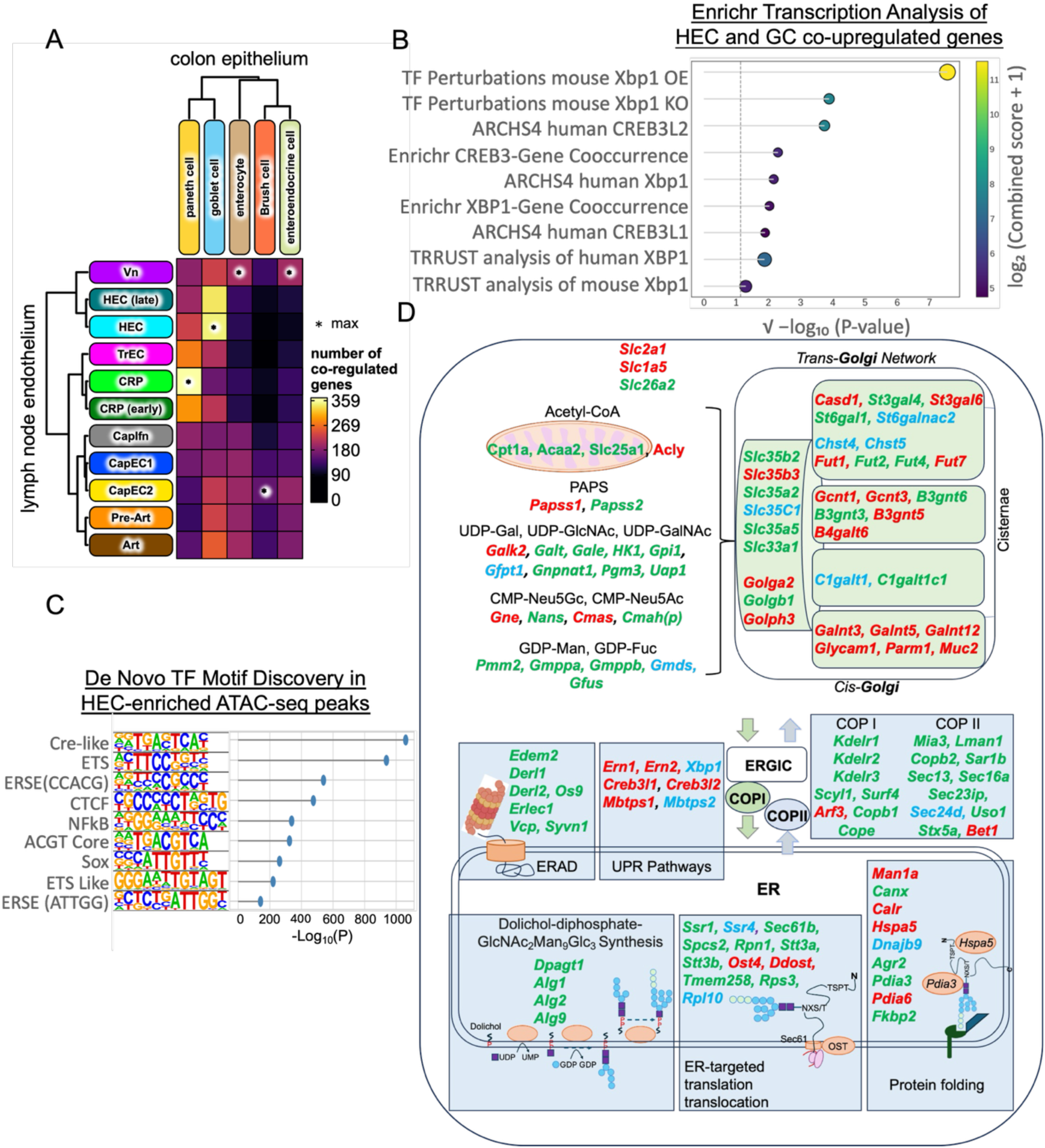
XBP1 and CREB3L2 Are Predicted to Link Conserved Networks Integrating PNAd Biosynthesis and Organelle Scaling. A. Gene set overlap of DEGs (top 2000 DEGs for individual cell subsets, computed independently for lymph node endothelium and intestinal epithelium) revealed that HECs and intestinal GCs share a transcriptional signature enriched for genes involved in integrating sulfated O-glycan synthesis with organellar machineries. B. Transcription factor regulated gene set enrichment among the 346 co-upregulated genes shared between HECs and GCs identified XBP1 and CREB3/CREB3L2 as the top candidate regulators. Enrichment was performed using the Enrichr platform across the indicated datasets (the list of 346 shared genes is provided in Table S2). Statistical significance is shown on the x-axis as √ (–log₁₀ *P*), compressing the dynamic range to facilitate comparisons. Dot size represents Odds Ratio (scaled linearly from ∼3.1 to 22.5), and dot color indicates log₂ (Combined Score + 1). Grey lines denote individual signatures, and the dashed vertical line indicates *P* = 0.05. C. *De novo* motif discovery using the HOMER platform on HEC-enriched ATAC-seq peaks (compared with capillary ECs) revealed significant enrichment of three major motif classes: a CRE-like motif (TGAN₁TCA) associated with CREB3L2 binding; canonical endoplasmic reticulum stress response elements (ERSE; ATTGG–N(≥1)–CCACG), which mediate binding of NFYA/B and spliced XBP1 (XBP1s); and a bZIP-family consensus motif containing a central ACGT core recognized by both XBP1 and CREB3L2), among the most significantly enriched features. Shown is a lollipop plot ranking the top motifs by significance (−log₁₀P). D. Genes harboring CRE–like motifs (TGAN₁–₂TCA) specific for CREB3L2 are highlighted in red, those containing the shared XBP1/CREB3L2 “ACGTG” core sequence in green, and genes with multiple conserved sites—including CRE–like, ACGTG core, and/or ERSE elements—are shown in blue. These conserved motifs, situated within noncoding regulatory modules, are provided at single-nucleotide resolution in Data S2.

To identify candidate upstream transcription factors (TFs), we performed enrichment analysis on these 346 shared genes using the Enrichr platform^63,64^. Across multiple datasets—including TF perturbation signatures, co-expression networks, and curated TF-gene associations— the bZIP family factors XBP1 and CREB3/CREB3L2 consistently emerged as the top regulatory candidates (Fig. 2B, Suppl. Fig. 3A, Table S2). Independent analyses of HEC- and GC-specific gene sets further supported a central role for XBP1 and CREB3L2 in coordinating the metabolic and organellar programs that enable sulfated mucin biosynthesis (Suppl. Fig. 3B–C, Table S3-4).

To identify candidate cis-regulatory elements underlying this transcriptional control, we performed de novo motif discovery using the HOMER platform on both HEC-enriched ATAC-seq peaks (Fig. 2C; Data S5) and HEC-enriched gene sets (Suppl. Fig. 3D)^65^. This analysis revealed significant enrichment of three motif classes: a CRE-like motif (TGAN₁TCA) associated with CREB3L2 binding; canonical endoplasmic reticulum stress response elements (ERSE; ATTGG–N(≥1)–CCACG), which mediate binding of NFYA/B and spliced XBP1 (XBP1s); and a bZIP-family consensus motif containing a central ACGT core recognized by both XBP1s and CREB3L2)^66,67^, among the most significantly enriched features. Comparative phylogenomics using Multiz alignments from 58 eutherian mammals and PhyloP conservation scores from the UCSC Genome Browser^68–70^, further demonstrated that conserved noncoding genomic regions across multiple gene sets harbor canonical sites associated with XBP1 and CREB3L2 activity. These include shared “ACGT” core motifs capable of binding both TFs)^66,67^, as well as factor-specific elements: ERSE motifs that selectively bind XBP1s^71^ but not CREB3L2, and CRE–like motifs (TGAN₁₋₂TCA) preferentially recognized by CREB3L2^72^ but not XBP1s. These elements were detected in multiple genes required for PNAd biosynthesis in HECs, sulfated mucin production in GCs, and secretory organelle function, as summarized in Fig. 2D.

Single-nucleotide–resolution maps of these conserved motifs, embedded within the relevant noncoding regulatory modules, are provided in Data S2 based on aligned genomic sequences from human, chimpanzee, monkey, rat, and mouse. Notably, XBP1- and CREB3L2-associated motifs frequently co-occur with binding elements for additional regulatory factors, including E-box, ETS, KLF/Sp1, and NF-κB, indicating integration with broader transcriptional architectures. Conserved XBP1- and CREB3L2-associated motifs are also present within genes encoding cell state–responsive regulators (Ltbr, Tnfrsf11a, Nfkb2, Relb, Nod1, Nod2, and Il13ra1) and lineage-determining factors (Nr2f2^73^ and Spdef^26^), suggesting that these bZIP-driven networks interface with context-dependent programs to shape HEC and GC specialization.

Beyond HECs and GCs, analysis of Human Cell Atlas single-cell transcriptomic data indicate that stomach plasma cells deploy a remarkably similar metabolic program for precursor nucleotide-sugar biosynthesis and cell-enriched glycosyltransferases including MGAT1, MGAT2, MGAT3, ST6GAL1 and FUT8, together with regulatory logic resembling that of HECs and GCs (Suppl.Fig. 4 and Table S6). This is consistent with prior studies showing that high glucose uptake and active hexosamine biosynthetic pathway flux are essential for persistence of long-lived antibody-secreting plasma cells^74,75^. A similar pattern was also evident in pituitary glandular cells, which produce sulfoglycosylated peptide hormones through glycosyltransferases such as B4galnt3/4 and Chst8^76^ (Table S6).

Together, these analyses identify key transcriptional regulators of the metabolic and secretory programs of HECs. The conservation of XBP1 and CREB3L2 binding motifs across genes embedded in these programs suggests that an integrated network links glycosylation capacity to secretory pathway expansion through evolutionarily conserved cis-regulatory modules. Consistent with this model, we identify conserved XBP1- and CREB3L2-binding sites in genes governing specialized glycosylation in pituitary glandular cells (e.g., B4galnt3/4, Chst8) and long-lived plasma cells (Fut8, St6gal1) (Data S3). Supporting this framework, prior studies in goblet, plasma B and pituitary secretory cells demonstrate essential roles for IRE1β (ERN2)^77,78^, XBP1^25,53^, CREB3L1/2^24,52^, and the S1P/S2P proteases (MBTPS1/2)^62,79^ in establishing lineage-specific secretory pathways and high-level production of glycosylated proteins.

### XBP1 and CREB3L2 Activate Conserved Enhancer–Promoter Elements of PNAd Biosynthetic Genes

Having identified conserved XBP1 and CREB3L2 binding motifs across genes involved in PNAd biosynthesis and organelle function, we next tested whether these motifs and transcription factors could directly regulate gene expression. We focused on glycosyltransferases and nucleotide sugar biosynthetic enzymes localized to the Golgi and cytosol, including *Chst4* (sulfation), *Fut7* (fucosylation), *C1galt1* (core 1 O-glycan synthesis), *Pgm3* (UDP-GalNAc production via the HBP)^80^, *Nans* (CMP-Neu5Ac synthesis)^81^, *Slc35c1* (GDP-Fuc transporter)^34^ and *Golph3* (a Golgi Resident Protein that maintains the spatial organization of glycosyltransferases) ^40^ (Fig. 3A).

**Fig. 3.**
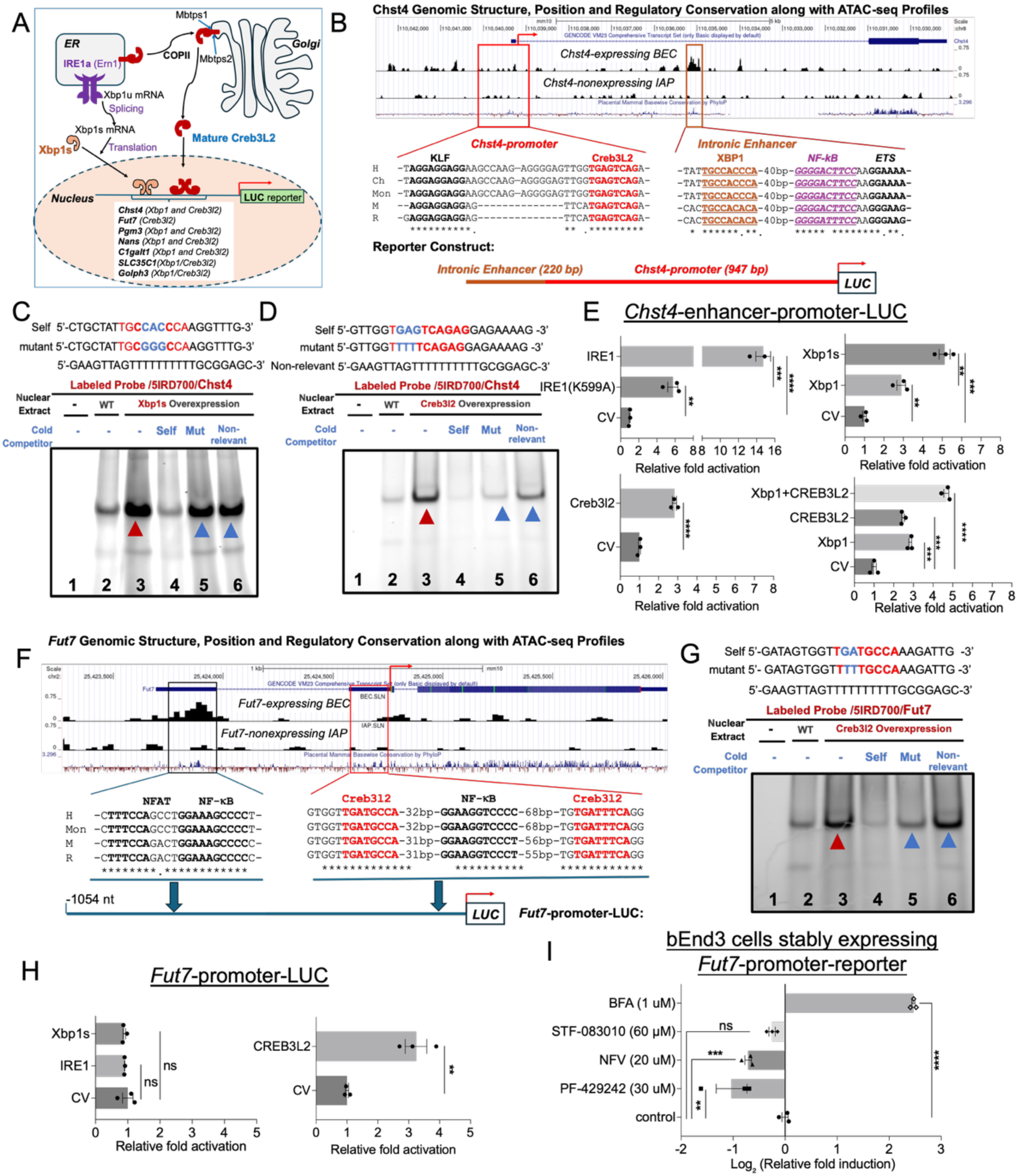
XBP1 and CREB3L2 Bind and Activate Conserved Regulatory Elements of PNAd Biosynthetic Genes. **A.** Diagram of the IRE1α–XBP1 and CREB3L2 activation pathways and their downstream transcriptional programs supporting PNAd biosynthesis. **B.** Evolutionarily conserved XBP1- and CREB3L2-binding motifs within the *Chst4* locus colocalize with ATAC-accessible chromatin in Chst4⁺ endothelial cells. The *Chst4* enhancer–promoter reporter construct positions the XBP1-dependent intronic enhancer upstream of the CREB3L2-binding promoter. **C–D.** EMSA using fluorescently labeled Chst4 probes demonstrated that XBP1 and CREB3L2 bind to their respective motifs within the *Chst4* regulatory region. In (C), a probe containing the candidate XBP1-binding motif formed a specific DNA-protein complex with nuclear extracts from wild-type 293T cells, which was further enhanced by XBP1s overexpression. In (D), a probe encompassing the predicted CREB3L2-binding motif produced a specific complex with nuclear extracts from CREB3L2-overexpressing cells. **Lane 1**, free probe; **Lane 2**, nuclear extracts from wild-type 293T cells; **Lane 3**, nuclear extracts from XBP1s-overexpressing (C) or CREB3L2-overexpressing (D) cells. Binding specificity was confirmed by competition with a 50-fold excess of unlabeled wild-type probe (Self, **Lane 4**), but not with unlabeled mutant (**Lane 5**) or non-relevant (**Lane 6**) probes. Probe sequences are shown above each gel image. Unbound labeled probe (**Lane 1**) migrated close to the dye front (cropped in image). Shifted protein–DNA complexes are indicated by arrowheads. **E.** IRE1, XBP1s, and CREB3L2 cooperatively activate the *Chst4* enhancer–promoter reporter in transiently transfected 293T cells. Results are shown as fold induction relative to CV (Control Vector). **F.** Conserved CREB3L2-binding motifs in the *Fut7* locus map to regions of open chromatin. The positions of these motifs are shown in the Fut7 promoter reporter construct. **G.** A labeled *Fut7* probe encompassing the predicted CREB3L2-binding motif formed a specific DNA–protein complex with nuclear extracts from CREB3L2-overexpressing 293T cells. EMSA conditions and lane assignments are as described in (C–D); shifted protein–DNA complexes are indicated by arrowheads. **H.** CREB3L2, but not IRE1α or XBP1s, selectively activates the *Fut7* promoter–driven luciferase reporter in transiently transfected 293T cells. Results are shown as fold induction relative to CV (Control Vector). **I.** CREB3L2 activation by brefeldin A (BFA) in endothelial cells increases *Fut7* reporter activity, whereas S1P/S2P protease inhibitors PF-429242 and nelfinavir suppress CREB3L2-dependent activation. In contrast, IRE1α inhibition (STF-083010) has no effect. Assays were performed in bEnd.3 cells stably harboring a Fut7 promoter reporter. Results are shown as log₂ (fold induction relative to control). **E, H** and **I**. Data are from three experiments (independent transfections) and are shown with mean ± S.E.M. Statistical significance was determined by unpaired two-tailed Student’s *t*-test comparing each condition to control vector (CV) (E, H) or vehicle control (I); Welch’s correction was applied when variances were unequal. *: p value < 0.05; **: p value < 0.01; ***: p value < 0.001; ****: p value < 0.0001.

To assess direct transcriptional regulation, we generated luciferase reporter constructs driven by enhancer and promoter regions from these genes, containing the evolutionarily conserved XBP1 and/or CREB3L2 binding sites identified in our motif analysis. Fig. 3B depicts the positions of XBP1 and CREB3L2 binding motifs within the *Chst4* locus. These elements coincide with regions of accessible chromatin in peripheral lymphoid, *Chst4*- and *Fut7*-expressing ECs, as indicated by ATAC-seq, but are absent in non-expressing ITGA7⁺ pericytes (IAPs). Electrophoretic mobility shift assays (EMSAs) demonstrated site-specific binding of recombinant spliced XBP1 (XBP1s) or active CREB3L2 to these motifs (Fig. 3C–D). Transient transfection–based reporter assays in 293T cells showed that overexpression of IRE1α, XBP1s, or an active form of CREB3L2 significantly transactivated the *Chst4* enhancer–promoter. In contrast, an IRE1α kinase mutant (IRE1α K599A^82,83^), or overexpression of unspliced Xbp1 exhibited markedly reduced activity (Fig. 3E). The IRE1α K599A mutation prevents activation of the IRE1 RNase domain required for XBP1 mRNA splicing. Protein expression of the IRE1α, XBP1, and CREB3L2 constructs used in the reporter assays was confirmed by western blot (Suppl. Fig. 5A, C–E). Wild-type IRE1α and IRE1α K599A were expressed at comparable levels; however, the mutant induced substantially less XBP1 mRNA splicing (Suppl. Fig. 5 B). Co-expression of XBP1s and CREB3L2 resulted in enhanced activation, indicating cooperative control of *Chst4* transcription by these two regulatory pathways (Fig. 3E).

The *Fut7* promoter also maps to regions of accessible chromatin in these lymphoid ECs (Fig. 3F) and contains two conserved CREB3L2-binding sites but lacks Xbp1 motifs. This specificity was supported by EMSA analyses showing selective CREB3L2 binding to the identified motifs (Fig. G) and confirmed selective *Fut7* promoter activation by CREB3L2 but not XBP1s in transient transfection-based reporter assays in 293T cells (Fig. 3H). Moreover, using bEnd.3 cells stably integrated with a *Fut7* promoter-driven reporter (Fig. 3I), we found that CREB3L2 activation by Brefeldin A (BFA)^84^, which induces ER-Golgi fusion and CREB3L2 processing, markedly upregulated reporter activity in bEnd3 cells. In contrast, the reporter activity was suppressed by S1P (PF-429242) and S2P (nelfinavir) protease inhibitors^85,86^, which block CREB3L2 activation; while the IRE1α inhibitor STF-083010^87^ had no effect.

Finally, XBP1s and CREB3L2 cooperatively enhanced transcription from reporter constructs driven by the Pgm3, Nans, and C1galt1 enhancer–promoter elements, each of which contains multiple conserved binding sites (Suppl. Fig. 6A–C). In contrast, overexpression of XBP1s and/or CREB3L2 did not produce additive or cooperative activation of enhancer–promoter constructs containing only a single binding site, such as the SLC35C1 and GOLPH3 reporters (Suppl. Fig. 6D–E). These findings suggest that XBP1s and CREB3L2 exert distinct, additive, or cooperative transcriptional effects depending on motif composition.

Together, these results demonstrate that XBP1s and CREB3L2 can directly regulate transcription of genes involved in precursor metabolism and glycan biosynthesis through conserved regulatory motifs, extending their regulatory scope beyond canonical ER-localized targets.

### Pharmacological Inhibition of IRE1α- or the S1P Protease Flattens HEVs, Reduces PNAd Expression, and Impairs Lymphocyte Homing

Having established that XBP1 and CREB3L2 can directly activate PNAd biosynthetic genes through conserved regulatory elements (Fig. 3), we next asked whether perturbation of these pathways disrupts HEV morphology and function in vivo. To test this, we pharmacologically perturbed IRE1α–XBP1 signaling and S1P-dependent transcription in wild-type C57BL/6 mice using inhibitors validated in our cell culture studies. Mice were treated with STF-083010, a selective inhibitor of IRE1α RNase activity that blocks generation of spliced Xbp1^87^, or PF-429242, an inhibitor of the Site-1 protease (S1P) required for proteolytic activation of several transcription factors, including CREB3 family members such as CREB3L2^88^.

Following treatment, peripheral lymph nodes (PLNs) were analyzed by whole-mount immunofluorescence (IF) using anti-CD31 to label the vasculature and MECA-79 to detect the extended core 1 epitope of PNAd^89,90^. PLNs from mice treated with STF-083010 every other day for one week or PF-429242 daily for two weeks displayed marked structural abnormalities, including reduced node size, flattened and elongated HEV walls, and diminished branching (Fig. 4A and D). PNAd expression was significantly reduced, as quantified by PNAd⁺ HEV surface area relative to total vasculature (Fig. 4B and E).

**Fig. 4.**
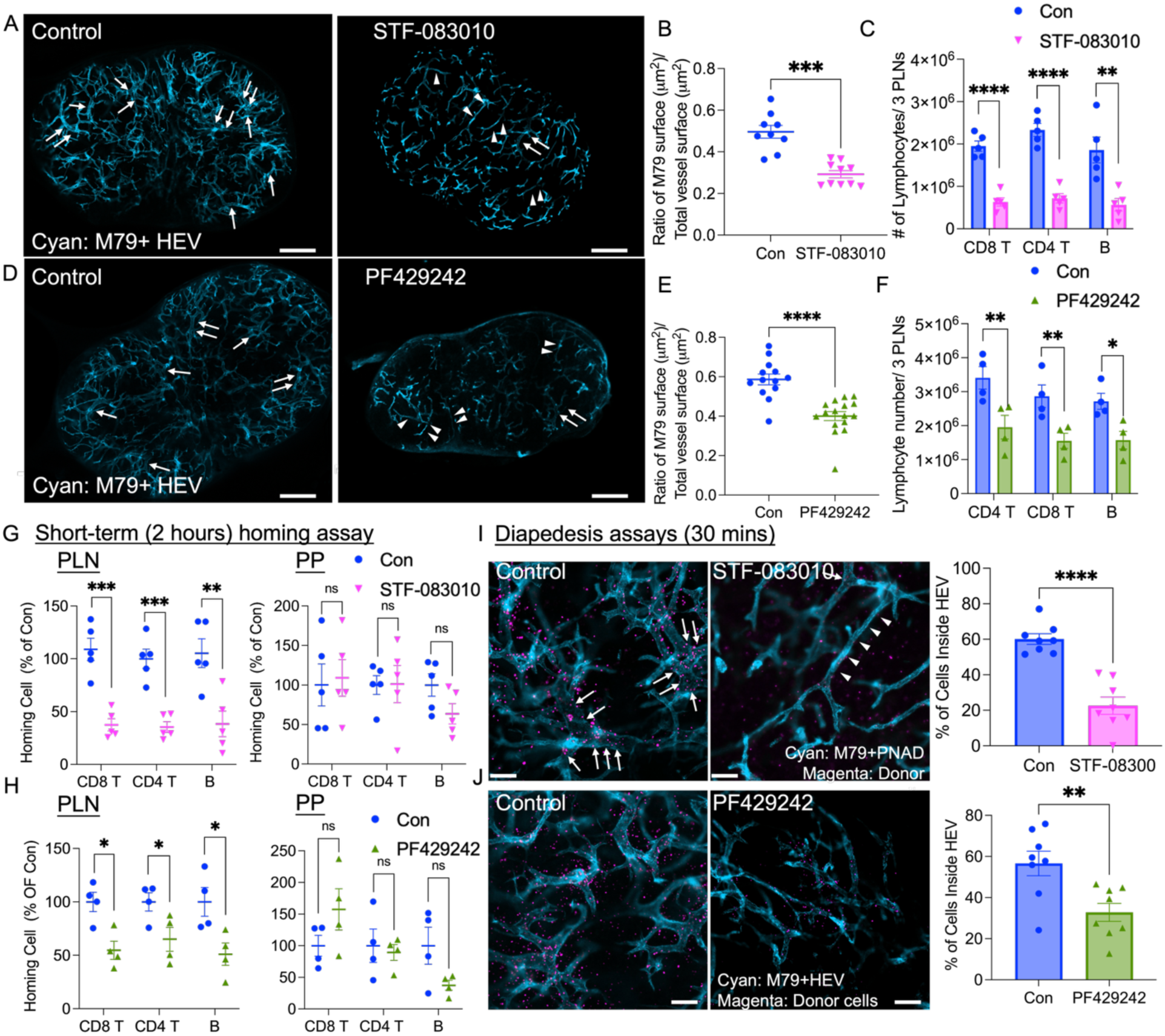
Pharmacological Inhibition of IRE1α–XBP1 Signaling or S1P-Dependent Transcription Flattens HEVs and Impairs Lymphocyte Homing A and. **D.** Representative whole-mount immunofluorescence (IF) images of PLNs from mice treated with IRE1α inhibitor STF-083010 (**A**), Creb3l2 pathway inhibitor PF429242 (**D**) and control mice. Mice were injected intravenously (i.v.) with antibodies to label PNAd (MECA-79, cyan). Arrows denote thick HEV segments; arrowheads indicate thin, elongated HEVs. Scale bar: 500 μm. **B and E.** Quantification of PNAd⁺ (MECA-79⁺) HEV surface area relative to total vasculature by *Imaris* analysis (See Star Methods). Data represent 1 field per PLN, 2–6 fields per mouse from 3 independent experiments (B: 1 control and 1 STF-083010–treated mouse per experiment; E: 1 control and 1 PF429242 treated mouse per experiment). **C and F.** Lymphocytes count from 3 pooled PLNs (Inguinal, Axillary, and Branchial PLN) per STF-083010-treated (**C**), PF429242-(**F**) and control mouse. **G and H.** Flow cytometry analysis of donor lymphocyte homing to PLNs and PPs 2 hours post-transfer in control vs. STF-083010– (**G**)and PF429242- (**H**) treated mice, expressed as percentage of control group mean localization ratio. **C and G:** Results are shown as mean ± SEM for n = 5 mice per group. **F and H:** Results are shown as mean ± SEM for n = 4 mice per group. **I and J. Left:** Representative IF images of PLNs 30 minutes after lymphocyte injection for short-term diapedesis assay. Donor lymphocytes (CFSE-labeled, magenta), PNAd (cyan). Arrows highlight thick HEV segments with bound donor cells; arrowheads denote thin, elongated HEVs. Scale bar: 100 μm. **Right:** Percentage of donor cells localized inside MECA-79⁺ HEVs, quantified by *Imaris*. Data collected from 1 field/PLN, 2–3 fields/mouse across 3 independent experiments (1 control and 1 STF-083010– (**I**) or PF429242– (**J**) treated mouse per experiment). All plotted data are shown as mean ± SEM. *: p value < 0.05; **: p value < 0.01; ***: p value < 0.001; ****: p value < 0.0001. Two-tailed unpaired t-test.

To determine how rapidly HECs lose their specialized phenotype upon pathway inhibition, we performed time-course analyses for each regimen. STF-083010 was administered every other day for one week and lymph nodes were analyzed on Days 1, 4, and 8. Despite rapid inhibition of Xbp1 mRNA splicing within ∼24 hours *in vivo*^87^, no overt changes were observed at day 1, whereas modest reductions in PNAd expression and early morphological alterations appeared by Day 4, followed by pronounced venular flattening and marked PNAd loss by Day 8 (Supplemental Fig. 7A). PF-429242 was administered daily for two weeks, with analysis on Days 1, 4, 8, 12, and 15. Similarly, PF-429242 rapidly inhibits S1P protease activity within ∼24 hours^85^, yet minimal changes were observed before Day 8, when flattening and loss of PNAd emerged and persisted through Days 12–15 (Supplemental Fig. 7B). Together, inhibition of either pathway induced progressive regression of the high endothelial phenotype, with reproducible loss of ‘highness’ and PNAd loss within ∼1 week, consistent with the timeframe reported following surgical occlusion of afferent lymphatic vessels^11^.

These defects in HEV were accompanied by a substantial decrease in total lymphocyte numbers within the PLNs (Fig. 4C and F). To test whether lymphocyte homing was impaired, we next assessed homing efficiency using adoptive transfer assays. Splenocytes were injected i.v. and quantified in recipient LNs 2 hours later. Lymphocyte recruitment to PLNs (Fig.4 G, H) and mesenteric lymph nodes (MLNs) (Suppl. Fig. 8A and C, left) were significantly reduced in both inhibitor-treated groups, whereas lymphocyte homing to the spleen, and T cell homing to Peyer’s patches (PPs)—where HEVs are MAdCAM-1⁺/PNAd⁻^91^—remained unaffected (Fig.4 G, H, Suppl. Fig. 8A and C, right). In contrast, B cell homing to PP was reduced by both inhibitors (Fig. 4G and H, right): Although this reduction did not reach statistical significance, it may reflect effects on IRE1α–XBP1 and S1P-dependent transcriptional programs involved in the synthesis of the B-cell mucosal addressin (BMAd), a St6gal1-dependent glycotope that binds CD22 and supports preferential B-cell homing to Peyer’s patches ^27,92^. Consistent with the role of HEV in determining endogenous as well as injected lymphocyte accumulation, the donor-to-recipient lymphocyte ratios in LN were comparable between treatment and control groups (Suppl. Fig. 8B, D). We also examined the effects of IRE1α and S1P inhibition on short-term lymphocyte arrest and diapedesis across HEVs^92^. CellTracker-labeled splenocytes were intravenously transferred into treated and control mice, and their localization (in HEV and in the surrounding parenchyma) was assessed 30 minutes later by confocal microscopy. LNs from both STF-083010- and PF429242–treated mice showed marked reductions in HEV-bound and extravasated lymphocytes (Fig. 4I for STF-083010; Fig. 4J for PF429242), and a significant decrease in the ratio of extravasated to HEV bound cells.

Because the S1P protease activates multiple transcription factors—including CREB3 family members, ATF6, and SREBPs—PF-429242 inhibition may affect several regulatory pathways. Among these, CREB3 family transcription factors have been implicated in secretory pathway regulation, whereas ATF6 primarily promotes ER proteostasis through induction of ER-associated degradation. Perturbation of these programs would be expected to impair secretory capacity and glycoprotein biosynthesis, consistent with the reduced PNAd expression and HEV flattening observed following PF-429242 treatment.

Taken together, these data indicate IRE1α–XBP1 signaling and S1P-dependent transcriptional programs are required to maintain PNAd expression and function, and the characteristic “high” endothelial morphology that defines HEV.

### Endothelial Xbp1 Preserves HEV morphology, Supports Lymphocyte Recruitment, and Sustains Inflammatory Immune Responses

We next wished to determine whether the IRE1α–XBP1 pathway functions cell-intrinsically in endothelial cells to maintain HEV structure and function. We generated inducible, endothelial cell–specific *Xbp1* knockout mice (iEC-Xbp1*-/-)* by crossing *Cdh5-CreERT2* mice^93^ with *Xbp1*^fl/fl^ mice^94^. Cre-mediated recombination was induced by tamoxifen administration in adult mice. Tamoxifen-treated *Cdh5-CreERT2*–negative littermates served as wild-type controls. All groups were analyzed 4–10 weeks after tamoxifen treatment. Deletion of Xbp1 in ECs resulted in a significant reduction in LN size (Fig. 5A, left; Suppl. Fig. 9A). Whole-mount IF imaging revealed diminished PNAd expression (Fig. 5B), reduced HEV branching (Fig. 5A, right), and characteristic HEV flattening in iEC Xbp1-deficient mice (Fig. 5A, C; Suppl. Fig. 9B).

**Fig. 5.**
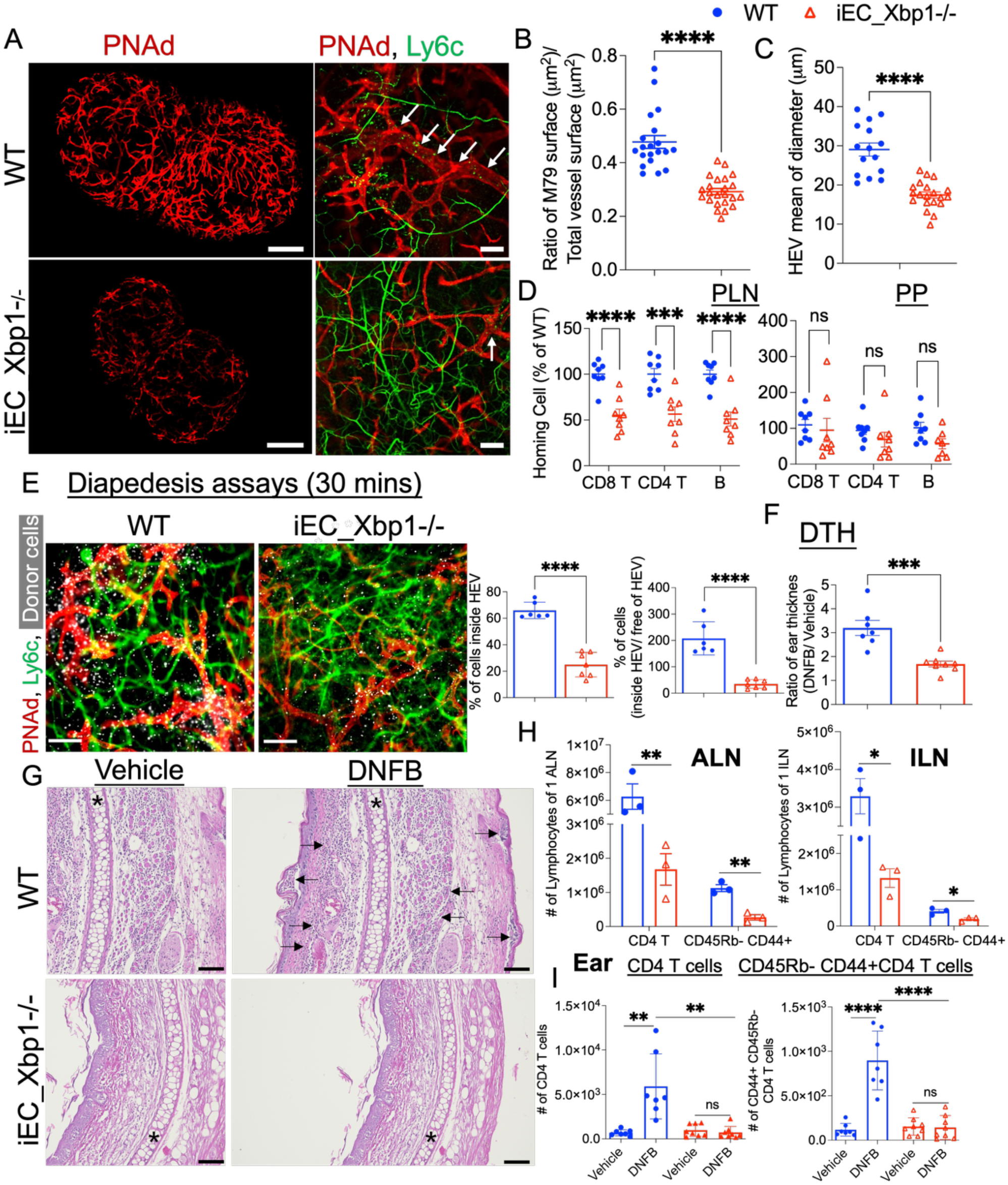
Endothelial Xbp1 Deletion Flattens HEVs and Reduces PNAd Expression, Lymphocyte Trafficking, and Inflammatory Immune Responses. **A.** Representative vascular IF images of PLN from WT control and iEC_Xbp1-/-mice. Mice were intravenously injected with antibodies labeling PNAd (MECA-79, red) and capillaries (Ly6C, green). Representative images from five independent experiments, each with one mouse per group per experiment. Arrows show thick HEV segments, arrow heads indicate elongated thin HEV. Scale bars: 100 μm. **B.** Reduced PNAd+ endothelium in EC-Xbp1 deficiency. PNAd⁺ HEV surface area performed relative to the total vascular surface area quantified using *Imaris*. **C.** Measurement of HEV diameter. Results are presented as the mean of 14-35 measurements per 10X field (∼4 mm2/field). (**B-C**): Data are shown as mean ± SEM from 2-3 fields per PLN, 1-2 PLN per mouse, n=5 per group. **D.** Donor lymphocyte localization in PLNs and Peyer’s patches (PPs) 2 hours post-transfer in control and iEC_Xbp1-/-mice. Results are shown as a percentage of the mean localization ratio to the WT group (mean ± SEM of with 8 mice per group). **E.** Left: Visualization of CFSE-labeled donor lymphocytes (white) within PLNs from control and iEC Xbp1-deficient mice, 30 minutes post transfer (i.v.). Vessels stained by i.v. injection of Mabs to PNAd (red) and capillaries (Ly6C, green). Scale bar: 100 μm. Right: Quantification of donor cell localization within PNAd⁺ HEVs and calculation of intra- vs. extravascular lymphocyte ratios using *Imaris*. Data are presented as mean ± SEM from 1–2 lymph nodes per mouse, n=4 mice per group. **F-I:** Attenuation of contact hypersensitivity responses in iEC_Xbp1-/-mice. **F.** Ear swelling ratio (DNFB/vehicle) measured 24 hours after challenge. n = 8 mice per group. **G.** H&E staining of ear tissue; stars indicate cartilage, arrowheads denote infiltrating leukocytes. Scale bar: 100 μm. **H-I.** Flow cytometry analysis of total and memory CD4⁺ T cells in auricular LN (ALN, H left), inguinal LN (ILN, H right) and inflamed ears (I). **H**: WT: n = 3; iEC_Xbp1-/-: n = 3. **I** WT: n = 7; iEC_Xbp1-/-: n = 8. All plotted data are shown as mean ± SEM. Statistical analysis was performed using two-tailed unpaired t-tests. *: p value < 0.05; **: p value < 0.01; ***: p value < 0.001; ****: p value < 0.0001.

Lymphocyte recruitment was severely impaired in iEC_Xbp1-/- mice. Total lymphocyte numbers in peripheral LNs were reduced by ∼65% compared to WT controls (Suppl. Fig. 9C), and short-term (2 hours) homing assays showed ∼50% reduction in lymphocyte accumulation in both PLNs and MLNs, while control organs were unaffected (Fig. 5D; Suppl. Fig. 9D). Donor-to-recipient lymphocyte ratios remained comparable between groups (Suppl. Fig. 9E), confirming that reduced recruitment was not due to altered donor cell viability. *In vivo* diapedesis assays further revealed a ∼60% decrease in lymphocyte adhesion to HEVs and an ∼85% reduction in extravascular to HEV-associated lymphocyte ratios in iEC Xbp1-deficient LNs (Fig. 5E), confirming compromised lymphocyte–HEV interaction and transmigration.

To assess whether EC-specific Xbp1 depletion impacts immune function, we evaluated contact hypersensitivity responses (DTH) in iEC_Xbp1-/- and WT littermate mice following sensitization and challenge with the hapten DNFB (2,4-dinitrofluorobenzene)^47^. Xbp1 deletion significantly attenuated ear swelling and leukocyte infiltration (Fig. 5F, G). CD4⁺ T cells, including memory subsets (CD4⁺CD44^highCD45RB^low), were markedly reduced in the draining lymph nodes (auricular and inguinal) and inflamed ear tissue of iEC_Xbp1-/- mice compared to controls (Fig. 5H–I). MECA-79⁺ HEV-like vessels were absent from DNFB-inflamed ear skin in both genotypes (data not shown). These findings support a critical role for endothelial Xbp1 in induction of inflammatory immune responses within draining lymph nodes, although we cannot exclude the possibility that Xbp1 also influences inflamed venular structures in extra-lymphoid tissues. Together, these results demonstrate that endothelial Xbp1 is required to maintain the specialized “high” endothelial phenotype, promote PNAd expression, support HEV integrity, and enable effective lymphocyte recruitment during both homeostatic and inflammatory immune responses.

### Xbp1 Cell-Intrinsically Coordinates Inter-Organelle Metabolic Programs, Organelle Integrity, and PNAd Biosynthesis in HECs

To define how Xbp1 sustains PNAd biosynthesis and organelle function *in vivo*, we performed single-cell RNA sequencing (scRNA-seq) of blood ECs isolated from lymph nodes of iEC Xbp1-deficient mice and littermate controls (Fig. 6A and B). UMAP analysis resolved nine EC populations consistent with our previously defined lymph node EC atlas^28^, including capillary ECs (capECs), venous ECs (Vn), HECs, and intermediate states (CRP, TrEC) along the capillary-to-HEC differentiation trajectory^28^ (Fig. 6A). As expected, *Xbp1* expression was reduced across all iEC-Xbp1-deficient EC subsets (Table S7). Compared with controls, fewer ECs were recovered from Xbp1-deficient mice, and the minor vnEC population was not represented (Fig. 6B).

**Fig. 6.**
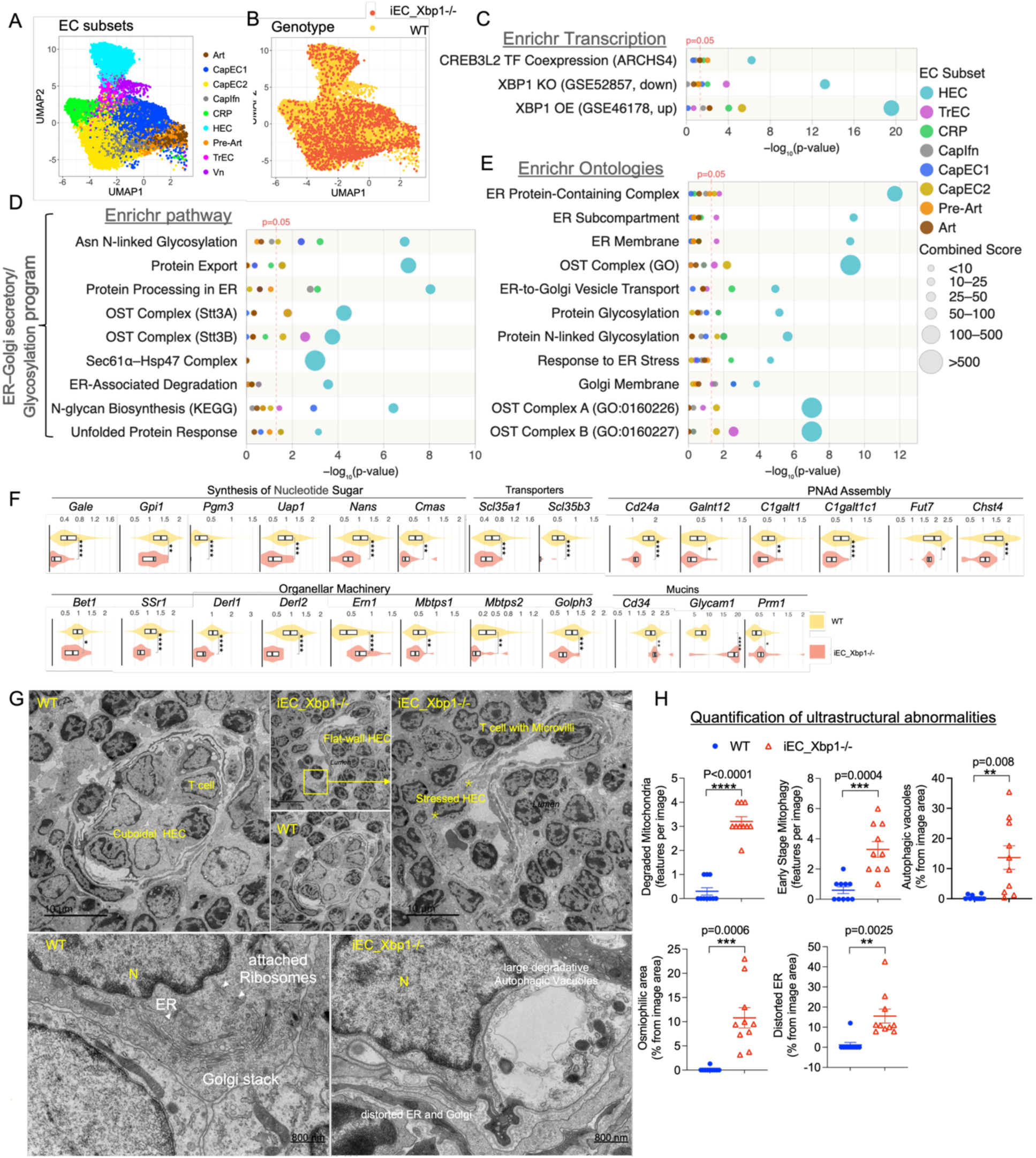
Endothelial Xbp1 Coordinates Metabolic Programs and Organelle Integrity to Sustain PNAd Biosynthesis in High Endothelial Cells. A–B. UMAP projections of lymph node blood endothelial cells colored by EC subset identity (A) and genotype (B; yellow, WT; red, iEC_Xbp1−/−), illustrating the transcriptional separation between conditions across all EC subsets. **C–E.** Integrated enrichment analyses of DEGs between LN blood ECs from WT littermates and iEC_Xbp1−/− mice (the top 2,000 DEGs per EC subset). Bubble plots display enriched terms from transcription factor–related libraries (C; ARCHS4, TF perturbations), pathway databases (E; BioPlanet, CORUM, KEGG, MSigDB Hallmark, WikiPathways), and Gene Ontology (GO) terms (D; Biological Process and Cellular Component). The x-axis represents term significance (−log₁₀ (P value)), with the dashed line marking the nominal significance threshold (P = 0.05). Bubble size reflects the enrichment combined score (integrating effect size and significance), and bubble color indicates EC subset identity. Redundant terms within each database were collapsed to the most significant representative. All panels share a unified x-axis and bubble size scale to allow direct comparison across datasets. Raw Enrichr results are provided in Table S8. **F.** scRNA-seq analysis of genes involved in PNAd biosynthesis and secretory organelle function in HECs from WT and iEC-Xbp1-/- PLNs. Violin plots show normalized expression values; box plots indicate median and interquartile range. n = 2 mice per group. Statistical testing was performed for exploratory purposes only due to limited sample size. *p < 0.05; **p < 0.01; ***p < 0.001; ****p < 0.0001. **G.** TEM of PLNs from WT and iEC-Xbp1-/- mice reveals pronounced ultrastructural abnormalities in iEC Xbp1-deficient HECs, including cellular flattening, distorted endoplasmic reticulum (ER) membranes with ribosome dissociation, and autophagic vacuoles containing mitochondria. N, nucleus; ER, endoplasmic reticulum. Scale bars: 2 μm, 10 μm, and 800 nm. **H.** Quantification of ultrastructural abnormalities in WT and iEC-Xbp1−/− HECs from TEM micrographs, including ER distortion, autophagic vacuoles, mitophagy-associated structures, degraded mitochondrial remnants, and osmiophilic cytoplasmic regions. Bars represent mean ± SEM (n = 10 micrographs per genotype), with each dot corresponding to an individual micrograph. For each genotype, 3 mice were analyzed; ≥2 tissue blocks per mouse were prepared; and ≥2 grids per block (10–12 sections per grid) were randomly sampled. Images were acquired from random fields on each section. Statistical comparisons were performed using unpaired two-tailed t-tests with Welch’s correction.

To assess transcriptional impact of Xbp1 deletion, differentially expressed genes (DEGs) were identified within each EC subset and analyzed using Enrichr (Table S7-9). HECs exhibited the most pronounced transcriptional response to Xbp1 loss (Suppl. Fig. 10A). HEC-DEGs (downregulated in iEC_Xbp1−/− HECs, defining endothelial Xbp1-induced or dependent genes in HECs), were associated with XBP1-regulated gene sets and CREB3L2-linked signatures (Fig. 6C) and showed pronounced disruption of glycosylation and ER–Golgi secretory pathways, including processes related to IRE1α chaperone activity, protein processing in the ER, N- and O-linked glycosylation, oligosaccharyltransferase (OST) complex function, N-glycan trimming, the calnexin/calreticulin cycle, protein export, ER-associated degradation, and Golgi organization (Fig. 6D and E, Suppl. Fig. 11 A). Consistent with these pathway-level changes, many genes essential for PNAd biosynthesis and secretory organelle function were downregulated, including genes spanning nucleotide sugar production (*Gale, Gpi1, Pgm3, Uap1, Nans, Cmas*), metabolite transport (*Slc35a1, Slc35b3*), and glycan assembly enzymes (*Galnt12, C1galt1, C1galt1c1, Fut7, Chst4*). Key components of the secretory machinery were similarly downregulated (*Ssr1*, *Bet1*, *Derl1*, *Derl2*, *Mbtps1*, *Mbtps2* and *Golph3)*. (Fig. 6F). In contrast, expression of genes encoding core PNAd scaffolding mucins^7^ was not decreased. Notably, Ern1 (IRE1α) expression was upregulated in iEC Xbp1-deficient HECs (Fig. 6F), a response previously associated with pro-apoptotic signaling and cell stress in other Xbp1 knockout models^95^.

PNAd synthesis-associated pathways were generally much less affected by Xbp1 loss in other EC subsets, but Xbp1 deficiency impacted their gene expression as well (Suppl. Figs. 12-18, Panels A; Table S8). DEG Genes downregulated by Xbp1 loss were enriched for immune and inflammatory programs in CapIfn, arterial, and TrEC populations; lysosomal and autophagy-associated processes in TrECs, and mitochondrial and energy homeostasis across capillary, pre-arterial, arterial, and CRP populations. Genes upregulated in KO were enriched in adaptive responses across EC populations, spanning stress, metabolic, and inflammatory associated programs (Suppl. Figs. 11–18, Panels B; Table S9). Within this broader endothelial context, HECs preferentially upregulated translational and ER stress–related pathways (Suppl. Fig.11B) and CapIfn cells exhibited enhanced interferon signaling (Suppl. Fig.18B). Notably, canonical EC subset identity marker genes were largely retained across all populations in iEC-Xbp1-deficient mice (Suppl. Fig. 10B) — including pan-EC markers (*Cdh5*, *Pecam1, Erg, Ets1*), arterial and capillary markers (*Bmx, Hey1, Dll4, Notch1, Notch4*), venous/HEC markers (*Nr2f2, Selp, Ackr1*), and CRP markers (*Cxcr4, Apln, Pgf, Arhgap18*) — indicating that endothelial lineage identity is preserved despite transcriptional perturbation, and that the observed gene expression changes reflect stress and adaptive responses rather than a general collapse of EC identity.

These findings indicate that, while Xbp1 broadly supports endothelial homeostasis, HECs depend on Xbp1-driven metabolic and secretory programs to sustain their functional specialization.

We next asked if the transcriptional changes in HEC translated into ultrastructural changes in organelle architecture and cellular integrity. Xbp1-deficient HECs exhibited pronounced morphological abnormalities compared to WT controls, including cellular flattening, distorted ER membranes with ribosome dissociation, and prominent autophagic vacuoles containing mitochondria (Fig. 6G and Suppl. 19A). Quantitative analysis confirmed significant increases in multiple ultrastructural stress features in mutant HECs (Fig. 6H), including ER distortion, autophagic vacuoles, mitophagy-associated structures, mitochondrial remnants, and osmiophilic cytoplasmic regions. Adherent platelets were observed (Suppl. Fig. 19B). These morphological abnormalities indicate severe organelle stress and compromised biosynthetic capacity, consistent with the transcriptional changes identified by scRNA-seq.

Together, these results demonstrate that endothelial Xbp1 is essential for coordinating inter-organelle metabolic and biosynthetic programs, preserving secretory organelle architecture, and sustaining PNAd biosynthesis and HEC survival.

### XBP1 Is Required for HEV Expansion, Ectopic PNAd⁺ Vessel Induction, and Functional Lymphocyte Homing during Inflammation

During immune activation, HEVs undergo dynamic expansion through immune-driven angiogenesis, facilitating enhanced lymphocyte recruitment and enlargement of draining LNs^96,97^. To determine whether Xbp1 activation is required for HEV expansion during immune stimulation, we immunized mice locally using topical oxazolone treatment. In contrast to the expanded HEVs observed in immunized wild-type lymph nodes (Fig. 7A-B and Suppl. Fig. 20A), iEC-specific deletion of *Xbp1* or pharmacological inhibition with STF083010 resulted in markedly thinner and sparsely branched PNAd⁺ vessels with substantially reduced PNAd intensity. Consistent with these changes, endogenous lymphocyte abundance and lymphocyte homing capacity were profoundly diminished (Suppl. Fig. 20B-C). These results demonstrate a key role for Xbp1 in HEV expansion and functional maturation during immune-driven angiogenesis. We next examined the role of Xbp1 in the atypical lymphoid structures of omental milky spots (MSs), immune aggregates within the peritoneal cavity that rely on induced PNAd⁺ vessels for lymphocyte recruitment and tissue organization^18,98^. Previous studies show that MS-associated HEVs are rapidly induced by microbial stimuli such as lipopolysaccharide (LPS). Intraperitoneal injection of LPS (two injections over 72 hours) triggered robust and rapid induction of PNAd⁺ vessels in MSs, and this response was significantly attenuated in iEC Xbp1-deficient mice (Fig. 7C–D).

**Fig. 7.**
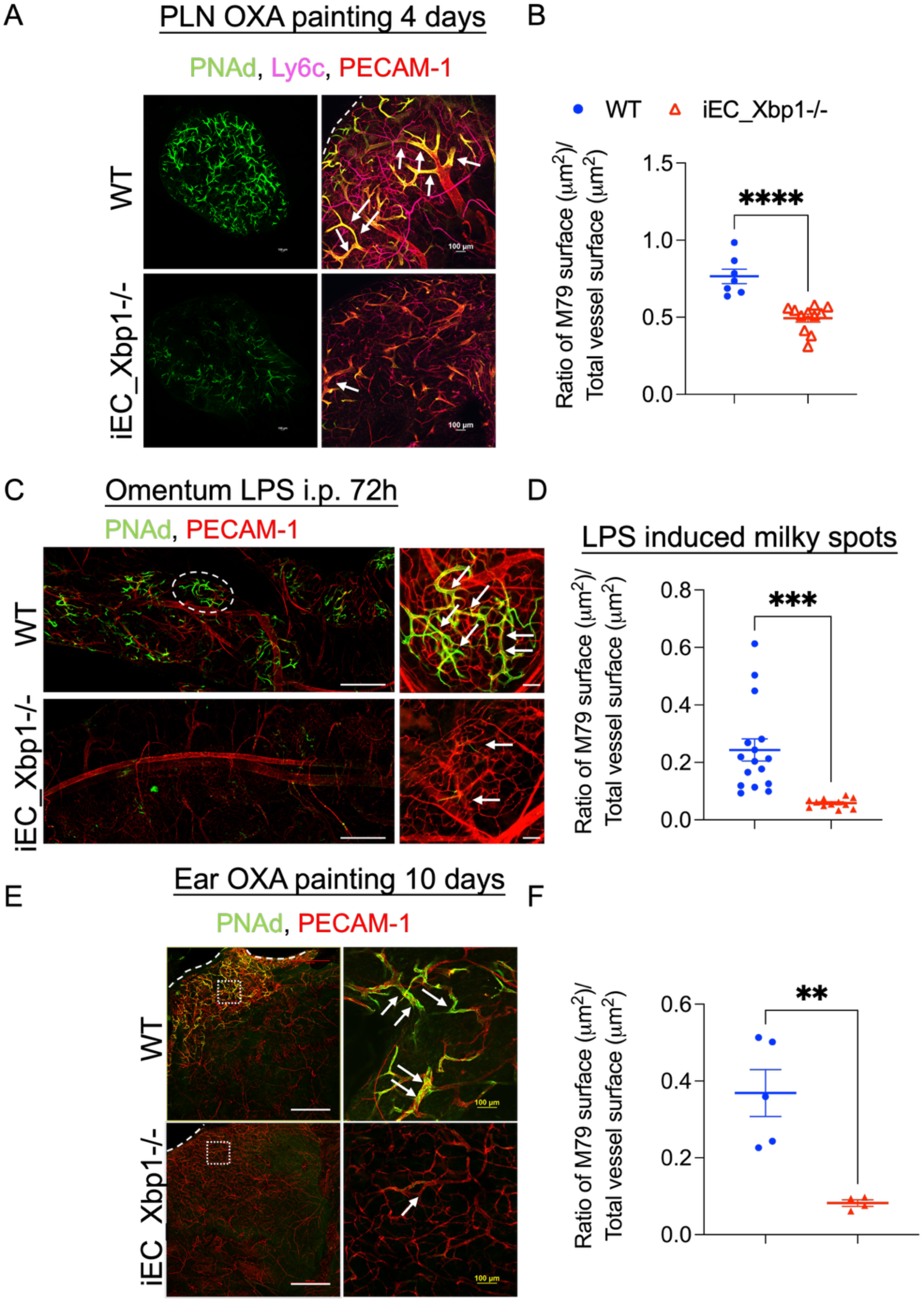
Endothelial Xbp1 is required for HEV expansion and Ectopic PNAd⁺ Vessel Induction during Inflammation. **A.** Representative whole-mount IF images of PLNs from WT and iEC_Xbp1-/- mice, four days after OXA-induced cutaneous inflammation. Intravenous labeling with antibodies against PNAd (MECA-79, green), capillary EC (Ly6C, pink), and CD31 (red) highlights HEV morphology and surface markers. Dashed lines indicate LN boundaries; arrows denote PNAd extension along vessel segments. Scale bars, 100 μm. **B.** Quantification of PNAd⁺ HEV surface area relative to total vessel area, measured using *Imaris* analysis. Data represent mean ± SEM from 2 fields per lymph node; 1–2 LNs per mouse (WT: n = 3; iEC_Xbp1-/-: n = 3). **C.** Left: Whole-mount omental staining of WT and iEC_Xbp1-/- mice following two intraperitoneal LPS injections over 72 hours. Right: Representative milky spots showing induced PNAd⁺ vessels (green) within CD31⁺ vasculature (red). Arrows highlight induction of PNAd+ HEVs. Scale bars, 100 μm. **D.** Quantification of the PNAd⁺/total vessel area ratio within milky spots, based on 10× fields (∼4 mm²/field), analyzed using *Imaris*. Data represent mean ± SEM from 3-4 fields per mouse: n = 4 mice per group. **E.** Whole-mount IF of ear vasculature ten days after bilateral OXA application. Left: overview of entire ear; right: higher-magnification views of inflamed skin showing PNAd⁺ vessels (green) and CD31⁺ endothelium (red). Dashed lines mark ear boundaries; arrows indicate PNAd⁺ HEV-like vessels. Scale bars, 100 μm. **F.** Quantification of PNAd⁺ vessel area in inflamed ear skin, analyzed using Imaris. Data are presented with mean ± SEM from 1 field from each ear of WT (n = 5) or iEC_Xbp1-/- mice (n = 3). B, D and F: Statistical analysis performed using unpaired two-tailed t-tests: *: p value < 0.05; **: p value < 0.01; ***: p value < 0.001; ****: p value < 0.0001.

HEV-like vessels also emerge *de novo* in sites of chronic inflammation^18,99^ where they are implicated in various human inflammatory diseases and cancer^100–103^. To model induction of PNAd in immunologically driven inflammation, we assessed HEV-like vessel induction in an oxazolone-induced model of atopic dermatitis^104^. We applied oxazolone to the ears of tamoxifen-treated iEC_Xbp1-/-and WT mice and analyzed vessels ten days later. WT mice developed prominent PNAd⁺ HEV like vessels in inflamed ear skin, whereas iEC Xbp1-deficient mice showed markedly reduced induction of these vessels (Fig. 7E–F).

Together, these findings establish Xbp1 as a critical regulator of PNAd⁺ HEV expansion in LN and and ectopic HEV formation in atypical lymphoid tissues and peripheral inflamed organs. By enabling the molecular specialization of HEV, Xbp1 promotes efficient lymphocyte recruitment and supports local tissue immune responses during inflammatory stimulation.

### S1P Protease Inhibition Prevents Ectopic PNAd⁺ Vessel Formation, HEV Immune Angiogenesis, and Functional Lymphocyte Homing during Inflammation

We next assessed the effects of the S1P protease inhibitor PF429242 in models of the MS-induced HEV neogenesis and lymph node immune angiogenesis. S1P inhibition markedly reduced the induction of PNAd⁺ HEVs in both MSs (Suppl. Fig. 21) and PLNs (Suppl. Fig. 22), recapitulating key features of Xbp1 deficiency. The inhibitor reduced generation of the canonical MECA79 glycotope (6-sulfo N-acetyllactosamine in O-linked branching structures), but even more dramatically suppressed reactivity with antibody S2 recognizing alpha 2,3neuraminic acid modified 6-sulfo LacNAc (6SO4 sialyl LacNAc), structures present on both O- and N-linked glycans. Consistent with a broader disruption of HEV glycan elaboration, staining with antibody F2, which recognizes sialyl Lewis X-containing N- and O-glycans, was also reduced in homeostatic lymph nodes following PF429242 treatment (Suppl. Fig. 23). These suggest that S1P-dependent transcription coordinates multiple glycosylation steps and is essential for ectopic HEV formation^90,105^.

To assess functional consequences, we performed homing assays in oxazolone-painted mice treated with PF-429242. Both endogenous lymphocyte accumulation and recruitment of intravenously transferred donor cells were significantly reduced in PF-treated mice (Suppl. Fig. 24), suggesting that SP1 - dependent HEV induction is essential for functional lymphocyte entry into inflamed tissues. Although S1P protease regulates multiple transcription factors — including CREB3 family members, ATF6, and SREBPs — and effects on other cells and pathways cannot be excluded, these in vivo findings are consistent with a role for S1P-dependent transcriptional programs in controlling HEV biosynthetic and functional gene networks. The specific contribution of CREB3L2 among these substrates will require future endothelial-specific genetic studies.

## Discussion

Continuous lymphocyte recruitment into lymph nodes through HEVs puts exceptional biosynthetic demands on the vasculature, and how these demands are met at the transcriptional level has remained poorly understood. Here we show that IRE1α–XBP1–centered transcriptional networks, together with CREB3L2-associated gene programs, orchestrate the metabolic and organellar infrastructure required for PNAd biosynthesis and lymphocyte homing. These networks scale nucleotide sugar and sulfate donor synthesis, direct precursor transport across endomembrane compartments, and deploy glycosyltransferases and sulfotransferases within the Golgi — thereby coupling metabolic flux to ER and Golgi expansion and enabling the high-output glycoprotein production that underlies HEV identity.

The mechanistic centerpiece of this regulatory logic is the coupling of nucleotide sugar and PAPS biosynthesis to the spatial organization of the Golgi glycosylation machinery. Our data imply that XBP1 and CREB3L2 engage conserved cis-regulatory modules across a coordinated suite of genes spanning nutrient uptake, precursor metabolism, endomembrane transport, and glycan assembly. The frequent co-occurrence of XBP1 and CREB3L2 motifs within evolutionarily conserved regions suggests that these transcription factors may act cooperatively as well as independently, a model supported by reporter assays demonstrating both combinatorial and factor-specific regulation of PNAd biosynthetic gene promoters and enhancers. The concomitant upregulation of ER and Golgi biogenesis and function genes in HECs, and their disruption upon Xbp1 deletion or pharmacologic IRE1α or S1P inhibition, suggest that these transcriptional programs drive the scaling of secretory organelle capacity required to process the resulting glycoprotein load. Consistent with this framework, genes involved in precursor synthesis, glycosylation, and ER–Golgi trafficking are broadly represented among established direct transcriptional targets of XBP1 and CREB3L family factors^66,67,106^. This systems-level coupling is consistent with a mechanism by which the characteristic “high” morphology of HEVs and their glycosylation output are maintained as a coherent, integrated cell state rather than as independent features, though whether transcriptional coordination of these programs is sufficient to establish HEC morphology independently of other signals remains to be determined. Activation of XBP1 through IRE1α-mediated mRNA splicing and of CREB3L2 through S1P/S2P-mediated proteolysis enables these pathways to dynamically respond to increased secretory demand.

*In vivo* perturbation experiments at steady state further support this regulatory framework. Pharmacologic inhibition of IRE1α RNase activity and endothelial-specific deletion of Xbp1 demonstrate that the IRE1α–XBP1 axis functions as a principal regulator coordinating metabolic flux, PNAd biosynthesis, and the characteristic “high” morphology of HECs. Disruption of this pathway reduces expression of key PNAd biosynthetic genes, alters secretory organelle architecture, diminishes PNAd expression, flattens HEC, and impairs lymphocyte homing and diapedesis. Consistent with this model, pharmacologic inhibition of the site-1 protease using PF-429242 — which prevents activation of CREB3L2 — also collapses HEC and reduces PNAd expression and lymphocyte recruitment. Because S1P inhibition also blocks other S1P/S2P-dependent substrates, including ATF6, further studies will be required to define the specific contribution of CREB3L2. Nevertheless, CREB3L2 or its closely related homolog CREB3L1 has been associated with transcriptional programs engaged during increased secretory and glycosylation demand^50,52,66,107,108^, and notably undergoes N-glycosylation itself, raising the possibility that its maturation and activation are coupled to the metabolic and structural requirements of glycan synthesis^109^. Through this dual control, these pathways can integrate ER adaptive capacity with metabolic and lineage-specific transcriptional programs, reinforcing glycosylation output and the scaling of secretory organelles that support PNAd production.

Beyond steady-state biosynthesis for HEC maintenance, these regulatory circuits also enable the adaptive remodeling of secretory capacity required for HEV expansion or generation in response to immune challenge. Pharmacologic inhibition of IRE1α or EC-specific deletion of Xbp1 impaired HEC remodeling and lymphocyte recruitment during oxazolone-induced inflammation. EC-specific deletion of Xbp1 also reduced ectopic induction of PNAd⁺ HECs in inflamed omental lymphoid tissues, a phenotype similarly observed following treatment with the S1P inhibitor PF-429242. These findings indicate that both the IRE1α–XBP1 and S1P-dependent transcriptional pathways are required not only for maintenance of steady-state HEC identity but also for inflammatory induction of specialized lymphocyte-recruiting endothelium. A parallel regulatory framework appears to operate in goblet cells, where XBP1 and CREB3L1 support sulfated mucin biosynthesis and expansion of secretory organelles, thereby sustaining mucosal barrier function under basal conditions and during microbial challenge^25,50,77^ ^26,62^. Together, these findings suggest that coupling metabolic input to organelle architecture through shared transcriptional networks is a conserved strategy for sustaining secretory output across distinct tissue contexts and challenges.

More broadly, our findings in HECs support a generalizable regulatory principle in which IRE1–XBP1-centered and CREB3 family networks coordinate metabolic and organellar programs to enable glycosylation-dependent specialization across diverse secretory cell types. Consistent with this framework, goblet cells and stomach-resident plasma cells engage parallel biosynthetic programs to support high-level production of sulfated mucins and specialized N-glycosylation of IgG, respectively (Suppl. Fig. 2 and 3), while pituitary glandular cells similarly enrich precursor-synthesis pathways and lineage-restricted glycosyltransferases required for glycosylated peptide hormone production^110,111^ (Table S6). XBP1 is established as a key regulator in plasma^53^ and pituitary secretory cells^52^, and recent studies further implicate CREB3L2 in both lineages^52,112^, suggesting a shared transcriptional architecture coupling secretory demand to lineage-specific glycan output. This regulatory logic may extend to additional specialized secretory cells, including chondrocytes and osteoblasts^50,113–116^, further supporting its generality across diverse biosynthetic contexts.

This regulatory architecture also appears to be evolutionarily conserved. A homologous mechanism may operate in *Drosophila*, where CrebA — the sole CREB3 family member — and Xbp1 together regulate secretory machinery in the salivary gland, suggesting deep evolutionary conservation of this regulatory architecture^66,117,118^. This conserved network may represent a core solution for scaling biosynthetic output of glycosylated macromolecules under both homeostatic and stress-adaptive conditions. Further investigation into how these transcriptional circuits interface with lineage-specific and signal-induced programs may yield insights into the modulation of secretory capacity in health and disease. Additionally, these pathways may provide strategic entry points for therapeutic glycoengineering or for manipulating vascular and epithelial remodeling in inflammation-associated conditions.

## Limitations of the study

Our study implicates IRE1α–XBP1–centered transcriptional networks in linking PNAd biosynthesis to metabolic reprogramming and secretory organelle expansion in high endothelial cells. Although downregulation of genes required for 6-sulfo sialyl Lewis X biosynthesis in iEC-Xbp1–deficient HECs correlates with loss of PNAd glycotopes and “HEVness,” our study does not exclude additional effects of Xbp1 loss on mucin precursor processing, the spatial organization of glycosyltransferases, or trafficking through the ER–Golgi system, which could also contribute to reduced PNAd expression.

Despite the profound impact of endothelial Xbp1 deficiency on HEC biosynthetic capacity, morphology, and lymphocyte recruitment, gene expression changes were also observed in other endothelial subsets, including subsets with HEC precursor potential. Thus, contributions from these subsets cannot be excluded. Definitive resolution of HEC-intrinsic requirements will require HEC-restricted genetic models such as CHST4-CreERT2^119^. Similarly, we emphasize that the in vivo functional requirement for CREB3L2 in HEV biology has not been formally demonstrated in this study. A specific contribution of CREB3L2 is suggested by transcription factor binding site motif enrichment in HEC-accessible chromatin and by transcriptional activation of PNAd biosynthetic gene reporters in vitro; however, confirmation of its in vivo role will require future endothelial-specific genetic studies.

In addition, how these transcriptional programs intersect with NF-κB signaling and are modulated by cytokine cues, metabolic inputs, and organelle remodeling across distinct immunological contexts remains incompletely understood. Addressing these questions will further refine our understanding of how transcriptional coordination of metabolic and organellar programs shapes immune cell trafficking and vascular specialization.

## Supporting information

https://drive.google.com/drive/folders/16LrJoVh-ugeEoXfNzdhAxGWC4Km_MaxT?usp=sharing

## Resource availability

### Lead contact

For more information and resource requests, please contact Yuhan Bi, who will collaborate with the Butcher Lab as needed.

### Materials availability

Reagent resources are available upon request to the lead contact.

### Code Availability

scRNA-seq analysis was performed using publicly available software. Code for computing imputed expression values is available: https://github.com/kbrulois/magicBatch.

### Data Availability

The authors declare that all data supporting the findings of this study are available within the article and its supplementary information files. Raw and processed scRNA-seq data generated in this study are available from the NCBI Gene Expression Omnibus repository under the accession number GSE296739. Source data are provided with this paper. De novo motif discovery on HEC-enriched genes is provided as a custom track in a UCSC Genome Browser session: https://genome.ucsc.edu/s/m.xiang/HEV_genes_conserved. Raw sequencing reads (FASTQ files) for ATAC-seq performed on PLN and PP HECs, as well as other endothelial cell subsets, are being deposited.

## Acknowledgments

Supported by NIH grants R01 AI130471 and R01 AI047822, grant 1903-03787 from The Leona M. & Harry B. Helmsley Charitable Trust to E.C.B. The Transmitted Electron Microcopy (TEM) study was supported, in apart, by NIH S10 Award Number 1S10OD028536-01, titled “OneView 4kX4k sCMOS camera for transmission electron microscopy applications” from the Office of Research Infrastructure Programs (ORIP). Y.B. and B.O. were supported by a Research Fellow Award of the Crohn’s and Colitis Foundation of America (835171 and 574148). M.X. was supported by the Tobacco-Related Disease Research Program of University of California (T31FT1867). We thank John J. Perrino for TEM sample preparation and imaging. We thank L. Magalhaes for administrative support.

## Author contributions

Y.B. conducted and oversaw experiments; K.B and Y.B. performed single-cell RNA-seq analysis. M.X., Y.B., J.P. performed genomic analyses. Y.B. A.A., R.B., B.O., M.K., T.D. and N.L. performed experiments. G.R. helped with mouse colonies. F.L. advised TEM imaging and analysis. H.K. provided mAbs. J.H.L. provided Xbp1tm2Glm fl/fl mice and scientific advice. Y.B., J.P. and E.C.B. conceived and supervised the study. Y.B., J.P. and E.C.B. wrote the manuscript.

## Supplemental information

Document S1. Supplemental Figs. 1–24.

Data S1. Immunohistochemical profiling from the Human Protein Atlas validated these transcriptomic predictions at the protein level, revealing selective enrichment in HECs of enzymes supporting metabolic flux, nucleotide sugar transporters involved in endomembrane trafficking, core components of the PNAd biosynthetic machinery regulators of secretory organelle homeostasis and vesicular transport, and unfolded protein response factors, related to Fig. 1

Data S2. Base-resolution phylogenetic DNA footprinting of genes regulating metabolic flux for sulfated mucin-type O-glycans and organelle machinery in HECs and GCs, related to Fig. 2D and Discussion.

Data S3. Phylogenetic DNA footprinting at base resolution reveals regulatory elements associated with specialized glycosylation in plasma cells and pituitary cells, related to Fig. 2D and Discussion.

Table S1. Multiple excel sheets of 2000 upregulated DEGs of each subset of lymph node endothelium and colon epithelium, related to Fig. 1 & 2.

Table S2-4. Integrated Enrichr TF enrichment analyses of HEC- and GC-enriched genes, related to Fig. 2.

Table S5. Integrated HEC-enriched ATAC-seq peaks (compared with capECs), related to Fig. 2.

Table S6. Single-cell transcriptomic data of Stomach Resident Plasma Cells and Pituitary Glandular Cells from Human Cell Atlas, related to Suppl. Fig. 4 and Discussion.

Table S7. Differential gene expression (DEG) analyses between wild-type (WT) and Xbp1-deficient ECs across all EC subsets, related to Fig. 6 and Suppl. Figs. 10-18.

Table S8-10. Enrichr analyses performed for each EC population using DEGs from WT and Xbp1-deficient ECs, related to Fig. 6 and Suppl. Figs. 10-18.

Table S11. Information for constructs, antibodies (Western Blot) and RT-PCR Primers.

## Supplemental Figures

**Suppl. Fig. 1.**
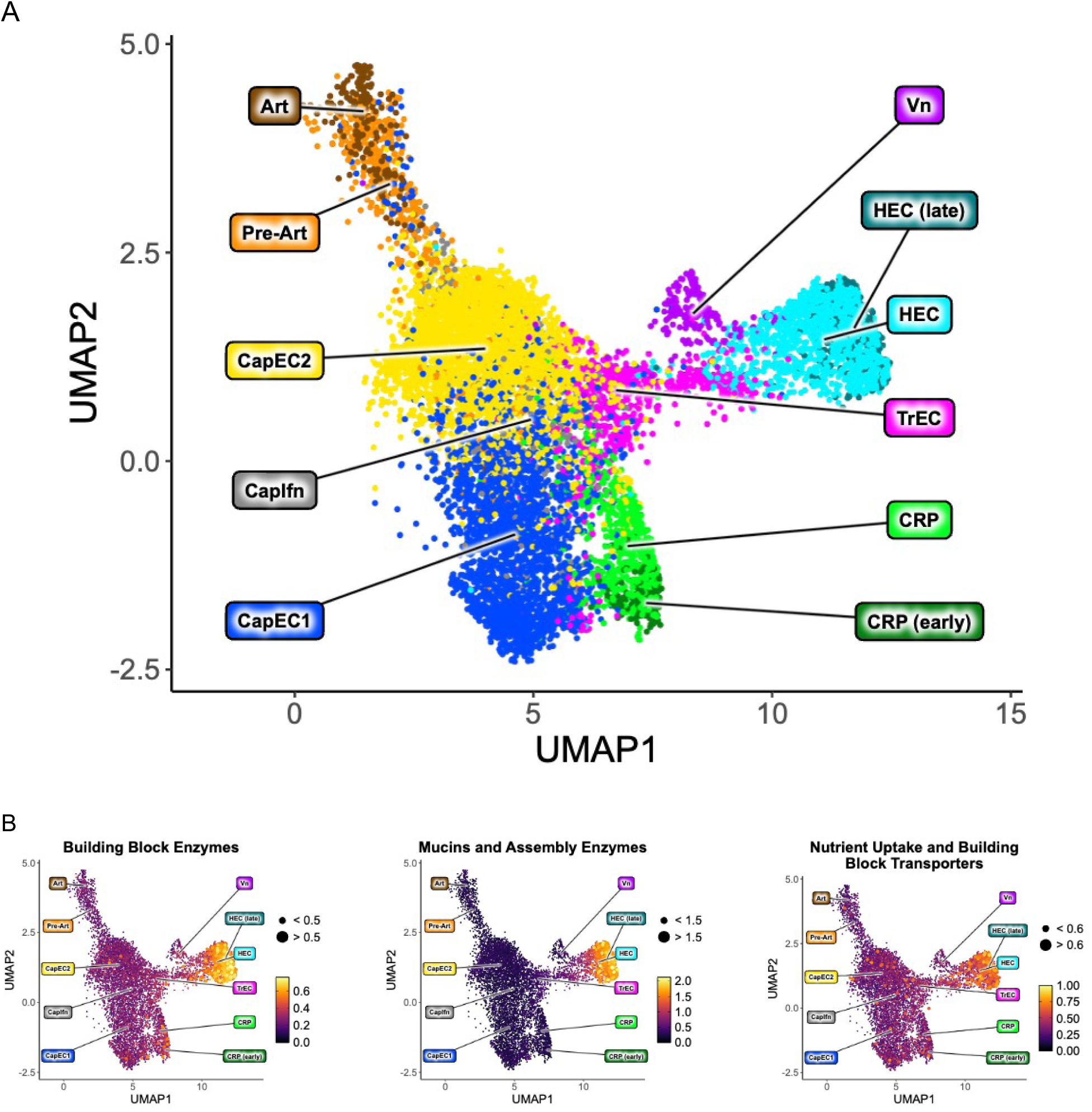
(linked to Figure 1)**. HECs Exhibit Enriched Expression of Metabolic and PNAd Biosynthetic Gene Programs. A.** UMAP projection of EC subsets from scRNA-seq profiling of murine peripheral lymph node ECs^26^. Cells are segregated by 11 subsets: arterial EC (pre-Art), high endothelial cells (HEC), non-HEC veins (Vn), and five capillary phenotype EC: CapEC1, CapEC2, capillary resident progenitors (CRP), transitional EC (TrEC), Interferon-stimulated gene-enriched CapEC (CapIfn). **B**. UMAP plots display module scores summarizing the expression of genes controlling building-block enzymes (left), mucins and assembly enzymes (middle), and nutrient-uptake/building-block transporters (right), all of which are significantly enriched in HECs compared with other EC subsets.

**Suppl. Fig. 2.**
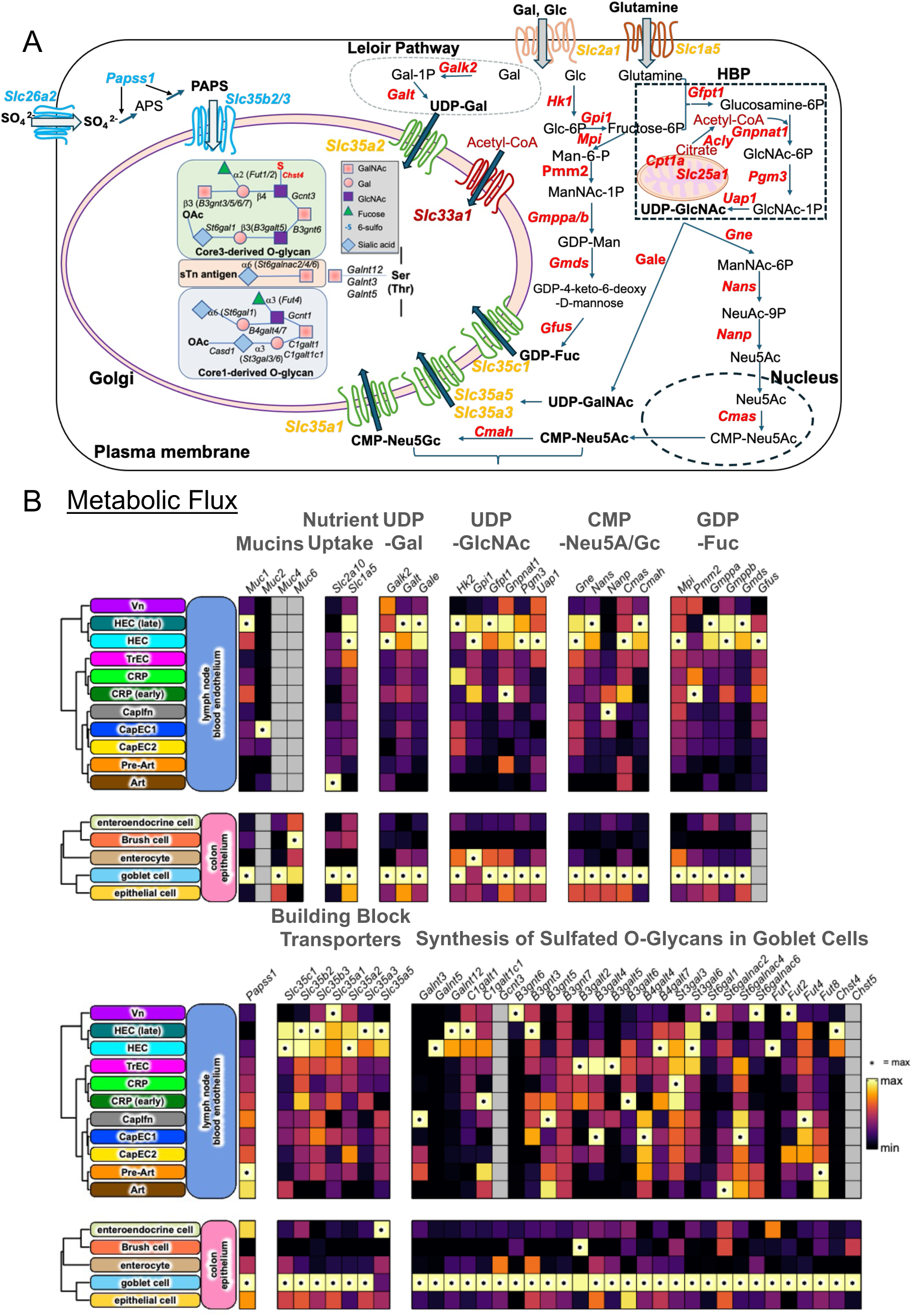
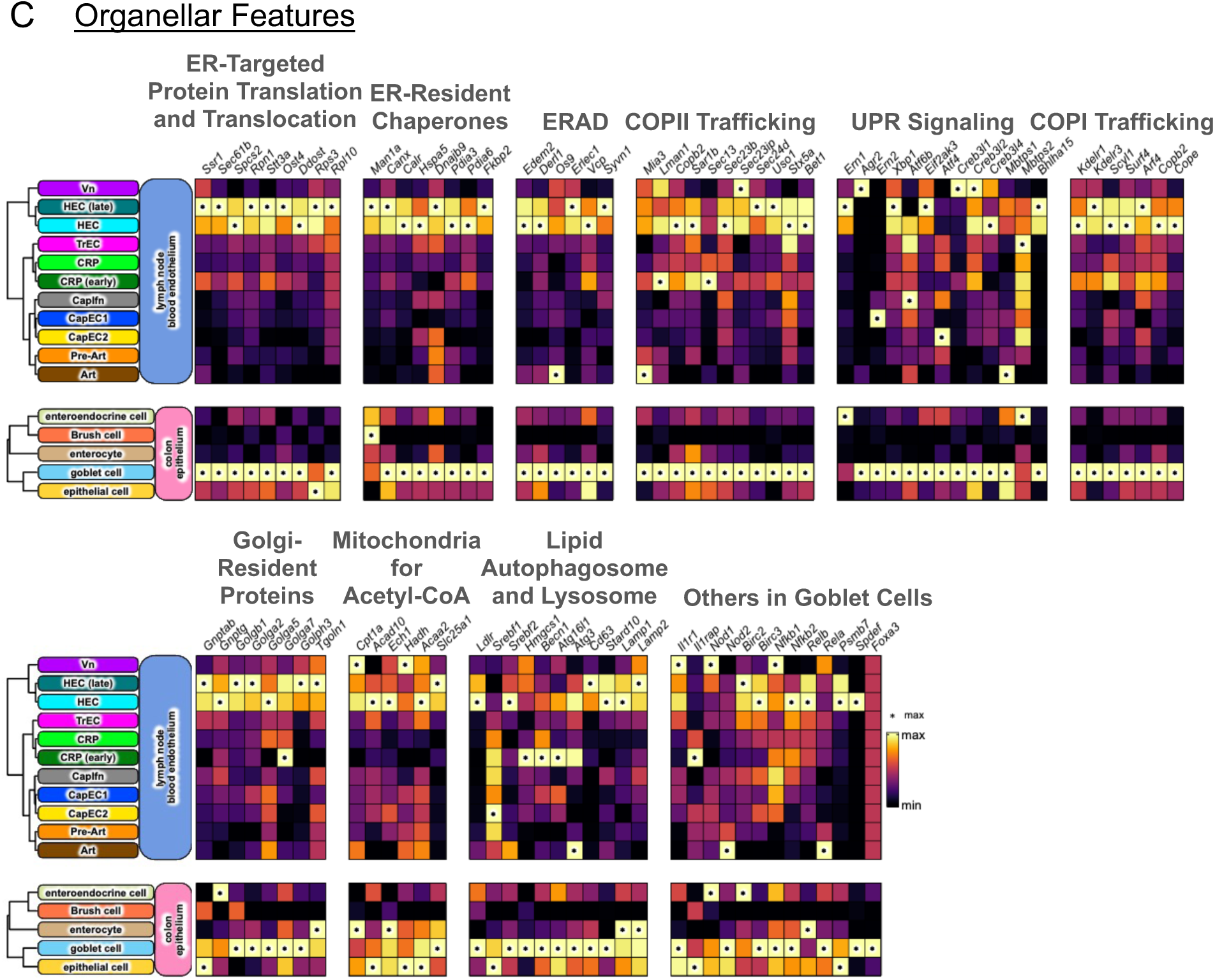
(linked to Fig. 1). **Goblet Cells Share Metabolic and Secretory Gene Programs with HECs for Sulfated Mucin Production. A.** A schematic chart illustrates the progression of metabolic fluxes, beginning with nutrient uptake and the production of building blocks, followed by the endomembrane transport of building blocks, and culminating in the biosynthetic assembly of sulfated O-glycans within the Golgi apparatus in GCs. The fluxes are regulated by specific transporters and enzymes encoded by the genes highlighted in yellow and red, respectively, with those specifically involved in sulfation shown in blue. **B–C.** Heat maps display GC-enriched expression (bottom panels) of genes involved in metabolic pathways for sulfated mucin biosynthesis (B) and organelle-associated protein machinery (C). The expression profiles of the same genes in HECs are shown in the top panels. Mean expression values in panels B and C are derived from three independent samples and are shown on a color scale from min to max, with an asterisk (*) indicating the maximum value.

**Suppl. Fig 3.**
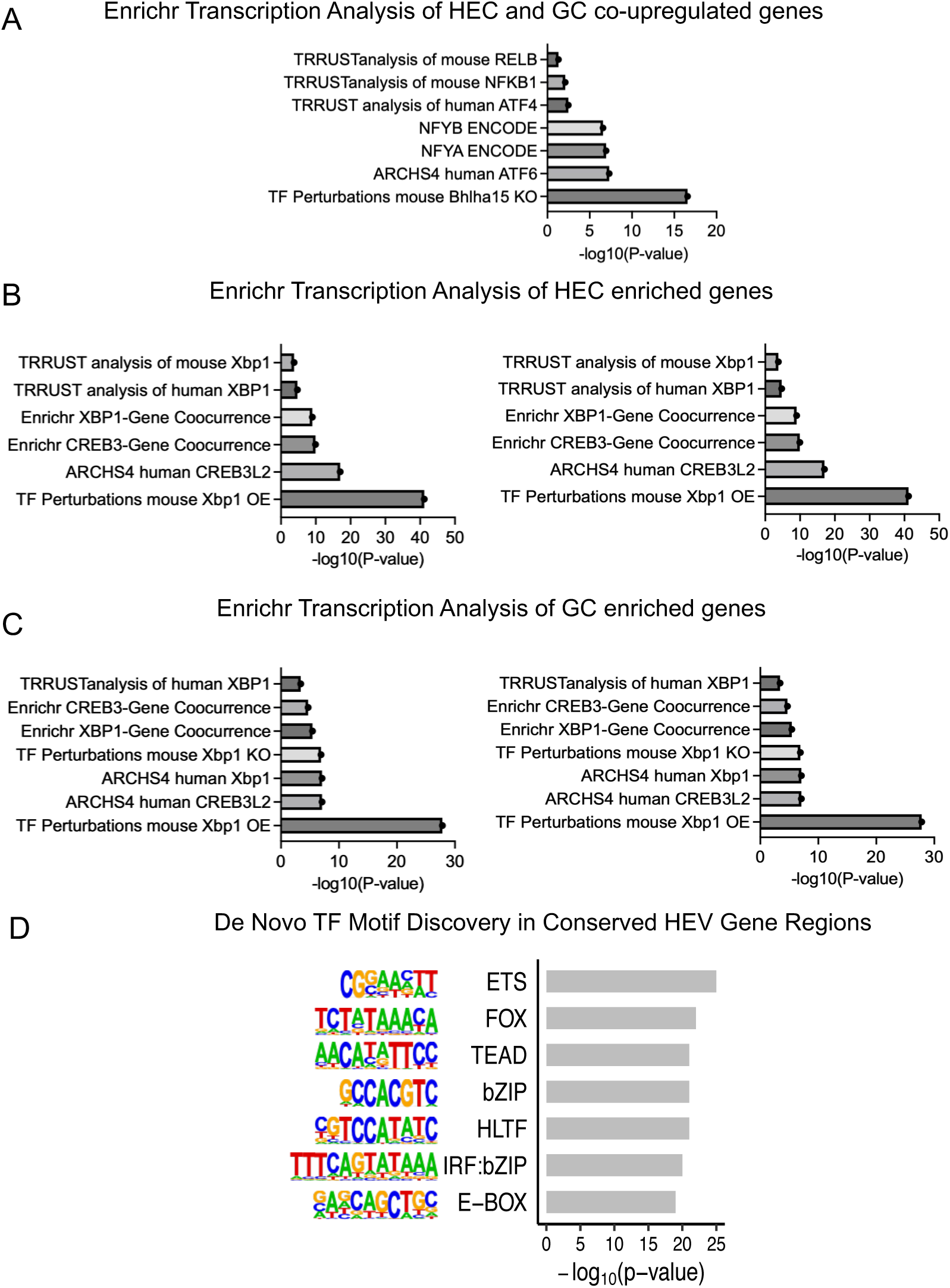
(Linked to Figure 2). **Motif and Enrichment Analyses Identify XBP1 and CREB3L2 as Candidate Regulators of HEC and Goblet Cell Programs. A**. Enrichr TF analysis of the up-regulated DEGs common to HEC and GC (Table S2). **B.** Enrichr TF analysis of HEV enriched genes (Table S3). **C.** Enrichr TF analysis of intestinal GC enriched genes (Table S4). **D.** De novo motif discovery in conserved regulatory regions flanking HEC-enriched genes identified bZIP family binding sites, including the consensus sequence for XBP1 and CREB3L2. Motif analysis was performed using HOMER on the mouse mm10 genome, with conserved regions defined by phastCons placental mammal scores.

**Suppl. Fig 4.**
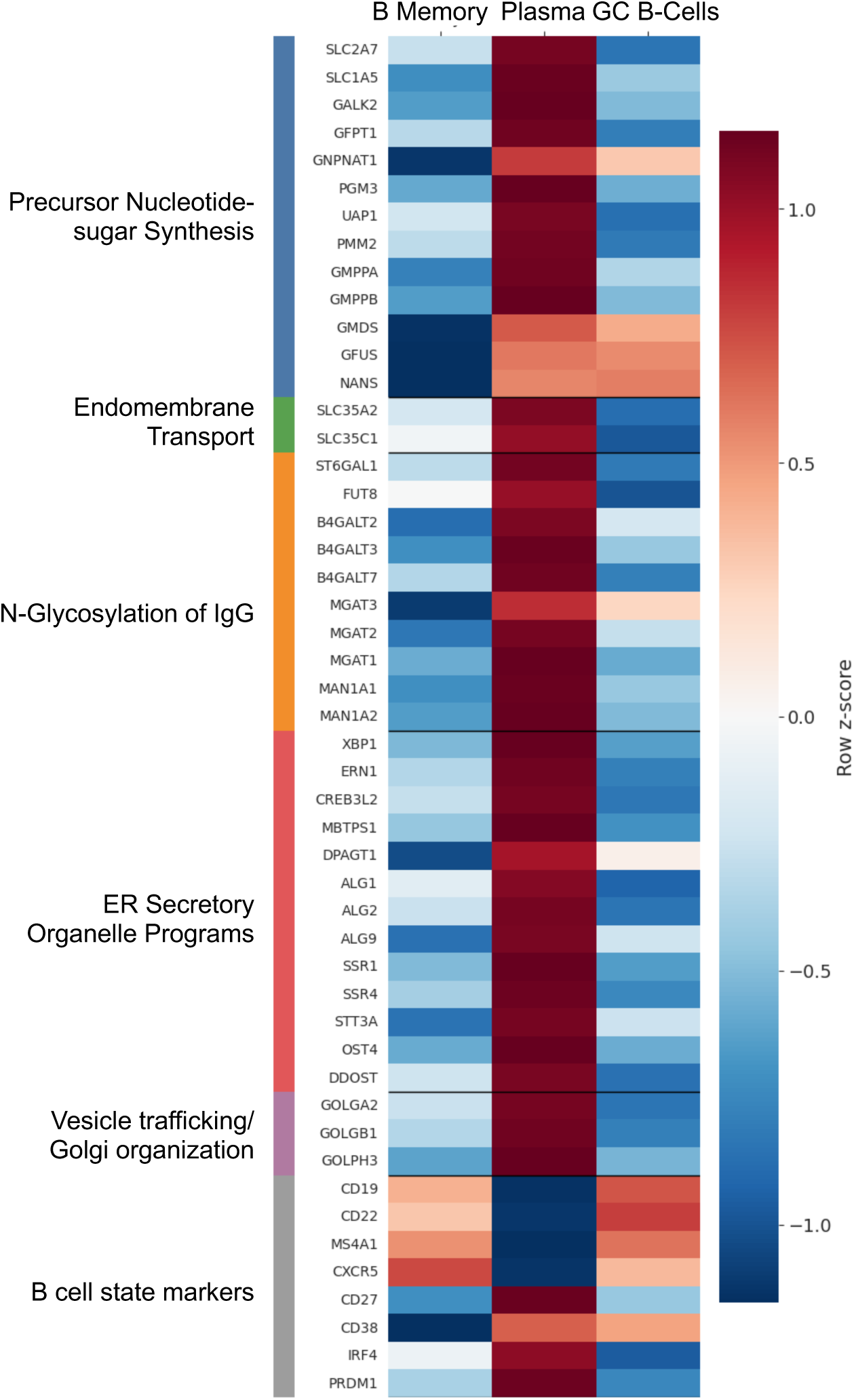
(Linked to Figure 2). **Stomach-resident plasma cells are enriched for secretory and specialized IgG glycosylation gene programs.** Heatmap of selected genes across B memory cells (C0), plasma cells (C4), and germinal center (GC) B cells (C13) isolated from human stomach. Expression values were obtained from the Human Protein Atlas (HPA) single-cell RNA-seq dataset and are reported as nCPM (TMM-normalized counts per million). Values were log2-transformed (log2[nCPM + 1]) and displayed as row-wise z-scores across cell types for each gene. Genes are grouped into functional modules, including metabolic input and precursor synthesis, endomembrane transport, IgG N-glycosylation, ER protein production and quality control, vesicle trafficking and Golgi organization, and B cell identity markers. Plasma cells, in contrast to memory B cells and GC B cells, exhibit coordinated upregulation of secretory pathway regulators and glycosylation-associated genes.

**Suppl. Fig 5.**
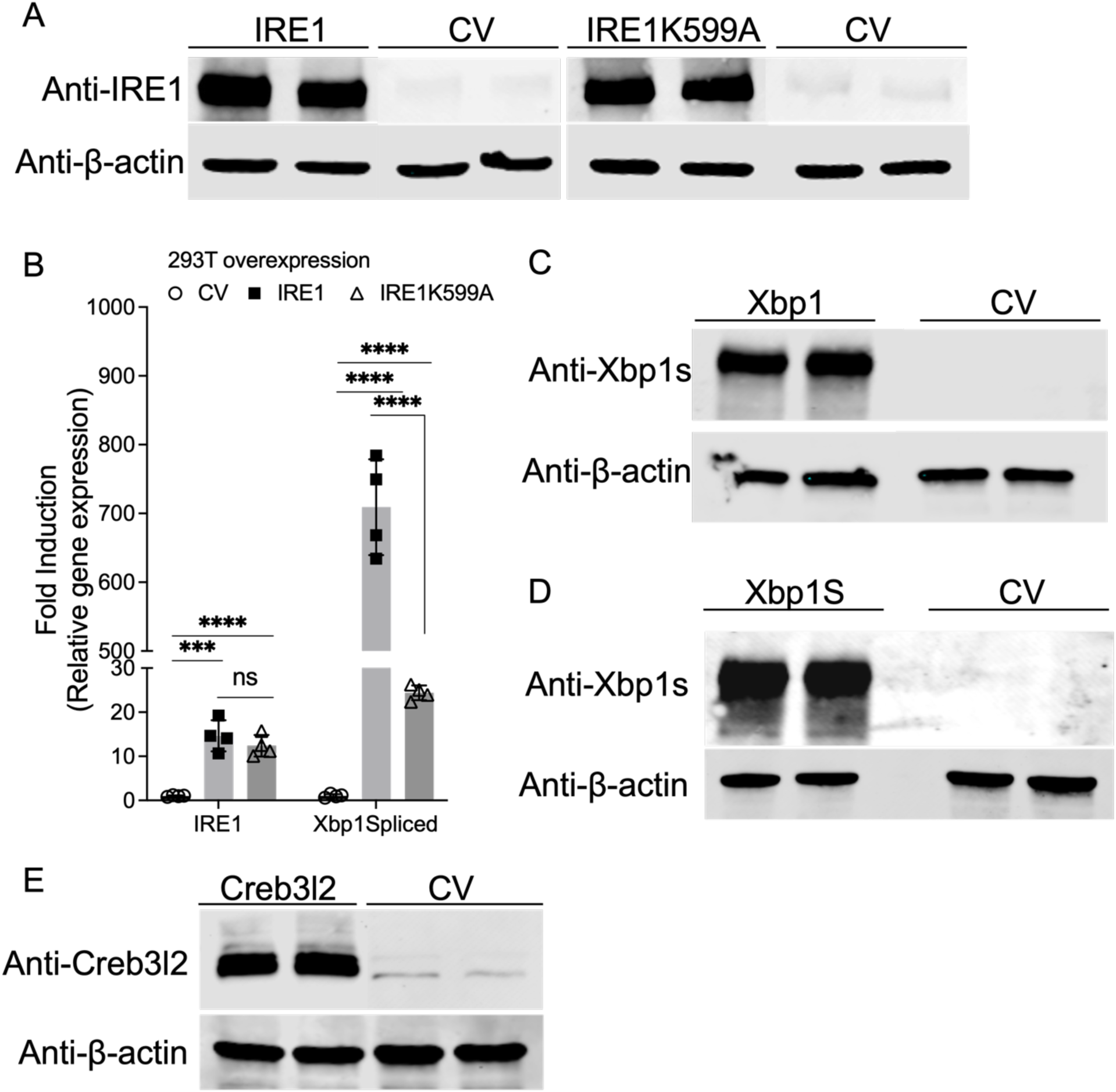
(linked to Figure 3). **Over expression of different IRE1a, XBP1 and Creb3l2 constructs in HEK293T cells. A.** Immunoblot analysis of IRE1 in HEK293T cells transfected with the indicated IRE1α, IRE1α K599A or control vector (CV) (2.5 μg DNA per well, 6-well plate). Cells were harvested 48 h post-transfection. Two representative samples are shown. **B.** RT-PCR analysis of cells transfected as in (A). Data are shown as mean ± SEM from four independent transfections. Statistical significance was determined by unpaired two-tailed Student’s t test versus control vector (CV). *: p value < 0.05; **: p value < 0.01; ***: p value < 0.001; ****: p value < 0.0001. **C–E.** Immunoblot analysis of XBP1s (C-D) and CREB3L2 (E) in HEK293T cells transfected with unspliced Xbp1 (C), spliced Xbp1 (D), or Creb3l2 (E). Cells were harvested 48 h post-transfection. Two representative samples are shown. RT-PCR primer sequences, antibody and construct information are provided in Table S11.

**Suppl. Fig 6.**
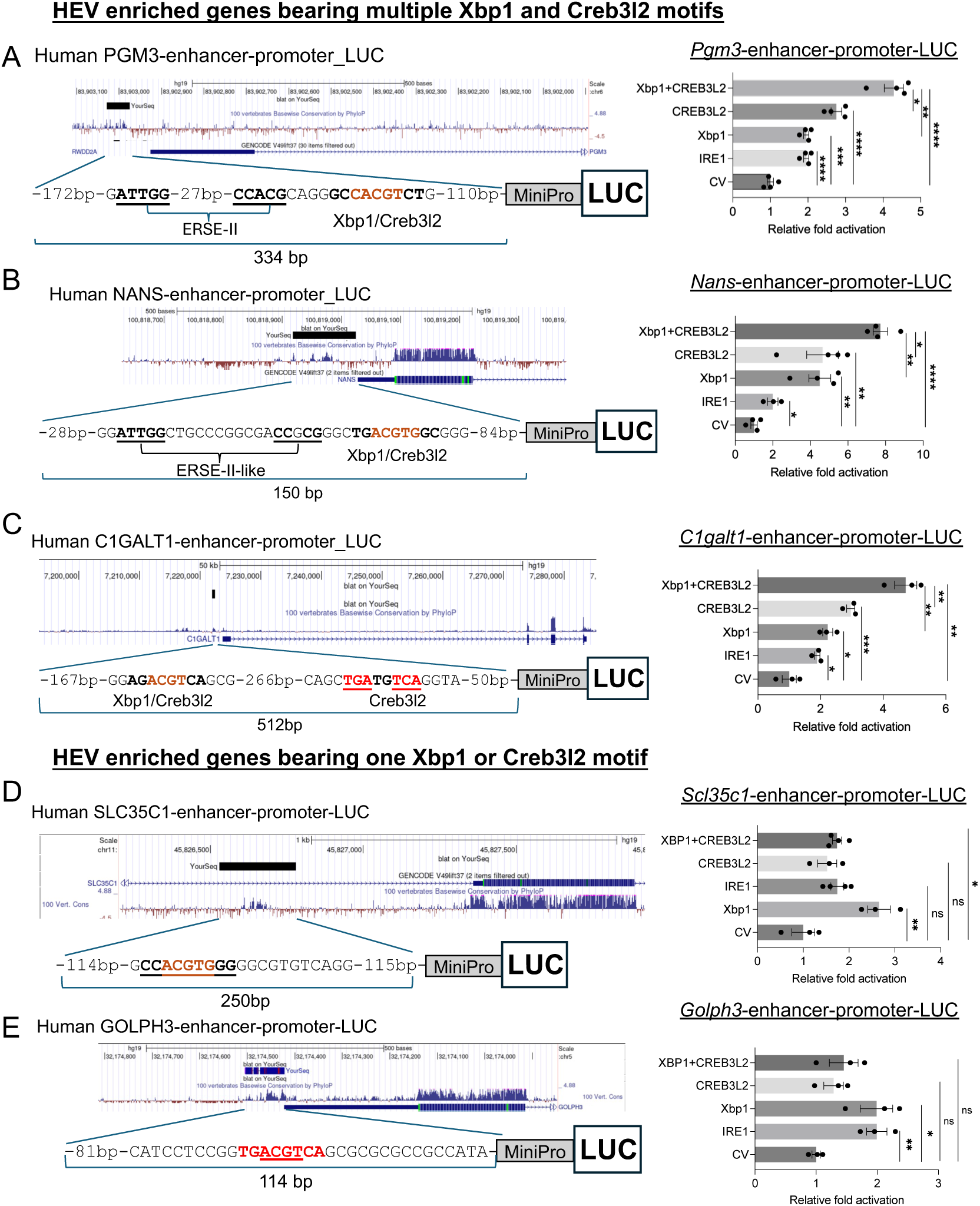
(linked to Figure 3). **XBP1 and CREB3L2 Regulate PNAd Biosynthetic Gene Reporters Through Conserved Regulatory Elements. A–C**. Overexpression of Xbp1s and/or activated CREB3L2 cooperatively enhances luciferase activity driven by the Pgm3, Nans, and C1galt1 enhancer–promoter elements in 293T cells. The locations of the predicted XBP1- and CREB3L2-binding sites, together with ERSEII-like motifs, within each reporter construct are indicated in the left panels. Results are shown as fold induction relative to CV (Control Vector). **D–E.** In contrast, overexpression of Xbp1s and/or activated CREB3L2 does not produce additive or cooperative transactivation of enhancer–promoter constructs harboring a single XBP1 or CREB3L2 binding site (the SLC35C1 and GOLPH3 reporters). The locations of the predicted XBP1-/CREB3L2-binding site within each reporter construct are indicated in the left panels. Results are shown as fold induction relative to CV (Control Vector). All plotted data are from three experiments and are shown with mean ± S.E.M. Statistical significance was determined by unpaired two-tailed Student’s t-test comparing each condition to control vector (CV); Welch’s correction was applied when variances were unequal. *: p value < 0.05; **: p value < 0.01; ***: p value < 0.001; ****: p value < 0.0001.

**Suppl. Fig 7.**
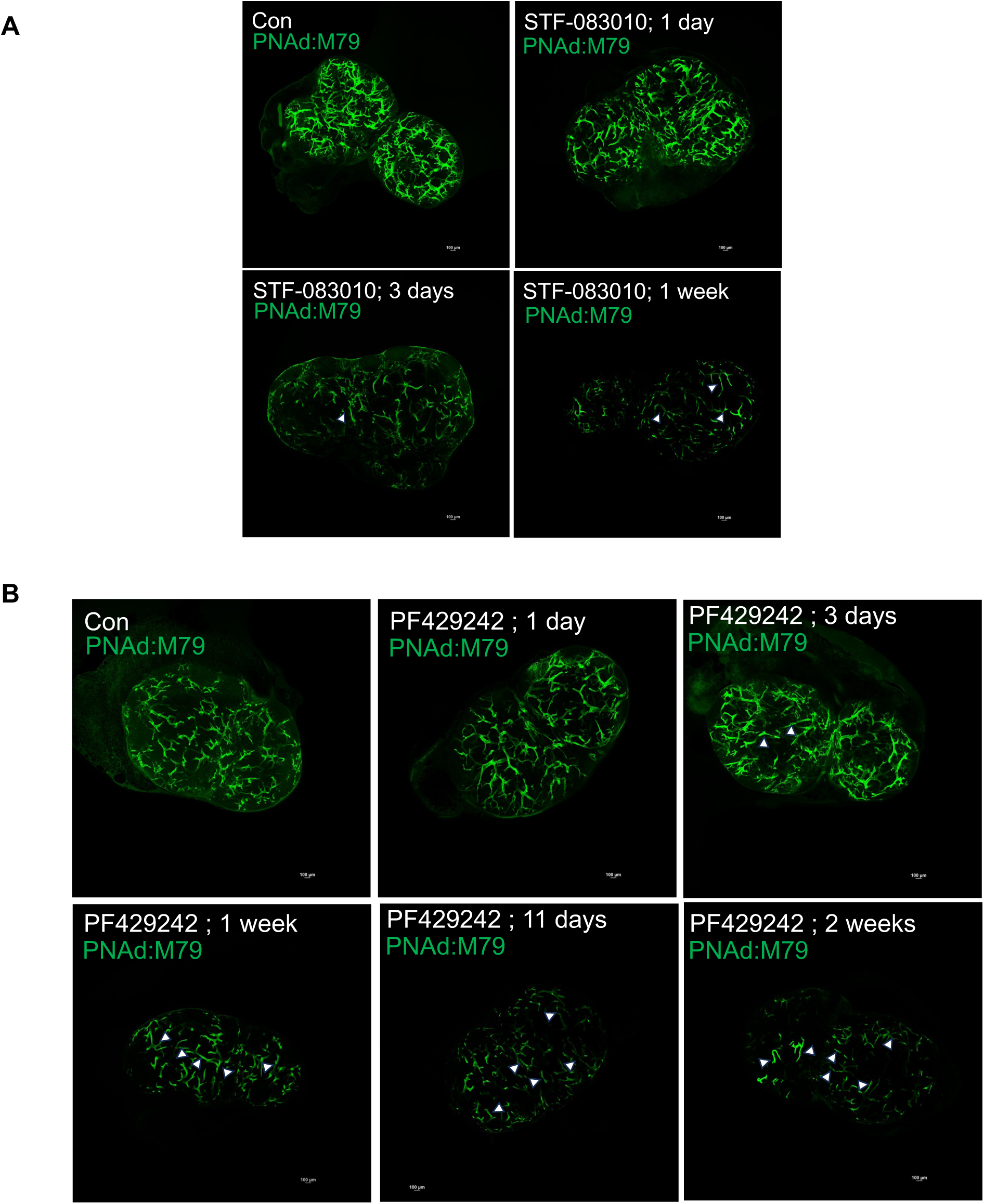
(linked to Figure 4). **Time-course analysis of HEV regression following inhibition of IRE1α and S1P protease activity in vivo, respectively.** Representative whole-mount IF images of PLNs from mice treated with the IRE1α inhibitor STF-083010 (**A**: 3 days and 1 week) or the S1P protease inhibitor PF429242 (**B**: 3, 7, 11 days and 2 weeks), alongside untreated controls. Mice received i.v. injections of antibodies labeling PNAd (MECA-79, green). Arrowheads indicate thin, elongated HEVs. Scale bar: 100 μm.

**Suppl. Fig. 8.**
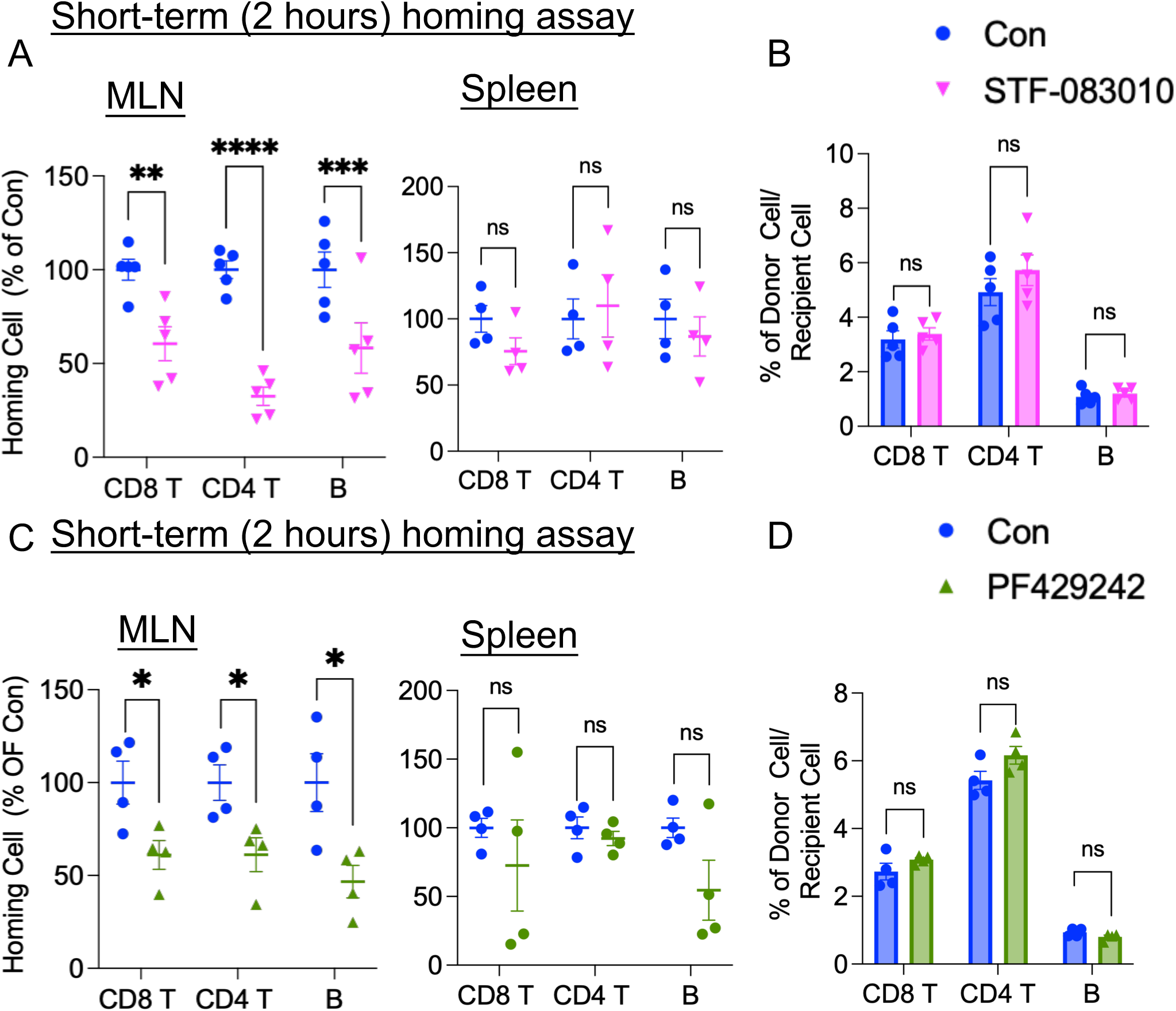
(Linked to Figure 4) **IRE1α or S1P Inhibition Reduces Lymphocyte Homing to Lymph Nodes. A, C.** Localization of donor lymphocytes in MLN and spleen 2 hours after homing in mice treated with STF-083010 (**A**) or PF429242 (**C**), compared to controls. Data represent the percentage of the mean localization ratio relative to control mice. **B, D.** Donor-to-recipient cell count ratios in PLNs from mice treated with STF-083010 (B) or PF429242 (D). **A–D**: Data are presented as mean ± SEM; n = 5 mice per group for A–B and n = 4 mice per group for C–D. Statistical significance was assessed using unpaired two-tailed t-tests. *p < 0.05; **p < 0.01; ***p < 0.001; ****p < 0.0001.

**Suppl. Fig. 9.**
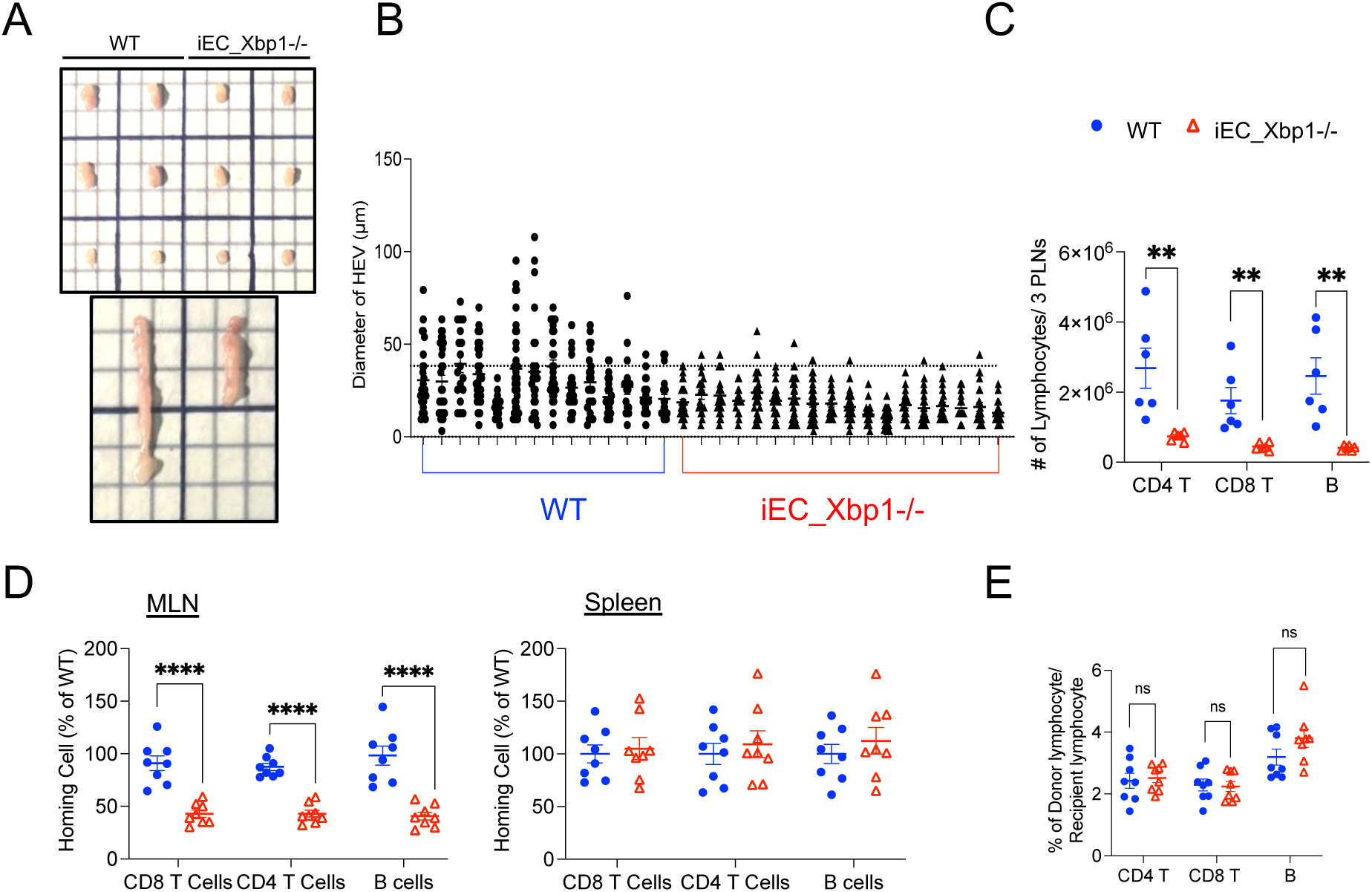
(Linked to Figure 5)**. Endothelial Xbp1 Deletion Alters HEV Morphology and Lymphocyte Accumulation. A.** Representative images of LNs. Upper panel: from top to bottom—inguinal PLN, axillary PLN, and brachial PLN. Lower panel: MLN. **B**. Quantification of HEV diameter. Data represent 14–35 diameter measurements per 10× field (∼4 mm²/field), averaged across 2–3 representative fields per PLN, with 1–2 PLNs analyzed per mouse (n = 5 mice per group). Each dot corresponds to an individual measurement. **C.** Total lymphocyte counts from three pooled peripheral lymph nodes (inguinal, axillary, and brachial) collected from each mouse. Each dot represents one mouse (WT: n = 6; iEC_Xbp1-/- n=5), and bars indicate the mean ± SEM. **D.** Localization of donor lymphocytes in the MLN and spleen of control and iEC Xbp1-deficient mice following a 2-hour short-term homing assay. Data are shown as the percentage of the mean localization ratio relative to the WT group. Each dot represents one mouse (n = 8 per group); bars indicate mean ± SEM. **E.** Quantification of donor lymphocytes as a percentage of total recipient lymphocytes recovered following the short-term homing assay. Each dot represents one mouse (n = 8 per group); bars indicate the mean ± SEM. **For panels C- E,** two-tailed unpaired t-tests were used for all comparisons. *p < 0.05; **p < 0.01; ***p < 0.001; ****p < 0.0001.

**Suppl. Fig. 10.**
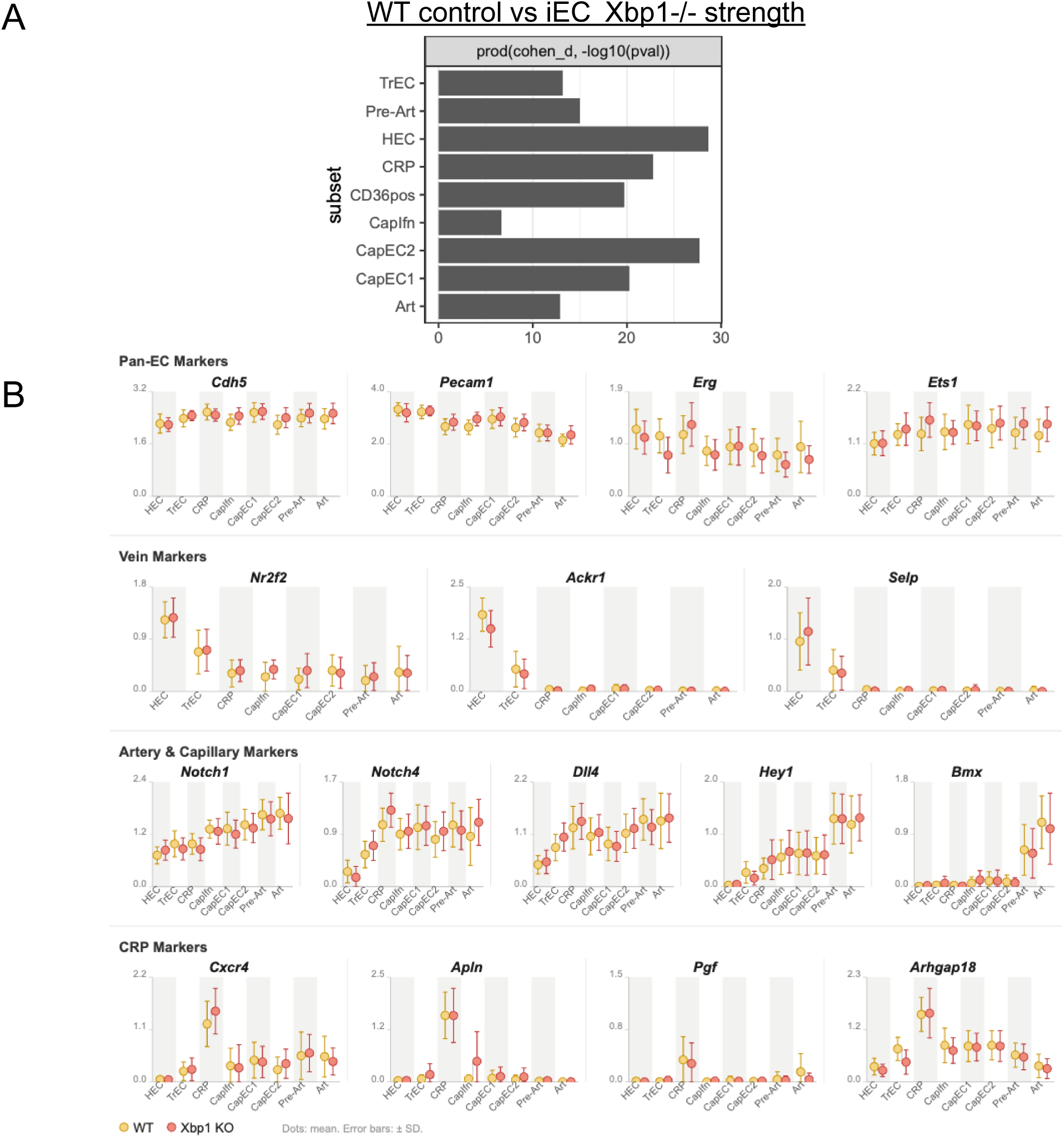
(Link to Figure. 6). **scRNA-seq transcriptomic analysis of endothelial subsets in LNs of WT littermates and iEC_Xbp1−/− mice. A.** Transcriptional perturbation across EC subsets, calculated as Cohen’s d × −log10(p value) using the top 100 DEGs per subset. **B.** Normalized mean expression of pan-EC markers (Cdh5, Pecam1, Erg, Ets1), vein markers (Nr2f2, Ackr1, Selp), artery and capillary markers (Notch1, Notch4, Dll4, Hey1, Bmx), and CRP markers (Cxcr4, Apln, Pgf, Arhgap18) across EC subsets from LNs. Each dot represents mean normalized expression. Error bars indicate ± SD. yellow, WT; red, iEC_Xbp1−/−. EC subsets: HEC, high endothelial cells; TrEC, transitional ECs; CRP, capillary-related population; CapIfn, interferon-responsive capillary ECs; CapEC1/2, capillary ECs 1 and 2; Pre-Art, pre-arterial ECs; Art, arterial ECs. n = 2 mice per group.

**Suppl. Fig. 11.**
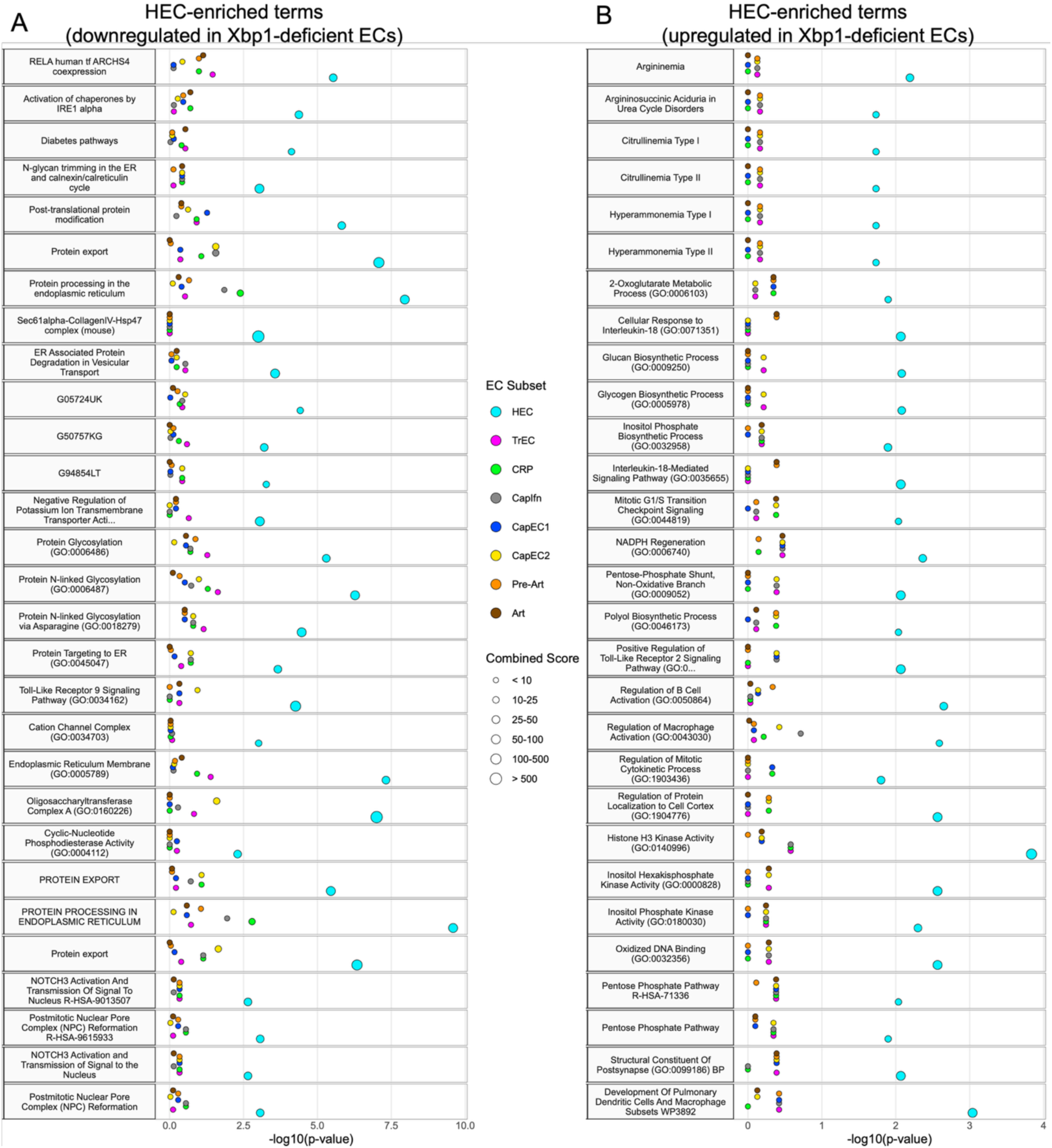
(Link to Fig. 6). **HEC-enriched gene programs across WT and iEC_Xbp1-deficient EC populations.** Enrichr-based pathway and Gene Ontology enrichment analysis of DEGs (top 2,000) in HECs from WT and iEC_Xbp1−/− LNs, showing enriched terms derived from genes downregulated (A) and upregulated (B) in Xbp1-deficient (KO) cells relative to WT. The top 30 enriched terms ranked by specificity score (spec) are shown for visualization and are provided in Table S10. Complete Enrichr results are provided in Table S8 (downregulated gene set) and Table S9 (upregulated gene set). The x-axis represents −log₁₀(P value). Dot size indicates the combined enrichment score, and dot color denotes EC subset identity.

**Suppl. Fig. 12.**
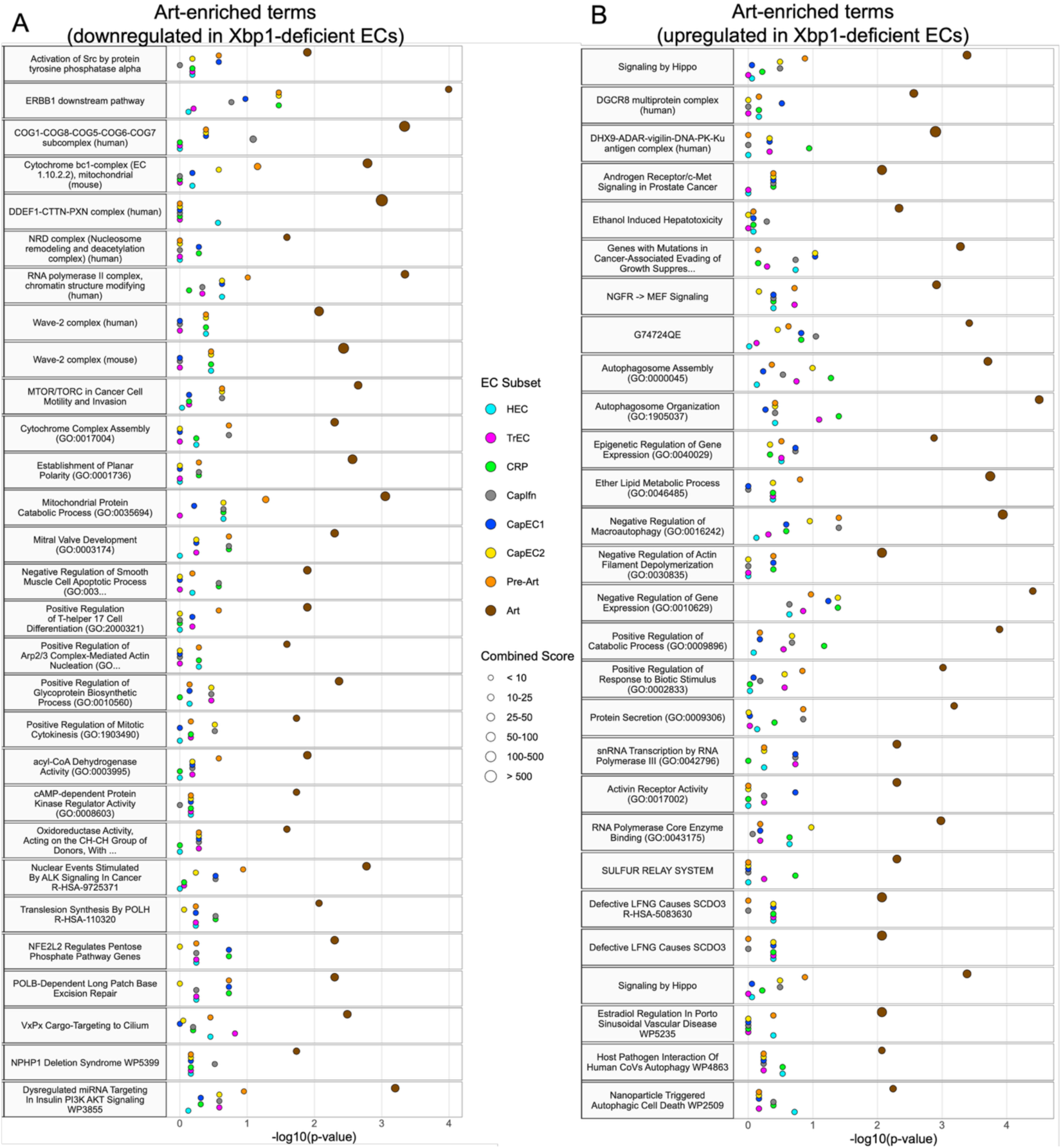
(Link to Fig. 6)**. Art enriched gene programs across WT vs iEC Xbp1-deficient EC populations.** Enrichr-based pathway and Gene Ontology enrichment analysis of DEGs (top 2,000) in arterial ECs from WT and iEC_Xbp1−/− LNs, showing enriched terms derived from genes downregulated (A) and upregulated (B) in Xbp1-deficient (KO) cells relative to WT. The top 30 enriched terms ranked by specificity score (spec) are shown for visualization and are provided in Table S10. Complete Enrichr results are provided in Table S8 (downregulated gene set) and Table S9 (upregulated gene set). The x-axis represents −log₁₀(P value). Dot size indicates the combined enrichment score, and dot color denotes EC subset identity.

**Suppl. Fig. 13.**
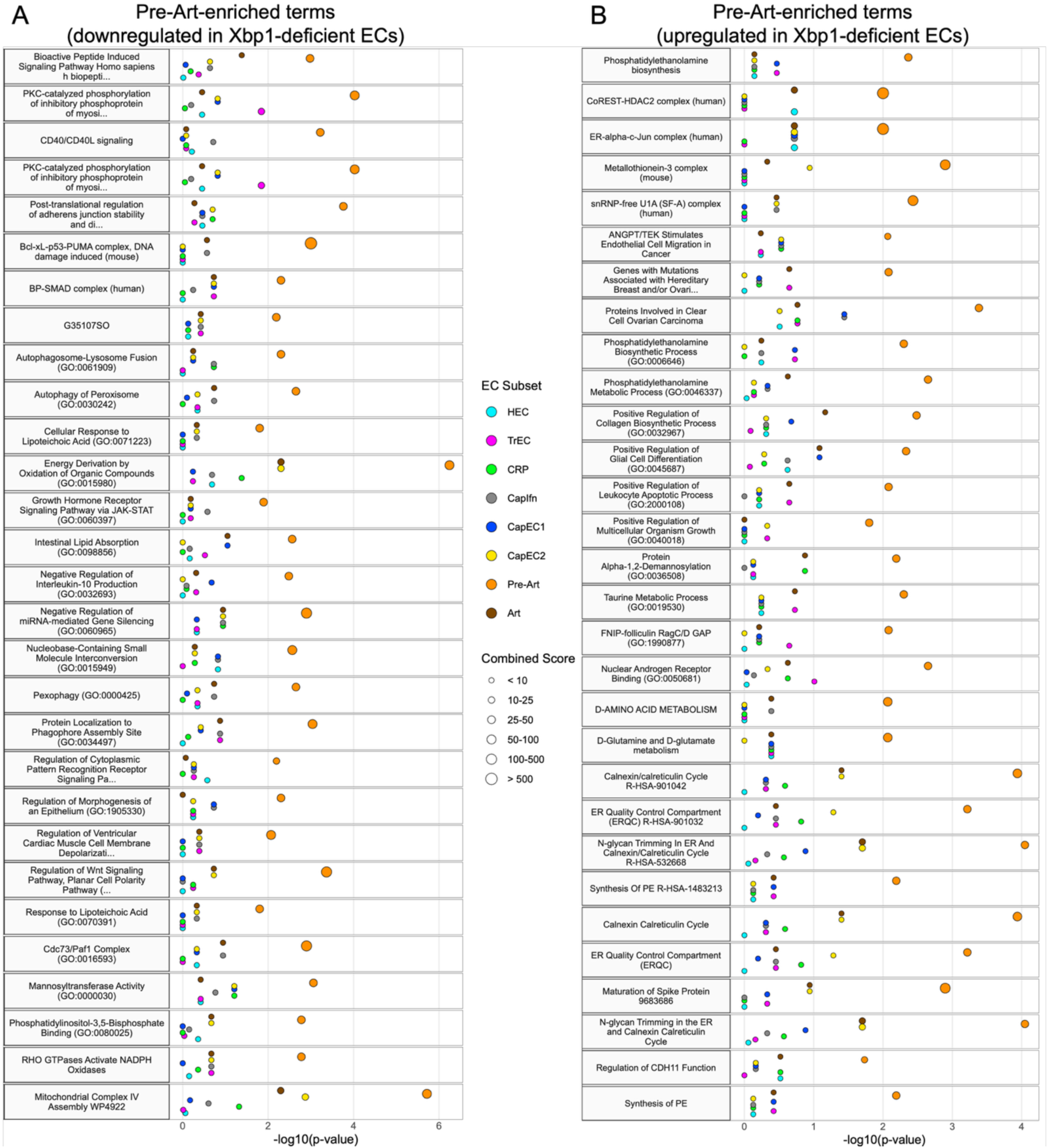
(Link to Fig. 6)**. Pre-Art enriched gene programs across WT vs iEC Xbp1-deficient EC populations.** Enrichr-based pathway and Gene Ontology enrichment analysis of DEGs (top 2,000) in pre-arterial ECs from WT and iEC_Xbp1−/− LNs, showing enriched terms derived from genes downregulated (A) and upregulated (B) in Xbp1-deficient (KO) cells relative to WT. The top 30 enriched terms ranked by specificity score (spec) are shown for visualization and are provided in Table S10. Complete Enrichr results are provided in Table S8 (downregulated gene set) and Table S9 (upregulated gene set). The x-axis represents −log₁₀(P value). Dot size indicates the combined enrichment score, and dot color denotes EC subset identity.

**Suppl. Fig. 14.**
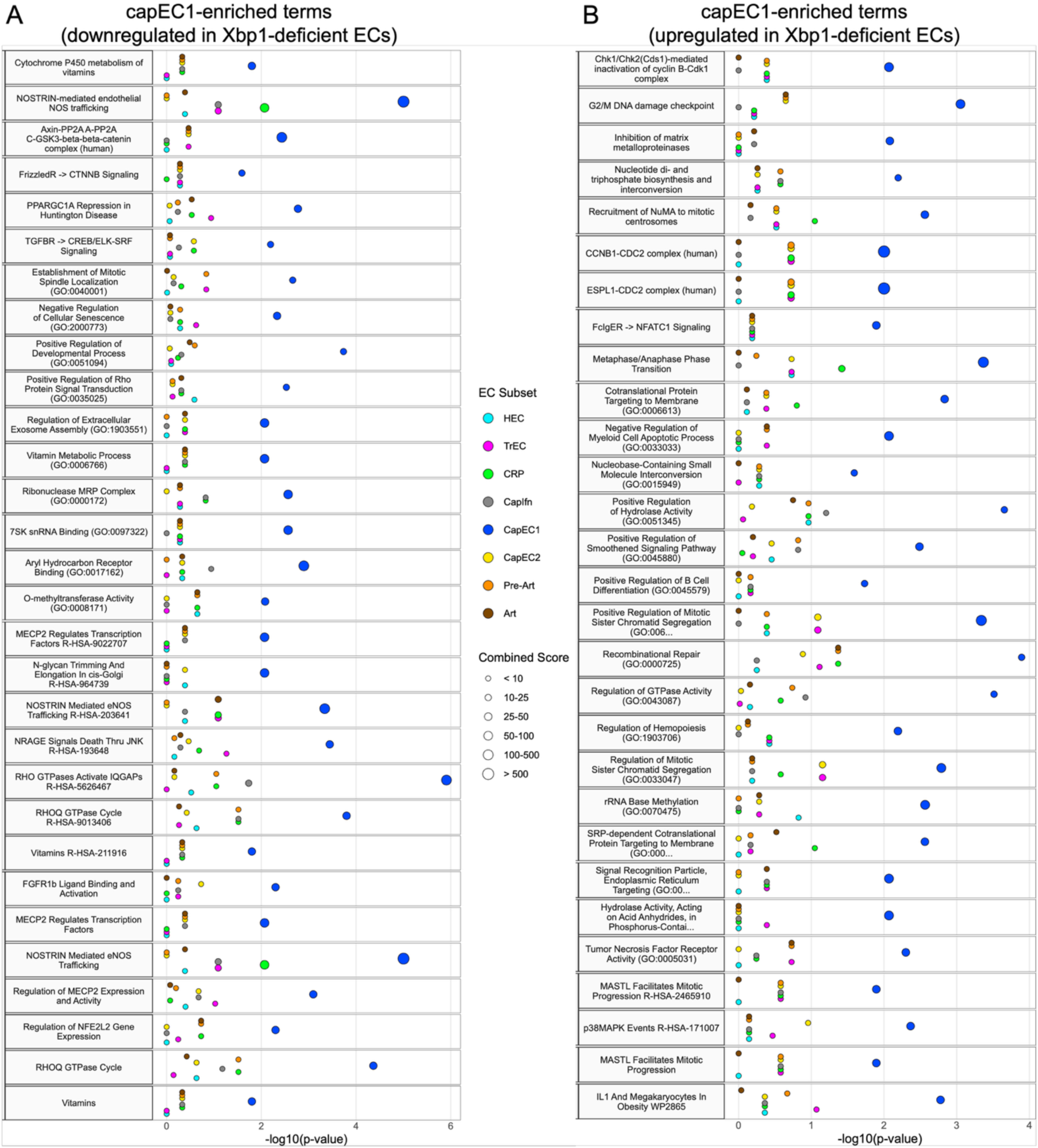
A. (Link to Fig. 6)**. capEC1 enriched gene programs across WT vs iEC Xbp1-deficient EC populations.** Enrichr-based pathway and Gene Ontology enrichment analysis of DEGs (top 2,000) in capEC1 from WT and iEC_Xbp1−/− LNs, showing enriched terms derived from genes downregulated (A) and upregulated (B) in Xbp1-deficient (KO) cells relative to WT. The top 30 enriched terms ranked by specificity score (spec) are shown for visualization and are provided in Table S10. Complete Enrichr results are provided in Table S8 (downregulated gene set) and Table S9 (upregulated gene set). The x-axis represents −log₁₀(P value). Dot size indicates the combined enrichment score, and dot color denotes EC subset identity.

**Suppl. Fig. 15.**
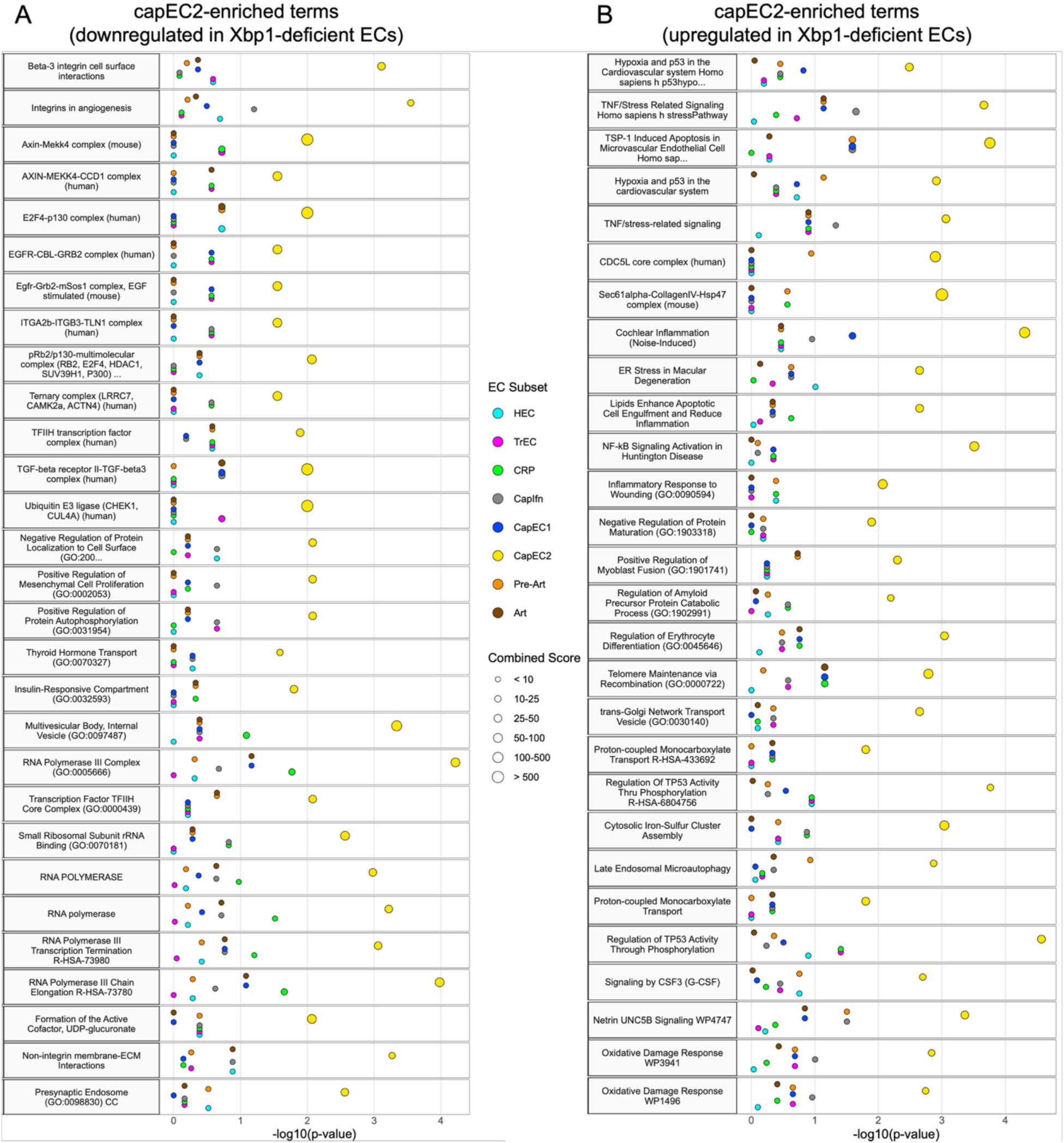
A. (Link to Fig. 6)**. capEC2 enriched gene programs across WT vs iEC Xbp1-deficient EC populations.** Enrichr-based pathway and Gene Ontology enrichment analysis of DEGs (top 2,000) in capEC2 from WT and iEC_Xbp1−/− LNs, showing enriched terms derived from genes downregulated (A) and upregulated (B) in Xbp1-deficient (KO) cells relative to WT. The top 30 enriched terms ranked by specificity score (spec) are shown for visualization and are provided in Table S10. Complete Enrichr results are provided in Table S8 (downregulated gene set) and Table S9 (upregulated gene set). The x-axis represents −log₁₀(P value). Dot size indicates the combined enrichment score, and dot color denotes EC subset identity.

**Suppl. Fig. 16.**
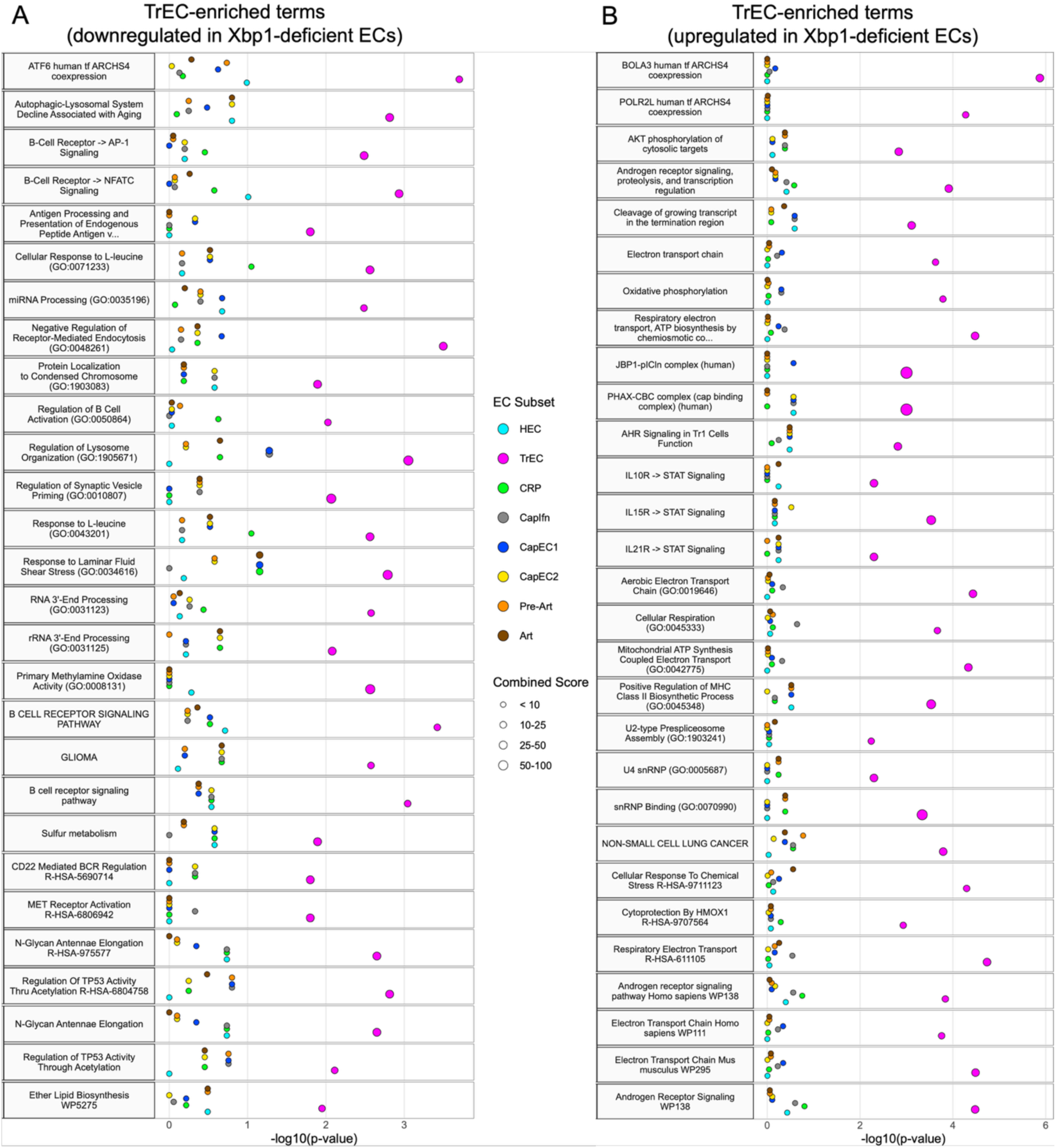
(Link to Fig. 6)**. TrEC enriched gene programs across WT vs iEC Xbp1-deficient EC populations.** Enrichr-based pathway and Gene Ontology enrichment analysis of DEGs (top 2,000) in TrECs from WT and iEC_Xbp1−/− LNs, showing enriched terms derived from genes downregulated (A) and upregulated (B) in Xbp1-deficient (KO) cells relative to WT. The top 30 enriched terms ranked by specificity score (spec) are shown for visualization and are provided in Table S10. Complete Enrichr results are provided in Table S8 (downregulated gene set) and Table S9 (upregulated gene set). The x-axis represents −log₁₀(P value). Dot size indicates the combined enrichment score, and dot color denotes EC subset identity.

**Suppl. Fig. 17.**
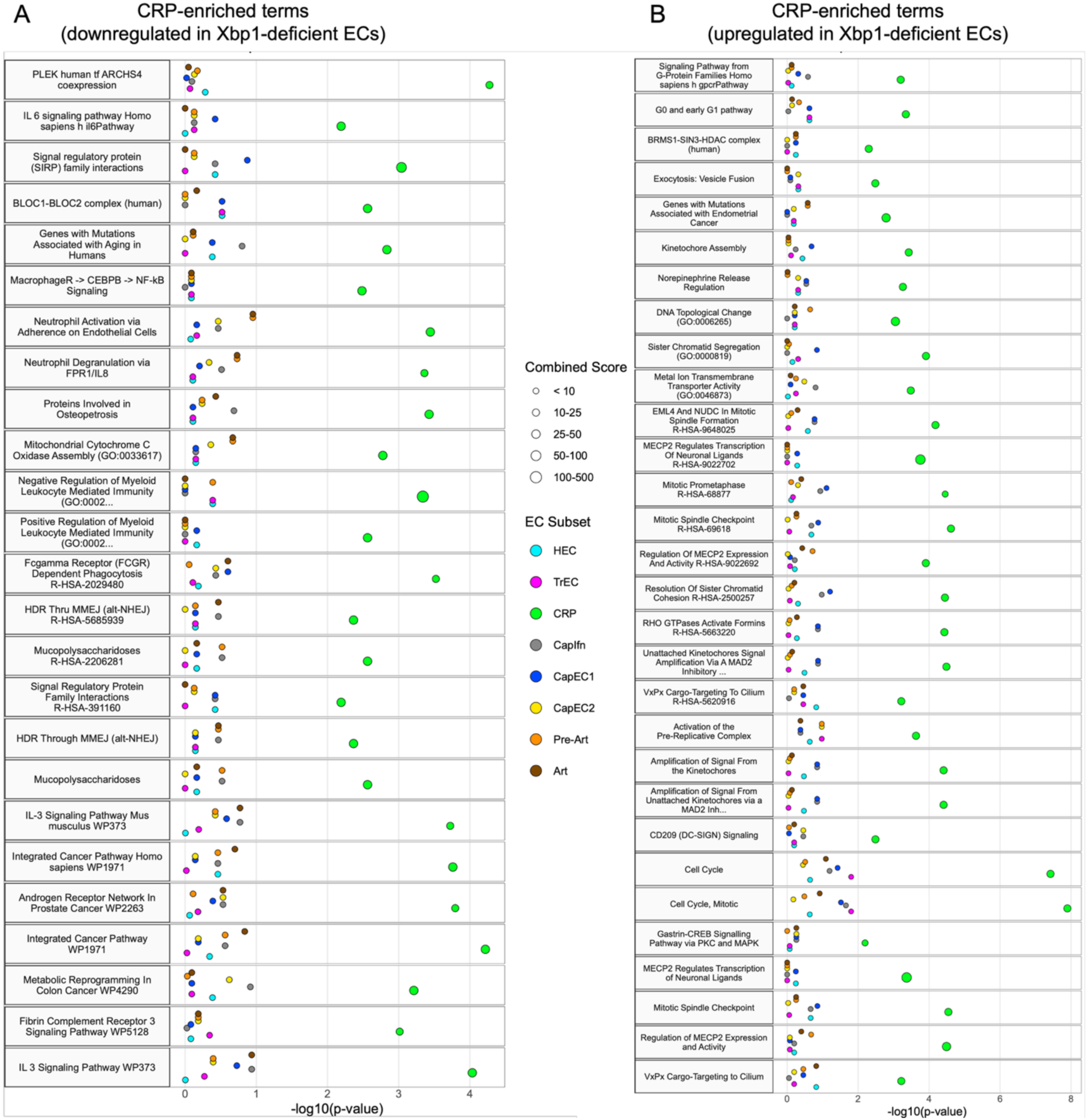
(Link to Fig. 6)**. CRP enriched gene programs across WT vs iEC Xbp1-deficient EC populations.** Enrichr-based pathway and Gene Ontology enrichment analysis of DEGs (top 2,000) in CRPs from WT and iEC_Xbp1−/− LNs, showing enriched terms derived from genes downregulated (A) and upregulated (B) in Xbp1-deficient (KO) cells relative to WT. The top 30 enriched terms ranked by specificity score (spec) are shown for visualization and are provided in Table S10. Complete Enrichr results are provided in Table S8 (downregulated gene set) and Table S9 (upregulated gene set). The x-axis represents −log₁₀(P value). Dot size indicates the combined enrichment score, and dot color denotes EC subset identity.

**Suppl. Fig. 18.**
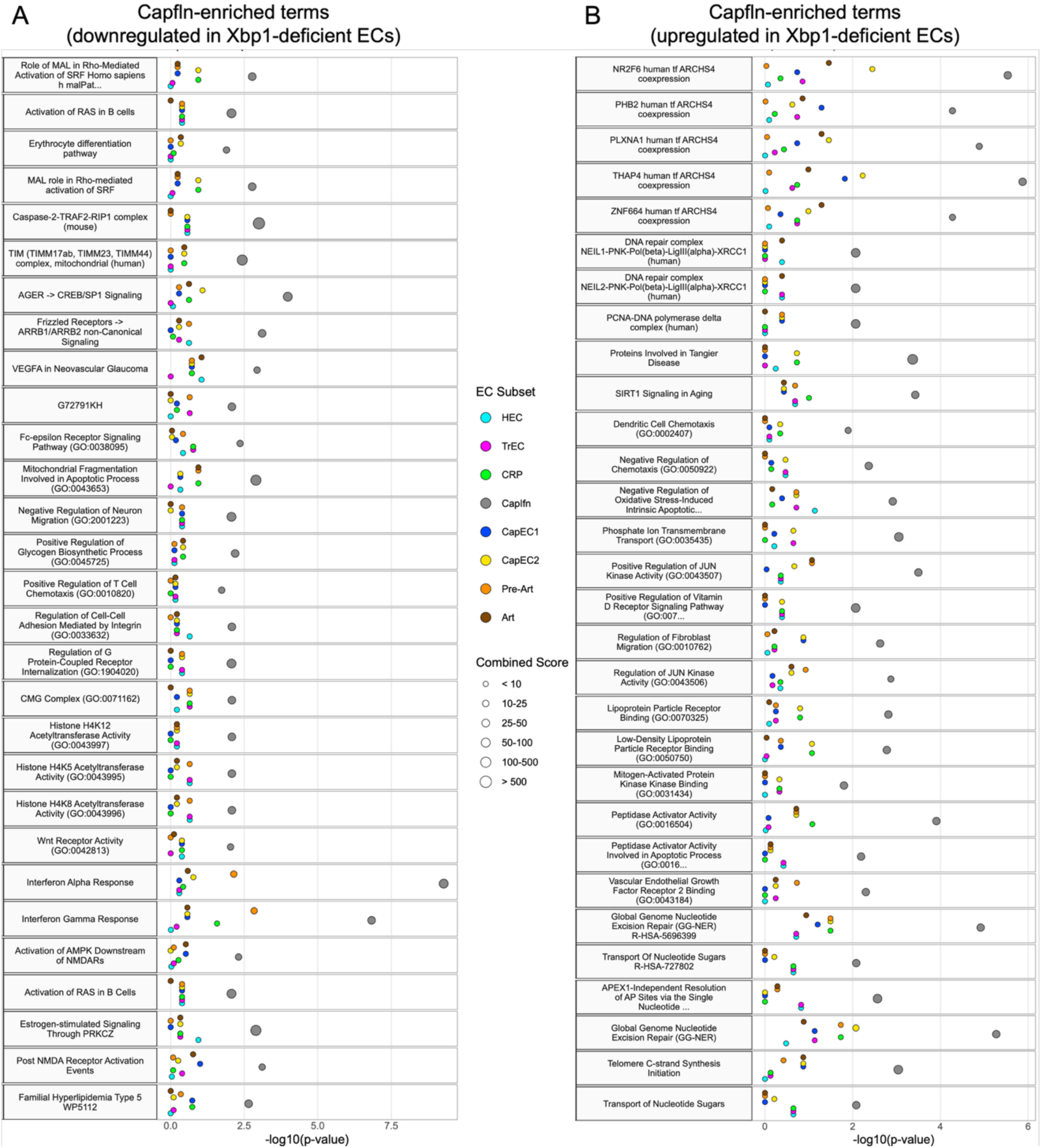
(Link to Fig. 6)**. Capfln enriched gene programs across WT vs iEC Xbp1-deficient EC populations.** Enrichr-based pathway and Gene Ontology enrichment analysis of DEGs (top 2,000) in Capfln from WT and iEC_Xbp1−/− LNs, showing enriched terms derived from genes downregulated (A) and upregulated (B) in Xbp1-deficient (KO) cells relative to WT. The top 30 enriched terms ranked by specificity score (spec) are shown for visualization and are provided in Table S10. Complete Enrichr results are provided in Table S8 (downregulated gene set) and Table S9 (upregulated gene set). The x-axis represents −log₁₀(P value). Dot size indicates the combined enrichment score, and dot color denotes EC subset identity.

**Suppl. Fig. 19.**
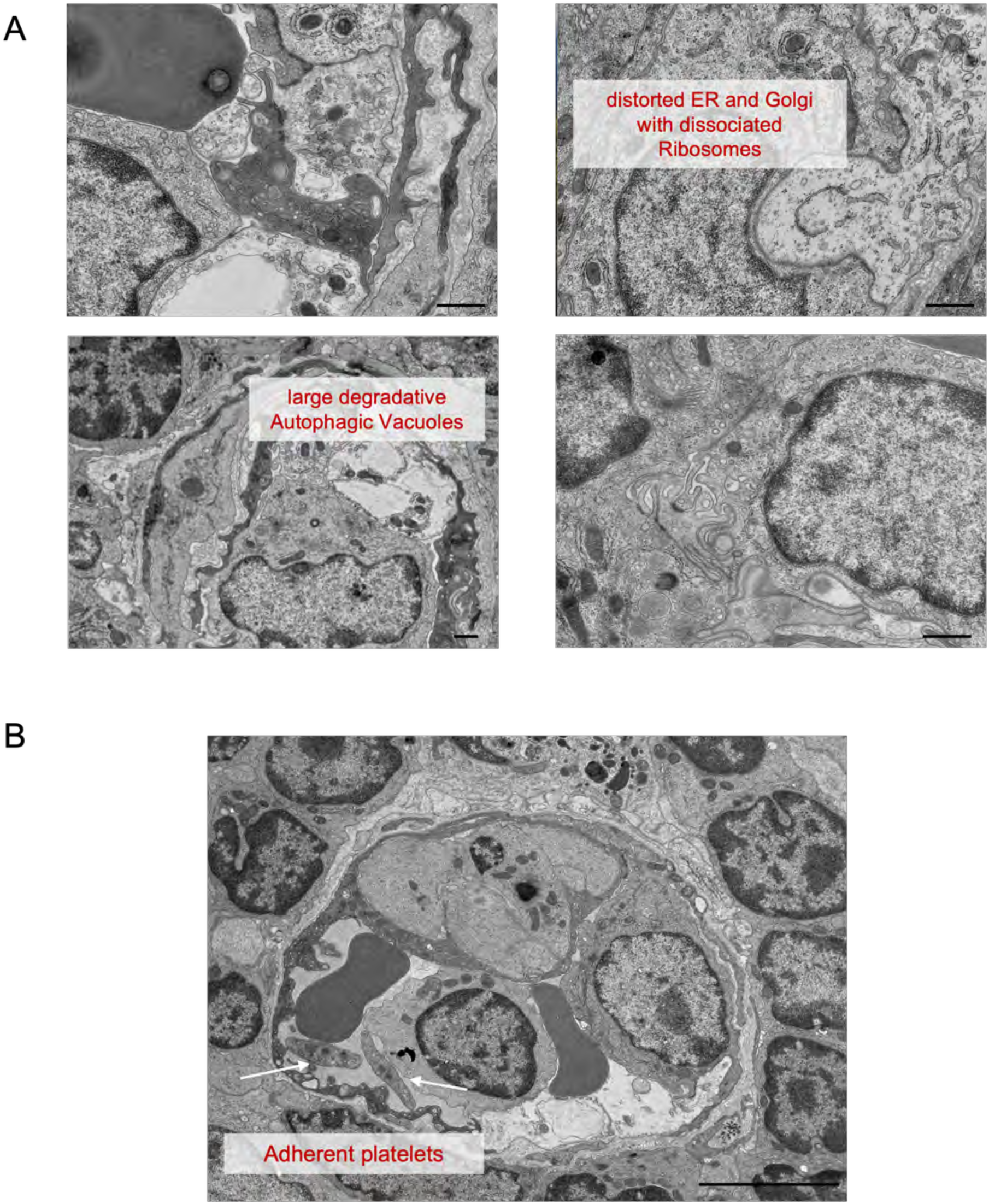
(Linked to Fig. 6). **Ultrastructural defects in iEC Xbp1-deficient HECs. A.** iEC Xbp1-deficient HECs exhibit large degradative autophagic vacuoles and markedly distorted ER and Golgi membranes with dissociated ribosomes. **B.** Adherent platelets (arrows) are indicated. Scale bars, 5 μm.

**Suppl. Fig. 20.**
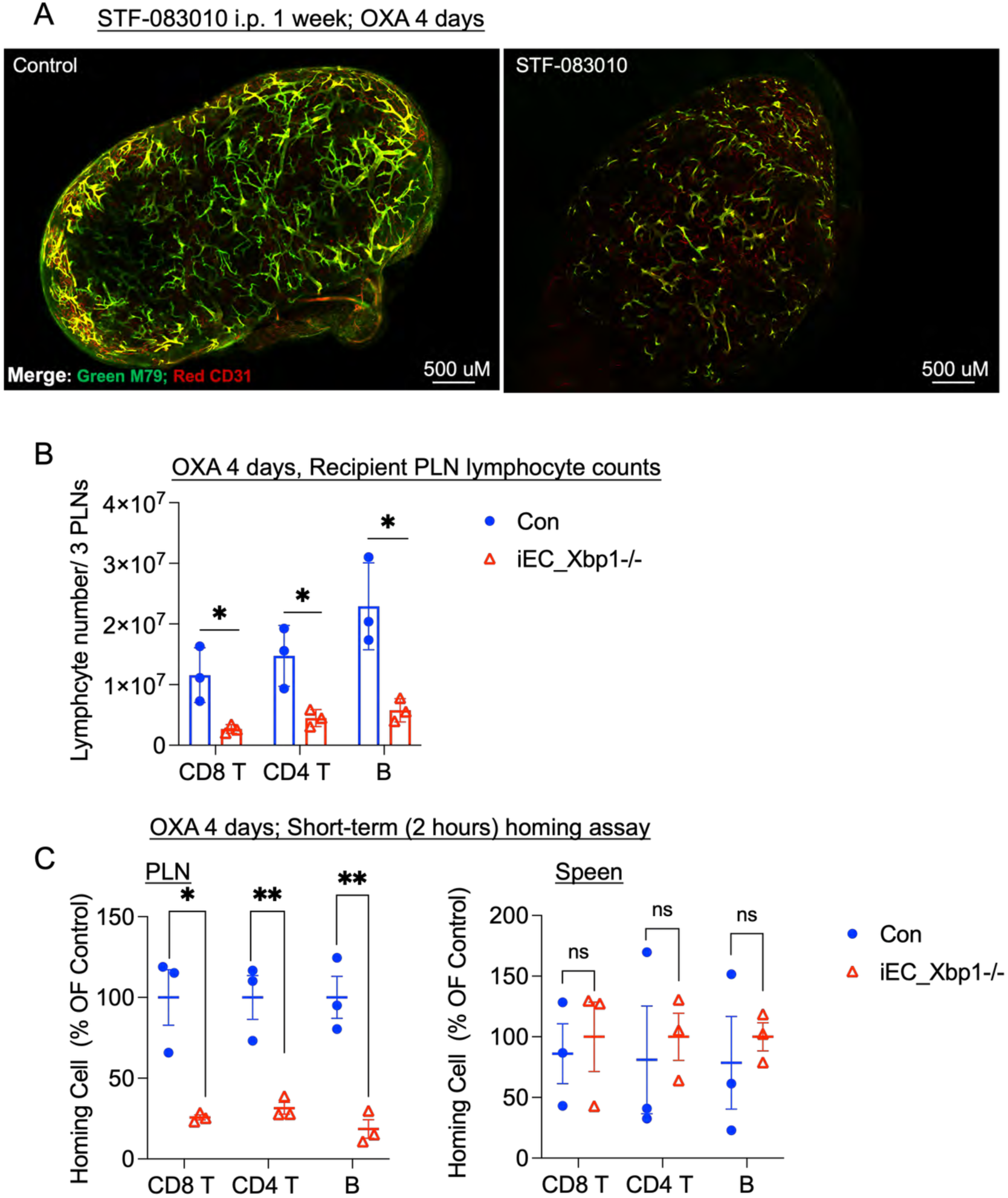
(Linked to Figure 7A)**. Xbp1 Deficiency Reduces Lymphocyte Recruitment to Inflamed Lymph Nodes during Immune Activation. A.** Representative whole-mount staining of OXA-painted lymph nodes on Day 4 in control and STF-083010–treated mice. Mice received i.v. injections of MECA-79 (green) and CD31 (red) antibodies to visualize PNAd⁺ HEVs and total vasculature, respectively. Data are representative of three independent experiments (each including one control and one STF-083010–treated mouse). **B–C.** Short-term homing of lymphocytes into OXA-painted lymph nodes on Day 4 in WT and iEC Xbp1-deficient mice. Mice were topically painted with OXA on Day 1, and homing assays were performed on Day 4. Fluorescently labeled lymphocytes were intravenously transferred, and homing was quantified after a 2-hour homing period. **B.** Total recipient lymphocyte counts from three pooled PLNs (inguinal, axillary, and brachial) collected from each mouse. Each dot represents one mouse (n = 3 per group); bars show mean ± SEM. **C.** Localization of donor lymphocytes in PLNs and spleen (control tissue) of WT and iEC Xbp1-deficient mice following the 2-hour homing assay. Data are presented as the percentage of the mean localization ratio relative to the WT group. Each dot represents one mouse (n = 3 per group); bars indicate mean ± SEM. **For panels B-C,** two-tailed unpaired t-tests were used for all comparisons. *p < 0.05; **p < 0.01.

**Suppl. Fig. 21.**
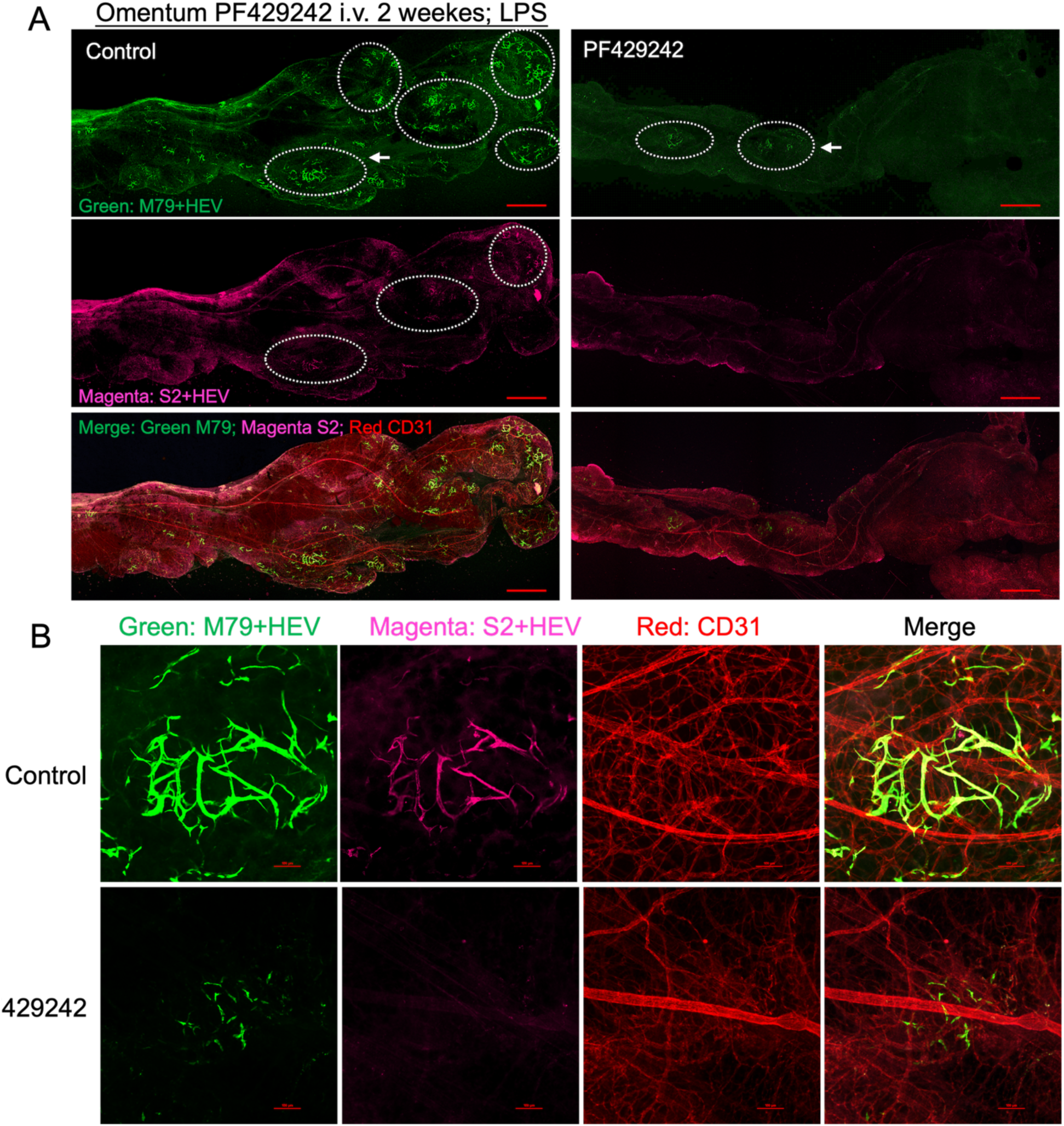
(Linked to Fig. 7, S1P Protease inhibition part) **S1P Protease Inhibition Reduces Inflammation-Induced PNAd⁺ Vessel Formation in the Omentum. A.** Representative whole-mount staining of the omentum from control and PF429242-treated mice following two intraperitoneal injections of LPS over 72 hours. To visualize HEVs, mice were injected intravenously with antibodies labeling PNAd (MECA-79: Green; S2: Magenta) and CD31 (red). Dashed circles indicate regions containing induced HEVs. **B.** Representative images of omental milky spots showing induction of PNAd⁺ vessels within CD31⁺ vasculature (red) after PF429242 treatment. Arrows in A highlight the location of induced HEVs shown in B. Scale bar, 100 µm. Data are representative of three experiments (one control and one PF429242–treated mouse per experiment).

**Suppl. Fig. 22.**
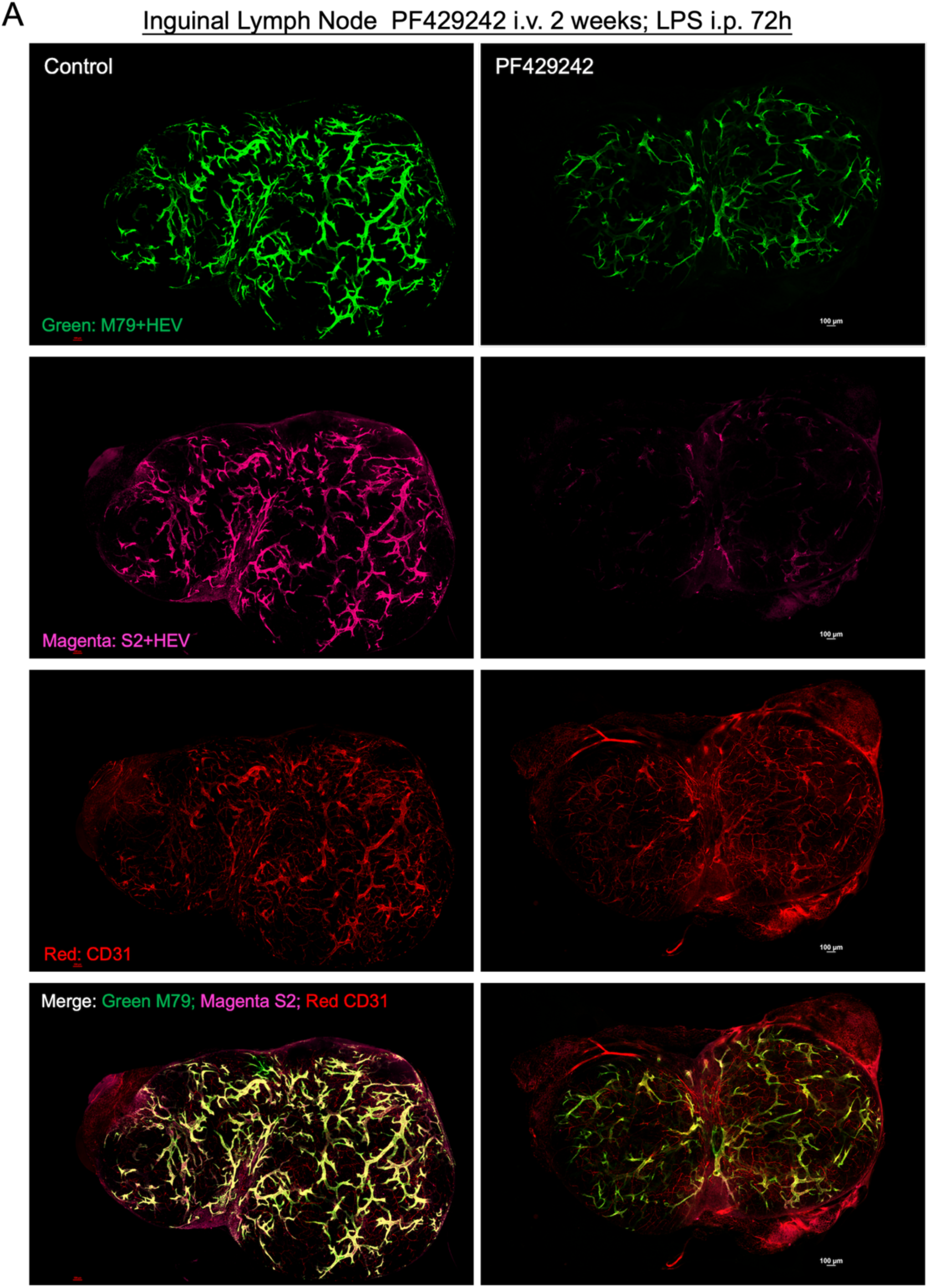
(Linked to Fig. 7, S1P Protease inhibition part)**. S1P Protease Inhibition Attenuates Remodeling of PNAd⁺ Vessels in Peripheral Lymph Nodes during Inflammation. A.** Representative whole-mount staining of the PLNs from control and PF429242-treated mice following two intraperitoneal injections of LPS over 72 hours. To visualize HEVs, mice were injected intravenously with antibodies labeling PNAd (MECA-79; green, S2; magenta) and CD31 (red). Scale bar,100 µm. Data are representative of three experiments (one control and one PF429242–treated mouse per experiment).

**Suppl. Fig. 23.**
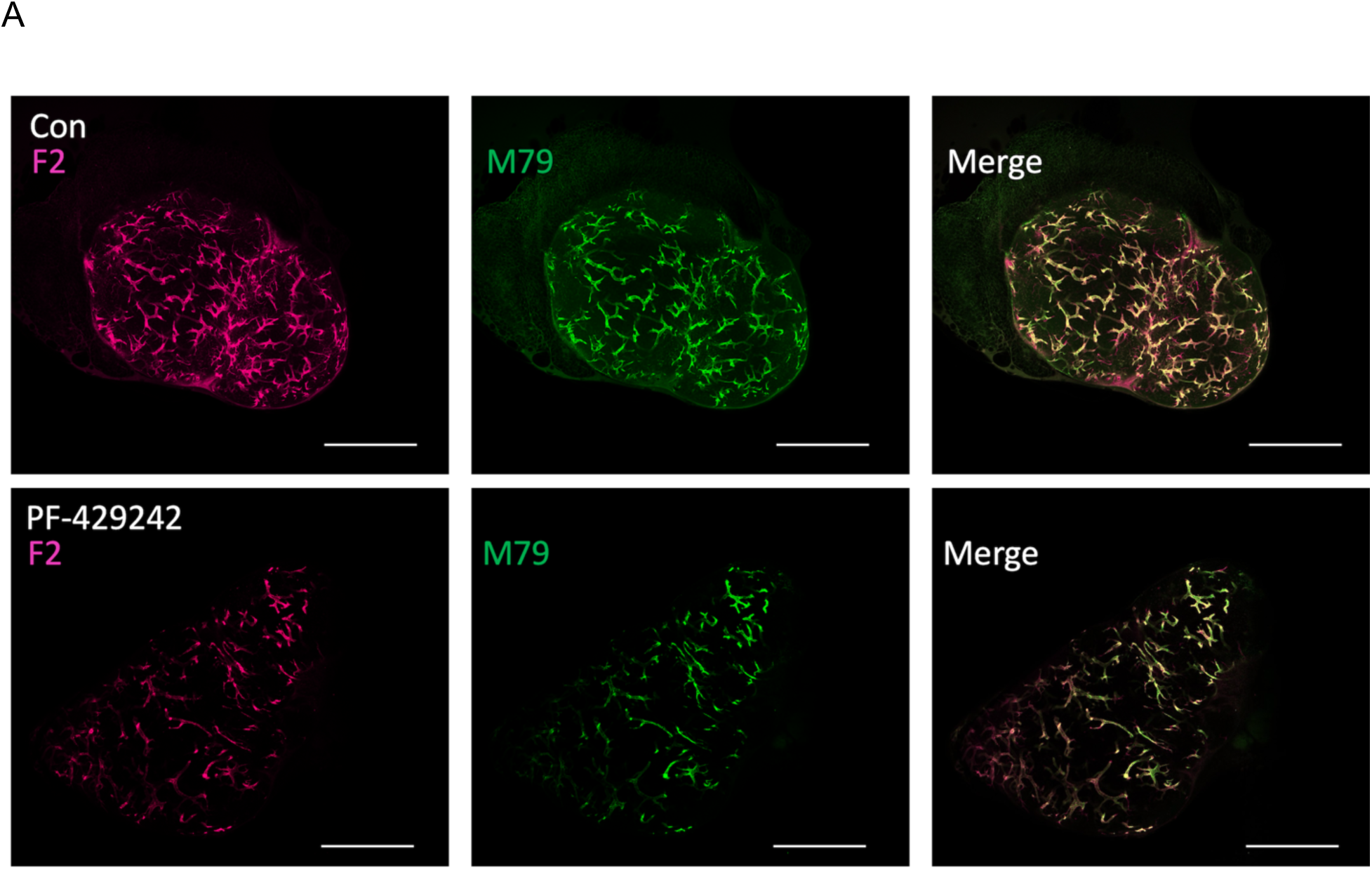
(Linked to Fig. 7, S1P Protease inhibition part)**. Differential PNAd glyco-epitope recognition following S1P inhibition. A.** Representative whole-mount IF images of PLNs from mice treated with the S1P inhibitor PF429242 (bottom; 2 weeks), alongside untreated controls (top). Mice were injected i.v. with antibodies labeling MECA-79 (green) and F2 (magenta). Scale bar: 1000 µm. Data are representative of three experiments (one control and one PF429242–treated mouse per experiment).

**Suppl. Fig. 24.**
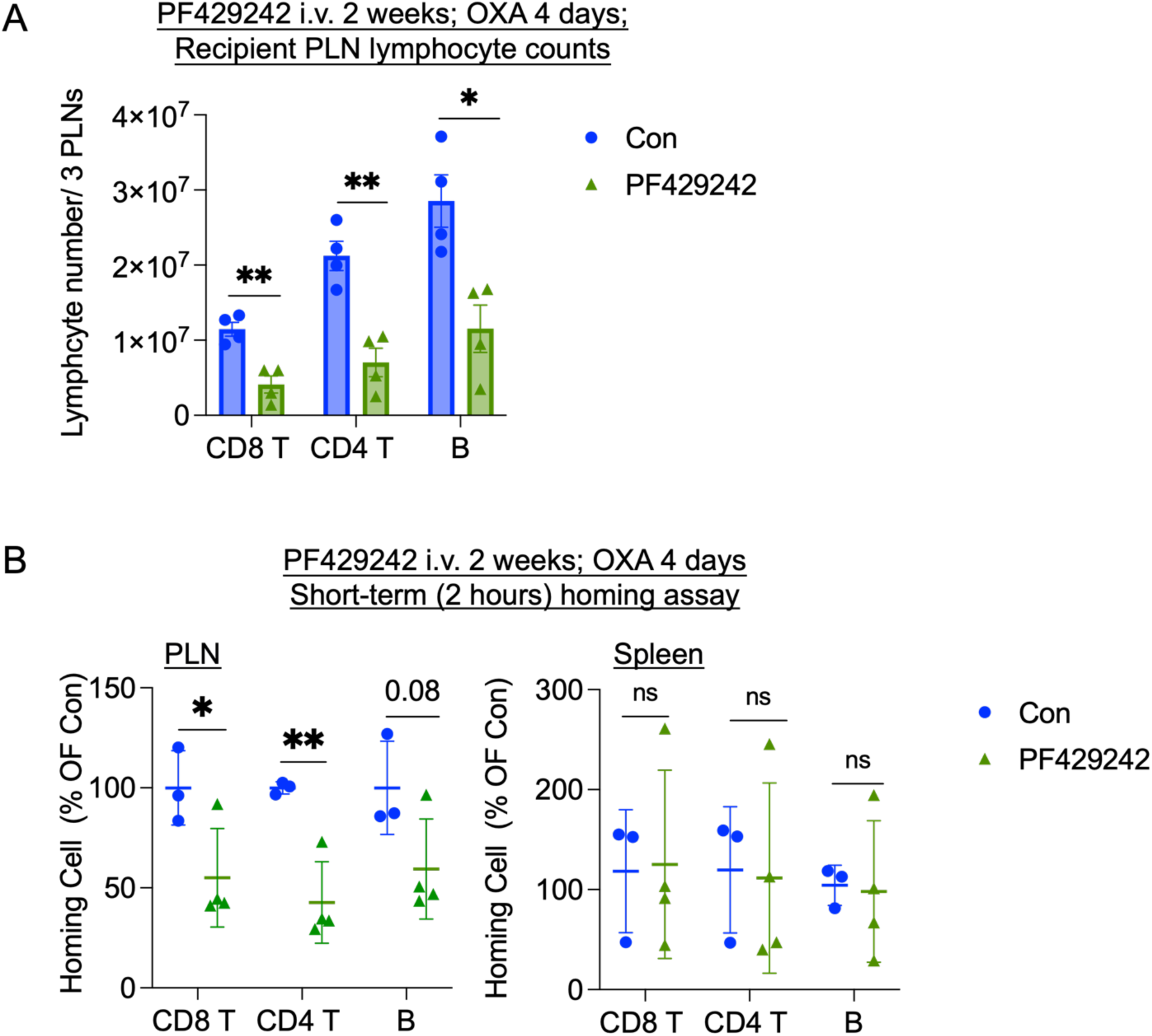
(Linked to Fig. 7, S1P Protease inhibition part). **S1P Protease Inhibition Reduces Lymphocyte Recruitment to Inflamed Lymph Nodes during Immune Activation.** Mice were topically painted with OXA on Day 1 and draining lymph nodes were analyzed on Day 4. CFSE-labeled lymphocytes were transferred i.v., and homing was assessed after 2 h. **A.** Total recipient lymphocyte counts from three pooled peripheral lymph nodes (inguinal, axillary, brachial PLN) per mouse. Each dot represents one mouse (n = 4 per group); bars indicate mean ± SEM. **B.** Localization of donor lymphocytes in PLNs and spleens of control and PF429242–treated mice after the 2-h homing assay, shown as percentage of the mean localization ratio relative to the control group. Each dot represents one mouse (n = 3 per group); bars indicate mean ± SEM. Two-tailed unpaired t-tests were used for all comparisons in panels A and B. *p < 0.05; **p < 0.01.

## Methods

### Mice

Xbp1^tm2Glm^ mice^1^ (provided by Dr. Jonathan Lin from Stanford University) were crossed with CDH5(PAC) Cre^ERT2^ mice^2^ to generate tamoxifen-inducible endothelial cell-specific Xbp1 knockout (KO) mice (Cre+ Xbp1^f/f^; iEC-Xbp1-/-). Littermates not harboring the Cre transgene were used as controls. Tamoxifen (75 mg/kg, Sigma-Aldrich, CAS # 10540-29-1) was administered via intraperitoneal (i.p.) injection four times, every other day, except for the omentum study, where tamoxifen was administered orally by gavage at the same dose. Wild-type (WT) C57BL/6J mice were obtained from the Jackson Laboratory (Bar Harbor, USA). Tamoxifen was prepared in corn oil at a concentration of 20 mg/ml by shaking overnight at 37°C in a light-protected vessel and stored at 4°C for the duration of the injections. Both male and female mice aged 6–20 weeks were used in experiments. Animals were housed under standard conditions (12 h/12 h light/dark cycle, 22 ± 2°C) with ad libitum access to food and tap water. All protocols discussed in the text and below have been approved or meet the guidelines of the accredited Department of Laboratory Animal Medicine and the Administrative Panel on Laboratory Animal Care at the VA Palo Alto Health Care System (VAPAHCS). Animals were sacrificed using approved procedures.

### Preparation of lymphoid tissue BECs and flow cytometry

Axillary, inguinal, and brachial peripheral lymph nodes (PLNs) from 10–20 adult mice were dissociated as previously described^3,4^. To minimize technical variation, male and female PLN samples were combined prior to processing. Post-sequencing, the samples were computationally separated using the **AddModuleScore** function in the Seurat package (v3.1.1), leveraging the enrichment of male-specific Y-chromosomal genes and the female-specific gene *Xist*. Endothelial cells (ECs) were isolated following the procedure outlined in the **online protocol**^3^. Briefly, PLNs were pooled in HBSS buffer and minced with scissors. The tissue was washed 2–3 times with HBSS, then resuspended in HBSS containing 0.2 mg/ml collagenase P, 0.8 mg/ml Dispase II, and 0.01 mg/ml DNase. Samples were incubated at 37 °C for 10 minutes, followed by gentle disruption through repeated pipetting with P1000 pipette tips of successively smaller bore sizes. After allowing tissue fragments to settle, released cells were transferred into ice-cold FBS (final concentration, 30%). The remaining tissue was subjected to a second round of digestion with fresh buffer. Digested cells were filtered, and hematolymphoid cells were depleted using anti-CD45 MicroBeads (Miltenyi). For Figure 1, the gene set module scores were calculated using **AddModuleScore** function in the Seurat package (v3.1.1).

Isolated mouse ECs were surface-stained with monoclonal anti-mouse antibodies: APC-conjugated anti-CD31 (390; 102410; 1:4000), peridinin chlorophyll protein–cyanine 5.5– conjugated anti-CD45 (30-F11; 103131; 1:400), peridinin chlorophyll protein–cyanine 5.5–conjugated anti-Ter-119 (TER-119 TER-119; 116227; 1:400), peridinin chlorophyll protein–cyanine 5.5–conjugated anti-CD11a (H155-78; 141007; 1:400), peridinin chlorophyll protein–cyanine 5.5–conjugated anti-CD326 (G8.8; 118219; 1:400), phycoerythrin-cyanine 7-conjugated anti-Gp38 (8.1.1; 127411; 1:100), and APC/Cyanine7-conjugated anti-Ly6c (HK1.4; 128026, 1:200) (Biolegend). Anti-PNAd (MECA79) were produced in-house from hybridomas; labelled with DyLight Antibody Labeling Kits and used at a concentration of 2mg/ml (1:400).

For additional barcoding, ECs from the PLNs of iEC-Xbp1-/- and control mice (4-5 weeks post-tamoxifen treatment) were further stained with TotalSeq™-A anti-mouse Hashtag antibodies (1:200) (BioLegend). Dead cells were excluded by staining with 4’,6-diamidino-2-phénylindole (DAPI) or the LIVE/DEAD Fixable Aqua dye (Invitrogen).

Approximately 5–10 × 10⁴ BECs (lin⁻Gp38⁻CD31⁺) were sorted into 100% fetal bovine serum using a FACS Aria (100 μm nozzle; ∼2500 cells/second). Freshly sorted cell suspensions were diluted with PBS to a final FBS concentration of ∼10%, then centrifuged at 400 × g for 5 minutes. The supernatant was carefully removed using micropipettes, and the cell pellets were resuspended in the residual volume (∼30–50 μl). Cells were counted using a hemocytometer, and the concentration was adjusted to 500–1000 cells/μl by adding PBS with 10% FBS, if necessary.

### Single-cell RNA sequencing

Single-cell gene expression was measured using the 10x Chromium v3 platform with the Chromium Single Cell 3’ Library and Gel Bead Kit v2 (10X Genomics, PN-120237), following the manufacturer’s guidelines. Male and female cohorts were processed together and computationally resolved post-sequencing. Libraries were sequenced on an Illumina NextSeq 500 using the 150-cycle High Output V2 Kit (Read 1: 26 bases, Read 2: 98 bases, Index 1: 8 bases). The Cell Ranger package (v6.0.1) was used to align high-quality reads to the mm10 transcriptome. Quality control and data analysis were performed as described previously7. Briefly, normalized log expression values were calculated using the scran package8. Imputed expression values were generated using a customized implementation of the MAGIC algorithm9 (Markov Affinity-based Graph Imputation of Cells) with optimized parameters (t = 2, k = 9, ka = 3) available at https://github.com/kbrulois/magicBatch.

Supervised cell selection was performed to exclude cells expressing non-blood endothelial gene signatures, including lymphatic endothelial cells (Prox1, Lyve1, Pdpn), pericytes (Itga7, Pdgfrb), fibroblastic reticular cells (Pdpn, Ccl19, Pdgfra), and lymphocytes (Ptprc, Cd52). The EC subsets clusters were identified based on canonical marker expression and their position in tSpace projections, yielding a total of 11 subsets7. Batch effects from technical replicates were corrected using the MNN algorithm as implemented in the batchelor package (v1.0.1) via the fastMNN function. Dimensionality reduction was performed using the UMAP algorithm. To account for cell-cycle effects, the data were split into dividing and resting cells, and the fastMNN function was used to align dividing cells with their resting counterparts. All subsets were represented by cells from both male and female mice across each condition. Heatmaps were created using the ComplexHeatmap package15, Cell subsets were clustered based on their mean expression profiles using ClustGeo and vegan packages. Violin plots were generated using the ggplot2 package, where y-axis units represent log-transformed normalized counts following imputation. Differential gene expression (DEG) analyses were performed using two-sided Student’s t-tests. For comparisons between endothelial cell (EC) subsets (e.g., blood endothelial subset^4^ vs intestinal goblet cell subset^5^), as well as between wild-type (WT) and Xbp1-deficient ECs, statistical analyses were conducted independently within each defined cell subset. Genes were considered differentially expressed if they met the following criteria: average log₂ fold change (log₂FC) ≥ 0.0586 and false discovery rate (FDR) ≤ 9.94 × 10⁻⁴⁴. The least significant DEG identified corresponded to a –log₁₀(P) value of 47.30. Complete DEG lists for EC subset comparisons are provided in Table S1, and DEG results for WT versus Xbp1-deficient ECs across individual EC subsets are provided in Table S7.

### Cell lines

bEnd.3 and HEK293T cells were obtained from ATCC (CRL-2299 and CRL-3216 respectively). HEK293T cells were grown in Dulbecco’s modified Eagle’s medium (DMEM) containing high glucose and L-pyruvate and supplemented with 10% heat-inactivated fetal bovine serum, at 37 °C in 5%CO2. bEnd.3 and DN-MAML bEnd.3 cells were grown in the same media supplemented with penicillin-streptomycin^6^. Stable bEnd.3 cell transfectants overexpressing Fut7 promoter reporter were generated by transfection and selected with puromycin (1ug/ml).

### Plasmid constructs

*Chst4* enhancer-promoter and *Fut7* promoter fragments were PCR-amplified fragment from mouse genomic cDNA and cloned into pGL4.23[*luc2P*/Puro] and pGL4.22[*luc2P*/Puro] respectively to generate the luciferase (LUC) reporter constructs. pGL4.23[*luc2P*/Puro] and pGL4.22 [*luc2P*/Puro] vectors were obtained from Promega. *Pgm3*, *Nans*, *Galt, Scl35c1 and Golph3* enhancer-promoter_LUC reporter constructs were made by Vector Builder custom service. Control Renilla (Ren) Luciferase vector is from Promega.

For co-transfection studies, mammalian expression vectors un-spliced Xbp1 (Xbp1), spliced Xbp1 (Xbp1s), IRE1 dominant negative kinase mutant (IRE1-Mut), IRE1, Creb3l2 were commercially obtained. Constructs information is provided in Table S7.

### Transient transfection

For luciferase reporter assays, 1.5 × 10⁵ HEK293T cells were plated in 24-well plates and transfected upon reaching ∼70–80% confluency. Respective luciferase reporters, transcription factors, and a Renilla luciferase control vector (Promega) were co-transfected using the Lipofectamine 3000 system (Thermo Fisher Scientific) according to the manufacturer’s instructions. Forty-eight hours post-transfection, cells were lysed using Lysis Reagent (Promega). Firefly and Renilla luminescence were measured using the Dual-Glo® Luciferase Assay System and read with a Turner Biosystems 20/20n luminometer. Luciferase activity was quantified as the ratio of Firefly to Renilla luminescence.

### Western blotting

Cells were lysed in RIPA buffer supplemented with protease inhibitors, and protein concentrations were determined by BCA assay. Equal amounts of protein were separated by SDS–PAGE and transferred onto PVDF membranes. Membranes were blocked and incubated with primary antibodies overnight at 4°C, followed by incubation with IRDye 680LT Goat anti-Rabbit IgG and IRDye 680LT Goat anti-Mouse IgG secondary antibodies (LI-COR Biosciences). Signals were detected using a LI-COR Odyssey imaging system.

### RT-PCR

Total RNA was isolated using the RNeasy Mini Kit (Qiagen) and reverse transcribed into cDNA using the High-Capacity cDNA Reverse Transcription Kit (Applied Biosystems) according to the manufacturer’s instructions. Quantitative PCR was performed using SYBR Green Master Mix on a real-time PCR system. Gene expression was normalized to Gapdh and calculated using the 2^−ΔΔCt method.

### Electrophoretic mobility shift assay (EMSA)

EMSAs were performed as previously described^6,7^. Double-stranded, fluorescence-labeled, and unlabeled probes were generated by annealing sense and antisense oligonucleotides (Sequences shown in Supplemental Fig. 4). Each binding reaction contained 1–2 pmol of labeled probe and 50 μg of nuclear extracts from 293T cells expressing recombinant spliced Xbp1 or CREB3L2. Gel shift reactions were conducted at 4 °C for 1 hour in 1.875 mg poly(d1-dC),100 mM NaCI, 10 mM Tris, 1 mM DTT, 1 mM EDTA, Protease inhibitors (1:1000), 1% BSA. In reactions with cold competitors, 50-fold molar excess of (”cold”) competitor unlabeled probes were added. Protein–probe complexes were separated on 5% Mini-PROTEAN® TBE gels (Bio-Rad) run in 4% TBE buffer (Bio-Rad). Gels were imaged using the Odyssey Imaging System (LI-COR).

### Pharmacological inhibition of IRE1α-Xbp1 and S1P Protease

For the pharmacological inhibition of IRE1α-Xbp1, WT C57BL/6J mice were injected with STF-083010 (30 mg/kg, i.p.) or DMSO, both given in 16% (vol/vol) Cremophor EL (Sigma-Aldrich) saline solution via i.p. injections every other day for four does^8^. For inhibiting active Creb3l2, WT C57BL/6J mice were administrated with PF-429242 (20 mg/kg, i.v.) or saline solution daily for two weeks^9,10^.

### Imaging

Peripheral lymph nodes (PLNs), omentum, and ear skin were imaged following retroorbital injection of fluorescently labeled antibodies. Antibodies (10–25 μg) were administered 20–30 minutes prior to sacrifice and tissue collection. To visualize the overall vasculature, tissues were gently compressed to a thickness of approximately 35–50 μm on a glass slide. Imaging was performed using a Zeiss LSM 880 laser scanning microscope and a Nikon ECLIPSE Ti2 microscope.

### Image quantification

Image analysis and parameter quantification were performed using *Imaris* 9.1. Parameters assessed included the total number of individual cell-sized particles expressing specific fluorescence signals and the volumetric surface area, representing the external surface of an isosurface. CFSE-stained cells were identified and counted using the **“Spots”** module. Volumetric measurements for **Ly6C⁺**, **MECA-79⁺**, and **CD31⁺** structures were obtained by generating 3D isosurfaces based on fluorescence detection with the **“Surfaces”** module. Total vascular surface area was measured from the **CD31⁺** isosurface. Surface statistics were generated automatically by the software and exported as CSV files for downstream quantification and statistical analysis. For analysis of cells within high endothelial venules (HEVs), a mask was applied to exclude voxels inside the region of interest (ROI), enabling quantification of cell distributions external to the surface objects using the **“Spots”** module.

### Short-term homing and short-term diapedesis assays

Donor splenocytes were isolated and were labeled with 2.5 μM CFSE (Invitrogen) or CellTrace Yellow dye (Invitrogen) for 15 min at 37°C in RPMI without fetal bovine serum (FBS). A total of 50–100 × 10⁶ donor cells were transferred into recipient mice (wild-type mice, wild-type mice treated with inhibitors, and iEC-Xbp1⁻/⁻ mice) via retroorbital injection.

For the homing assay, peripheral lymph nodes (PLNs; pooled axillary, inguinal, and brachial nodes; average of 3 PLNs per mouse), mesenteric lymph nodes (MLNs), Peyer’s patches (PPs; average of 3 per mouse), and spleens were harvested 2 hours after donor cell injection. Single-cell suspensions were surface-stained for flow cytometry using the following monoclonal antibodies: PC-conjugated anti-CD4 (RM4-5; 100516; 1:200), BV421-conjugated anti-CD8β (YTS156.7.7; 126629; 1:200), PE-Cy7-conjugated anti-CD3 (145-2C11; 100320; 1:200), PE-conjugated anti-CD19 (6D5; 115508; 1:200), and FITC-conjugated anti-IgD (11.26c.2a; 11-5993-82; 1:100). Appropriate isotype controls were included. The Fc receptors were blocked using Rat Serum and anti-CD16/32 prior to the staining. Dead cells were excluded using Fixable Aqua Dead Cell Stain (Invitrogen), and counting beads were added to determine absolute cell numbers. T and B cell homing efficiencies were calculated as the ratio of recovered cells to input cells and normalized relative to the mean ratio of the wild-type control group.

For the diapedesis assay, tissues were collected 30 minutes after donor cell injection. Fluorescently labeled antibodies (10–25 μg) were administered retroorbitally 15 minutes prior to sacrifice. To visualize the overall vasculature, tissues were gently compressed to a thickness of approximately 35–50 μm on a glass slide. Imaging was performed using an LSM 880 laser scanning microscope (Zeiss) and an ECLIPSE Ti2 microscope (Nikon).

### Delayed-Type Hypersensitivity (DTH) Assay

The DTH reaction in WT and iEC-Xbp1⁻/⁻ mice was induced as previously described^11^. Briefly, the DTH response was triggered through repeated epicutaneous application of 2,4-dinitrofluorobenzene (DNFB; Sigma-Aldrich). Primary immunization was achieved by applying 0.5% DNFB (50 μL in acetone: olive oil, 4:1) to bare abdominal skin on Day 1. On Day 7, mice were challenged by applying 0.5% DNFB (20 μL) to both sides of the left ear to elicit a secondary DTH response, while the right ear received 20 μL of solvent (acetone: olive oil, 4:1) as a control. 24 hours post-challenge, the thickness of both the left and right ears was measured with Ames calipers. The inflamed skin along with the draining lymph nodes (auricular and inguinal LNs) was harvested for FACS and H&E analysis.

### Preparation of lymphocytes from skin and flow cytometry

Skin-infiltrating T cells were released from the connective tissue after separating the epidermal sheets as described^11–13^. T cells were surface-stained with monoclonal anti-mouse antibodies for flow cytometry, including BV421-conjugated anti-CD4 (RM4-5, 100563, 1:200); PE-Cy7-conjugated anti-CD3 (145-2C11; 100320; 1:200); BV650-conjugated anti-CD45Rb (C363-16A; 740449; 1:150); APC-conjugated anti-CD44 (IM7; 560567; 1:2000). The Fc receptors were blocked using Rat Serum and anti-CD16/32 prior to the staining. Fixable Aqua dye was applied for live/dead cell staining, and counting beads were included to calculate absolute cell numbers.

### Transmission Electron Microscopy

One inguinal lymph node from each wild-type or iEC-Xbp1⁻/⁻ mice was isolated and washed with cold PBS. The whole lymph nodes were fixed in Karnovsky’s fixative: 2% Glutaraldehyde (EMS Cat# 16000) and 4% formaldehyde (EMS Cat# 15700) in 0.1M Sodium Cacodylate (EMS Cat# 12300) pH 7.4 for 1 hr. then placed on ice 24 hours. The fix was replaced with cold/aqueous 1% Osmium tetroxide (EMS Cat# 19100) and were then allowed to warm to Room Temperature (RT) for 2 hrs rotating in a hood, washed 3X with ultrafiltered water, then en bloc stained in 1% Uranyl Acetate at RT 2hrs while rotating. Samples were then dehydrated in a series of ethanol washes for 30 minutes each @ RT beginning at 50%, 70% EtOH then moved to 4°C overnight. They were place in cold 95% EtOH and allowed to warm to RT, changed to 100% 2X, then Propylene Oxide (PO) for 15 min. Samples are infiltrated with EMbed-812 resin (EMS Cat#14120) mixed 1:2, 1:1, and 2:1 with PO for 2 hrs each with leaving samples in 2:1 resin to PO overnight rotating at RT in the hood. The samples are then placed into EMbed-812 for 2 to 4 hours then placed into molds w/labels and fresh resin, orientated and placed into 65° C oven overnight.

Sections were taken around 80nm using an UC7 (Leica, Wetzlar, Germany) picked up on formvar/Carbon coated 100-mesh copper grids, stained for 40seconds in 3.5% Uranyl Acetate in 50% Acetone followed by staining in Sato’s Lead Citrate for 2 minutes. Observed in the JEOL JEM-1400 120kV. Images were taken using a Gatan OneView 4k X 4k digital camera.

### Electron microscopy sampling and quantification

For each genotype, 3 animals were analyzed. From each animal, ≥2 tissue blocks were prepared, and from each block, ≥2 grids (10–12 sections per grid) were randomly selected. Images were acquired from a random start field on each section at predefined magnifications. Ultrastructural features were quantified in ImageJ, including autophagic vacuoles, osmiophilic cytoplasm, and distorted ER, which were measured as percent area per micrograph, as well as mitophagy events and degraded mitochondria, which were quantified as the average number per micrograph. Data are presented as mean ± SEM, with n denoting the number of micrographs analyzed.

### Lymph node and Ear immunization

Mice underwent a cutaneous immune challenge by applying 20 μL of 3–5% 4-Ethoxymethylene-2-phenyl-2-oxazolin-5-one (Sigma–Aldrich) dissolved in a 1:2 acetone: oil mixture. For lymph node analysis, the solution was applied to bare abdominal skin, while for skin analysis, it was applied to both sides of the ears. Peripheral lymph nodes (axillary, brachial, and inguinal) were collected on Day 4, and ear tissues were harvested on Day 10 following inflammation induction for imaging purposes.

### HEV induction in Omentum

Six-week-old iEC-Xbp1⁻/⁻ and control mice were administered tamoxifen (75 mg/kg) orally via gavage every other day for a total of four doses. Four to five weeks after tamoxifen treatment, mice received LPS (5 mg/kg) intraperitoneally twice over a 72-hour period, followed by omentum vasculature imaging.

### De novo TF motif discovery

ATAC-seq on purified HECs and Capillary ECs (CapECs) were performed essentially as previously described^14^. HEC-enriched ATAC-seq peaks were identified by differential accessibility analysis between HECs and CapECs. Peaks uniquely enriched in HECs (HEC>CapEC) were used for downstream motif analysis.

Evolutionarily conserved regulatory regions associated with HEV-enriched genes were defined as the gene body ±5 kb and intersected with conserved genomic elements scoring >300 in the UCSC Genome Browser phastCons Placental Elements track, which aligns 60 vertebrate genomes and reports nucleotide-level conservation^15,16^. All conserved regions of HEV genes are provided as a custom track in a UCSC Genome Browser session: https://genome.ucsc.edu/s/m.xiang/HEV_genes_conserved.

De novo transcription factor (TF) motif discovery was performed on HEC-enriched ATAC-seq peaks as well as on conserved regulatory regions of HEV-enriched genes in the mouse *mm10* genome using HOMER^17^, applying default background models and standard significance thresholds.

### TF regulation analysis

List of HEC and GC co-upregulated DEGs, HEC- and GC- enriched genes were analyzed to identify candidate upstream TF regulators using Enrichr^18–20^. Results of TF regulators from select databases are shown. Detailed analysis, including the gene lists used, is provided in Data 2-4.

### Comparative Genomics and Phylogenomic Analysis

Comparative genomics and phylogenetic footprinting were performed using the UCSC Genome Browser to identify evolutionarily constrained regions across candidate genes, including promoter regions, 5′ and 3′ untranslated regions (UTRs), and intronic sequences. Analyses were based on the 58 eutherian mammals Multiz Alignment and PhyloP Basewise Conservation tracks to evaluate sequence conservation. To identify putative transcriptional regulatory elements, we utilized the TFBS Conserved track and the JASPAR TFBS database, focusing on conserved binding motifs for XBP1 and CREB3L2 within the identified regulatory modules.

### Data and statistical analysis

Results are expressed as mean ± SEM and individual data points represent biological replicates. Statistics were calculated using GraphPad Prism, with differences among means tested for statistical significance as indicated in the corresponding figure legend. Statistical significance is indicated by the following: *P ≤ 0.05, **P ≤ 0.01, ***P ≤ 0.001 and ****P≤0.0001. No statistical significance is denoted as ‘na’.

