## Supplementary material for "Coordinated Transcriptional Networks Program Organelle Expansion and Metabolic Flows for High Endothelial Morphology and Function": https://drive.google.com/drive/folders/16LrJoVh-ugeEoXfNzdhAxGWC4Km_MaxT?usp=sharing

**Data S1:**

Immunohistochemical profiling from the Human Protein Atlas validated these transcriptomic predictions at the protein level, revealing selective enrichment in HECs of enzymes supporting metabolic flux (PASS2, PGM3, PMM2, CMAS), nucleotide sugar transporters involved in endomembrane trafficking (SLC35B2, SLC35C1), core components of the PNAd biosynthetic machinery (PARM1, GALNT1, CHST4), regulators of secretory organelle homeostasis and vesicular transport (EDEM2, STX5A, KDELR2, GOLPH3), and unfolded protein response factors (SLC35B1, HSPA5) (Data S1). Representative immunohistochemical images from the Human Protein Atlas illustrating this protein-level enrichment are shown below.

PAPSS2 – Sulfation Pathway

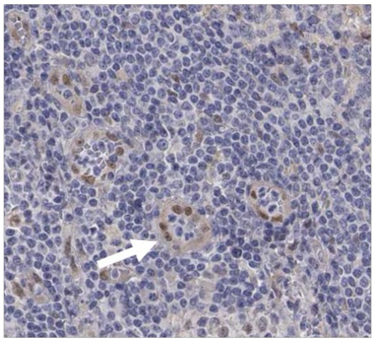

PGM3 – HBP Pathway for synthesizing GlcNAC

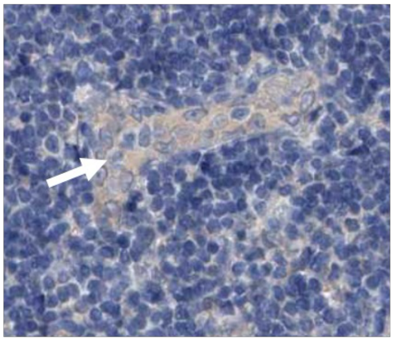

PMM2 – Fucosylation Pathway

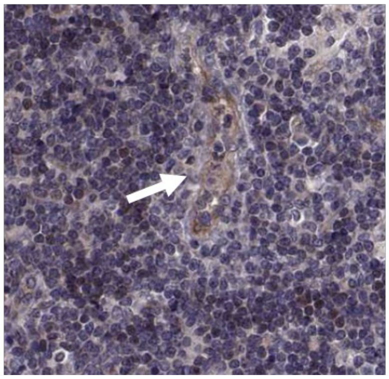

CMAS (nuclear staining) –Sialylation Pathway

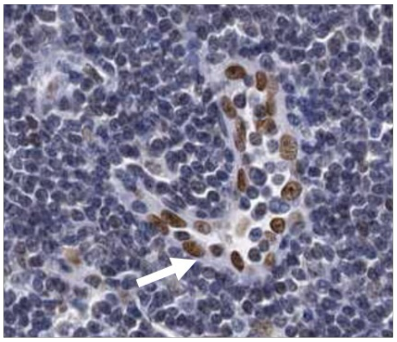

Slc35b2 for transporting PAPS, a universal sulfate donor

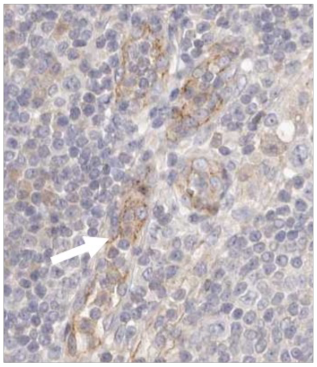

Slc35c1 for transporting GDP-Fucose

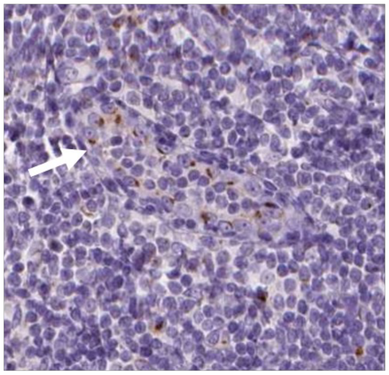

Genes for PNAd assembly

PARM1

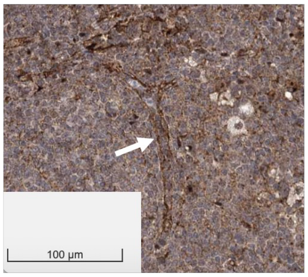


Galnt1

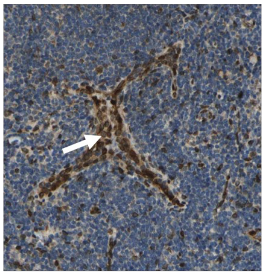

Chst4

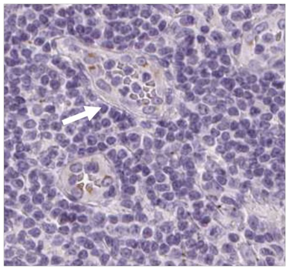

Slc35b1, importing ATP from cytosol into ER lumen to support protein folding

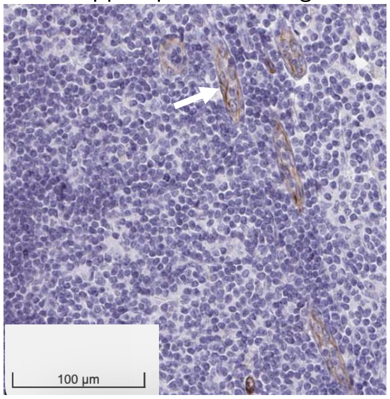


HSPA5 – a regulator for UPR

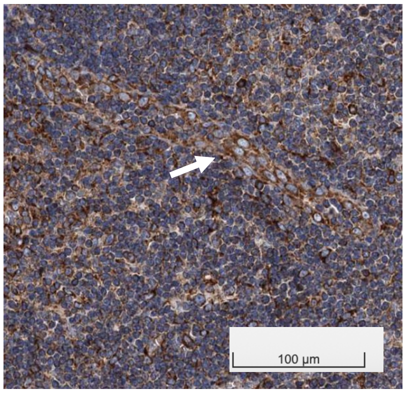


### Organelle Proteins

Stx5a

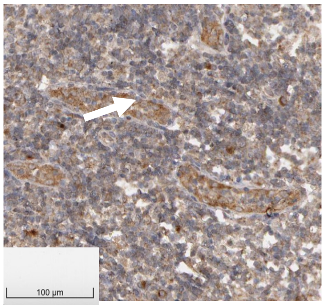


KDELR2

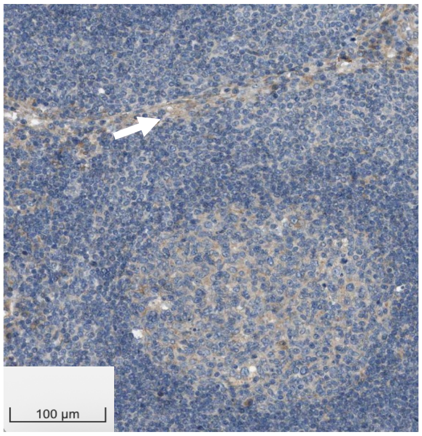


Golph3

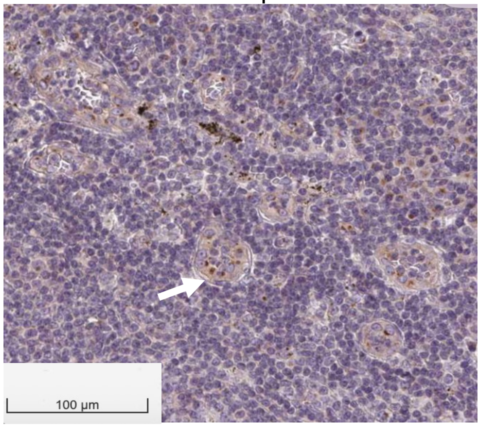


EDEM2

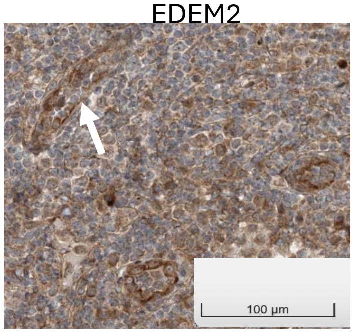
