## Supplementary material for "Coordinated Transcriptional Networks Program Organelle Expansion and Metabolic Flows for High Endothelial Morphology and Function": https://drive.google.com/drive/folders/16LrJoVh-ugeEoXfNzdhAxGWC4Km_MaxT?usp=sharing

**Data S2:**
Using phylogenetic DNA footprinting, we identified evolutionarily conserved binding motifs for XBP1 and CREB3L1/2 within cis-regulatory modules of genes implicated in coordinating inter-organelle metabolic fluxes that underpin the biosynthesis of sulfated mucin-type O-glycans and the maintenance of organelle architecture. These regulatory elements were enriched in lineage- and state-specific expression programs of high endothelial cells (HECs) and intestinal goblet cells (GCs), supporting a shared transcriptional logic that governs secretory cell specialization through organelle adaptation and glycan remodeling.

**Inter-Organelle Metabolic Fluxes for Biosynthesis of Sulfated Mucin-Type O-Glycans:**
*Mucins:*
Glycam1
Cd34
Parm1
Podxl
Cd24a
Muc2
*Metabolic Fluxes for PAPS, UDP-Gal, UDP-GlcNAc, UDP-GalNAc, GDP-Fus, CMP-Neu5Ac*
Papss1
Papss2
Galk2
Galt
Hk1
Hk2
Gpi1
Gfpt1
Gnpnt1
Pgm3
Uap1
Gale
Mpi
Pmm1
Pmm2
Gmppa
Gmppb
Gmds
Gfus
Gne
Nans
Nanp
Cmas
Cmah
*Genes for the biosynthesis of Dolichol-diphosphate GlcNAc2Man9Glc3, a precursor for N-linked glycosylation*
Dpagt1
Alg2
Alg6
Alg9
*Mitochondria for producing Acetyl-CoA*
Cpt1a
Acaa2
Slc25a1
Acyl
*Transporters*
Slc2a1
Slc1a5
Slc26a2
Slc35b2
Slc35b3
Slc35a2
Slc35c1
Slc35a3

Slc35a5
Slc35a1
Slc33a1
*Sulfated Mucin-Type O-Glycan Biosynthesis in Golgi Stacks:*
Galnt3
Galnt5
Galnt12
C1galt1
C1galt1c1
Gcnt1
Gcnt3
B3gnt3
B3gnt5
B3gnt6
B4galt6
St3gal4
St3gal6
St6gal1
St6galnac6
Chst2
Chst4
Chst5
Fut1
Fut2
Fut4
Fut7

**Genes for Organellar Architecture:**
*ER-targeted translation, translocation, and N-glycosylation*
Ssr1
Ssr4
Sec61b
Spcs2
Rpn1
Stt3a
Stt3b
Ost4
Ddost
Tmem258
Rps3
Rpl10

*Protein Folding*
Man1a
Canx
Calr
Hspa5
Dnajb9
Agr2
Pdia3
Pdia6
Fkbp2

*COP II trafficking*
Mia3
Lman1
Copb2
Sar1b
Sec13
Sec16a
Sec23ip
Sec24d

Uso1
Stx5a
Bet1

*COP I Trafficking*
Kdelr1
Kdelr2
Kderlr3
Scyl1
Surf4
Arf3
Copb1
Cope

*ERAD*
Edem2
Derl1
Os9
Erlec1
Vcp
Syvn1

*UPR pathways*
Slc35b1
Hspa5
Agr2
Pdia3
Pdia5
Pdia6
Ern1
Ern2
Xbp1
Creb3l1
Creb3l2
Mbtps1
Mbtps2

*Golgi-resident Proteins*
Golga2
Golgb1
Golph3

***Lineage-specific and State-specific genes in HECs and GCs***
Ltbr
Tnfrsf11a (Rank)
Nfkb2
Relb
Bcl3
Bhlha15 (mist1)
Nod1
Nod2
Atg16l
Il13ra1
Nr2f2
Spdef

### Mucins:

#### *Glycam1*

```
                                   Creb3l2-like
hGlycam1        TAATTAACACCCGCCCTGTTAATGATTAAATCTAATGACTTTGACTTGCGTGTGGGAGGA
chimp           TAATTAACACCCGCCCTGTTAATGATTAAATCTAATGACTTTGACTTGCCTGTGGGAGGA
rhesus          TAATTAACACCCGCCCTGTTAATGATTAAATCTAATGACTTTGACTTGCGGGTGGGAGGA
mouse           TAATTAACACAAACCCTGTTAATGATTAAATCTAATGACTTTGACTTAGAGGTGGGATGA
rat             TAATTAACACACACCCTGTTAATGATTAAATCTAATGACTTTGACTTAGAGGTGGGATGA
                **********...**********************************.   ****** **
                    NF-kB
hGlycam1        TGGAAAATTCCCCAGCCACAGGACGGGATGAGGGTAGGTTTCCAGGAATGGGAGGGCAAG
chimp           TGGAAAATTCCCCAGCCACAGGACGGGATGAGGGTAGGTTTCCAGGAATGGGAGGGCAAG
rhesus          TGGAAAATTCCCCAGTCACAGGACAGGATGAGGGTAGGTTTCCAGGAATGGGAGGTCAAG
mouse           TGGAAAATTCCCTAGCCACAGGATAGAATGAGGGTAGGTTACCAGGAGTAGGGAGGCCAA
rat             TGGAAAATTCCCTAGCCACAGGACCGAATGAGGGTAGGTTACCAGGAGTGGGGAGGCCAA
                ************ ** *******  *.*************:******.*.**..* *.*.
```

---

#### *Parm1*

```
                                                            Creb3l2
hParm1          TTCAGTTCT----TAATTTCTGCCCATTCCAGGGCTTTATAACATGAGTCACACCCAGTC
chimp           TTCAGTTCT----TAATTTCTGCCCATTCCAGGGCTTTATAACATGAGTCACACCCAGTC
rhesus          TCCAGTTC----TTAATTTCTGCCCATTCCAGGGCTTTATCACATGAGTCACACCCACTC
mouse           TCCAGTTCTCTTTTAAGGTCTGCCCATCCCAGAGCTTTATGGCATGAGTCACACCTGGTC
rat             TCCAGTTCTCTTTTAAGGTCTGCCCATCCCAGAGCTTTATGGCATGAGTCACACCAGGTC
                * ******     ***  ********* ****.******* .************* . **
                                      ETS                      ETS
hParm1          TCTAACTCACAAATCCCAATGAGGAAGAAATGAAACGCCAGCCACCTTCCCCATGTCTCC
chimp           TCTAACTCACAAATCCCAATGAGGAAGAAATGAAACGCCAGCCACCTTCCCCATGTCTCC
rhesus          TCTAACTTACAAATCCCAATGAGGAAGAAATGAAATGCGAGCCACCTTCCCCGTGTCTCC
mouse           TGAGAATAATGGGCCCTGAGGCGGAAGAACTAAAATGCCAGCAAACTTCCCCATAGCCCT
rat             TGTGAATAATGGACCGTGAGGCGGAAGAAATAAAATGCCAGCAAACTTCCCACTG-CCCT
                * :.*.* * ... *  .* *.*******.*.*** ** ***.*.******. *. * *
```

---

#### *Muc2:*

```
                                                     Creb3L2
hMuc2promoter      CCAGGGAGCCATAAAGAGATGACCTCCGATAACCTGAATCAATATTTCCCCATTGGGGCT
chimp              CCAGGGAGCCATAAAGAGATGACCTCCGATAACCTGAATCAATATTTCCCCACTGGGGCT
rhesus             CCAGGGAGCCATAAAGAGATGACCTCTGATAACCTGAATCAATATTTCCCCACTGGGGCT
mMuc2promoter      CCAGGGAGTCATATAAAGATAAACTCAGATAACCTGAATCAATATTTCCTCTCTGGGACC
rat                CCAGGGAGTCATATAAAGATAAACTCAGATAACTTGAATCAATATTTCCTCTCTGGGACC
                   ******** ****:*.****.*.*** ****** *************** *: ****.*
                                E-box                             KLF
hMuc2promoter      CGGG---CCCCCGCAGCTGTCTTCTTGATCATCTGGCAGATGCCACACCCACCCTTG-GC
chimp              CAGG---CCCCCGCAGCTGTCTTCTTGATCATCTGGCAGATGCCACACCCACCCTTG-GC
rhesus             CGGG---CCCCTGCAGCTGTCTTCTTGATCATCTGGCAAATGCCACACCCACCCCTTGAC
mMuc2promoter      CATGGAGCCCCCACAGCTGTTTTCCTGATAACTTGGCAAATGCCCCACCCACCCTTCGCA
rat                TGTGGAGCCCCCACAGCTGTCTTCCTGATAACTCGGCAAATGCCCCACCCACCCTTCGCA
                    . *   **** .******* *** ****.*   ****.*****.********* *   .
```

---

#### ***Metabolic Fluxes for PAPS, UDP-Gal, UDP-GlcNAc, UDP-GalNAc, GDP-Fus, CMP-Neu5Ac***

#### *Papss1:*

```
                                                  GATA               E-box
hPapss1         TATAGACTATAAAATCAGGAACATCCTGT----TGATAACCAAATAGGCTTAACAGCTGC
chimp           TATAGACTATAAAATCAGGAATATCCTGT----TGATAACCAAATAGGCTTAACAGCTGC
rhesus          TACAGACTGGAAAATCAGGAACATCCTGT----TGATAACCAAATAGGCTTAACAGCTGC
mouse           TATAGACTGGAAAATGAGGAACATCCTGTTGATAGATAAGCGCGCAGCCTCAGCAGCTGT
rat             TATAGACTAGAAAACGAGGAACATCCTGT----TGATAACCTCGCAGCCTTAGCAGCTGT
                ** *****. ****  ***** *******     ***** * .. ** ** *.******
                                      XBP1   (ERSEII-like)   NF-YA/B
hPapss1         CCATGGTTCTGAAAAGCATTTGTGCTAAGGAAACAAATCTGGTACCAATGCCTTCTAAAT
chimp           CCATGGTTCTGAAAAGCATGTGTGCTAAGGAAACAAATCTGGTACCAATGCCTTCTAAAT
```

```
rhesus      TCATGGTTCTGAAAAGCATTTGTGCTAAGGAAACAAATCTGGTACCAATGCCTTCTAAAT
mouse       GCATTGTTCTGAAGTGCATATGTGCTATGGAACCAAATCTGGTACCAATGCCTGCTAAAC
rat         TCATTGTTCTGAAGCGCATCTGTGCTAAGGAAACAAATCTGGGACCAATGCCTTCTAAAC
             *** ********. **** *******:****.********* ********** *****

hPapss1     GACCTACAGAAAATGATTAAAGGAAGAGTCGTTGTTTTTATTTTATTATTATTAGTTGGC
chimp       GACCTACAGAAAATGATTAAAGGAAGAGTCGTTGTTTTTATTTTATTATTATTAGTTGGC
rhesus      GACCTACAGAAAACGATTAAAGGAAGAGTCATTGTTTTTATTTTATTATTATTAGCTGGC
mouse       AACC--CAGAGAGCCATTAAAG-GAGAATCACTGTTTGTATTTTATTATGACAATTTGGC
rat         AACC--CAGCAG-CGATTAAAG-GAGAATCGCTGTTTGTATTTTATTATGACAATTTGGC
            .***  ***...   ******* .***.**. ***** *********** * :*  ****
                                    Creb3l2   NFAT
hPapss1     AAATAAGCATTACTGACAGACTTTTTCCTGATGTCACTGGAAAAGCTACAAAAATGAGAG
chimp       AAATAAGCATTACTGACAGACTTTTTCCTGATGTCACTGGAAAAGCTACAAAAATGAGAG
rhesus      AAATAAGCATTACTGACAGACTTTTTCCTGATGTCACTGGAAAAGCTACAAAAATGAGG-
mouse       AAGTATGTCTTATTGGCAGCCTTGTTCTTGATGTCACTGGAAAAACTAGAAAAACGAGGG
rat         AAATATGTGTTGTTGGCAGCCTTGTTCCTGATGTCACTGGAAAAACTAGAAAAATGAGGG
            **.**:*  **. **.***.*** *** ****************.*** ***** ***.
```

***Papss2:***

```
                                              Relb/NFkB2-p52
hPAPSS2     CTGGCGGAGCGCGCGCCCGCGAGTAGGGGCCGGGCCGGGGACCCCGCCTAGGCGG-CGGC
chimp       CTGGCGGAGCGCGCGCCCGCGAGTAGGGGCCGGGCCGGGGACCCCGCCCAGGCGAGCGGC
rhesus      CTGGCGGAGCGCGCGCCCGCGAGTAGAGGCCGGGCCGGGGACCCCGCCCAGGCGG-CGGC
mouse.      CTGGCGGAGCGCCG-------------------GCGGGGAACCCCGCCCTGGCGG----C
rat         CTGGCGGAGCGCCG-------------------GCGGGGAACCCCGCCCAGGCGG----C
            ************                     ** ***.******** :****.    *
                 Relb/NFKB2-p52
hPAPSS2     GGCCGGGTCCCCAAGGCTGGGCGCTGCTTGCGGAACCGACGGGGCGGAGAGGAGCGTGGC
chimp       GGCCGGGTCCCCAAGGCTGGGCGCTGCTTGCGGAACCGACGGGGCGGAGAGGAGCGTGGC
rhesus      GGCCGGGTCCCCGAGGTTGATCGCTGCTTGCGGAACCGACGGGGCGGAGAGGAGCGTGGC
mouse.      TGCCGGGTCCCCGGGGCTGGGCGCTGCTGGCGGAGCCGACGGGGCGGAGAGGAGCGCGGC
rat.        TGCCGGGTCCCCGGGGCTGGGCGCTGCTGGCGGAGCCGACGGGGCGGAGAGGAGTGCGAC
             ***********..** **. ******* *****.****************** * *.*
                                        Coup/ETS    XBP1/Creb3l2
hPAPSS2     GGGAGGAGGAGTAGGAGAAGGGGGCTGGTCAAGGGAAGTGCGACGTGTCTGCGGAGCCTT
chimp       GGGAGGAGGAGTAGGAGAAGGGGGCTGGTCAAGGGAAGTGCGACGTGTCTGCGGAGCCTT
rhesus      GGGAGGAGGAGTAGGAGAAGGGGGCTGGTCAAGGGAAGTGCGACGTGTCTGCGGAGCCTT
mouse       GGGAGGAGGAGTAGGAGAAGGGGGCCGGTCAAGGGAAGTGCGACGTGTCTGAGGAGCCTT
rat.        GGGAGGAGGAGTAGGAGAAGGGGGCCGGTCAAGGGAAGTGCGACGTGTCTGAGGAGCCTT
            ************************* ************************.********
```

***Galk2:***

```
                                                        NFAT
hGalk2      AGCCAAGGAGACATAAATAACCATAAACCACAAAATACAAAGATGGAAAGTGGATAG-AG
chimp       AGCCAAGGAGACATAAATAACCATAAACCACAAAATACAAAGATGGAAAGTGGATAG-AG
rhesus      AACCAAGGAGACATAAATAACCATAAACCACAAAATACAGAGATGGAAAGTGGATAG-AG
mouse       AACCTG------------------AAACCACAAAACACAAAGATGGAAAGTGGACGAGAG
rat         AACCCG------------------AAACCACAAAACACAAAGATGGAAAGTGGACAG-AG
            *.**  .                 *********** ***.**************  .. **
                      Creb3l2
hGalk2      GAATGTTACATGACTTAGGCGAATGGAGAAAGGTCAGAGCCAAGTGCCTGCGGAGGGAGT
chimp       GAATGTTACATGACTTAGGCGAATGGAGAAAGGTCAGAGCCAAGTGCCTGCGGAGGGAGT
rhesus      GAATGGTACATGACTTAGCAGAATGGAGAAAGGTCAGAGCCAAGTGCCTGTGGAGGGAGT
mouse       AACGGTTGCATGACTTAGCAGAAGGAAGAAAGGTCAAAGCTAAGTGCCATGGGGAGGAGA
rat         AGCGGCT-CATGACTTAGCAGAAGGAAGAAAGGTCAAAGCTAAGTGCCAGG-AGAGCAGA
            ...  * * ********** .*** *.**********.*** *******:    ...* **:
                              Creb3l2
hGalk2      TGCAGATGGAAAGGCAGCTGACTCATCCCACAGCTCTCTGGCAAGGCTTAGGAAGAAAGC
chimp       TGCAGATGGAAAGGCAGCTGACTCATCCCACAGCTCTCTGGCAAGGCTTAGGAAGAAAGC
rhesus      TGTAGATGGAAAGGCAGCTGACTCATCCCACAGCTCTCTGGCAAGCCTTAGGAAGAAAGC
mouse       CATAGATAGAAG-CAAGCTGACTCATCCTAGT---CTCAGGGAA-GGATAGAAAGAAAGC
rat         CATAGACAGAAG-CAAGCTGACTCATCCTAGT---CTCAAG-----GATAGAAAGAAAGC
             . *** .***.  .************* * :   ***:.*      :***.********
```

------

***Galt:***

```
                                                                      KLF
hGaltPromoter    AGGATCATCAATGTCGGCCCCCAGGG------TTCACAGCTGTTCTGAGCCCCGCCCCCA
chimp            AGGATCATCAATGTCGGCCCCCAGGG------TTCACAGCTGTTCTGAGCCCCGCCCCCA
rhesus           GGGATCATCAATGTCGGCCCCCAGGG------TTCACAGCTGTTCTGAACCCCGCCCCCA
mouse            AGAATCATCAAA----ACAACCTGGGGAAGTGAGCGTGGCGGTCCCGAGCCCCGCCCTCA
rat              AGAATCATCAAA----ACATCCTGGGGAGGTGTGCG--GCGGTCCCGAGCCCCGCCCTCA
                 .*.********:    .*. **:***       : *.   ** ** * **.******** **
                                                   XBP1   (ERSE I) NF-YA/B
hGaltpro         GGTGGCAGGGCAG--CCCAGTCAGTCAGTCACGTGCTGGCGGCTGGCCAATCATCGGGGG
chimp            GGTGGCAGGGCAG--CCCAGTCAGTCAGTCACGTGCTGTCGGCTGGCCAATCATCGGGGG
rhesus           GGTGGCAGGGCAG--CCGT------CAGTCACGTGCTGGCGGCTGGCCAATCATCGGGGG
mouse            AGTGGCAGGCTGTGCCCCCA-----CCGTCACGTGTTGGCGACTAGCCAATCATCCAGGG
rat              AGTGGCAGGCTGT--CCCCC-----TCGTCACGTGTTGGCGGCTAGCCAATCATCCAGGG
                 .********   .    **          .******** ** **.**.********** .***
```

## *Hk1:*

```
                                                      AHR
hHK1intron       CCCCGCCTGGCACGGCCCACCCGGGCACGGCGCCGCCACGCGTGAAGGCCCCGCATCGGCT
Gorilla          CTGCGCCTGGCACGGCCCACCCGGGCACGGCGCCGCCACGCTTGAAGGCCCCGCATCGGCT
rhesus           CCCCGCCTGGCCCGGCCTACTCGGGCACGGCGCCGCCACGCGTGAAGGCCCCGCATCGGCT
mouse            CCCACTCTGGCTTG----------GCACTGTGCCACCACGCGTGACCGCGCG------CT
rat              CCCGTTCAGGCATG----------GCACTGTGCCGCCACGCGTGACCGTGCG------CT
                 *     *:***  *           **** * ***.****** ***. *   **      **
                 XBP1/Creb3l2  XBP1/Creb3l2
hHK1intron       CCCTACGTGGGGGACGTGCAGGATGATGCGGGGTCGGGGGGGATTTCGCTGCGTCGCCCG
Gorilla          CCCTACGTGGGGGACGTGCAGGATGATGCGGGGTCGGGGGGGATTTCGCTGCGTCGTCCG
rhesus           CCCTACGTGGGGGACGTGAAGGGTGATGCGGAGCATGGGGGGATCTCGCTGCGTCGCCCG
mouse            CCTTACGTGGGGAACGTGCAGGGTGAT---------------------------------G
rat              CCTTACGTGGGGGACGTGCAGGGTGCT---------------------------------G
                 ** *********.*****.***.**.*                                 *
                                      KLF
hHK1intron       CCCCCTTTCCGGCGCAGGAGGGAGCGGCCGCCGCGTCCCCGCT
Gorilla          CCCCCCTTCCGGCGCGGGAGGGAGCGGCCGCCGCGTCCC----
rhesus           CCCCCTTTCCGGCGCGGGAGGGAGCGGCCGCCGCGTCCCAGCT
mouse            CCTCCTTTCCGGCGCGGGAGGGAGCGGCCGTCATTCTCCTG--
rat              CCTCCTTTCCGGTGCGGGAGGGAGCGGCCGTCTTTCTCCTG--
                 ** ** ****** **.************** *     **
```

#### *Gfpt1:*

```
                                    Creb3l2
hGfpt1           ATCCACGTGTACAGAAGATGATTCAAGAGGTTATAAATAACTCATATGAGGTAA
Rhesus           ATCCACATGTACAGAAGATGATTCAAGAGGTTATAAATAACTCATATGAGGTAA
Chimp            ATCCACGTGTACAGAAGATGATTCAAGAGGTTATAAATAACTCATATGAGGTAA
Mouse            ACCCAAG---CATGTGGGTGACTCAGGGTGGCATAAATAATTCGTGAGAGGAAG
Rat              ACCCAAG---AGGGTGGGTGACTCAGGGTGGCACCTCAGGATGGTGTAAATAAT
                 * ***..    .   *:.*.*** ***.*.  *   *  .:.:.. *  .*.:.*. :*
```

```
                                                                             Ets
hGFAT1promoter       CTCCCTCTCGGCTCGGTCCCTCCCGCGGCTCGCCCCGGGGCGGGCGTCTCCAGGAACTCC
chimp                CTCCCTCTCGGCTCGGTCCCTCCCGCGGCTCGCCCCGGGGCGGGCGTATCCAGGAACTCC
squirrelMonkey       CTCCCTCTCGGCTCCTTCCCTCCCGTGGCTCGTCCGTGGGAGGGCGTCTCCAGGAACTCC
mouse                ------CTCGGCTCG------------------CCCTCAGCGGGCGGCTCCAGGAACCTA
rat                  ------CACGGCTCG------------------CCCTCAGCCGGCGGCTCCAGGAACCTG
                           *:******                  **    .*.  ****  .*********
                                                 Xbp1/Creb3l2
hGFAT1promoter       AGGCGGACGCGGCCACGAGCCACTGCCACGTCGCCGCCGCCAGCAGTCCAGCGCCTGCGT
chimp                AGGCGGACGCGGCCACGAGCCACTGCCACGTCGCCGCCGCCAGCAGTCCAGCGCCTGCGT
squirrelMonkey       AGGCGGACGCGACCACGAGTCTCTGCCACGTCT---CCGCCAACAGTCCAGCGCCTGCGT
mouse                AAGCGGATGCTGG-TCGAGCCCCTGCCACGTCG---CTACCCGCAGCTCTGCGTCTGCG-
rat                  AAGCGGATGCTAG-TCCAGACCCTGCCACGTCGTCGCTGCCCGCAGCTCTGCGTCTGCG-
                     *.***** **  .   :* ** * **********     *  .**..***  *:*** *****
                                                            AHR
hGFAT1promoter       GGGCTCGGGGGGCGGGGCAGCGCTGGGAGGCGGGACTTGCGCGTGAGCGGCGCGTGAGCG
chimp                GGGCTCGGGGGGCGGGGCAGCGCTGGGAGGCGGGACTTGCGCGTGAGCGGCGCTTGAGCG
squirrelMonkey       GGAGTCGGGGGGCGGGGCAGCGCTGGGAGGCGGGACCTGCGCGTGAGCGGCGCTTGAGCA
mouse                -----CGCTGGGCGGG--------------CGATGCCTGCGCGTGAGCGGCGCGGGCGCG
rat                  -----CGTGGCAGGGG--------------CTAGACCTGCGCGTGAGCGGCTCGGGCGCG
```

```
                    **   *  .  ***               *  .  .*  **************  *   *.**.
```

***Gnpnt1:***

```
                       ETS              XBP1/Creb312
hGnpant1           TTGGGAACCG-CGGCCCAGACGTGGCAGCGCCAACGCCTCCACCTCGCCTCTGCCCCCTCACGCAGG
chimp              TTGGGAACCG-CGGCCCAGACGTGGCAGCGCCAACGCCTCCACCCCGCCTCTGCCCCCTCACGCAGG
GreenMonkey        TAGGGAACCG-CGGCCGAGACGTGGCAGCGCCAACGCCTCCACCCCGCCTCTACCCCCTCACGCAGG
mouse              -CCGGAACCGCCAGGCCAAACGTGGCCGGACCAACGCCTCTGCCCTGCCCCG---CCCTCACGCACC
rat                -CCGGAACCGCCTGGCCGAACGTGGCGGGACCAACGCCTCCGCCCTGCCCCG---CCCTCACGCACC
                      ******* * * *  ..******* *  .**********  .**   *** *     **********
```

***Pgm3:***

```
                                                NFYA/B          ERSE II-like
hPgm3             --GCTGGCGGATAGTCCTCTGCCGTGATTGGCCAGG------------------------
rhesus            --GCCGGCGGACCTTCCTCCGTCGTGATTGGCCGGG------------------------
chimp             --GCTGGCGGATAGTCCTCTGCCGTGATTGGCCAGG------------------------
mouse             --GGCCACAGACACCAACCCGCCGCGATTGGTCGGGGCGGGGCTTGGTGGTGCAGGGGCG
rat               GGGGTCACAGAAACCAACTCGCCGCGATTGGCCGGGGCGGGGCCTGGTGATGCAGGGGCG
                    *    .*.**  .   ..    *  **  ****** *.**
                                                              Xbp1        ETS
hPgm3             ---------------------------------GGGCGTGGCGACGAGCCGGAAGCCACG
rhesus            ---------------------------------GGGCGCGGCGACAAGCCGGAAGCCACG
chimp             ---------------------------------GGGCGTGGCGACGAGCCGGAAGCCACG
mouse             GGTTCAGCGGCCCTTCCGGGCTGTGATTGGTGTGGGCGGGGCGTCGGGCCGGAAGCAAAG
rat               GGGCCATCGGACCTTCCCAGCCGTGATTGGTGGGGGCGGGGCGTCGGCCCGGAAGCCAGG
                                                   *****  ****:*..  ********.*  *
                        XBP1/Creb312
hPgm3             CAGGGCCACGTCTGCGG-TTCTGAGGACTGGGTTTG---GGTGCAGACGTTGTTGCTTGG
rhesus            CAGGGCCACGTCTACGG-TTCTGAGGACTAGGTCTGGCTGCTGCAGACGTTGTCGCTTGG
chimp             CAGGGCCACGTCTGCGG-TTCTGAGGACTGGGTTTG---GGTGCAGACGTCGTTGCTTGG
mouse             CTGGGCCACGTCTGCGAGGTCTGAG---AGAGTTGGTTTGGTGCGCAC-----TGCGCGG
rat               CTCGGCCACGTCTGCGAGATCTGAG---TGAGCTGGTTAGGTGCACAC-----TGCGCGG
                  *:  **********.**.   ******    :..*    *    * ***.  **       **   **
```

***Uap1:***

```
                                XBP1/Creb312          ETS
hUAP1             TGTGCTCCCGGCGCTGACGTGTCTGGGCGGTCGGCTTCCACTCCTTCAGGCGTCGGCAGC
chimp             TGCGCTCCCGGCGCTGACGTGTCTGGGCGGTCGGCTTCCACTCCTCCAGGCGCCGGCAGC
rhesus            TGCGCTCCGGACGCTGACGTGTCTGGGCGGTCGGCTTCCACTCCTCCGGGCGCCGGCAGC
mouse             -------CGGTCGCTGACGTGTCGGGCCGGCAGGCTTCCGCTCCGCCCCTGGGCCGCAGC
rat               -------CGGTCGCTGACGTGTCCAGCCGGCAGGCTTCCGCTCCGCCCCTGGGCCGCAGC
                         * * ************ .* ***  .*******.****   *     * * *****
                                       SP1
hUAP1             CACTAGTCGTGGCGAGAGGGGCGGGGTGGCCGGGGCTGGCGCTCCACTTGGCCCCCGCTC
chimp             CACTAGTCGTGGCGAGAGGGGCGGGGTGGCCGGGGCTGGCGCTCCCCTTGGCCCCCGCTC
rhesus            CACTAGTCGTGGCGAGAGGGGCGGGGTGGCCGGGGCTGGCGTTCCCCTTGGCCCCCGCTC
mouse             CACTCACCGCGGCGGGAGGGGCCGGGTGGCCGGGGCCCGCGCTCCCCTCGGCCGCTGCTC
rat               CACTCACCGCGGCGAAAGGGGCGGGGTGGCCGGGGCCCGCGCTCCCCTCGGCTGCTGCTC
                  ****.. ** ****..****** *************  *** ***.** ***  * ****
```

***Gale:***

```
                                                                     KlF        E-box
hGalepromoter       GGGGCTTCCTCCCCGTACTACCCG-------CAGAGTTTGTTCAGGGTGGAGGTGCGCAC
chimp               GGGGCTTCCTCCCCGTACTACCCG-------CAGAGTTTGTTCAGGGTGGAGGTGCGCAC
rhesus              GGGGCTTCCTCCCCGTCCCGCCCGTTCTGCCCGCAGTTTGTTCAGGGTGGAGGTGCGCAC
mouse               GG-ACTGCCTCTCTGTCCGCCCAG---------CAGTTTGCTCAGGGTGGGGATGCACAC
rat                 GG-ATTTCCTTCCTGTCCGCCCAG---------CAGTTTGCTTAGGGTGGGGATGCACAC
                    **  .  * ***   *  **.*   **.*        ****** * *******.*.***.***
                                              Xbp1/Creb312
hGalepromoter       CTG----------------GCCAAGTCCACGTCTACACGG-CCCACCCAGAGTAGCCTAAG
chimp               CTG----------------GCCAAGTCCACGTCTACACGG-CCCACCCAGAGTGGCCTAAG
rhesus              CTG----------------GCCAAGTCCACGTCTACACGG-CCTACCCAGAGAGGCCTAAG
mouse               CTGGCAGCGGAGAGCGAG-CCAAATCCACGTCGACACCGGCTCACTTGGTGCTGCCCTCC
rat                 CTGGCAGTGGAGTGCGAGGCCAAATCCACGTCGACACG-----ACTCCGTGCTGCCCTAG
```

```
                ***             ****.******** ****      **   *:*  *** :.
```

***Pmm2:***

```
hPmm2          ATGGGAACGGAGTCCCCTCCTCTTCCCGACGTGCCCTGCGACTCAGCGGCCGAACCCGGA
chimp          ATGGGAACGGAGTCCCCTCCTCTTCCCGACGTGCCCTGCGGCTCAGGGGCCGAACCCGGA
rhesus         GTGGGAACGGAGTCTCCTCCTCTTCCCGACGTGCCCTGCGGCTCGGGGGCCGAACTCGGA
mouse          CCATTAGGG-AATCTTCCCATGGTGCCTTGGAACTCCAGGGCTCGGAG--CGGACCCGGA
rat            CCATTAGAG-AATCTTCCCATCGTGCCTTGGAGTTCCGGGGCTCGGAG--CGGACCCGGA
                 .  *. * *.**  * *.*  * ** : *:.  * . *.***.* *  **.** ****
                NFKB
hPmm2          AGTTCCGGGCCGAGTTCCTCGTGCCAACGTGTCTTGTAAGGTG-CGGCTAGAAACTGGGG
chimp          AGTTCCGGGCCGAGTTCCTCGTGCCGACGTGTCTTGTAAGGTG-CGGCTAGAAACTGGGG
rhesus         AGTTCCGGGCCGAGTTCCTGATGCCAACGAGTCTTGCGAGGTG-CGGTTAGACGCTGGGG
mouse          AGTTCCGGGTAGAGTTCCGGGT---AGAGTTCCTGGTGCGGTCGAGGTTTCTGACTGGAG
rat            AGTTCCGGGTGGAGTTCCGG---------------GTGCCAACGAGTCTTCTGACTGGAG
               *********  *******                  * .. .:  .*  *: : .****.*
                                                         Xbp1/Creb3l2
hPmm2          ACATGGCAGCGCCTGGCCCAGCGCTCTGCCTCTTCGACGTGGATGGGACCCTCACCGCCC
chimp          ACATGGCAGCGCCTGGCCCAGCGCTCTGCCTCTTCGACGTGGATGGGACCCTCACCGCCC
rhesus         ACATGGCAGCGCCTGGCCCAGCGCTCTGCCTCTTCGACGTAGATGGGACCCTCACCGCCC
mouse          AAATG------------GCCACTCTCTGTCTCTTCGACATGGATGGGACCCTGACTGCCC
rat            AAATG------------GCCACGCTCTGTCTCTTTGACATGGATGGGACCTTGACTGCCC
               *.***            *..* ***** ***** ***.*.********* * ** ****
```

***Gmppa:***

```
hGMPPApromoter   TCTTTCCTCTCCCATAAAG--TTAGAAAGCGGGTAAAGGGTCAGACTACAATTCCCGGCA
chimp            TCTTTCCTCTCCCATAAAG--TTAGAAAGCGGGGAAAGGGTCAGACTACAATTCCCGGCA
rhesus           GCTCTCCTCTCCCATTAAG--TTTGAAAGCGGTGAAAAGTTCAGACTACAATTCCCGGCA
mouse            TTTCCCAAGTGTCTTATTGGAGTTGGAGAGAGAACGTTTTCAAAACTACAATTCCCAGCA
rat              CTTCCCAACTGTCTAATAAGAGTTGGAGAGAGGACATTTGCAAAACTACAATTCCCAGCA
                   *  *.: *  *:::::.  *:*.*.. .*  ..:   .*.***********.***
                                                    XBP1/Creb3l2  ETS
hGMPPApromoter   GACCCTGCGAATGAGTTCTCGCGACTTGCGAGAAGACACGTGCGCGGAAGGAGCTTGCAG
chimp            GACCCTGCGAATGAGTTCTCGCGACTTGCGAGAAGACACGTGCGCGGAAGGAGCTTGCAG
rhesus           GAACCTGCGAATGAGTTCTCGCGACTTGCGAGAAGACACGTGCGCGGAAGAAGCTTGCAG
mouse            GCCCCCGCGAAAGAGCTCTCGCGGGTTGTAAGGAGACACGTACGCGGAAGGAGCAAGCAG
rat              GCCCCTGCGAAGGTGCTCTCGCGGGTTGTAAGGAGACACGTACACGGAAGGAGCAAGCAC
                 *..** ***** *:* *******. *** .**.********.*.******.***::***

hGMPPApromoter   TAGCGGGCGGCAGAGCTGGAGTGAAGGGAGCTAGTGGTAAAGGGAGCTGGTGG-----AG
chimp            TAGCGGGCGGCAGAGCTGGAGTGAAGGGAGCTAGTGGTAAAGGGAGCTGGTGG-----AG
rhesus           TAGCGGGCGGCAGAGCTGGAGTAAAGGGAGCTAGTGGTAAAGGGAGCTGGTGG-----AG
mouse            TGGCAGGCGGCAGAGCTGGCGTAAAGGGAGCTAGTGGTAAAAGGATCTACATGGTGGAGG
rat              TGGCAGACGGCAGAGCTGGCGTAAAGGGAACTAGTGGTAAAAGGACCTAAATGGTGGAGG
                 *.**.*.***********.**.******.***********.*** **. : *     .*
```

**Gmppb:**

```
                                                   NF-YA/B (ERSE II)  XBP1
hGMPPB          CGGACCCGGCGCGGGCAGTGACGCGACCACCGCGGGCCAATCGGCTGCCGCGTACGTGGC
chimp           CGGACCCGGCGCGGGCAGTGACGCGACCACCGCGGGCCAATCGGCTGCCGCGTACGTGGC
greenMonkey     CGGACCCGGCGCGGACAGTGACGCGAGCACCGCGGGCCAATCGGCTGCCGCGTACGTGGC
mouse           CGGAGGCGGCGCAGCCAGTGACGTAACAGGG-AGAGCCAATC----------TACGTGGC
rat             CGGAGGCGGCGCATCCAGTGACGTAACAGGGGCGAGCCAATCGGCTGTTAACTACGTGGC
                ****  ******.  ******** .*  ..   .*.*******          ********

hGMPPB          ACGGCCGCCGGCGTCGGAACG---
chimp           ACGGCCGCCGGCGTCGGAACG---
greenMonkey     ACGGCCGCCGGCGACGGAGCG---
mouse           AGGGCCGCCCGCAGCCGAGCG---
rat             AGGGCCGCCCGCAGCCGATCGCCC
                * ******* **. * ** **
```

***Gmds:***

```
                                                      Creb3l2
hGMDS_DHS          ATGACATTTTGACTGGGCAGGCTCCAGTTCTGGACAGGTCTTGATTTATTGCGACTGAGT
chimp              ATGACATTTTGACTGGGCAGGCTCCAGTTCTGGACAGGTCTTGATTTATTGCGACTGAGT
mouse              ATGACATTTTGACTGGGCAGGCTCCAGCTCTGGACAGGTCTTGATTTATTACGACTGAGT
```

```
rat                  ATGACATTTTGACTGGGCAGGCTCCAGCTCTGGACAGGTCTTGATTTATTACGACTGAGT
rhesus               ATGACATTTTGACTGGGCAGGCTCCAGTTCTGGACAGGTCTTGATTTATTGCGACTGAGT
                     *************************** **********************.*********
                                 Creb3l2
hGMDS_DHS            CCCAACCAACTTTGATTTATTTCAGCTTAGGGGAGGAAGGCAAGCTTGGAACAGAAAAGC
chimp                CCCAACCAACTTTGATTTATTTCAGCTTAGGGGAGGAAGGCAAGCTTGGAACAGAAAAGC
mouse                CTCAACCAACTTTGATTTATTTCAGCTTTGGGGAGGAGGGCAGGCTCAGAACAGAAAAGC
rat                  CTCAACCAACTTTGATTTATTTCAGCTTTGGGGAGGAGGGCAGGCTCAGAACAGAAAAGC
rhesus               CTCAACCAACTTTGATTTATTTCAGCTTAGGGGGGGGAGGCAAGCTTGGAACAGAAAAGC
                     * **************************:****.**..****.***  .***********
                                     Xbp1/Creb3l2  (Overlapping)
hGMDS_DHS            GCTGTTGAATGTTGAGCACGTGATGTCAGGACAGGCCCTCCAGACTTCCAGGGGCACGG-
chimp                GCTGTTGAATGTTGAGCACGTGATGTCAGGACAGGCCCTCCAGACTTCCAGGGGCACGG-
mouse                CCTGTTGATTGTCGAGCACGTGATGTCAGGACAAGACCTCCAGACTTCCTAGGGCTGTAG
rat                  GCTGTTGATTGTCGAGCACGTGATGTCAGAACAAGACCTCCAGACTTCCTAGGGCTGTAG
rhesus               GCTGTTGAATGTTGAGCACGTGATGTCAGGACAGGCCCTCCAGACTTCCAGGGCCACGG-
                      *******:*** ****************.***.*.*************:.** *:   .
```

***GFUS:***

```
                          XBP1/Creb3l2       CEBP
hGFUSexon            GGATGACGTGTGTGGGTTGGACCTTCTCAAACA
Chimp                GGATGACGTGTGTGGGTTGGACCTTCTCAAACA
Rhesus               GGATGACATGCGTGGGTCGGACTTTCTCAAACA
SquirrelMonkey       GGATGACGTGCGTGGGTCGGACCTTCTCAAACA
Mouse                GGATGACGTGGGTGGGCTGTACTTTCTGGAAGA
Rat                  GGATGACGTGGGTGGGCTGTACCTTCTGGAACA
                     *******.** *****  * ** **** .** *
```

***Nans:***

```
                                                 NFYA/B ERSE II-like Xbp1
hNans                ------GGCTGTCGGGAGAGAGGCGGGGCCTAGGGGATTGGCTGCCCGGCGACCGCGGGC
chimp                --GATTGGCTGTCGGGAGAGAGGCGGGGCCTAGGGGATTGGCTGCCCGGCGACCGCGGGC
rhesus               ------GGCTGTCGGGAGCGAGGCGGGGCCTAGGGGATTGGCTGAGCGGCGTCCGCGGGC
mouse                CCGATTGGCTGTCG--AGCGAGACGGGGCCCGCGG-ATTGGCTGCGCGACAGCCGCCGGC
rat                  CCGATTGGCTGTGG--AGAGAGACCGGGCCTGCGG-ATTGGCTGCGCGGCGGCCGCCGGC
                           ****** *  **.***.* ***** . ** ********. **.*. **** ***
                         Xbp1/Creb3l2
hNans                TGACGTGGCGGGGCTGGCGTGTGGGTCTCGCAGCGTTGCTCACAGAACAGAGTAGAGGCG
chimp                TGACGTGGCGGGGCTGGCGTGTGGGTCTCGCAGCGCTGCTCACAGAACAGAGTAGAGGCG
rhesus               TGACGTGGCGGG-CAGGCGTGTGGGTCTTGTAGCGCTGCTCACAG--AAGAGTAGAGACG
mouse                TGACGTGGCGGGGCTG-------AGACCTGCGGGCCTGC------------------GCG
rat                  TGACGTGGCGGGGCCG-------AGACCTGCGGGCCTGC------------------GCG
                     ************ * *        .*:*  * .*   ***                  .**
```

***Cmas:***

```
                                 TEAD
hCMAS                GAGGCATCAGGTTCATGAATGGAGTTGAGTGTCGGGGTGTCAGAGCTGAATTTCAAGCTC
chimp                GAGGCATCAGGTTCATGAATGGAGTTGAGTGTCGGGGTGTCAGAGCTGAATTTCAAGCTC
rhesus               GAAGCATCAGGTTCGTGAATGGAGTTGAGTGTCGGGGTGTCAGAGCTAAATTTCAAGCTC
mouse                GAGGCAGCAGGTGCATGAATGGAGTCGAGT--TGGGGCTCCC-----------------C
rat                  GAGGCAGCAGGTGCATGAATGGAGTCCAGAGTTGGGGCTCTG-----------------C
                     **.*** ***** *.**********  **:    ****                    *

hCMAS                AGCTTGGAGATCTTTTCCTAACCTTTG------AAAAATACTCCAGCTATCTTCCTGTGA
rhesus               AGCTTGGAGATCTTCTCCTAACCTTTG------AAAAATACTCTAGCTATCTGCCTGTGA
chimp                AGCTTGGAGATCTTTTCCTAACCTTTG------AAAAATACTCTAGCTATCTGCCTGTGA
mouse                AGCTTAGAGAGTCTTCCCTCACCTTTGAGAAAAAAATATTCTCTGGCTGTGAGGCTGTGA
rat                  AGCTTAGAGAGTCTTCCCTCACCTTT---GAAAAAAAATTCTCTGGCTGTGAGGCTGTGA
                     *****.****   *  ***.******       ***:**:*** .***.* :  ******
                      Creb3l2
hCMAS                GTCAGCACAATCTAGAAAA-CTCCCTACATTTATAAATACGTGAGACTCAGTGGGAAGGT
chimp                GTCAGCACAATCTAGAAAA-CTCCCTACATTTATAAATACGTGAGACTCAGTGGGAAGGT
rhesus               GTCAGCACAATCTAGAAAG-CTCCCTACATTTATAAATACGTGAGACCCAGTGGGAAGGT
mouse                GTCAGCACAAAGAGGAAATGGTCCCTACGTGTG----TACGAGG----------------
rat                  GTCAGCACAAAGAGGAAATGCTCACTACGTGTG----TGCGAGG----------------
                     **********: :.****   **.****.* *.    *.**:*.
```

*Genes for the biosynthesis of Dolichol-diphosphate GlcNAc2Man9Glc3, a precursor for N-linked glycosylation*

***Dpagt1:***

```
                                                    Xbp1/creb3l2
hDpagt1              CCGGTTCCAAGATGAATGCCTCCTTGTACCACGTGTCTGACACCGCTCACATTCCGGTTC
```

```
chimp                CCGGTTCCAAGATGAATGCCTCCTTGTACCACGTGTCTGACACCGCTCACATTCCGGTTC
rhesus               CCGGTTCCAAGATGAATGCCTCCTTGTACCACGTGTCTGATACAGCTCACATTCCGGTTG
mouse                CCAGTTCCAACATGAACGCTTCCTCGTACCACGTGGCTGACACCGCTAGTTCTGTGGTTC
rat                  CCAGTTCCAACATGACCGCTTCCTC-TATCACGTGGCTGACACCGCTTGTTCTGTGGTTC
                     **.******* ****. ** ****  ** ****** **** **.***  . : *   ****
                     Xbp1/Creb312
hDpagt1              CACGTCAGTTTTAGCTCCGTCCGCCTCCATAGGTCAAGCTTAAAGGGCCCGTACCTCTCC
chimp                CACGTCAGTTTTAGCTCCGTCCGCCTCCATAGGTCAAGCTTAAAGGGCCCGTACCTCTCC
rhesus               CACGTCAGTTTTAGCTCAGTCCGCCTCCGTAGGTCAATTTTAAAGGGACCGTACCTTGCC
mouse                CACGTCGT-------------CTCCCTCGCAGGTCGGGTTTAAAGGGCTAGCAGCCTGAG
rat                  CACGTCG---------------TCCCCCGCAGGTCGGGTTTAAAGGGCTAGCAGCTT---
                     ******.                ** *. *****.. ********. .* * *
```

#### *Alg9:*

```
                                                         Creb312      ERSE II
hAlg9                CCTATTCCTCCAGGGGAGAGAAAGCACACTGAGAATTGACGTCA-GACCCAATTCCATTC
Mouse                GCTATTCCTCCAGGGGAGAGAAAGCACACTGAGAATTGACGTCA-GACCCAATTCCATTC
Chimp                CCTATTCCTCCAGGGGAGAGAAAGCACACTGAGAATTGACGTCA-GACCCAATTCCATTC
Rhesus               CCTATTCCTCCAGGGGAGAGAAAGCACACTGAGAATTGACGTCA-GACCCAATTCCATTC
Rat                  GCCATTCCTCCAGGGGAGAGAAAACACACTGAG-ATTGGCGTCA-GACCCAATTCCATTC
                      * ********************.********* ****.***** ***************
                      Xbp1
hAlg9                CACATAAC
Mouse                CACATAAC
Chimp                CACATAAC
Rhesus               CACATAAC
Rat                  CACATAAC
                     ********
```

<u>*Mitochondria for producing Acetyl-CoA*</u>

#### *Cpt1a:*

```
                                     KLF                        XBP1/Creb312
chimp                CTGCGGCTTCGCGCCCCGCCCCTTCCCCCAAGCCCCGGGCGG-CGTCACGTGACGGCTGA
mouse                CAGCGGCTTCCCGCCCCGCCCCTTCCCCTA----CCCAGCCG-GGTCACGTGACGGCTGA
rat                  CCGCGGCTTCCCGCCCCGCCCCTTCCACCCA--CCCGGCCAG-GGTCACGTGATGGCTGA
hCpt1a               CTGCGGCTTCGCGCCCCGCCCCTTCCCCCAAGCCCCGGGCGG-CGTCACGTGACGGCTGA
rhesus               CTGCGGCTTCGCGCCCCGCCTCTTCCCCCAAGCCCCGGGCGGGCGTCACGTGACGACTGA
                     * ******** ********* *****.*  .     **  .  * *  ********* *.****
                                                          KLF
chimp                GAAAAGTAGGGGAAAGGTCAGCAGCAGCCCCGCCCCCGGGCTGGCGCGTCCCAGGCGCTC
mouse                GAAAAGTAGGGGAAAGGTTA-CGGGTGCCCCGCCCCCGTGCTGGCGCGTCCGGGGCGCTC
rat                  GAAAAGTAGGGGAAAGGTTA-CGGGAGCCCCGCCCCCGTGCTGGCGCGTCCGGGGCGCTC
hCpt1a               GAAAAGTAGGGGAAAGGTCAGCAGCAGCCCCGCCCCCAGGCTGGCGCGTCCCAGGCGCTC
rhesus               GAAAAGTAGGGGAAAGGTCAGCAACAGCCCCGCCCCCGGGCTGGCGCGTCCCAGGCGCTC
                     ****************** * *.. :***********. ************ .*******
                                                              ETS
chimp                GGATCCTGTTCCACCCGCCGTGCCCACAGAGACCTTCCGGAGCGAGTTGGCGCGCGTCCC
mouse                GGATCCTGTTCCACCGGCCGAGCCCACGCAGGCCTTCCGGAGCGA-------CGCGTCGC
rat                  GGATCCTGTTCCACCGGCCGAGCCCACGCAGGCCTTCCGGAGCGA-------CGCGTCGC
hCpt1a               GGATCCTGTTCCACCCGCCGTGCCCACAGAGACCTTCCGGAGCGAGTTGGCGCGCGTCCC
rhesus               GGATCCTGTTCCACCCGCCGTGCCCACAGAGACCTTCCGGAGCGAGCTGGCGCGCGTCCC
                     *************** ****:******. **.*************       ****** *
```

#### Slc25a1:

```
                                                                  NFY/A/B
hSlc25a1promoterR    TCCCGCCGCTCGCGGTCGCCCAGGGACTCGGCCTCGGCCATTGGCTGCGCTTTGG-CCCC
chimp                TCCCGCCGCTCGCGGTCGCCGAGGGACTCGGCCTCGGCCATTGGCTGCGCTTTGG-CCCC
greenMonkey          CCCCGCCGCTCGCGGTCGCCCAGAGACTCG------GCCATTGGCTGCGCTTTGG-CCCC
mouse                ---CGGGGTTCTCCGTCTCCCCGGAATCCAGCACCGGCCATTGGCTGCTCTCAGCTCTCC
rat                  ---CGGTGTTCTCCATCTCCCCGGAACCCAGCACCGGCCATTGGTTGATCTCAGCTCTCC
                         **  * ** * .** ** .*..*  *.       ******** **. ** :*  * **
                          XBP1
hSlc25a1promoterR    GCCCCTACGCCTGCCTCGGCGCGCCCCGCCCAGGCCTCAGTTTCCCGGCCCGCACGGGGC
chimp                GCCCGTACGCCTGCCTTGGCGCGCCCCGCCCAGGCCTCAGTTTCCCGGCCCGCACGGGGC
greenMonkey          GCCCCTACGCCTGCCTCGGCGCGCCCCGCCCCGGCCTCAGTTTCCCGGCCCGCTTGGGGC
mouse                GCCCCGACGCGTGCCCCAGTGCATCCCGCCCAAGCCTCAGTTTCCCGGCTGGCACGGGGC
rat                  GCCCCGACGCGTGCCTCGGTGCATCCCGCCCAAGCCTCAGTTTCCCGGCTGGCACGGGGC
                     ****  **** ****   .* **. *******..****************  **: *****
                            KLF                                         XBP1/Creb312
```

```
hSlc25a1promoterR   GGGCTGGGGGCGGGGCCTGGCTCGGACCACGCGGGGCGGGACCTGG-AGCTGACGCGGCC
chimp               GGGCTGGGGGCGGGGCCTGGCTCGGACCACGCGGGGCGGGACCTGG-AGCTGACGCGGCC
greenMonkey         GGGCTGGGGGCGGGGCCCGACTCGGACCACGCGGGGCGGGACCTGG-AGCTGACGCGGCC
mouse               GTGGCGTGGGCGGGGCTCAGCTCAGGCCACGCGGGGCGGAGCCCGGGAGCTGACGTGACC
rat                 GTGGCGTGGGCGGGGCTCAGCTCAGGCCACGCGGGGCGGAGCCGGGGAGCTGACGTGACC
                    * *  * *********  ..***.*.*************..** ** ******** *.**
```

---

***Acly:***

```
hACLY               GATTCCCTGGCGGCGGCTGTGGCGAGACAGCCCTCATCCTAGTAAGTCCCAAAACCCCCG
chimp               GATTCCCTGGCGGCGGCTGTGGCGAGACAGCCCTCATCCTAGTAAGTCCCAAACCCCCCG
mouse               GGGTCCCTGGCGGCAGCAAAGGCGAGACAGCCCTCATCCTGAGCCCCCCGCCTGCTTCCC
rhesus              GATTCCCTGGCGGCAGCCACGGCGAGACAGCCCTCATCCCGGTAAGTCCCAAACCCCCCG
rat                 GGGTCCCTGGCGGCAGCAAAGGCGAGACAGGCCTCATCCCCAGTCCCCCCACCTGCCCCC
                    *. ***********.** . ********** ********  .  .  ** ..     **

                                        Creb3l2
hACLY               CCTGGCCTGTGCGTCACTGCTGAGTCAGGGACCAGACAGCAATCCATAAGAGGTAATGAG
chimp               CCTGGCCTGTGCGTCACTGCTGAGTCAGGGACCAGACAGCAATCCATCAGAGGTAATGAG
mouse               GCTCGCCG-TGCGTCACTGCTGAGTCAGGGACCAGACAGTGATCCATCAGAGGTAATGAG
rhesus              CCCAGCCTGTGCGTCACTGCTGAGTCAGGGACCAGACAGCAATCCATCAGAGGTAATGAG
rat                 CGCCTGCCGTGCATCACTGCTGAGTCAGGGACCAGACAGTGATCCATCAGAGGTAATGAG
                          *  ***.************************** .******.************

hACLY               CCACTCCTGCTGG-CCCAACGCTGCTGCCAG-CTCAGATGTCTGATG--TCCCACCTTCC
chimp               CCACTCCTGCTGG-CCCAACGCTGCTGCCAG-CTCAGATGTCTGATG--TCCCACCTTCC
mouse               CCGCTCCAGCCAGGTCTGACACTGCTGCCAAACTCAGATGTCCGTGA--CATCTCCCTCC
rhesus              CCGCTCCTGCTGG-CCCAAGGCTGCTGCCAG-CTCAGATGTCTGATGTCCCCCACCCTCC
rat                 CCGCTCCAGCCAGGTCTGACACTGCTGCCAAGCTCAGATGTCCGTG--ACATCTCCCTCC
                    **.****:** .*  * .* .*********. ********** *:     . *:** ***
```

<u>*Transporters:*</u>

***Slc2a1*/Glut1**

```
                                   Creb3L2
Hslc2a1             -TAACAG----TCCAAAACCTGACCTTAGGGTGACCTCAAGCATTAAGTGCTCCTGAAGC
rhesus              -TAACAG----CCCGAAGCCTGACCTTAGGGTGACCTCAAGCATTAAGTGCTCCTGAAGC
chimp               -TAACAG----TCCAAAACCTGACCTTAGGGTGACCTCAAGCATTAAGTGCTCCTGAAGC
mouse               TAATCCGGGAACACCAACACTGACCTTCATGTGACCTTAGGCCCTGATTCCTCTT----C
rat                 TAATCTGGGAACACCAATGTTGGCCTTCATGTGACCTTAGGTCCCCATTCCTCTT----C
                     :*:* *     .* **   **.****.. ******* *.*  .   * * *** *    *
                       ETS                Creb3l2           Creb3l2
Hslc2a1             CACTTCCTTTAAGCCTTCCAATGATATCAAGACCCTTGCATGACTGAACCTCCAGACACC
rhesus              CACTTCCTTTAAGCCTTCCAGTGATGTCAAGACCCTTGCATGACTGAACCTCCCGACACC
chimp               CACTTCCTTTAAGCCTTCCAATGATGTCAAGACCCTTGCATGACTGAACCTCCAGACACC
mouse               AATTTCCTTAAAACCTTCTAGTGATGTTCAGAT--TTGCATGACTGAC-----------T
rat                 AATTTCCTTAAAGCCTTCTAGTGATGTTAAGAT--TTGCATGACTGAC-----------T
                    .* ******:**.***** *.****.* .***   ************
```

---

***Slc1a5:***

```
                                Creb3l2
hSlc1a5             GA-GAACCCTGTTCCGTGACTCATCATTTCCTTCCCCCAAC-------------------
chimp               GG-GAACCCTGTTCCGTGACTCATCATTTCCTTCCCCCAAC-------------------
rhesus              GGGAAACCCTGTTCCATGACTCATCACTTCCTCCCCTCAAC-------------------
mouse               AGGATGCCCGGTTTTCTGACTCACGGTTTCCACATCCCCACCACCCCACCCACCTC----
rat                 AGGATGACCTGTTTTCTGACTCACTGTTTCCACATCCCCACCCCCACCCTCACCCCGCCC
                    .. .:..** ***   *******  .  ****: .  *  *.**
                                        NFkB
hSlc1a5             ---------------TTCCAGGAAGTCCCCAACGTGGAGACACTGAATGAGCAAGCCCTA
chimp               ---------------TTCCAGGAAGTCCCCAACGTGGAGACACTGAATCAGCAAGCCCTA
rhesus              ---------------TTCCAGGAAGTCCCCAACCTGGAGACACTGAATCAGCAAGCCCTA
mouse               --ACTTCTCTCTTCCTTCCAGGAAGTCCCCAATCTGAAGATACTGAATCAG-AGGCCCTC
rat                 CCGCCTCTCTCTTCCTTCCAGGAAATCCCCAATCTGAAGATGCTGAAACAG-AGGGCCTC
                                   *********.*******  **.*** .*****: ** *.* ***.
```

---

***Slc26a2***

```
hSlc26a2            GGGCAGCCAATCGCGAGGAGGAGAGTAGTCAGAGGGCGGCACCCTCCCCGGTCTGGGCCG
chimp               GGGCAGCCAATCGCGAGGAGGAGAGTAGTCAGAGGGCGGCACCCTCCCCGGTCTGGGCCG
rheusus             GGGCAGCAAATCGCGAGGAGGAGAGTAGTCAGAGGGCGGCACCCTCCCCGGTCTGGGCGG
mouse               GAACAGCCAATCGGTAAGAAGCAAGCGGCCGAGGGGCGGGGCTGTCGCCGG-CGGGCCCC
rat                 GGACAGCCAATCACGAAGAAGCAAGCGGTCGAGGGGCGGGGTTGTCGCCGG-CCGGCCCA
```

```
                 *..****.****.  *.**.*..** .* *...****** .   ** **** * ** *
                                                          XBP1/Creb3l2
hSlc26a2         AGTTATTGGCTGGTGGTAGCGTCGCTCGCCCGGCAGTACCACGTGACGGCCTCGGCCGCG
chimp.           AGTTATTGGCTGGTGGTAGCGTCGCTCGCCCGGCAGTACCACGTGACGGCCTCGGCCGCG
rheusus          AGTTATTGGCTGGTGGTAGCGTCGCTCGTTCGGCAAAACTACGTGACGGCCTCGGCCGCG
mouse            GGTGATTGGCCAGGGG---------CCATGCCGCCACGCCACGTGACTGCCACG-CCGCG
rat.             GGTGATTGGCCGGCGGT--------CCACACCACCATGCCACGTGACTGCCACG-CCGCG
                 .** ****** .* **          *.  * .*.. .* ******* ***:** *****
```

#### *SLC35b2*

```
                                             KLF
hSLC35b2         CGACGGCCGAG-----CAGCCGGGACCCCACCCGGCCCCCG-TGCTGAGGCGCGGCAGTC
Gibbon           CGACGGCCGAG-----CAGCCAGGACCCCACCCGGCCCCCG-TGCCGAGGCGCGGCAGTC
GreenMonkey      TGACGGTCGAG-----CAGCCAGGACCCCACCCGGCCCCCAGTGCAGATGTGCGGCAGTC
mouse            CTGCGGCTGGA-----GGACC----CCCCACCCCGCCTCCGGAGCCCTGGGGTCTCGGTC
rat              CTGCGGCCGGGGGACCCCTCCAACCCCCCACCCCGCCTCCCCAGCGCTGGGGACTCGGTC
                   .***   *..         **     ******** *** **   :**   : * *    *.***
                  XBP1/Creb3l2       XBP1/Creb3l2                 KLF
hSlc35b2         ACGTGGCCCGGGGG-CTCGGTCACGTGAGCTCGCTGCTGCCCTAG-CCCCCACCTGGCGT
Gibbon           ACGTGGCCCGGGGG-CTCGGTCACGTGAGCTCGCCGCTGCCCTAG-CCCCCACCTGGCGT
GreenMonkey      ACGTGGCCCGGGGG-CTCGGTCACGTGAGCTCACCGCTGCCCTAG-CCCCCACCTGGCGT
mouse            ACGTGGCCCCGGGGTCTCGGTCACGTGGGC-CATGGCTGCCCGAGCCCCCCACCTGGCGT
rat              ACGTGGCCCCGGGATCTCGGTCACGTGGAC-CATGGCTGCCTGAGCCCCCCACCTGGCGT
                 ********* ***. ************..* *.   ******  ** **************
```

#### Slc35b3

```
                                             Creb3l2
hSLC35b3Intron   GCTAGGGTGATATGGTGTGCTCAAGCTGACTCACCCCTTTGATGTGCAG
Chimp            GCTAGGGTGATATGGTGTGCTCAAGCTGACTCACCCCTTTGATGTGCAG
Rhesus           GCTAGGGTGATACGGTGTGCTCAAGATGACTCACCCCTTTGATGTGCAG
Mouse            GCTGAAGAGGTCTGGTGTGCTTAAGCTGACTCATCCTCCGGACATGGAA
Rat              GTTGAAGTGGTCTGGCACGATTAAGTTGACTCACCCTCCTGACACGAAG
                 * *...*:*.*. ** . *.* *** ******* **    ** . * *.
```

#### Slc35c1

```
                                                            XBP1/Creb3l2
hSlc35C1promoter GCTTTAAG-GGCAAGGCGGGGCGTGCGGGCCCTTTAAGGCCACGTGGGGGCGTGTCAGGA
chimp            GCTTTAAG-GGCAAGGCGGGGCGTGCGGGCCCTTTAAGGCCACGTGGGGGCGTGTCAGGA
rhesus           GCTTTAAG-GGCAAGGCGGGGCGTGCGGGCCCTTTAAGGCCACGTGGGGGCGTGCCAGGA
mouse            GCGGGAAGTGGCAACGGCCTGCGAGCTGGCCCTTTAAGGCGGCTCGTAGGCGTGTCAGGA
rat              GCG-AAGTGGGAAAGGCCCTGAGAGCTGGCCCTTTAAGACTGTTCGTAGGCGTGTCAGGA
                 **    *.  **.** *    *.*:** ***********.*  .    * .****** *****

hSlc35C1promoter AGTGAGTCCAGGGCCCGCCTCCCGGGGAGTCGGCCTCGGATGTCCGGAGGCTCCTGGGCT
chimp            AGTGAGTCCAGGGCCCGCCTCCCGGGGAGTCGGCCTCGGATGTCCGGAGGCTCCTGGGCT
rhesus           AGTGAGTCCCGGGCCCGCCTCGCGGGGAGTAGGCCTCGGCTGTCCGGAGGCTCCTGGGCT
mouse            AATGCGCGCAGGGCCCGCCCTGCTCGGTAAGTGGCCCG--GGACCCGCGTCGCTGAGCCG
rat              AATGCTT-CTGGGCCCGCCCCGCTCGGTAAGTGGCCCG--GGACTGGCGACGCTGAGCCG
                 *.**.   * *********   *  **:.:  * * **   *:*  *.* * *   .* *
```

#### *Slc35a5:*

```
                                          ETS    ETS
hSlc35a5         CCTCGCTTTG-CTTCACT-CGCGCCCCTTCCGGCTTCCCTT---CCTTCTCACTCTCTTG
chimp            CCTCGCTTTG-CTTCACT-CGCGCCCCTTCCGGCTTCCCTT---CCTTCTCACTCTCTTG
rhesus           CCTCGCTTTG-CTTCACT-CGCGCCCCTTCCGGCTTCCCCTT--CCTTCTCACTCTCTTG
rat              CGTCCCCCAAGCTCCACT-TGAACCCCTTCCGGCTTCCCCTTCCTCTTCTCACACTCTCC
mouse            CTTCCCCGGAGCTCCACTGAACACCCCTTCCGGCTTCCCCTTCCTCTTCTCACACTCTCC
                 * ** *    . ** ****  ...**************** *    ********:****
                                        XBP1/Creb3l2        XBP1/Creb3l2
hSlc35a5         CCGTAGCTGCGCCGCCACCGGGG-CCTCACGTGACACTAGACTCTCCCGGCACGTGACGT
rhesus           CCGTAGCTGCGCCGCCACCTGGG-CCTCACGTGACACTG-ACTCTCCCGGCACGTGACGT
chimp            CCGTAGCTACGCCGCCACCGGGG-CCTCACGTGACACTAGACTCTCCCGGCACGTGACGT
rat              CAACAGCCCCGCCGCCTCCTAGG-CCTCACGTGACTCGAATCTCCCGGGACACGTGACAG
mouse            CAACAGCCCCGCCGCCACCTAAGGCCTCACGTGACTCGACTCTCCCAGGACACGTGACAA
                 *.. ***  *******:** ..* ***********:* . :*** *  *.********.
```

### *Sulfated Mucin-Type O-Glycan Biosynthesis in Golgi Stacks:*

#### *Galnt12:*

```
                                                                         TEAD
hGalnt12upstream      GCCTAATTGCATTTCTCTTTTCTCTGTATCTGAACCTCTATTGAGAAGCATTCCAATTTG
chimp                 GTCTAATTGCATTTCTCTTTTCTCTGTATCTGAACCTCTATTGAGAAGCATTCCAATTTG
mouse                 ------------TTTTTTTTCCATC----CTAAGCCTCTGCAGGGAAGCATTCCAATTTG
rat                   ---------ATTTTTTTTTTCCACC----CTAAGCCTCTGTGGAGAAGCATTCCAATTTG
                                  ** * *** *:      **.*.*****.  *.****************
                                                                          Creb3l2
hGalnt12upstream      GCAAGCGGATCCAAGACATTATGAATTGTAATCCTTGTCTGTTTTCTAATCTGACCGCAG
chimp                 GCAAGCGGATCCAAGACATTATGAATTTTAATCCTTGTCTGTTTTCTAATCTGACCGCAG
mouse                 GCAG-CCAATCCAAGCCCTTCTGAATTGTAACCCTTGTCTGTTTTCCAATCTGACCGCAG
rat                   GCAG-CCGATCCAAGCTCTTCTGAACTGTAATCCTTGTCTGATTTCCAATCTGACCACAG
                      ***. * .*******. .**.**** * *** *********:**** *********.***

hGalnt12upstream      ATCTTAAATGTGTTGGGTGTCTGAACAGAGGCCAAAGGGCTCAGGTCCAGACCTACCTAA
chimp                 ATCTTAAATGTGTTGGGTGTCTGAACAGAGGCCAAAGGGCTCAGGTCCAGACCTACCTAA
mouse                 ATCTTAACTAT---------CTGGAGACTGGCCCAAAGATGAGGCCCTAGGCTTTCCTAT
rat                   ATCTTAACTAT---------CTGGAGGCTGGCCCAAAGGTGGGGCCCTAGGCTTTCCAAT
                      *******.*.*         ***.* . :****.**.*.   .*  * **.* *:**:*:
```

#### *C1galt1*:

```
                    NF-YA/B  ERSE I   XBP1
hC1galt1        CGGCAGCCAATCCCCGCTTCCTCACGTCGCCGTAT-------------CCCCGCCCACCG
chimp           CGGCAGCCAATCCCCGCTTCCTCACGTCGCCGTAT-------------CCCCGCCCACCG
gorilla         CGGCAGCCAATCCCCGCTTCCTCACGTCGCCGTAT-------------CCCCGCCCACCG
mouse           CGGCGGCCAATCCCCGCATCCGCAGGTCGCGGTG-------------TCCCCGCCCACT-
nakedrat        CGGCAGCCAATCGTCGCGCCCGCACGTCTCCGGAGGCGGGGAGGCCCGCCCCAACCTGC-
                ****.*******  ***  ** ** *** * * .              ****..**:
                                 ETS
hC1galt1        CCCCTCCCGCTCCGCCCTTCCCTGGGGCTGGT----------------GCGGCGGCGGAGA
chimp           CCCCTCCCGCTCCGCCCTTCCCTGGGGCTGGT----------------GCGGCGGCGGAGA
gorilla         CCCCTCCCGCTCCGCCCTTCCCTGGGGCTGCT----------------GTGGCGGCGGAGA
mouse           CTGCCCAGGCCCCGCCCTTCCCCGGGGCTGCTGCTGCTGCTGCAGCTGCTGCAGCCGGGA
nakedrat        CCCCTCCGGCTCCGCCCTTCCCCGGGGCTGGG-----------TGGAGGCGGTGC--CGA
                *  * *. ** *********** *******                 *  *  **   **
CREB3l2
hC1galt1        CGTCAGCG-CCAGGAGACTTCGGG-TGGCGGCTGCGCCATAGGCGGGCCACGG-CCTGGA
chimp           CGTCAGCG-CCAGGAGACTTCGGG-TGGCGGCTGCGCCATAGGCGGGCCACGG-CCTGGA
gorilla         CGTCAGCG-CCAGGAGACTCCGGG-TGGCGGCTGCGCCATAGGCGGGCCACGG-CCTGGA
mouse           CGTCAGCATCTCCGAGACCCCTGGGTGGCGGCTGCGCAGTAGGCGGGCCGTGGTCCAGGC
nakedrat        CGTCAGCG--TTGGAGACCCCTGGGTGGTGGCGGCGCCGTGGGCGGGCCGCGG-CTTGGA
                *******.     *****  * ** *** *** ****..*.********. ** * :**.
```

#### *C1galt1c1:*

```
                                                  Xbp1/Creb3l2
hC1galt1c1      AGGAAAGCGCAGCCGCGCACGGGTT--TCCTCTCACGTTGGCGCACCACTCCG
Chimp           GAGAAAGCGCAGCCGCGCACGGGTT--TCCTCTCACGTTGGCGCACCACTCCG
Gorilla         AGGAAAGCGCAGCCGCGCACGGGTT--TCCTTTCACGTTGGCGCACCACTCCG
Prairie         AGGGAAGCACAGCCTGACGCGGCTT--TCCTCTCACGTCCACCGGCCCCTCAG
Rat             AGGAAAGCGCTGCCTTACGCCGCTT--TCCTCTCACGTCCACGGCCCACTCGG
Naked           AGGAAACCGCAGCGGTGCACAGCTC--TCCTTTCACGTTAGCGCGCCACTCGG
Mouse           AGGAAAGCGCTGCCTCACGAGGCTTGGACGTGAGAGGTCCAAGGCCCACTCTG
                ..*.** *.*:**   .*.. * *   :* * : * **  ..   **.*** *
```

#### *Gcnt1:*

```
                   ETS
hGcnt1          GAGGAAACACCAAGGAAAAATGGAAAGATGGAAAGCTGACGGATGAGTGTGAGCTGACCC
chimp           GAGGAAACACCAAGGAAAAATGGAAAGATGGAAAGCTGACGGATGAGTGTGAGCTGACCC
rhesus          GAGGAAACACCAAGGAAAAATGGAAAGATGGAAAGCTGACGGATGAGAGTGAGCTGGCCC
mouse           GAGGAAAACTCAGGAAGGCTTGGAGTCTTTGTAAGGG-AGGGGAGAGCGTTTACTGACAG
moleRat         GAGGAAATAGCAAGGAAGAACTAGAATATGAAAAGCG------TGAGTGTGAACTGACCG
                ******* . **.*.*...:  ...: :* .:***        :*** ** :.***.*.
                                                             Creb3l2
hGcnt1          AGCAGATTCGGTGAGGCCAACACCTAAGCTGGCAGTGAGGGAAAGTGAGTCATCCATCCT
chimp           AGCAGATTCGGTGAGGCCAACACCTAAGCTGCCAGTGAGGGAAAGTGAGTCATCCACCCT
rhesus          AGCAGAT-CAGCGAGGCAAACACCTCAGCTGCTAGCAAGGGAAAGTGAGTCATCCATCCT
mouse           AG---ACTCAAT-AAACTGAAACCCACATTGGCAGGGCTGGAGAATGAGTCATTTGTCAG
moleRat         AGCAGACTCTGG-AAGTGACAACCCAAGCTGGCAGTGGG-AAAAGTGAGTCATCCATCAT
```

```
                    **   *  * .  *..  ...*** ... **  ** .   .*.*.******** . *.
```

***Chst4 promoter:***

```
                    KLF                          CREB3L2
hCHST4Pro      TAGGAGGAGGAAGCCAAG--AGGGGAGTTGGTGAGTCAGAGGAG-AAAAGCGCATGGCCC
chimp          TAGGAGGAGGAAGCCAAG--AGGGGAGTTGGTGAGTCAGAGGAG-AAAAGCGCATGGCCC
rhesus         TAGGAGGAGGAAGCCAAG--AGGGGAGTTGGTGAGTCAGAGGAG-AAAAGCGCACGGCCC
mouse          -AGGAGGAGGAG---------------TTCATGAGTCAGAGAGAGAACGGTGCTGAGCCT
rat            GAGGAGGAGGAG---------------TTCATGAGTCAGAGAAAGAAAGGTGCTCAGCCT
                **********.               :* .**********... **..* **:  ***
```

***Chst4 intronic enhancer:***

```
                                                          Xbp1 (variant)
hCHST4Intron    GTTATTATGAATGGAAGATAAGGGG------TTTTTCTGCTATTGCCACCCAAGGTTTGG
chimp           GTTATTATGAATGGAAGATACGGGG------TTTTTCTGCTATTGCCACCCAAGGTTTGG
rhesus          GTTATTATGAATGCAAGAAAAGGGG------TTTTTCTGCTATTGCCACCCAAGGTTTGG
mouse           CTGGCAGTGAACGCGAGAGATGGTGGGGGTTGGGGGGTGCCACTGCCACACAAAGTCTCG
rat             GGAAGAGAGATGGGGAGGGATGGAGGAG-AAGGAAGGTGCCACTGCCACACAG--TCTGG
                   . :.:**: * .**. * ** *          *** * ******.**.  * * *
                  Creb3l2           ETS/COUP     NF-κB      ETS
hCHST4Intron    CTTGAGTCATTACATACTTCCCAGAAGGTTCAGGGGACTTCCAAGGAAAACCCACGAGCT
chimp           CTTGAGTCATTCCATACTTCCCAGAAGGTTCAGGGGACTTCCAAGGAAAACCCACGAGCT
rhesus          CTTGAGTCATTTCATACTTCCCAGAAGGTTCAGGGGACTTCCAAGGAAAACCCACGAACT
mouse           TGAGGACCATGACACACTTCCAGAATGTTCAAGGGGACTTCCAAGGGAAGCTGAG-AGCT
rat             TGAGGATCGTGACACACTTCCAGAATGTTCAAGGGGATTTCCAAGGGAAGCCCAAGAGCT
                  :*.. *.*  ** ******...*:* * .****** ********.**.*   *   *.**
                                  ETS
hCHST4Intron    GGCAAAGCATGC--TTTCCCTTCAGGAAGAAAGCTATAAATGCCAAGAGTCTCCTTTGTG
chimp           GGCAAAGCATGC--TTTCCCTTCAGGAAGAAAGCTATAAATGCCAAGAGTCTCCTTTGTG
rhesus          GGCAAGGCATGC--TCTCCCTTCAGGAAGAAAGCTATAAGGGCCAAGAGTCTCCTTTGTG
mouse           GGCAAAGCATGCCCTAACTCTTGAGGAAACAAGTTACCGACTCCAAGAATCACCTCTGCG
rat             GGCAAAGCATGCCCTACCTCTTGAGGAAATAAGTTACAAACTCCAAGAATCCCCTCTGTG
                *****.******  *  * *** *****. *** **  ...  ******.** *** ** *
```

***Chst3:***

```
                                       Creb3L1          SREBF
hCHST3            -GCCACACCTAGAAACGGACTATGAATCAAGCCAGGGGCCCAGTGCCAGCTTCCTTGGCC
Chimp             -GCCACACCTAGAAACGGACTATGAATCAAGCCAGGGGCCCAGTGCCAGCTTCCTTGGCC
Rhesus            -GCCACACCTGGAAACGGACTATGAGTCAAGCCAGGGGCCCAGTGCCAGCTTCTTTGGCC
Gorilla           -GCCACACCTAGAAACGGACTATGAATCAAGCCAGGGGCCCAGTGCCAGCTTCCTTGGCC
Orangutan         -GCCACACCTGGAAACGGACTATGAGTCAAGCCAGGGGCCCAGTGCCAGCTTCCTTGGCC
SquirrelMonkey    -GCCACACCTGGAAAAGGAGTATGAGTCAAGCTAGGGGCCCAGTGCCAGCTTCCTTGGCC
Mouse             GGTCAGACCTGGAAAAAGAATATGAGTCAGGTTGGGG-CGGGCTGCCAGCTTCCTCGGCC
Rat               GGTCAGCCCTGGAAAAAGAACATGAGTCAGGCTGGGG-CGAGCTGCCAGCTTCCTCGGCC
ChineseHamster    GGTCAGACCTGGAAAAAGAATATGAGTCAGGCCGGGG-CTAGGTGCCAGCTTCCTCGGCC
                   * ** .***.****..**  ****.***.*  .*** *   . ********** * ****
```

***Fut4:***

```
                          Xbp1/Creb3l2
hFUT4          CGGCGCCTTCATCCACGTGGACGACTTCCCAA
Rhesus         CGGCGCCTTCATCCATGTGGACGACTTCCCAA
Chimp          CGGCGCCTTCATCCACGTGGACGACTTCCCAA
Gorilla        CGGCGCCTTCATCCACGTGGACGACTTCCCAA
Mouse          TGGCGCCTTCATCCACGTGGACGATTTCCCTA
Rat            TGGTTCCTTCATCCATGTGGATGATTTCCCTA
                **  ********** ***** ** *****:*
```

***FUT7***

```
                              Creb3l2                KLF
hFUT7promoter  GGTGGGGACAGTGGTTGATGCCAAAGGTTGTGGGGGCAGGGGCGGGGCAGGAGCAGGAAG
baboon         GGTGGGGACAGTGGTTGATGCCAAAGGTTGTGGGGGCAGGGGCGGGGCAGGAGCAGGAAG
rhesus         GGTGGGGACAGTGGTTGATGCCAAAGGTTGTGGGGGCAGGGGCGGGGCAGGAGCAGGAAG
mouse          GTTG--GATAGTGGTTGATGCCAAAGATTGAAGGGGTAGGG-CGGGGCAGAAGTGGGAAG
rat            GTTG--GATGGTGGTTGATGCCAAAGATTGAAAGGGTAGGG-TGGGGCAGAAGTGGGAAG
               * **  ** .***************.***:..*** ****  *******.** .*****
               NF-κB
```

```
hFUT7promoter   GTCCCCTGAGTTCCCTCACCTTGGGCAGAGATAAAAGGAGCACAGTTCCAGGCGGGGCTG
baboon          GTCCCCTGAGTTCCCTCACCTTGGGCAGAGATAAAAGGAGCACAGTTCCAGGCGGGGCTG
rhesus          GTCCCCTGAGTTCCCTCACCTTGGGCAGAGATAAAAGGAGCACAGTTCCAGGCGGGGCTG
mouse           GTCCCTG--GCTTCCTCACCTTGGTAGATGGTGAGGAGCCCCAG-----AGGTTGAGCTG
rat             GTCCCTG--GCTTCCTCACCTTGGTCGATG-AAACAAAAGCATC-----AGGCTGAGTTG
                *****    * * *********** ...:* :.* .... *.       ***  *.* **
                                Creb3l2
hFUT7promoter   AGCTAGGGCGTAGCTGTGATTTCAGGGGCACCTCTG-----GCGGCTGCCGTGATTTGAG
baboon          AGCCAGGGCGCAGCTGTGATTTCAGGG-CATCTCTG-----GCGGCTGCTGTGATTTGAG
rhesus          AGCCAGGGCGCAGCTGTGATTTCAGGG-CATCTCTG-----GCGGCTGCTGTGATTTGAG
mouse           AGCAG-----CAGCTGTGATTTCAGGG-TGCCTCTGTTGGAGAGGCTGCTGTGATTTGAA
rat             AGCAG-----TAGCTGTGATTTCAGGG-TGCCTCTGTTGGAGAGGCTGCTGTGATTTGAA
                *** .      ****************  . *****     *.****** *********.
```

***ST3GAL4:***

```
                                       E-box                            NFAT
hST3GAL4        -------CTTGGCAATTGCTTCCATCTGGCCTGCCTGCAG---CTCCTATTCCTTTGGAA
chimp           ----TCCCTTGGCAATTGCTTCCATCTGGCCTGCCTGCAG---CTCCTATTCCTTTGGAA
rhesus          -----CCCTTGGCAATTGCTTCCATCTGGCCTGCCTGCAG---CTCCTATTCCTTTGGAA
mouse           -----CCGCCGGTAATTACTTCCATCTGCCCCGCCTGCCTGCTCCCCTTTT-CCTTGGAA
rat             CTGCCCCGCCGGTAATTACTTCCATCTGCCCTGCCTGCCTGCTCTTCTTATTCCTTGGAA
                          ** ****.********** ** ******.    *  **::* * ******
                                                                      XBP1/Creb3l2
hST3GAL4        ATGTGGCTGCCGCCCACATGGCTGCAAG-----------GCAACCACGGCCACACGTGGG
chimp           ATGTGGCTGCCGCCCACATGGCTGCAAG-----------GCAACCACGGCCACACGTGGG
rhesus          ATGTGGCTGCCGCCCACATGGCTGCAAG-----------GCAACCACGGCCACACGTGGG
mouse           ATGCAGCCGGCTCCCACAAGGCTGAAAAGACTTTCTTGGGCAACCGTTGAGACACGTGGG
rat             ATGCGGCCGCCGCCCACAAGGCTGAAAAGACTTTCTTGGGCAACCGCTGGGACACGTGGG
                *** .** * * ******:*****.**.           ******.  *  *********
                      KLF
hST3GAL4        GAAGGGGAGGGGCCCTATGGGGTCTAAGCACAGGGAACCCAGCCAAAGTGGCTGCTCCTG
chimp           GAAGGGGAGGGGCCCTATGGGGTCTAAGCACAGGGAACCCAGCCAAAGTGGCTGCTCCTG
rhesus          GAAGGGGAGGGGCCCTATGGGGTCTAAGCACAGGGAACCCAGCCAAAGTGGCTGCTCCTG
mouse           GAAGGGGAGGGGCTCTATTGCTGTCTGAACCTGGGCACCCACCCAG--------------
rat             GAAGGGGAGGGGCTCTGTTGGTGTCTAAGCCCGGGCACCCACCCAGG--------TCACC
                ************* **.* *     :.. .* ***.***** ***.
```

***St6galnac2:***

```
hSt6galnac2         GAGCGGTAGGAGCGGGGCTGGGAGTAGTAGCGGGAGAAGGGGGTGTAGAAGGTGGCAGGC
orangutan           GAGCGGTAGGAGCGGGGCTGGGAGTAGTAGCGGGAGAAGGGGGTGTAGAAGGTGGCAGGC
squirrelMonkey      GAGCGGTAGGCTCGGGGGTGGGAGTAGTAGCGGGAGAAGGGGGTGTAGAAGGTGGCAGGC
mouse               -AGCGGTAGGGCCGGGGCTGGGAGTAGAAGCGGGAGAATGGGGTATAGAAGGTGGCTGGT
rat                 -AGCGGTAGGGCCGGGGCTGGGAGTAGAAGCGCGAGAACGGGGTATAGAAGGTGGCTGGT
                     *********  ***** *********:**** ***** *****.************:**
                                    Xbp1/Creb3l2
hSt6galnac2         AGTACGTTGGGGACCACGTGCGGCTTCCCGCTCACCAGGACCTCCACGGGCGGGGTGTAG
orangutan           AGTACGGTGGGGACCACGTGCGGCTTCCCGCTCACCAGGACCTCCACAGGCGGGGTGTAG
squirrelMonkey      AGCGAGGTGGGGACCACGTGCGGCTTCCCGCTCACCAGGACCTCGATGGGTGGGGTGTAG
mouse               AGCACAGATGGAACCACGTGCGGCTTTCCCTTCAGCAGGACCTCCGTCGGTGGAGTGTAG
rat                 AGGACCGACGGTACGACGTGCGGCTTCCCCTTCAGCAGGACCTCCGTCGGTGGAGTGTAG
                    ** ..  : ** ** *********** **  *** ********* .  ** **.******

hSt6galnac2         GAGAGAGAGGAGGGGGGCATGAACAGACCGGGGGGAGTTGGGAAGGGGCAGTTAAACTGG
orangutan           GAGAGAGAGGAGGGGG-CATGAAGAGACCTGGGGGAGTTGGGAAGGGGCAGTTAAACTGG
squirrelMonkey      GAGAGAGAGGAGGGGGGCGTGAACAGAGCGGGGGGAGTTGGGAAGGGGCAGTGAAACTGG
mouse               GAGAGGGAGGATGGGGGTGTGAACAGGGCTGGGGGTGTGGGGAAGGAGCACTGGAACTGG
rat                 GAGAGGGAGGATGGGGGTATGAACAGGGCTGGGGGTGTGGGATAGGAGCACTGGGACTGG
                    *****.***** ****  .**** **. * *****:** **.:***.*** * ..*****
                                                           Creb3l2
hSt6galnac2         GGGTGGTGGGGGGGCGGGGGAAGGAGAGGTAAGTAGAGTGAGTCAGGAGGGGGTGG----
orangutan           GGGTGGTGGGGGGGTGGGGGAAGGAGAGGTAAGTAGAGTGAGTCAGGAGGGGGTGG----
squirrelMonkey      GGTTGGTGGGGGGGTGGGGGAAGGAGAGGTAAGTAGAGCGAGTCAGGAGGGGGTGA----
mouse               GGGATGTGTGGAGGGGGTGGCAGGAGGGGGAAGTAGGTTGAGTCAGTAGGTGGCG-----
rat                 GGAACGTGGGGGGGAGGTGGCAGGAGGGGGAAGTAGAGAGAGTCGGGAGGTGGTGGTGGT
                    ** : *** **.** ** **.*****.** ******.  *****.* *** ** *
                                                                   CEBP
hSt6galnac2         --------GGGCTTCCGGGCAGGGCAGGGAGAGGGACAGGCAAAGCATGGGTTTTGGGGA
orangutan           --------GGGCTTCTGGGCAGGGCAGGGAGAGGGACAGGCAAAGCATGGGTTTTGGGGA
squirrelMonkey      --------GGGCTTCTGGGTGGGGCAGGGACAGGGACAGGCAAAGCATGGGTTTTGGGGA
```

```
mouse           -GCTGCCTGGATTGCTGGGCCTGGCACGGGAAAGGGCAGGGGAAACACCCGTTTTGTGGA
rat             GGCTGCCTGGTCTGCTGGGCCTGGCACGGGAAGGGGCAGGGGAAGCACCGATTTTGTGGA
                        **  * * ***   **** **. *.**.**** .**.**   .***** ***
                                       NFAT                CEBP
hSt6galnac2     AGTGGTGCTGTGGGGGGCTGGAAAAAGGGGGGCTCCTTCAGAAAGCGGGGCACATAGGG
orangutan       AGTGGTGCTGTGGGGGGCTGGAAAAAGGGGGGCTCCTTCAGAAAGCGGGGCACATAGGG
squirrelMonkey  AGTAGTGCTGTGGGGGGCTGGAAAAAGGGAGGCTCCTTCAGAAAGCGGGAAACATAGGG
mouse           AGAGGTGCTGTAGGGGGCTGGAAATAGGGGGGCTCTTTCATGAAGCGCGGCACATAGGG
rat             AGAGGTGCTGTGGGGGGCTGGAAATAAGGTGGCTCTTTCATGAAGCGTGGCACATAGGG
                **:.*******.***********:*.** ***** **** .***** *..********
```

#### *St6gal1:*

```
                                                XBP1/Creb3l2
hst6gal1        AGCGGAGGGG--GCACAGGCACCATAACAAACGCACGTGCGCCCTGCCCGGAGAGCTGGA
chimpst6gal1    AGCGGAGGGG--GCACAGGCACCATAACAAACGCACGTGCGCCCTGCCCGGAGAGCTGGA
rhesus          AGCGGAGGGG--GCACAGGCACCATAACAAACGCACGTGCGCCCTGCCCAGAGATCTGGA
mouse           AGGAGAGGGGGAGCGCAGGCACCACAACAAACGCACGTGTGCCCCG--------------
rat             GGGAGAAGG--AGTGCAGGCACCACAACAAACGCACGTGTGCCCCGCCAGGG---ATCGG
                .* .**.**    *  .********* ************** **** *
```

#### *Casd1:*

```
                                                                     Creb3l2
hCasd1          TTTGGGTTAACACTGGTTATTGCATGCTGATCAGAAATTCTAATTTGCTGGTAATGACAT
rhesus          TTTGGGTTAACGCTGGTTACTGCATGCTGATCAGAAATTCTAATTTGCTGTTGATGACAT
mouse           TGGGGGTTAGCTCTGGTTATTGAAGGTGGATCAGAAATTCTAATTTGGTGTTGATGACAT
squirrel        TGTGGGCTAACACTGGTTATTGAAGTCTGATCAGAAATTCTAATTTGCTGTTGATGACAT
chimp           TTTGGGTTAACACTGGTTATTGCATGCTGATCAGAAATTCTAATTTGCTGTTGATGACAT
                *  *** **.* ******* **.*    ****************** ** *.*******
                          Spdef                                        GATA
hCasd1          CATAG-AAATGTCTAGGATGTGATTGGCAGACCATATATTGTTGGCAGAG----TTATCT
rhesus          CATAG-AAATGTCTAGGATGTGACTGGCAGACCATATATTGTTGGCAGAG----TTATCT
mouse           CATAGTATGTGTCTAGGATGTGATTGGCAGGCCACATATTGTTAGCCCAG----TTATCT
squirrel        CATAG-AAATGTCTACGATGTGATTGGCAGACCATATATTGTTAGCAGAGTTATTTATCT
chimp           CATAG-AAATGTCTAGGATGTGATTGGCAGACCATATATTGTTGGCAGAG----TTATCT
                ***** *:.****** ******* ******.*** ********.**. **     ******
                                                E-box
hCasd1          GGTTTTTACAATACCC-----AATTCTTACAACATGTGATCAACCTCTGATCAGAA---T
rhesus          GGTTTTTACAGTATCC-----AATTCTTACAGCATGTGATCAACCTCCGATCAGAA---T
mouse           GGTTGTTGTCGTATCTGATTCAATTCTGCCAGCATGTGATCAGCCTCAGATCAGAAGAAA
squirrel        GGTTTTTACAATATCTGATTTAATTCTGCCAGCATGTGATCAACCTCAGATCAGAAGAAT
chimp           GGTTTTTACAATACCT-----AATTCTTACAACATGTGATCAACCTCCGATCAGAA---T
                **** **.  ..** *      ****** .**.**********.**** ********   :

hCasd1          GACAGCCAAAATTAAAAGACTATTTGCAGGGAACAGTGCCTAAGTGTTTGTACACTTATT
rhesus          GACAGCCAAAATTAAAAGACTATTTGCAGGGAACAGTGCCTAAGTGTTTGTACACTTATT
mouse           G--AGCCAAAATTAAGCGACCACTTGCAGGGAACAGTGCCTAAGTGTTTGTACACTTATT
squirrel        GACAGCCAAAATTAAGCGACCATTTTCAGGGAACAGTGCCTAAGTGTTTGTACACTTATT
chimp           GACAGCCAAAATTAAAAGACTATTTGCAGGGAACAGTGCCTAAGTGTTTGTACACTTATT
                *  ************..*** * ** ***********************************
                    SRF                                  Creb3l2
hCasd1          TAACTTTATTGCTTATTAGCATTACATCCATGACAGTGATGTATGTGATTTCATTTAAGT
rhesus          TAACTTTATTGCTTATTAGTATTACATCCATGACAGTGATGTATGTGATTTCATTTAAGT
mouse           TAACTTTATTGCTTATTAGCATTATATCTATGACAGTGATGTATGTGATTTCATTTAAGC
squirrel        TAACTTTATTGCTTATTAGCATTACATTCATGACAGTGATGTATGTGATTTCATTTAAGT
chimp           TAACTTTATTGCTTATTAGCATTACATCCATGACAGTGATGTATGTGATTTCATTTAAGT
                ****************** **** **  *****************************
```

**Genes for Organellar Architecture:**

*ER-targeted translation, translocation, and N-glycosylation*

#### *Ssr1:*

```
                                                  XBP1/Creb3l2
hSSR1promoter   -----------GCCACAAAGAG-GCAGCAGCCGGCCACGTGACTTCCCGGTGGCCTGCTC
chimp           -----------GCCACAAAGAG-GCAGCAGCCGGCCACGTGACTTCCCGGTGGCCTGCTC
rhesus          -----------GCCACAGAGAGGCCAGCAGCCGGCCACGTGACTTCCGGGCGGCCGGCTC
mouse           GCCCGTAGGCTCCCCTGGTCCCCGCCACCGCCGGCCACGTGATCTCCCG---GCCCGCAG
```

```
rat            ------------ACCGGGACCCCGCCCCCCGCCGGCCACGTGACCGCGCG---GACCGCAG
                            .*.  ..: .    *.  *.*************    *   *    *.*  **:
                                      XBP1/Creb312                    SP1
hSSR1promoter  TCCCCACCTCTCCGCGCTGAGCAGCCACGTCGGCGGCGGTCCCTGGTCCAGGGAGGGGCG
chimp          TCCCCACCTCTCCGCGCTGAGCAGCCACGTCGGCGGCGTTCCCTGGTCCAGGGAGGGGCG
rhesus         TACCTAAGTCTCCGCGCTGAGCAGCCACGTCGGCGGCGTTTCCCGTTCAGGGGAGGGGCG
mouse          GCCCCGCCCCTGGGCGCTTAGAGGCCACGTCGACGGTGG---------CGGGGAGGGGCG
rat            GCCCCGCCCCTGGGCTCCTAGAGGCCACGTCGACGGTAG---------CGGGGAGGGGCG
                .**  ..   **   ** *   **..*********.*** .          ..**********
                                          SP1
hSSR1promoter  TGGCAAAGGCCCGTGCGCG-GTACGTGTCCCGCCCCTCGCTGCCCGGAGCCCGGATGAAG
chimp          TGGCAAAGGCCCGTGCGTG-GTACGTGTCCCGCCCCTCGCTGCCCGGAGCCCGGATGACG
rhesus         TGGCGAAGGCCCGTGCGTG-GTACGTGTCCCGCCCCTCGCCGCCCGGAGCCCGGATGAAG
mouse          ACGGATGG---CGCGCGTGCGTGCGTGCCCCGCCCCTC-CACCGCG--GCCCGGATGAAG
rat            ACGGGTGG---CGCGCGT----GCGTGCCCCGCCCCTC-CTGCGCG--GCCCGGATGAAG
               : *  .:.*    ** ***      .**** ********** *   * **   **********.*
```

***Ssr4:***

```
                                                              Xbp1/Creb312
chimp      ----------------------CGACCAGCTGGGCCCCTGGCTCAGGGAGGGGCCACGTC
rat        AAGGCTGAGAAGGAGGGAAGTGCGACCGGTTGGGACCCTGGCTAGGGAAACGGACACGTC
mouse      AAGGCTGAGAAGGAGGGAAGTGCGACCGGTTGGGACCCTGGCTAGGGAAGCGGACACGTC
hSSR4      ----------AGGGGGGGACCGCGACCAGCTGGGCCCCTGGCTCAGGGAGGGGCCACGTC
rhesus     ----------ATGGGGG-ATCGCGACCAGCTGGGCCCCTGTCTCGGGGAGGGGCCACGTC
                                 *****.* ****.***** **..**.*.  **.******

                  Creb312                          NFYA/B ERSE II-like
chimp      AGTGCTGCCAGAGACGTCACAATGCGG-CCCCAGCCCTTCGGTGCGCGATTGGCTGCCGC
rat        AGTGCCGCCAGCGACGTCACCATGCCCGACCCAGCCCGCCGACGCACGATTGGGTGCTGA
mouse      AGTGCCGCCAGCGACGTCACCATGCCCGACCCAGCCCGCCGACGCACGATTGGGTGCTGA
hSSR4      AGTGCTGCCAGAGACGTCACAATGCCG-GCCCAGCCGTTCGGTGCGCGATTGGCTGCCGC
rhesus     AGTGCTGCCAGAGACGTCACAATGCCG-GCCCGGCCCTTCGGTGCACGATTGGCTGCCGC
           ***** *****.********.****     ***.***    **.  **.******* *** *.
              Xbp1           ETS
chimp      TGCCACTTACGCGTCGCTCTTCCTCGTTTGCCCCTCGTGTTCATGGGAGCTCGTTTTCTT
rat        GGCCGCTTACGCGTCGCGCTTCCTCGTCTGCTCCGCATGTTCAGGGCGGCTTGCTCTCTT
mouse      GGCCGCTTACGCGTCGCGCTTCCTCGTCTGCTCCGCATGTCCAGGGTGGCTTGCTCTCTT
hSSR4      TGCCACTTACGCGTCGCTCTTCCTCGTTTGCCCCTCGTGTTCATGGGAGCTCGTTTTCTT
rhesus     TGTCACTTACGCGTCGCTCTTCCTCGTTTGCCCCTCGTGTTCATGGGAGCTCGTTTTCTT
            * *.************ ********* *** **  *.*** ** **  .*** * * ****
```

***Sec61b:***

```
                                          Xbp1/Creb312
hSec61b    GTGAAGATTGACAAAGGCAAAATGAGCTTCTAGTGACGTGGCCGTGGGAGTAGTTAAAGG
chimp      GTGAAGATTGACAAAGGCAAAATGAGCTTCTAGTGACGTGGCCGTGGGAGTAGTTAAAGG
rhesus     ATGAAGATTGACAGGGGCAAAATGAGCTTCTAGTGACGTGGCCATGGGAG----TAAAGG
mouse      GGAAAGATTAATAGGGGCCGAGTGAGCCCGAGCTGACGTGGCCATCTAGTTAGTTAAAGG
rat        GGGAAGATTCATAGGGTCCGGGTGAGCCCGAGCTGACGTGGCCATCCAGTTAG-------
           . .****** * *..* *....*****    :. ***********.*   ..
```

***Rpn1:***

```
                                            NF-kB
hRPN1promoter  CGGGTCGGCTTCCCGGTTGCCCCGGAGACAGGGTCCTTCCTTGTTCGGCGCCGCCATCTT
rhesus         CGGGTCGACTTCCCGGTTGCCCTGGAGACAGGGTCCTTCCTTGTTGGGCGCCTCCATCTT
chimp          CGAGTCGGCTTCCCGGTTGCCCCGGAGACAGGGTCCTTCCTAGTTCGGCGCCGCCATCTT
mouse          CCCACGATCG-CCCGGTTGCCCTGGAGACGGGGTCCTTTCTGACCGGGCCCAGCCATCTT
rat            CCCATGGGCG-CCCGGTTGCCTTGGAGACGGGGTCCTTCCTGACCCGGCCCAGCCATCTT
               *   .   . *    **********   ******.******** **  .    *** *.  *******
                                NF-YA/B   ERSE II  Xbp1
hRPN1promoter  GTGTCACCAGCCCTTGCCCAATGAACGGCTGCTCAGTGGCGGGAGGGCGGGCTACCGCGC
rhesus         GTGTCACCAGCCCTTGCCCAATGAGCGGCTGCTCAGTGGCGGGAGGGCGGGCTACCGCGG
chimp          GTGTCACCAGCCCTTGCCCAATGAACGGCTGCTCAGTGGCGGGAGGGCGGGCTACCGCGC
mouse          GTGGTGCTG----TTGGCCAATCAGCGGTTGCGCCGTGGCGGGGGGG-------------
rat            GTGGTGCTG----TTGGCCAATCAACGGTTGCGCCGTGGCGGGGGGGGGG----------
               ***   .* .     *** ***** *.*** *** *.********.***

hRPN1promoter  CCGGGCCCTACCCGCCCTGGCGCACTGCTCTTCCCGGTCATGGAGGCGCCAGCCGCCGGC
rhesus         CGGGGCTCTGCCCGCCTTGGCCCTCTGCTCTTCTCGGTCATGGAGGCGCCAGTCGCCCGC
chimp          CGGGGCTCTACCCGCCCTGGCGCTCTGCTCTTCCCGGTCATGGAGGCGCCAGCCGCCCGC
mouse          CGGGGCGCGTGTCG--GTGGCGCTCAGCTCG---CGGTCATGGAGTCGCCGGTCGCCTTG
rat            CGGGGCGCGTGTCG--GTGGCGCTCAGCTCG---CGGTCATGGAGGCGCCGATCGTCTTG
```

```
                   * **** *    **   **** *:*:****    *********** ****.. ** *
```

***Stt3a:***

```
                                XBP1/Creb312
hStt3aPromoter     GCAAACCGACACGTCACTCCTCGGCCCTCAACCAAACTGCTTGGGGCACGTTCTTTCCGC
rhesus             GCCGACCGACACGTCACTCCTCGGCCCTCAACCAAACTGCTTGGGGCACGTTCTTGCCGC
rat                GCAGACCGACACGTCACTTCTCGGCTCTCAACCAAACTGTTCAGAGCACGCTCTCTCCGC
chimp              GCAAACCTACACGTCACTCCTCGGCCCTCAACCAAACTGCTTGGGGCACGTTCTTTCCGC
mouse              GCAGACCGACACGTCACTGCTCGGCTCTCAACCAAACTGTTCAGAGCACGCTCTTTCCGC
                   **..*** ********** ****** ************* *  .*.***** ***  ****
```

***Stt3b:***

```
hSST3bpromoter     CGGGCTGCTGAACCCCTCGGCCCGACCCACGCGCCGAGGGCTAGGATAATGCGCCTGCGC
chimp              CGGGCGGCTGAACCCCTCGGCCCGACCCACGCGCCGAGGGCTAGGATAATGCGCCTGCGC
squirellMonkey     CGGGCCGCAGAGCCCATCCGCCAGACCCACGCGCCGAGAGGTGGTGTACCGCGCCTGCGC
mouse              TGGCACGTTCAGCACAGCGGCTGGACCCACGCGCCTAGGGCCCCG-CACCGCGCCTGCGC
rat                -----------------CAGCTGGACCCACGCGCCCAGGACCCTG-CACCGCGCCTGCGC
                                    * **   ************ **..        *. **********

                               XBP1/Creb312
hSST3bpromoter     GGCC-GTCACGTGGTGAGGGCTCCGCGCAAAGCGGGTTAAAAGTCACGACCGCGGAGAGC
chimp              GGCC-GTCACGTGGTGAGGGCTCCGCGCAAAGCGGGTTAAAAGTCACGACCGCGGAGAGC
squirellMonkey     GTCC-GTCACGTGGTGGGGGCTCAGCGCACAGCGGGTTAAAAGTCACGACCGCGGACAGC
mouse              GCCCCGTCACGTGGCGGGCGCCCCGCGCTGAGCGGGTTAAAAGTCACGGCCGCCGAGCAC
rat                GCCCAGTCACGTGGCGGGCGCTCCGTGCTGAGCGGGTTAAAAGTCACGGCCGCCGAGCAG
                   * *  ********* *.* ** *.* **: *****************.**** **  ..
```

***Ost4:***

```
                             Creb312
hOst4          TACCCTCCATCTTGCCTCAGACTATGTCCTGGAACCCTAGAAAAGAGGGGGAAAGACCTG
chimp          TACCCTCCATCTTGCCTCAGACTATGTCCTGGAACCCTAGAAAAGAGGGGGAAAGACCTG
rhesus         TACCCTCCATCTTGCCTCAGACTATGTCCTGGAACCCTAGAAAAGAGGGGGAAAGACCTG
rat            CACCCTCCATCTTGCCTCAGACTATGTCCTGGAAGCCTAGAAAAAAAGGGG-AAAGACCCG
mouse          CACCCTCCATCTTGCCTCAGACTATGTCCTGGAACCCTAGAAAAAAAGGGG-AAAGACCCG
                ********************************* *********.***** ******* *

                    ETS
rhesus         AGGTAAGGAAAATGCTGCAGCTTTCCT
rat            AAGTAAGGAAAATGCTGCAGCTTCCTT
chimp          AGGTAAGGAAAATGCTGCAGCTTTCCT
hOst4          AGGTAAGGAAAATGCTGCAGCTTTCCT
mouse          AAGTAAGGAAAATGCTGCAGCTTCCTT
               *.********************* * *
```

***Tmem258:***

```
                                       ETS                 Xbp1/Creb312
hTmem258       GCGGGTGGCCTGACGTTCAGCCGCCTTCCAAAGCCCGCGCTCCCGTCACGTGACCTGCTC
chimp          GCGGGTCGCCTGACGTTCAGCCGCCTTCCAAAGCCCGCGCTCCCGTCACGTGACCTGCTC
rhesus         GCGGGTCGCCTGACGCTCAGCCGCCTTCCAAAGCCCGCGCTCCCATCACGTGACCTGCCC
mouse          ----ATCGTTCCCAGCTTCGCCGCCTTCCAAAGCCCGCGCTCTCGTCACGTGATCTGCCA
rat            ----ATCGTTTCCAGCTTCGCCGCCTTCCAAAGCCCGCGCTCTCATCACGTGACCTACCA
                   .* *    ..* * .*********************** *.******** **.*  .

hTmem258       GCCACGCACAGCCTCTTGGGGCGCGTAGTGCAGGCTGCGTCCCCTCAGACGGCTGCTCTC
chimp          GCCACGCACAGCCTCTTGGGGCGCGTAGTGCAGGCTGCGTCCCCTCAGACGGCTGCTCTC
rhesus         GCCACGCACAGCCTCTTGGGGAGTGTAGTGCAGGCTGCGTCCCCTCAGACGGCTACTCTC
mouse          GGCGCGAACAGCCTCCTGGGAAATGTAGTGTATGCCCCACCAGGTCCGATAGCAGCTTTC
rat            GGCGCGAACAGCCTCCTGGGAAGTGTAGTGTAGGCCGTGCCACGTCCGATAGCAGATTTC
               * *.**.******** ****... ****** * **    . *.   **.** .**:..* **
```

***Rps3:***

```
                                 Xbp1/Creb312
hRps3          CCCTTCGATGAAGAGATGATGACGAGTCTGACTTGGGGATG-TTCTCTTT
Chimp          CCCTTCGATGAAGAGATGATGACGAGTCTGACTTGGGGATG-TTCTCTTT
Rhesus         CCCTTCGATGAAGAGATGATGACGAGTCTGACTTGGGGGTG-TTGCCTTC
Rat            CCCTTCGATGAAGAGGTGATGACGAGTCTGACTAGGAGGTATTTGTCTTT
Mouse          CCCTTCGATGAAGAGGTGATGACGAGTCTGAGTAGGAAGTG-TTGTCTTT
               ***************.*************** *:**...*. **  ***
```

***Rpl10:***

```
                                                             Xbp1/Creb312
hRpl10         GCGCCAGACGCATTTCCACTCGGCTTGGAGGTGGATTTAGTGCCACGTGCCCGAAA
Chimp          GCGCCAGACGCATTTCCACTCGGCTCGGAGGTGGATTTAGTGCCACGTGCCCGAAA
Rhesus         GCGCCAGACGCCTTTCCACTCGACTCGGAGGTGGATTTAGTGCCACGTGCCCGAAA
Mouse          ACGCCATAT---TCTCTTAACGTGGCTAAGGCGGGTCTTCTGTCACATGCTAGGAA
```

```
Rat             GCGCCATAT---TCTCCTCACATGGCTAAGGCGGATATTCTGTCACATGCTCTGAA
                .***** *    * ** :.:*.     .*** **.* *: ** ***.*** . .**


                               Creb3l2
HRpl10          TTCCCTTCGGTGTGGTGAGTAAGCGCAGTTGTC------
Chimp           TTCCTTTCGGTGTGGTGAGTAAGCGCAGTTGTC------
Rhesus          TTCCCTTCGGCGCGGTGAGTAAGCGCGGTTGTA------
Green           TTCCCTTCGGCGGGGTGAGTAAGCTCAGTTGTA------
Prairie         TTTCCTCCGGCGCGGTGAGTGAGTTGTCTGGTTAGTGT-
Mouse           TTTCCTCTGGCGCGGTGAGTGAGTT---CGGTCACGGTT
                ** * *  ** * *******.**       **
```

#### *Protein Folding*

##### *Man1a1:*

```
                                       Xbp1/Creb3l2
hMan1a1         TGAGG-CTAACTCAAATCTGACTGACGTGCAACTGAAATATGTGACAGTTTT
Rat             TGAGC-CTGACGTAAATCTGACTGACGTGCCGTTGAAATACGTGACCGTGTT
Chimp           TGAGG-CTAACTCAAATCTGACTGACGTGCAACTGAAATATGTGACAGTTTT
Rhesus          TGAGG-CTAACTCAAATCTGATTGACGTGCAACTGAAGTATGTGACAGTTTT
Mouse           TGAGC-CTGATGTCAATCTGACTGACGTGCGGTTGGAATACGTGACTACGTT
                ****  **.*   .******* ******** . **.*.** ***** .  **
```

##### *Canx:*

```
                                             Ets              NFYA/B ERSE I  Xbp1
chimp              CACCTCCCGCCCAACCAACAGCCACTTCCTGCCTCACTCCCGGCCAATCTTCACACTCCA
rhesus             CACCTCCCGCCCAACCAACAGCCACTTCCTGCCTCACTCCCGACCAATCGTCACACTCCA
hCANXpromoter      CACCTCCCGCCCAACCAACAGCCACTTCCTGCCTCACTCCCGGCCAATCTTCACACTCCA
mouse              TCCTTCATACCCAATCAGCAACCACTTCCTGCCTCGCTATTGACCAATCACCACGCTCCA
rat                TTCCTCATACCCAATCAGCAACCACTTCCTGCCTCACTCTTGTCCAATCATCACGCTCCA
                     * **. .***** **.**.**************.**.  * ******  ***.*****

chimp              CGAAGAATAGCCGTAGGGGGCGTATGGGACGGTGCGCAGAGCGATGCCCACGCCGGCCAA
rhesus             CGAAGAATAGCCGTAGGGGGCGTATGGGACGGTGCGTAGAGCGATGCCCACGCCGGCCAA
hCANXpromoter      CGAAGAATAGCCGTAGGGGGCGTATGGGACGGTGCGTAGAGCGATGCCCACGCCGGCCAA
mouse              CGAAGAATAGCCCTAGGGGGCGTATGGAAGGCGCTTCGAGCGATGGCTGTATCTACCAA
rat                CGAAGAATAGCCCTAGGGGGCGTATGGAAAGAGCTTTGAGCGATGGCCGTATCTACCAA
                   ************ ****************..* **   ******** * . . * .****

chimp              CCGGATGTCGGGGCTTGCGCGGGAGGGCGGGACTTGGCGCCGGCTGTGGCTACTCAGGGG
rhesus             CCGGATGTCGGGGCTTGCGCGGGTGGGCGGGACTTGGCGCCGGCTGTGGCTACTGAGGGG
hCANXpromoter      CCGGATGTCGGGGCTTGCGCGGGAGGGCGGGACTTGGCGCCGGCTGTGGCTACTCAGGGG
mouse              TAAATTCTTAGATCTTGCGCAAAGGGGCGGGATCTGACGTTCGTTTTAGCCACTCAGGAA
rat                TAGATTCTCAGATCTTGCGCAAAGGGACGGGAGTTGACGTTTGTTGTAGCCACTCAGGAA
                    ...:* * .*. *******... **.*****  **.**   * * *.** *** ***..
                                                  EGR
chimp              CCAGGGGCGGGCACAGGGCCGGGCTTCGTGCGGTGGGGCTCGCTCGCGCGGCAGCGGTGG
rhesus             CCAGGGGCGGGCGCAGGGCCGGGCTTAGTGCGGTGGGGCTCGCTCGCGCGGCAGCGGTGT
hCANXpromoter      CCAGGGGCGGGCACAGGGCCGGGCTTCGTGCGGTGGGGCTCGCTCGCGCGGCAGCGGTGG
mouse              CGAGGGGCGGACGCGGGGTTGGGCTTCGTGCGGTGGGGCTCGCTCGCGCGGCGGCCGTAG
rat                CGAGGGGCGGACGCGGGGTTGGGCTTCGTGCGGTGGGGCTCGCTCGCGCGGCGGCGGTAG
                   * ********.*.*.***  ******.*************************.** **.
                                      Xbp1/Creb3l2
chimp              CCGAGGCCTCTTGGTTCTGCGGCACGTGACGGTCGGGCCGCCTCCGCCTCTCTCTTTACT
rhesus             CCGAGGCCTGTTGGTTCTGCGGCACGTGACGGTCGGGCCGCCTCCGCCTCTGTCTTTACT
hCANXpromoter      CCGAGGCCTCTTGGTTCTGCGGCACGTGACGGTCGGGCCGCCTCCGCCTCTCTCTTTACT
mouse              CCGAGGCCTCTTAGTTCTGCGGCACGTGACGGTCGGGCCGCCTCTGCCGCTGTCTCCACT
rat                CCGAGGCCTCTTAGTTCTGCGGCACGTGACGGTCGGGCCGCCTCTGCCGCTGTCTCCACT
                   ********* **.******************************* *** ** ***  ***
```

##### *Calr:*

```
                                                                  Creb3l2
hCalr           CCAGATG----GGCAACGACGCGCGCGGACGAGGGCGGGGTTGGGTTCAGGTCTGGTCAC
chimp           CCAGATG----GGCAACGACGCGCGCGGACGAGGGCGGGGTTGGGTTCAGGTCTGGTCAC
rhesus          CCAGATG----GGGAACGACGCGCACGGGCAAGGGCGGTCTTAGGTTCAGGTCTGGTCAC
mouse           CTCCTAG-CGAGCCAGAGACTCTCAGCAGCAAGGGCGGGGTTGGGCTGAGGTTCAGTCAC
rat             CTTCATGGCGAGCAAGGGACTCTCACCAGCAAGGGCGGGGTTGGGCTGAGGCTCAGTCAC
                *   ::*    *  *. *** * *.  ..*.*******  **.** * ***   .*****

hCalr           ATGACCTGGCCTGAG-GTGCTCGCGGCCCCCACCCCACCAGTGGGCGTCCCCCCC-ACGC
```

```
chimp            ATGACCTGGCCTGAG-GTGCTCGCGGCCCCCACCCCACCAGTGGGCGTCCCCTCCAACGC
rhesus           ATGACCTGGCCTGAG-GGGCTCGCGGCCCCCACCTCACCAGTGGGCGTCCCCCACAACGC
mouse            GTGACCGTGCCTGAGTGGGCTAGCGGCCCCCACCCCACCAGGGGGCGTCCCCCACAACGC
rat              GTGACCGCGCCTGAGTGGGCTCGCGGCCCCCACCCCAACAGGGGGCGTCCCCTACAACGC

                 .*****  ******* * ***.************ **.*** ********** .* ****
                         ERSE I              ERSEII-like
hCalr            GTGGTCGACCATCATTGGTCGGTGGTGAGGCCAATAGAAATCGGCCAT
chimp            GTGGTCGACCATCATTGGTCGGTGGTGAGGCCAATAGAAATCGGCCAT
rhesus           GTGGTCGACCATCATTGGTCAGTGGTGAGGCCAATAGAAATCGGCCAT
mouse            GTGGTCGACCCTCATTGGCCCATAGTGCGACCAATAGAAATCAGCCAT
rat              GTGGTCGACCCTGATTGGCCCAGGGTGCGGCCAATAGAAATCAGCCAT
                 **********.* ***** *  . .***.*.************.*****
```

#### *Dnajb9:*

```
                                                                        Creb3l2
hDnajb9          CGAGCCCCTGCGCCTGCGCTAGCATTCTGCCGGG--AAAGCCGCCTCGTCTGTCGACTCA
rhesus           GGAGCCACTGCGCCTGCGCTGGCATCCG-CCGGG--AACACCGCCTCGTCTATCGACTCA
chimpDnaj        CGAGCCCCTGCGCCTGCGCTAGCATCTG-CCGGG--AAAGCCGCCTCGTCTGTCGACTCA
mouseDNA         CCCTCTAAAGCGCCTGCGCG-CGATGCGCCCCAGGGAAGGATGAGGAAATCGTCGACTCA
rat              CCATTTACAGCGCCTGCGCAGCGATGCGCCCCAGGGAAGGATGAGGAAATCGTCGACTCA
                   .   ..:**********    **    ** .*  ** .. *.  ..:  .********
                  ETS                                                 ETS
hDnajb9          CTTCCGCCTCCTCCGGTTTAAGCACCGCCTTTTCGGGGGTTGAAGCCGGAAGTGGCGCAC
rhesus           CTTCCGCCTCCGCCGATTTAAGCACCGCCTTTTCGGGGGTTGGGGCCGGAAGTGGCGCAC
chimpDnaj        CTTCCGCCTCCTCCGGTTTAAGCACCGCCTTTTCGGGGGTCGAAGCCGGAAGTGGCGCAC
mouseDNA         CTTCCGCTTCCGCACACCTTGGCACCGCCCCTCGGAGGGTGG-CGACGGAAGTGACGCAA
rat              CTTCCGTTTCCTCACATTTAG-CACCGCCCCCTGGAGGGTGG-CAACGGAAGTGACGTAG
                 ******  *** *. .  *:. *******     *.**** *  ..********.** *
                                             Xbp1/Creb3l2              Xbp1
hDnajb9          AGCCCTAGCAGCAACAACAGTTTTCCACGTGCGCGTAGGGCGCCGGGATCACGTGGGGAG
rhesus           GGCCCCAGCGGCAACAACAGTTTTCCACGTGCGCGTAGGGCGCCGGGGTCACGTGGGGAG
chimpDnaj        AGCCCTAGCAGCAACAACAGTTTTCCACGTGCGCGTAGGGCGCCGGGATCACGTGGGGAG
mouseDNA         GGACCAAACGGCAACAACAGTTTTCCACGTGCGCGTAGGGCGCCAAAGTCAGGTGGGCCG
rat              GGATCAAACGGCAACAACAGTTTTCCACGTGCGCGTAGGGCGCCAAAGTCAGGTGGGCCG
                 .*. * *.*.*********************************....*** ***** .*
                       NFYA/B              CEBP
hDnajb9          GCAGAAGCCAATGGGGAAGCGTTTCGTGTAGGTCTTCTG---AGGTGGTGGCGCCAGCGG
rhesus           GCAGAAGCCAATGGGGAAGCGTTTCGTGTAGGTCTTCTG---AGGTGGTGGCGCCAGCGG
chimpDnaj        GCAGAAGCCAATGGGGAAGCGTTTCGTGTAGGTCTTCTG---AGGTGGTGGCGCCAGCGG
mouseDNA         GCGGGAGCCAATGGGGAAGCGTTTCGTGTAGGTCGGCTCTATAAGCGGTCGCGCCAGCCA
rat              ACGGGAGCCAATGGGGAAGCGTTTCGTGTAGGTCTTCTCTAGAGGCGGTTGTGCCAGCAA
                 .*.*.****************************  **     *.* *** * ****** .
```

#### *Pdia3:*

```
                                                               KLF
hPdia3promoter       CCCGCCCCGCGCTCCGGCCACTCGGCGGTAACGAGTTGGTCCCGCCCGCCCGACGAAGGC
chimp                CCCGCCCCGCGCTCCGGCCACTCGGCGGTAACGAGTTGGTCCCGCCCGCCCGACGAAGGC
mouse                CTCGCCCCGCGCGCCCGCCAATCGGCAGTTACCAGCTG-TTCCGCCCGCCCGACGCTG-C
rat                  CTCGCCCCACGCGCCCGCCAATCGGCAGTTACCAGCTGGTTCCGCCCGCCCGACGCAG-C
                     * ******.*** ** ****.*****.**:** ** ** * **************.:* *
                                       ERSE I
hPdia3promoter       GACGCGCAGCCAATCAGCGGCTGCCACACAGCGG--CCCAAGCCGGGTTTGGGGGTTGGG
chimp                GACGCGCAGCCAATCAGCGGCTGCCACACAGCGG--CCCGAGCCGGGTTTGGGGGTTGGG
mouse                GTGACGCAGCCAATCGGCGGCTGCCACACTGCGGGCCCCGAGCAGGCCTAGGGGGTTGGG
rat                  GTGACGCAGCCAATCGGCGGCTGCCACACTGCGGGCCCG-AGCAGGCCTAGGGGGTTGGG
                     *:  .***********.*************:****   **   ***.**   *:**********

hPdia3promoter       ACCTCCGGCTGCAGGTCCGCCTGGGCCAGACGCGCGAGCGCAAGCAGCGGGTTAGTGGTC
chimp                ACCTCCGGCTGCAGGTCCGCCGGGGCCAGACGCGCGAGCGCAAGCAGCGGGTTAGTGGTC
mouse                ACCTCGG-CAGCGGGTCTGCCCGGGCCAGACGCGCGAGCGCAGGCAAGCGGCTGCAGATT
rat                  ACCTCGG-CAGCAGGTCTGCCCGGGCCAGACGCGCGAGCGCAGGCAG-CGGCTGCAGATT
                     ***** * *:**.**** *** ********************.***.   ** *.  :*.*
```

#### *Pdia6:*

```
                          ERSEI
Human            CCGTGGCTGCGGCTCATTGGTCCGGGC
Chimp            CCGTGGCTGCGGCTCATTGGTCCTGGC
Baboon           CCGTGGCTGCGGCTCATTGGCCCTGGT
Orangutan        CCGTGGCTGCGGCTCATTGGTCCTGGA
Mouse            CCGTGGCCGCCTTTCATTGGTCTCTGT
Rat              CCGTGGCCGCCGTTCATTGGTCTCTGT
```

```
                    ******* **    ******* *    *
```

#### *COP II trafficking*

##### ***Mia3:***

```
                                   XBP1/Creb312
hMia3           TGAGGTGCGCCAGGGGCAGTGGCTGATGACGTGTTTTATAGGCGGCATAAAGCGAAAGTG
chimp           TGAGGTGCGCCAGGGGCAGTGGCTGATGACGTGTTTTATAGGCGGCATAAAGCGAAAGTG
rhesus          TGACATGCGCCAGGGGCCGTGGCTGATGACGTGCTTTATAGGCGGCATAAAGCGAAAGTG
mouse           TGACGTCGTTCCTCGGCCGTGTCTGATGACGTGTTTCACCGGAGGCCTAGTGCGGAAGGT
rat             TGACGTTGCTCCTCGGCCGTGTCTGATGACGTGTTTTACCGGAGGCCTAGTGCGGAAGGT
                *** .*    *.  ***.*** *********** ** *  .**.***.**.:***.***
                            ETS
hMia3           CAGGCAGCTGTGGAAGGCGAAGTTCAATCCCAGAGTCCG--------CCCCCTGAATTGG
chimp           CAGGCAGCAGTGGAAGGCGAAGTTCAATCCCAGAGTCCG--------CCCCCTGAACTGG
rhesus          CCGGCAGCGGTGGAAGGCGAAGTTCAATCCCAGAGTCCG--------CCCCCTGAACTGG
mouse           GCCGCGGCCGGGGAAGGCGAAGTTCATTCCCCGTGTCCG--------CCCCCCGGAGCCG
rat             TCCGCAGCCCGGGAAGGCGAAGTTCATTCCCAGTTTCCGCCCCCCCCCCCCCCGGAGCCG
                 . **.**   ***************:****.*.: ****        ***** *.*   *
                               ETS
hMia3           GGCCTTTCCGGAGGAGGAAGCTCTGAAAAACAGGGGGCCCAGTGCCATTCCGCAGGGAAT
chimp           GGCCTTTCCGGAGGAGGAAGCTCTGAAAAACAGGGGGCCCAGTGCCATTCCGCAGGGAAT
rhesus          GGCCTTTCCGCAGGAGGAAGCTCTGAAAGACAGGGGACCCAGTGCCATTCCCCTGGGAAT
mouse           GTCCTTCCCG-AGGCGGAAGCGGCGAAAGCCGG----CTCGGTGCCG-TCCGCAGG-AAG
rat             GTCCTTCCCG-AGGCGGAAGCGGTGAAAGCCGG----CTCGGTGCTG-TCCGCAGG-AAA
                * **** *** ***.******   ****..*.*    * *.****  .  *** *:** **
                             E-box
hMia3           TGTCGCTTGCGTTCAGCTG--TTCTACACAATGGACTCAGTACCTGCC
chimp           TGTCGCTTGCGTTCAGCTG--TTCTACACAATGGACTCAGTACCTGCC
rhesus          TGTCGCTTGCATTCAGCTG--TTCTACACAATGGACTCAGTACCTGCC
mouse           CGTCGTCTCCGCGCAGCTGCCCCCCATACGATGGACTCACTGCCTGCC
rat             CGTCGTCTCCGCGCAGCTG-CCCCCACACCATGGACTCATTGCCTGCC
                 ****   * *.  ******    * * ** ********* *.******
```

##### ***Lman1:***

```
                               Xbp1/Creb312
hLman1intron      TCATGTTTCACAACGCCACGTCTCTAGGAGAAATG--TGCCCTACCCCCAGCCATCCCAA
chimp             TCATGTTTCACAACGCCACGTCTCTAGGAGAAATG--TGCCCTACCCCCAGCCATCCCAA
rhesus            TCATGTTTCACAACGCCACGTCTCTAGGAGAAATG--TGCCCTACCCCCGGTCATCCTAA
mouse             CCTTACACCCTAATGCCACGTCTCATGGG--------TACACTTTCCGTGGCCATAGCAA
rat               CCCTACACCCGAATGCCACGTCTCAAGGG--------TACACTTTCCGTGGCCATAGCAA
                   * *. : *. ** **********::**.        *.*.**: **  .* ***.  **
                              CEBP
hLman1intron      CCTGAGACATGTTCTAGAAGCCCCC
chimp             CCTGAGACATGTTCTAGAAGCCCCC
rhesus            CCTGAGACATGTTCTAGAAGCCCCC
mouse             TCTGAGACCAGTTCCAACAGCCTCC
rat               TTTGAGACCAGTTCCAACAGCCTCC
                    ******.:**** *..**** **
```

##### ***Copb2:***

```
                          ETS                 ETS        KLF
hCopb2            CGG----TCTGGAAGGCCACGTTGCCTAGGAAACTGGGGCGGGGCTTGGGAAACGACGGC
rat               AGA----CGCGGAAGGCCGCGTTACCTAGGAAACTGGGGCGGGACTACG-CGCAGATTGC
chimp             CGG----TCTGGAAGGCCACGTTGCCTAGGAAACTGGGGCGGGGCTTGGGAAACGGCGGC
rhesus            CGG----TCTGGAAGGCCACGTTGCCTAGGAAACTGGGGCGGGGCTTGGGAAAGGATGGG
mouse             CGGGAGACGCGGAAGTCCGCGTTACCTAGGAAACTGGGGCGGGACCATG-CACGGACCGC
                  .*.       ***** **.****.******************.*  :  *  ...  *.  *

hCopb2            CCTGCGCGCAGTTTGCGCAGGCGCTCCGGCCCTGGCCTCTGACTATCGCGAGAATCGGCT
rat               CTTACGGAGAATTTGCGCAGGCGCAGAG--CCTG-GTTCCGAGTGTCGCGAGAGCCGCTT
chimp             CCTGCCCGCAGTTTGCGCAGGCGCTCCGGCCCTGGCCTCTGACTATCGCGAGAATCGGCT
rhesus            TCTGCGCGCCGTTTGCGCAGGCGCTTCGGCCCTGGCGTCTGACTATCGCGAGAATCGGCT
mouse             CTTACGGAGGGTGTGCGCAGGCGCAGAG--CCTGGC-TCCGAGTACCGCGAGAGTCGCTT
                    *.*   .  .* ***********: .*   ****    ** ** *. *******. **  *
                        ETS      XBP1/Creb312
hCopb2            CCCGGAAGTTCCACGTCAGTCAGTCTGACGGTCAGTGGATCGGTGGGTTTATCTCAAGGC
rat               CCCGGAAGTTCCACGTCAGTGGGTCCACTGCTCCGTGGGTCTGTGTGTTCTTG--AGTAC
```

```
chimp           CCCGGAAGTTCCACGTCAGTCAGTCTGACGGTCAGTGAATCGGTGGGTTTATCTCAAGGC
rhesus          CCCGGAAGTTCCACGTCAGTCAGTCTGACGGTCAGAGGTTCGGTGGCTTTCTCTCAGGGC
mouse           CCCGGAAGTTCCACGTCAGTGGGTTGGCCGGTCCACGGGTCGGTGGGTTCCTGAGTGTGT
                ******************** .**  .. * **.. *. ** ***  **  *   :. .
```

***Sec13:***

```
                                             XBP1/CREB3L2
hSec13          TGAGCTGTTCCGAGGCGCCGCCGGGAGCTGCCACGTCCGAGACCTGGAGCAGCCACCGCC
chimp           TGGGCTGTTCTGTGGCGCCGCCGGGAGCTGCCACGTCCGTGTCCTGGAGCAGCCACCGCC
rhesus          TGAGCTATTCGGAGGCGCCGCCGGGAGCTGCCACGTCCGAGACCCGGAGCAGCCACCGCC
mouse           TGAAGCGTTTTGAG-------------CTGCCACGTCCGAGGCCGTGGGTCGCCGCTGCT
rat             TGAAGCGTTCTGAG-------------CTGCCACGTCCGAGGCTGTGAGTCGCCGCTGCT
                **..  .**  *:*             ************:* *   *.* .***.* **

chimp           GCAATCATGGTGAGTGTTGGTACTGAGGAGGCCTAGCAGGGTGAAGGCCACGGTCGCGG
rhesus          ACAATCATGGTGAGTGTTGGGACTGAGGAGGCCGAGCAGGGTGAAGGCCACGGTCGCAG
hSec13          GCAATCATGGTGAGTGTTGGTACTGAGGAGGCCTAGCAGGGTGAAGGCCACGGTCGCGG
mouse           GCAGCTATGGTAAGTGTCG-------------CGATCCCTGCGATAGTCGTGG------
rat             GTAGCTATGGTAAGTGATG-------------CGGTCCCTGAGATAGTCGTGG------
                . *.  *****.****: *             * . *.  * **:.* *. **
```

***Sec16a:***

```
                                 XBP1/Creb3l2
HumanSec16a     GGCCGCCGCGCCGCCGACGTGTCCGGCT
Baboon          GGCCGCCGCGCCGCCGACGTGTCCGGCT
GreenMonkey     GGCCGCCGCGCCGCCGACGTGTCCGGCT
Mouse           AGGTGCCG--CCGCCGACGTGTCCGAA-
Rat             AGGTGCCG--CCGCCGACGTGTCCGAC-
                .*  ****  ***************..
```

***Sec23ip:***

```
                     Xbp1/Creb3l2
hSec23ip        GGTGTCACGTGCTAGACGCGGGCGTGAAGACAGTGTCGCCATGTTGATATCTGGCGTCGC
chimp           GGTGTCACGTGCTAGACGCGGGCGTGAAGACAGTGTCGCCATGTTGATATCTGGCGGCGC
rhesus          GGTGTCACGTGCCAGACTCGGGCGGGAAGACATTGCCGCCATGTTGATTTCTGGCGGCGC
mouse           GCTGTCACGTGATAGGCTTGGGTCTGAAGCCGATTCCACCTTATTGATTTTGGGCGGCGT
rat             GGTGTCACGTGATAGGCTTGGGTCTGAAGCCGATTCCACCTTATTGATTTTGGGCGACG
                * *********. **.*  ***   ****.*. *  *.**:*.*****:*  **** **

hSec23ip        CGTGTAGGGGAAGGGAGAGGGCGGCT----GACTAGCTACCCGTAATTTCGGCTCCTGTG
chimp           CAGGTAGGGGAAGGGAGAGGGCGGCT----GACTAGCTACCCGTAGTTTCGGCTCCTGTG
rhesus          TGGCTGGGGGAAGGGAGAGGGTGGCTGACTAACTAGCTACCCGTAGTTCCGGCTCCTGTG
mouse           CAAATTCCGGAAGAAAACCGATCCCG----GAGTCGCTGGCCTGTG-----GCACCTGTG
rat             CAGCTTCCGGAAGAAAATCGGTCCCT----TAGTCGCTGTCCTGTG-----GTCACCGTG
                 .  *   *****..*.  *.   *      * *.***. **  :.     *  .* ***
                          ETS                            ETS
hSec23ip        CGCAAGTGCGGAAGTGCGCTGGCCAGGAGCGTGATCGGTTTCCGGTCAGTGGTGTGGTAC
chimp           CGCAAGTGCGGAAGTGCGCTGGCCAGGAGCGTGATCGGTTTCCGGTCAGTGGTGCGGTAC
rhesus          CGCAAGTGCGGAAGTGCGCTGGCCGGGAGAGTGATCGGTTTCCGGTCAGTGGTGCGGTAC
mouse           CGCAAGTGCGGAAGTGCGGTGGCATCCTCGCTAACCGGTTTCCGGTCTGTG-TGCAGCAC
rat             CGCAGGTGCGGAAGTGCGGTGGCATGCTCAG-GACCGGTTTCCGGTCTGTG-TGAGGCAC
                ****.************* ****.   :    .* ************:*** ** .* **
```

***Sec24d:***

```
                        ETS                Xbp1/Creb3l2 ETS
hSec24d         GAATCAGCTTCCTCACGCAACCCAGTTTGCCACGTCAAACTTCCTCACACAGAGCACCAA
chimp           GAATCAGCTTCCTCACGCAACCCAGTTTGCCACGTCAAACTTCCTCACACAGAGCACCAA
rhesus          GAATCAGCTTCCTCACGCTACCCAGTTTGCCACGTCAAACTTCCTCACACAGAGCACCAA
rat             GAATTTACTTCCTCGCCCTACTC----TGCCACGTCAAACTTCCTCACCCAGAACACCAA
mouse           GAATTCGCTTCCTCACCCTTCTC----TGCCACGTCAAACTTCCTCACCCAGTACACCAA
                ****  .*******.* *::* *    *********************.***:.******
                            Creb3l2                            E-box
hSec24d         GTTAAAGAAACCTGACTCAGCTTCAAGCGGCGACACGGCAGAAAGCCACTTGGTATAAGC
chimp           GTTAAAGAAACCTGACTCAGCTTCAAGCGGCGACACGGCAGAAAGCCACTTGGTATGAGC
rhesus          GTTAAAGAAACCTGACTCAGCTTCAAGCGGCGACATGGCAGAAAGACACTTGGCATGAGC
```

```
rat        GTTGCAGAAACCTGACTCAGCTGCAAGCGGCGACATGGCAGAGAGCCACTTGGCACGAAC
mouse      GTTGCAGAAACCTGACTCAGCTGCAAGCGGCGACATGGCAGAGAGCCACTTGGCACAAAC
           ***..***************** ************ ******.**.******* *  .*.*
```

***Uso1:***

```
hUSO1      ACCCTGCAAGTCGAAACGCGTAGTCTAATCAAATCCTCTCCAGCCTGCAGGGCCGGGCTT
rhesus     ACCCTGCAAGTCGAAACGCGTAGTCTAATCAAATCCTCTCCAGCCTGCAGGGCCGGGCTT
chimp      ACCCTGCAAGTCGAAACGCGTAGTCTAATCAAATCCTCTCCAGCCTGCAGGGCCGGGCTT
mouse      -ACCTACCAGCCGATGCTCCTAGTCTAATCAGAGCCTCGGGAGCCGACTGGGCAGGGCTG
rat        -ACCTACCAGCCCAAGCGCGTAGTCTAATCTGAGCCTCCGGAGCCGACTGGGCAGGGCTG
            .***.*.** * *:.* * **********:.* ****    **** .*:****.*****

                 XBP1/Creb3l2
hUSO1      CCACCACGTGGGTGGCTGCGGCCAGCAGGATCAATCCGCGAAGGGTGGGGTAGGAGGCG
rhesus     CGACCACGTGGGTGGCTGCGGCCAGCAGGATCAATCCGCGAAGGGCGGG---AGGAGGCG
chimp      CCACCACGTGGGTGGCTGCGGCCAGCAGGATCAATCCGCGAAGGGTGGGGTAGGAGGCG
mouse      CGACCACGTGGGTGACCGCGGCCAGCAGCAGCAAGCAGCGCAAGGAGGAGG-AGGGGGCG
rat        CGACCACGTGGGTGACCGCGACCAGCGGCAGCGAGCAGAGCAAGGAGGAGG-AGGGGGCG
           * ************.* ***.*****.* * *.* *.*.*.*.** **.    ***.****
```

***Stx5a:***

```
                                                    Xbp1/Creb3L2
HumanStx5  CTGAAGCCGCCGAAACCCGACCAAAGACTGGAAGCGGCCACGTCACCTACACGTAACCCC
Chimp      CTGAAGCCGCCGAAACCCGACCGAAGACTGGAAGCAGCCACGTCACCTACACGTAACCCC
Rhesus     CTGAGGCCGCCGAAACCCGACC--AGACTGGAAGCGGCCACGTCACCTACACGTAACCCC
Mouse      TGTAAGCAGTGGAAGCCGGACCAAAGAACAGGAGCCGCCACGTCACCTAGACGTACTCCT
Rat        TGTAAGCAGTGGAAGCCGGACCAAAGAACTGGAGCCGCCACGTCACCTACACGTACCCCT
              *.**.*  ***.** ****  ***.  *.*** ************ *****. **
```

***Bet1:***

```
                                     Stat3/5
hBet1      GTTATGCG----TTGGTGAACTACATTTCCCAGAATGCCTTTC-AACCTTCTTCCCAGGG
chimp      GTTATGCG----TTGGTGAACTACATTTCCCAGAATGCCTTTC-AACCTTCTTCCCAGGG
rhesus     TTTATGCG----TCGGTGAACTACATTTCCCAGAATGCCATTC-AACCTTCTTCCCAGGG
mouse      GTGCCCCAGAAGACGGAAAACTACATTTCCCAGAATGCAACTAGTAATCGCATTTCAGGA
rat        GTGCCCCAGAAGCCCGAAGACTACATTTCCCAGAATGCAACTA-AAATCTCATTGCAGGG
            *  .  *.        *:..*******************.: *. :*.    *:*  ****.

                        CREB3L2                                Creb3l2
hBet1      CTCCAATGAGCGCCGTGGTGACGCCATCATCCGGCGCGGTGTTTAGACTCCACTGATGTC
chimp      CTCCAATGAGCGCCGTGGTGACGCCATCATCCGGCGCGGTGGTTAGACTCCACTGATGTC
rhesus     CTCCAATGAGCGCCCTGGTGACGCCATCAGCAGGCGCGGTGTTTAGACTACACTGATGTC
mouse      CGCCTATATGTGTCCGAGTGACGACATAAACCGGCGCGGTGTTTAGACTTTACTGATGTC
rat        CGCCTATATGTGTCCGGGTGACGACATAAACCGGCGCGGTGTTTAGACTTTACTGATGTC
           * **:**.:* * *  .******.***.* *.********* *******  *********

hBet1      GTGGCGCTTTAGGGGAAGAAGTTGGTGTTTCGCTGGGCCCTGGTACTGAAGACGCGGTCC
chimp      GTGGCGCTTTAGGGGAAGAAGTTGGTGTTTCGCTGGGCCCTGGTACTGAAGACGCGGTCC
rhesus     GTGGCGCTTTAGGGGAAGAAGTTGGTGTTTCGCCCGGCCCTGGTACTGAAGACGCGGTCG
mouse      GTGGCGCTCTGAGGCAAAAAGTTGTTGCTCTTAGGAGCCCTGGGACTGAAGCCGAGGTCG
rat        GTGGCGCTCTGAGGCAAAAAGTTGTTTCTCTGAGGAGCCTTAGGACTGAAGCCGAGGTTG
           ******** *..** **.****** *   *    .  .*** *.* *******.**.***
```

*COP I Trafficking*

***Kderlr1:***

```
                                                        Creb3l2
hKdelr1Intron  CCGCAGTCTGGACCTGAGGTGAGGCCTAGGGAGGACCTTCCCTGACCTCAGGCCCCGGAT
rhesus         CTTCAGTCCGGACCTGAGGTGGGGCCTAGGGAGGACCTCCCCTGACCTCAGCCCCCCGAT
chimp          CCGCAGTCTGGACCTGAGGTGAGGCCTAGGGAGGACCTTCCCTGACCTCAGGCCCCGGAT
mouse          TGACAGTCTAGACCTACACTGGGACCTAGGGAGAGCCCTGCCTGACTTCAGCCTCCCAGT
rat            TGACAGTCTGGACCTACATTGGGACTTAGGAAAAGCCCTGCCTGACTTCAGCCTCCCAGT
                 ***** .*****... **.*.* ****.*...**   ****** **** * **  ..*

                                      ETS          XBP1/Creb3l2
hKdelr1Intron  CCGGTCACCTCCCTCCCGTTCCCTTCCTGGCCG-CCACGGTCCACGTCACCAACTCACAG
rhesus         CTGGTCACCTTCCTCCCGTTCCCTTCCTGGCCG-CCACGGTCCACGTCACCAACTCACAG
chimp          CCGGTCACCTCCCTCCCGTTCCCTTCCTGGCCG-CCACGGTCCACGTCACCAACTCACAG
mouse          CG-GTCACCTTCCTCCCGCTCCCTTCCTGGCCTTCCACAGTCCACGTCACCAACTCCCAG
rat            CG-ATCACCTTCCTCCGGCTCCCTTCCTGGCCTTCCACAGTCCACGTCACCAACTCCCAG
               *  .****** ***** * ************  ****.****************.***
```

```
                             KLF
hKdelr1Intron       CCACCTCCCCGCCTGCCATGGTTCCCTGTTCCCGGTCCATGAACCCTCG-GGCTGCCCAG
rhesus              CCACCTCCCCGCCTGCCATGGTTCCCTGTTCCCGGTCCGTGAACCCTCG-GGCTGCCCAG
chimp               CCACCTCCCCGCCTGCCATGGTTCCCTGTTCCCGGTCCGTGAACCCTCG-GGCTGCCCAG
mouse               CCACCTCCCCGCCTGCCATGGCTCCTGGTTCCCAGCCCCGGAACACCCAGAGCTGTCCCA
rat                 CCACCTCCCCGCCTGCCATGGCTCCTGGTTCCCAGCCCCGGAACACCCATAGCTGTCCCA
                    ********************* ***   ******.* **   ****.* *.  .**** **..
```

***Kdelr2:***

```
hKdelr2          TTAATGCTGCTTTGGGAAACGATCTGGTTTTTAA---------GTGCCTCCATCTAATAA
chimp            TTAATGCTGCTTTGGGAAACGATCTGGTTTTTAA---------GTGCCTCTATCTAATAA
rhesus           TTAATGTTGCTTTGGGAAACGATCTGGTTTGTCAAACTCTTAAGTGCCTCCATCTAATAA
mouse            TTCATGTTACTCTGAGAAGCAAGCTAGTTTTTCATCCTTG--AGTGTGGCCGTCTAATAA
rat              TTTATGCTACTCTGAGAAGCAAGCTAGTTTTTCATTCTTG--AGTGTGGCCGTTTAATAA
                 ** *** *.** **.***.*.* **.**** *.*         ***   *  .* ******

mouse            CTGGTTCTCTTAAACTGCCAGCGCTGGG-TACCACAGAGGCCTCTGTTCTCGGCTAGTA-
rat              CTGGTTCTCTTAAACTGCAAGCACTGGG-TACCACTGAGGTCTCTGCTCTCATCTAATG-
rhesus           TGGGTTCTCTTAAAACTGTAGCGCCCGGGGGCCGCCGAGGCCTCCTATGGGTTCCTGCAG
hKdelr2          TGGGTTCTCTTAAAACTGCGGCGCCCGGGGACCGCCGAGGCCTCCTATGGGCTCCTGCAA
chimp            TGGGTTCTCTTAAAACTGCGGCGCCCGGGGACCGCCGAGGCCTCCTATGGGCTCCTGCAA
                   ************.     .**.*  **   .**.* **** ***    *      * :. .
                                 Xbp1/Creb3l2
mouse            -TCCTTGCTCCACGTCACCATTGAGTTGGCGCTGTACCTCCCTCAAACTACAT-------
rat              -TCCTGGCTCCACGTCACCAGTGAGTTGGCGCTGCACCTCCCTCGAACTACATGCATATT
rhesus           TTTCTTACTCCACGTCACCAGTGAGTTGGCTGCTGCCCTGTTCAGGAGTACAG----ATG
hKdelr2          TTCGTTGCTCCACGTCACCAGTGAGCTGGCTGGTTCCCTGATCAAGAGTACAG----ATT
chimp            TTCGTTGCTCCACGTCACCAGTGACCTGGCTGGTTCCCTGATCAAGAGTACAG----ATT
                  *  *  .************* ***   ****      .***     ...* ****
```

***Kdelr3:***

```
                                    ETS                             XBP1/Creb3l2
hKelr3promoter    GATCTCCGGGCCGCGCCGCTTCCTGGCTCCCCACCCTGCGCCGGCGGCCGCCCTGGCCAC
chimp             GATCTCCGGGCCGCGCCGCTTCCTGGCTCCCCACCCTGCGCCGGCGGCCGCCCTGGCCAC
rhesus            GAACTCCAGGCCGCGCCGCTTCCTGGCTCCCCACCCTGCGCCGGCGGCCGCCCTGGCCAC
mouse             -------------CGCTGCTTCCTGGCTCCTGTCCCG-CGCCT--GGCCGCCCC-GCCAC
rat               GATCTCCCGGCGGCGCTGCTTCCTGGCTCTCCACCCTGTGCCCG--GCCGGCCACGCCAC
                               *** ************    :***    ***     **** **   *****

hKelr3promoter    GTCACCGCCCCGGCCAAGAGTGCGTGGGCGGCGGCGCG-CGGGTGCGATCGCGGAGCTGTG
chimp             GTCACCGCCCCGGCCAAGAGTGCGTGGGCGGCGGCGCG-CGGGTGCGATCGCGGAGCTGTG
rhesus            GTCACCGCCCCGGCCAAGAGTGTGTGGGCGGCGGCACGGCGGGTGCGAACGCGGAGCTGTG
mouse             GTCACCGCGCGGCCAAGGGTGCG----------CGCGGCGGGAGCGCGCCAG---CAAGG
rat               GTCACCGCCCCGGCG----------------------GCCGGGAGCG--CGCGGAGCCGGT
                  ******** ****                        *  ****:***   *  .*    *  .
                                                             XBP1/Creb3l2
hKelr3promoter    AGGCGCAGGCAGGGCTCTGGGGCACCTAGAG-ACCGGGGCCGGAGACGTGGCAGCCGCCC
chimp             AGGCGCAGGCAGGGCTCTGGGACACCTAGAG-ACCGGGGCCGGAGACGTGGCAGCCGCCC
rhesus            AGGCGCAGGCAGGGCTCTAGGGCACCTAGAG-ACCAGGGCCGAGGACGTGGCAGCCGCCC
mouse             AGGCGCGG-CACCGGCCTGGGGCACCGCGAGGAGCAGGGACGGGGACGTGGCAGCG-CCC
rat               AGGCGCGGGCTGG-GCCTGGGGGGCTGCGAG-GCCGCGGCCGGGGACGTGGCAGCCGCCC
                  ******.* *:     **.**.  .*   .***  . *.  **.**..***********   ***
                          NF-KB
hKelr3promoter    TGCCCGCCAGAAAGTTTCCTAGAAG--TTTGCT-GGGCGCGGGCGCACGACTGACTGGCT
chimp             TGCCCGCCAGAAAGTTTCCTAGAAG--TTTGCT-GGGCGCGGGCGCACGACTGACTGGCT
rhesus            TGCCCGCTAGAAAGTTTCCTAGAAG--TTTGCT-GAGCGCGGGCGCACGACCGACGGGCC
mouse             CGCGCAACAGAAAGTTCCCTGGGAAGTTCTGCG-CGACACGG-CGCGGGAGCGGCGGGCG
rat               TACGCATCAGAAAGTTCCCTGAAAG--TTCGCTGCTGGGCACGCACAGGACCGGCGGGCT
                   .* *.   ******** ***...*.   *   **     .  .*.   *.*.  **   *.* ***

hKelr3promoter    GGACCATGAACGTGTTCCGAATCCTCGGCGACCTGAGCCACCTCCTGGCCATGATCTTGC
chimp             GGACCATGAACGTGTTCCGAATCCTCGGCGACCTGAGCCACCTCCTGGCCATGATCTTGC
rhesus            GGACCATGAACGTGTTCCGAATCCTCGGCGACCTAAGCCACCTCCTGGCCATGATCTTGC
mouse             GGACCATGAACGTGTTCCGAATCCTCGGGGACCTGAGCCACCTCCTGGCTATGATCTTGC
rat               GGGCAATGAACGCGTTCCGAATCCTCGGCGATCTGAGCCACCTCCTGGCCATGATCTTGC
                  **.*.******* *************** ** **.************* *********
```

***Scyl1:***

```
                                                                  XBP1/Creb3l2
hScyl1intron     TGGGCCTTGCCTATTCCGCGCCGCCGACCCCGTCGCGTTGCGCCCGGCCTGACGTGGCGG
chimp            TGGGCCTTGCCTATTCCGCGCCGCCGGCCCCGTCGCGTCGCGCCCGGCCTGACGTGGCGG
greenMonkey      TGGGCCTTGCCTAGTCCGCGCCGCCGGCCCCGCCGTGTCGCGCCCGGCCTGACGTGGCGG
```

```
rat              TCGGCCTTTCCTAGTCTGAATG-CCGGCTTCGCTGTGTCGCGCCGGGCCTGACGTGGCGG
mouse            TCGGCCTTTCCTACTATGTATG-CCGGCTTCGCTGTGTCGCGCCGGGCCTGACGTGGCGG
                 * ****** **** *. *  .   ***.*  **  * ** ***** ***************
                                                     KLF              Xbp1/Creb3l2
hScyl1intron     CGGAACCAAG-----GGGTAGCAGGCCGGGGAG-GGGGCGGGCAGTGCCTACGTGGCGGG
chimp            CGGAACCAAG-----GGGTAGCAGGCCGGGGAG-GGGGCGGGCAGTGCCTACGTGGCGGG
greenMonkey      CGGAGCCTCG-----GGGTAGCGGGCCGGGGAG-GGGGCGGGCAGTGCCTACGTGGCGGG
rat              CG--ACCAAAGATCCCGGCAACGGGCTGGGGAGTGGGGCGACCAGTGACGACGTGGCGGG
mouse            CG--ACCGAGAATCCCGGCAACGGGCTGGGGAGCGGGGCGAGCAGTGACGACGTGGCGGG
                 **   .** ..      ** *.*.*** ****** ******. *****.* **********
                   KLF                             XBP1/Creb3l2
hScyl1intron     CGGAGGGACCGAGGGACCGACAGGCGGACAGGGCCGGGGTCACGTGGCCCGGCGAAGTG
chimp            CGGAGGGACCGCGGGACCGACAGGCGGACAGGGCCGGGGTCACGTGGCCCGGCGAAGTG
greenMonkey      CGGAGGGACCGAGGGACCGACAGGCGGACAGGGCCGGGGTCACGTGGCCCGGCGAAGTG
rat              CGGAAGAACCGAGGGATGGACAAGCAAACCGGACGGGGGTCACGTGGTCTGGCTAG--A
mouse            CGGAAGGACCGAGGGACGGACAAGCAAACCGGACGGGGGTCACGTGGCCTGGCTAG--C
                 ****.*.****.****  ****.**..**.**.* ************* * *** *.
```

***Surf4:***

```
                                        ETS          SP1
hSurf4       CCGCACCCGCTGCGGCCTCCACAGGAAGTGCCCGGCGGCCGGCCTCGCTCCGCGTCGGCT
chimp        CCGCACCCGCTGCGGCCTCCACAGGAAGTGCCCGGCGGCCGGCCTCGCTCCGCGTCGGCT
mouse        TCGCGCCTGCTGCGGCCTCCACAGGAAGTACCCGGCGGCCGGCCTCGCTGAGCCGCGGCT
rat          TCGCGCCTGCTGCGGCCTCCACAGGAAGTACCCGGCGGCCGGCCTCGCTGAGCCGCGGCT
monkey       CCGCGCCCGCTGCGGCCTCCACAGGAAGTGCCCGGCGGCCGGCCTCGCTCCGCGTCGGCT
              ***.** *********************.*******************  .**   *****
                                XBP1/Creb3l2
hSurf4       GCGGCTCCAGCGGCTGCCACGTAGGCCAAGCCTTAAAGGGGCCG
chimp        GCGGCTCCAGCGGCTGCCACGTAGGCCAAGCCTTAAAGGGGCCG
mouse        ---GCTCTAGCGGCTGCCACGTAGGCCAAGCCTTAAAGGACCG
rat          ---GCTCTAGCGGCTGCCACGTAGGCCAAGCCTTAAAGGGGCCG
monkey       GCGGCTCCAGCGGCTGCCACGTAGGCCAAGCCTTAAAGGGGCCG
                **** *******************************.***
```

***Arf3:***

```
hArf3-3'     GCCTCCCCTCCTTTTTCCTGTCCACCTATATGACCAATCCCTAATTGCTGTCCTGATGAT
chimp        TCCTCCCCTCCTTTTTCCTGTCCACCTATATGACCAATCCCTAATTGCTGTCCTGATGAT
rhesus       GCCTCCCCTCCTTTTTCCTGTCCACCTATATGACCAATCCCTAATTGCTGTCCTGATGAT
rat          GCCTCCCGCCCCCTTCCTTGTGCACTGACACGGTCAG-CCCCCACTGCTGTCCTGATGAT
mouse        TCCTCCCTCCTCCCTCCCTGTACACTGACACGGTCAGCCCCCAACTGCTGTCCTGATGAT
              ******  *     * * *** ***   *  *  *. **. ***  .* ***************
            Creb3l2 NF-YA/B
hArf3-3'     GAGTCATTCCAATTTACTGGATTTAAAACAAAAAG----TTGTTACGGTTTCATGGGGTG
chimp        GAGTCATTCCAATTTACTGGATTTAAAACAAAAACAAAGTTGTTACGGTTTCATGGGGTG
rhesus       GAGTCATTCCAATTTACTGGATTTTAAACAAAAACAAAGTTGTTATGGTTTCATGGGGTG
rat          GAGTCATTCCAATCTACTGGGTTTAAACCAGAAGG-AGGCTGTTATGGTTTCGTGGGGT-
mouse        GAATCATTCCAATTTACTGGATTTAAAACAGAATG-AAGCTGTTACGGTTTCGTGGGGTG
             **.********** ******.***:**.**.**       ***** ******.******

hArf3-3'     GCCCCTCTCTCTCTCACCCTCTGGGTTTCGGGAGGTCGAGTGGG---GTATTCTCT----
chimp        GCCCCTCTCTCTCTCACCCTCTGGGTTTCGGGAGGTCGAGTGGG---GTATTCTCT----
rhesus       GCCCCTCTCTCTCTCACCCTCTGGGTTTCGGGAGGTCGAGTGGG---GTATTCTCT----
rat          ----------------CCCCCTTGGTTTCAGGAGGAAGAGTGGGGACGTTTCCTCTGTTT
mouse        G--------------TCCCCATTGGTTTCAGGAGGAAGAGTGGG---GTGTTCTCTGTTT
                             ***  .* ******.*****:.*******    ** * ****
                                                 E-box
hArf3-3'     -TTGTTTGGCAGTGGTTGCTGTCCTCTTGGCATCTGAGTCCTTTCCCTGTCCCCACAAGC
chimp        -TTGTTTGGCAGTGGTTGCTGTCCTCTTGGCATCTGAGTCCTTTCCCTGTCCCCACAAGC
rhesus       -TTGTTTGGCAGTGGTTGCTGTCCTCTTGGCATCTGAATCCTTTCTCTGTCCCCACAAGC
rat          GCTGTTTGGCACTGGTTGCTGTGCTCTTGGCATCTGAGTCCCTTCCCTGAGTCCCTGG-C
mouse        GCTGTTTGGCACTGGTTGCTGTGCTCTTGGCATCTGAGTCCCTTCCCTGTGTCCCTGG-C
               ********* ********** **************.*** *** ***:   **.  .. *
```

***Copb1:***

```
                      Xbp1/Creb3l2
HumanCopb1       AGTCCTTGGCTGACGTGGAAGGGGGTGGGGTAACGACCGAAGCCAC
```

```
Chimp        AGTCCTTGGCTGACGTGGAAGTGGGTGGGGTAACAACCGAAGCCAC
Rhesus       AGTCCTTGGCTGACGTGGAAGGGGGTGGGGTAACAACCAAAGCCAC
Mouse        AGTCCTTGGCTGACGTGGAAGGGGGCGGGGAGACCAGCAAGGATGC
Rat          AGTCCTTGGCTGACGTGGAAGGAGGCGGGGAGG--AACTAGGATGC
             ********************  .** ****:..  * * *.*. .*
```

***Cope:***

```
hCope        ATTGGCTTACTTCGCCGCCGCTCGGAAAAAGTAAACTACATTTCCTAGCGTGCCC-GTG-
chimp        ATTGGCTTACTTCGCCGCCGCTCGGAAAAAGTAAACTACATTTCCCAGCGTGCCC-GTGT
rhesus       ATTGGCTTACTTCGCCGCCACTCGGAAAAAGTAAACTACATTTCCCAGCGTGCCC-GTGA
mouse        -TTCTTTTTCATCGCCTCTGATTTGTCCCATAGAACTACGTTTCCCAGAGTTCCTGACGA
rat          -TTCTGTTTCAACGCCTCCTATTGGTCCCATAGAACTACGTTTCCCATAGTTCCCGGCGG
              **   **:*::**** *  .*  *:...* :.******.***** * .** **  . *
                       ETS   Xbp1/Creb3l2
hCope        -----TCTTGCTTCCGGCTGACGTGTCTTTCAGGAAGAGGAGCTGGTGAGAAGACAGCGA
chimp        AGGTGTCTTGCTTCCGGCCGACGTGTCTTTCCGGAAGAGGAGCTGGTGAGAAGACAGCGA
rhesus       AGGTGTCTTGCTTCCGGCCGACGTGTCTTTCAGGAAGAGGAGCCGGTGAGAAGACAGCGA
mouse        CGGGGACCTGCTTCCGGCCTACGTGTC--CCAG-AAGAAGGTTGGTTAGAGGTGCGGTGA
rat          CAGGTCCCTGCTTCCGGCCTACGTGTCTCCCAGGAAGAAGACCGGTTAGAGGGGTGGTGA
                   * **********  *******   *.* ****.*.   * *..... . .* **

hCope        AATGGCGCCTCCGGCCCCCGGCCCGGCCTCC
chimp        AATGGCGCCTCCGGCCCCCGGCCCGGCCTCT
rhesus       CATGGCGCCTCCGGCCCCCGGCCCGGCGTCC
mouse        CATGGCTCCTCCGGTTCCTGGCGCGGTCTCT
```

*ERAD*

***Edem2:***

```
                                                  XBP1/Creb3l2
hEdem2       GATGGCCAAGAG-GTGATGGAAAGCACAGT------GTGACGTGTCTGGGTGTGAACAGT
chimp        GATGGCCAAGAG-GTGATGGAAAGCACAGT------GTGACGTGTCTGGGTGTGAACAGT
rhesus       GATGGCCAAGAGGCTGGTGGAAAGCACAGT------GTGACGTGTCTGGGTGTGAACAGT
mouse        GGTGGCCAAGAAGCTAGCGGAGTGCATGCAGCCGCAGTCACGTGTCTGGGCGTGAGAT--
rat          --------AGGAGCTAGCCGAGTGCATGCAGTGGCAGTCACGTGTCTGGGTGTGAGAT--
                     **..  *..  **.:*** . :      ** *********** ****..:
                                           NR3C (glucorticoid receptor)
hEdem2       CCTCAGAGCCTCCACCACACCCTGAAGGGAACACAGTGTACAGTTGATGCTACTGAGCCT
chimp        CCTCAGAGCCTCCACCACACCCTGAAGGGAACACAGTGTACAGTTGATGCTACTGAGCCT
rhesus       CCTCAGAGCTTCCACCACACCCTGAAGGGAACACAGTGTACAGTTGATGCTACTGAGCCT
mouse        -CCCTAGACTTCTACCACACCCAGAAGGGAACACAGTGTCCAGCTGACACGGCCGAGCTC
rat          -CCCTAAACTTCTGCCACACCCA--CGGGAACACAGTGTCCAGCTGATGCTGCCCAGCTG
              * *:...* ** .********:  .*************.*** *** .* .*  ***
                                        FOXO
hEdem2       GGTGTTACTG-CCCCACTAGACAAAAACATG-GAGGCTTGACCTGCAGCCCTGGACACAG
chimp        GGTGTTACTG-CCCCACTAGACAAAAACATG-GAGGCTTGGCCTGCAGCCCTGGACACAG
rhesus       GGTGTTACTG-CCTCACTAGACAAAAACACG-GAGGCTTGGCCTGCAGCCCTGGACACAG
mouse        AGCGTTGATGGCCCCACTGGATAAAAAACAGTGAGGCTTGGCTTGAAGTCCTGAACATGG
rat          GGCGCTGATGGCCCCACTGGATAAAAACAAT-GAGGCTTGGCTTGAAGTCCTGAACATGG
             .* * *..** ** ****.** *****..   ********.* **.** ****.*** .*
```

***Derl1:***

```
                     Xbp1/Creb3l2         Dual ETS motif
hDerl1       ACTTCTGGTCACGTCGTCCGCGGTCGCCGGAAGGGGAAGTTTCGCCTCAGAAGGCTGCCT
chimp        ACTTCTGGTCACGTCGTCCGCGGTCGCCGGAAGGGGAAGTTTCGCCTCAGAAGGCTGCCT
rhesus       ACTTCTGGTCACGTCGTCCGCGGCCGCCGGAAGGGGAAGTTTCGCCTAAGAAGGCTGTCT
mouse        ACTTCTGGTCACGTGGACCGCGGCCACCGGAAGGGGAAGTTTCTCTTGTGATCGCTGCCT
ratDerl1     ACTTCTGGTCACGTGGCCCGTGGCCACCGGAAGGGGAAGTTTCTCTTGTGATCGCTACCT
             ************** * *** ** *.***************** * :**: ***. **
                    Pax-binding     Xbp1/Creb3l2
hDerl1       CGCTGGTCCGAATTCGGTGGCGCCACGTCCGCCCGTCTCCGCCTTCTGCATCGCGGCTTC
chimp        CGCTGGTCCGAATTCGGTGGCGCCACGTCCGCCCGTCTCCGCCTTCTGCATCGCGGCTTC
rhesus       CGCTGGTCCGAATTCGGTGGCGCCACGTCCTCCCGTCTCCGCCTTCTGCATCGCGGCTTC
```

```
mouse           AGCTTGTCCGAATTCGGTGGCGCCACGTCCGTCGGTCTCCGCCTCCTGCACAGCGGCTGC
ratDerl1        AGCTCGTCCGAATTCGGTGGCGCCACGTCCGTCGGTCTCCGCCTCCT-----------GC
                .*** **************************   * ********** **          *
```

***Os9:***

```
                                                                  Xbp1/Creb312
hOs9            AGCGATTGATATTTCTAATCGTCGTGATCCACCTCCCCAGGA-CCTTGGAGCCACGTTTA
chimp           AGCGATTGATATTTCTAATCGTCGTGATCCACCTCCCCGGGG-CCTTGGAGCCACGTTTA
rhesus          CGCAATTGATTTTTCTAATCGTCGTGATCCACCTCCCCGGGG-CCTTGAAGCTACGTTTA
mouse           AGCTATTGATTTTTACTAACAATGCGACCCGCCTCCCTGGAGGCTTTGACGCCACGTTTA
rat             AGCTATTGATTTTTACTACCAGTGCGACCCGCCTCCCTGGAGGCTTTGACGCCACGTTTA
                .** ******:***. :* *.  * ** **.****** .*.. * ***..** *******
                                                              ERSE II
hOs9            CAAATAGGAATAGGGTACGTGGGAGGGATAGAACGTACAGCCAATAAAATCATGTGGCGC
chimp           CAAATAGGAATAGGGTACGTGGGAGGGATAGAACGTACAGCCAATAAAATCATGTGGCGC
rhesus          CAAATAGGAATAGGGTACGTGGGAGGGATAGAACATATAGCCAATAAAATCATGTGGCGC
mouse           CAAATGAGGATGCGTTACGTAAGAGAGAGCTCCCGGACAGCCAATGAAAGCTTGGGGCGC
rat             CAAATGAGGATGAGTTACGTAAGAGAGAGCGCCCGGGCAGCCAATGAGAGCTTGAGGCGC
                *****..*.**. * *****..***.** . ..*. . *******.*.* *:** *****
                                                    ETS
hOs9            CGATGGGCGTGTTGAGGCCGCTGCCTGGCTTAGGGCGGAAACAGATTCTCTGCATAAGAA
chimp           CGATGGGCGTGTTGAGGCCGCTGCCTGGCTTAGGGCGGAAACAGATTCTCTGCATAAGAA
rhesus          CGATGGGCGTGTTGAGGCCGCTGCCAGGCTTAGGGCGGAAACAGATTCTCTGGATAAGCG
mouse           TAATGGGCG-GGTGAGGCCGCTGCCAGGTTTGGGGCGGAAGGGACATCTCTAGCAGAGAC
rat             TAATAGACG-GGTGAGGCCGCTGCCAGGTTTGGGGCGGAAGGAACATCTCTAGCAGAGAC
                 .**.*.** * *************:** **.********. ...:*****. .:.**.
```

***Erlec1:***

```
                                 Xbp1/Creb312
hErlec1         CGGGCTCTCCGGAAGGAGACGTGGCGGCGG---
Chimp           CGGGCTCTCCGGAAGGAGACGTGGCGGCGG---
Rhesus          CGGGCTCTCCGGAAGGAGACGTGGCGGCGG---
Mouse           CCAGCCGTCTGCAAGGAGACGTGGCGGCGGCGG
Rat             CCAGCCGTCCGGAAGGAGACGTGGCGGCGG---
                *  .**   ** * ******************
```

***Vcp:***

```
                                          Xbp1/Creb312 NF-YA/B
hVcp            CCCGCCTCAGTGATGCGACACGCCTAGTAACGCCACGTCAATTGGCCGCAACGC
Mouse           CCCGCCTCAAGGCTGCAACACGCCTAGTAACGCCACGTCAATTGGCCGCAGCAC
Chimp           CCCGCCTCAGTGATGCGACACGCCTAGTAACGCCACGTCAATTGGCCGCAACGC
Rhesus          CCCGCCTCAGTGATGCGACACGCCTAGTAACGCCACGTCAATTGGCTGCAACGC
Rat             CCCGCCTCAAGGCTGCAACACGCCTAGTAACGCCACGTCAATTGGCCGTAGCAC
                *********. *.***.***************************** * *.*.*
```

***Syvn1:***

```
hSyvn1          GGAAAACAAAAATTTGTCTTCTCCATG--------ATCACCCCCACAAGTTGTGTGGGAG
chimp           GGAAAACAAAAATTTGTCTTCTCCATG--------ATCACCCCCACAAGTTGTGTGGGAG
rhesus          GGAAAACAAAAATTTGTCTTCTCCATG--------ATCACCCCCCAAG-TTGTGTGGGAG
mouse           AGAAAACAGATATAGGTCTTCCCAGGTTACCT---CCCCCCAACGCAGGTTGTGTAGGAG
rat             GGAAAACAGATATATATCTTCCCAGATCACCTCCCCCCCCCAACGCAGGTTGTGTAGGTG
                .*******.*:**: .***** *..          . *.**..* .*. ******.**:*
                                   ERSE I                        ERSE II-like
hSyvn1          TGAGTTGGGTCCCGTGGCTGCCACTGATTGGTTGCGATAAG-TTACGCGGCCCAATAAGA
chimp           TGAGTTGGGTCCCGTGGCTGCCACTGATTGGTTGCGATAAG-TTACGCGGCCCAATAAGA
rhesus          TGAGTTGGGTCCCGTGGCTGCCACTGATTGGTTGCGATAAG-TTACGCGGACCAATAAGA
mouse           TGAGTTTGTTCCCGTGGCTACCGCCGATTGGTTGCGATAAAGTTACGCGGACCAATGAGA
rat             TGAGTTTGTTCCCGTGGCTACCGCTGATTGGTTGCGGTAAAGTTACGCGGACCAATGAGA
                ****** * **********.**.* ***********.***. ********.*****.***
                                                           E-box
hSyvn1          AAAAGAATGTACCTTTGCTGGCCTATAGAAAGGGGAAAGCAGGTGAGTGGTTGTTAGGGG
chimp           AAAAGAAAGTACCTTTGCTGTCCTATAGAAAGGGGAAAGCAGGTGAGTGGTTGTTAGGGG
rhesus          AAAAGAAAGTACCTTTACTGGCCTATAGAAAGGGGAAAGCAGGTGAGTGGTTGTTAGGGG
mouse           AAAGAAAGGAAAAATTTCTGGCCTATAGGGAAAGAAAAGCAGGTGAGTGGTTGCTAGGGT
rat             AAAGAAAAGAAAAATTTCTAGCCTATAGAGAAAGAAGAGCAGGTGAGTGGTTGCTAGGGT
                ***..** *:*..:** **. *******..*..*.*.**************** *****
```

*UPR pathways*

**Slc35b1:** an ATP/ADP antiporter, importing ATP into the ER lumen for energy/protein folding.

```
hSlc35b1        CG-ACCACACCTCCTCCCGCCCCGCCTACCTGTCCGTCTTCGCACAGCGCAGCCAATTGG
rhesus          CG-GCCGCACCTCCTCCTGTCCCGCCTACCCGTCCGAGTTCGCACAGCGCAGCCAATTGG
chimp           CG-GCCACACCTCCTCCTGCCCCGCCTACCTGTCCGTCTTCGCACAGCGCAGCCAATTGG
mouse           TG-GCCTCACCGCCTCTTTCTCCGCCCATCTGTCTATTTTCGCTCTGCGGAGCCAATTGA
rat             TAGACCTCACCGCCTCTTTCTCCGCCCAGCTGTCTATTTTCACTCCTCGTAGCCAATTGA
                  .  .** **** ****     ***** *  * ***  .: ***.*:*   ** *********.
              ERSE-I  (NFYA/XBP1)  Creb312
hSlc35b1        TGGCCTCCACATCCTTATACGTCATTATTCTACGCCCAGGTGGC-TCTAGGACACCCTGT
rhesus          TGGCCTCCACATCCCTATACGTCATTATTCTACGCCCGGGTGGC-TCTAGGACACCCTGT
chimp           TGGCCTCCACATCCTTATACGTCATTATTCTACGCCCAGGTGGC-TCTAGGACACCCTGT
mouse           TAACATCCACATACTTATACGTCATCACTATACGCCGCGTAATCGCCTAGGCCTCCCCAC
rat             TAACATCCACATACTTGTACGTCATCACTATACGCCGCGTAAACTCCTAGGCCTCCCCGC
                *..*.*******.* *.******** * *.******   *  :.  *   *****.*:***  .

hSlc35b1        GGTGCTCTGGGTCGCAGTTGTCCTCAGAGTTGCCTGATTGGAAAAATCTCCAGGGCGAGG
rhesus          GGTGCTCTGGATCGCAGTTGTCCTCAGAGTTGCCTGATTGGAAAAATCTCCAGGTCGAGA
chimp           GGTGCTCTGGGTCGCAGTTGTCCTCAGAGTTGCCTGATTGGAAAAATCTCCAGGGCGAGG
mouse           CGAGCTCCAGCTTGTAGATGTCCTCGGCGTTGCCTGATCGGAAAAACCTAAAAGAGGAGA
rat             CGAGCTCCGGGTTGTAGGTGTCCTAGGCGTTGCCTGATCGGAAAAATCTAAAAGATGAGA
                 *:****  .*  *  *  **  ******..*.********** ******* **..*.*   ***.
```

---

**Ern1:**

```
                           E-box for Mist1            NF-KB      ETS
hErn1          GCCCTAGGGTTCTCCAGCTGTGCCTCGTCTTCTCTGGGG-CCCCTTTCTTCCTTTTCCTT
chimp          GCCCTAGGGTTCTCCAGCTGTGCCTCGTCTTCTCCGGGG-CCCCTTTCTTCCTTTTCCTT
rhesus         GCCCTAGGGTTCTCCAGCTGTGCCTCGTCTCCTCCAGGG-CCTCTTTCTTCCTTTTCCTT
mouse          -GACGAGGGTCCTCCAGCTGTGCCTCATCCAGCCCGAAGGCCCCTTTCTTCCCTAGCCTT
rat            -GAGGAGGATCCTCCAGCTGTGCCTCATCCAGCCCGAAGGCCCCTTTCTTCCCTAGCCTT
                 .  ***.* ***************.**     *  ...* ** ********* *: ****
                          NF-kB                            ETS
hErn1          G-CCCTCAGGAATTCCCCTCAAGTCATATCCACCAGCCATCACTTCCGCTTGGAATGAAC
chimp          G-CCCTCAGGAATTCCCCTCAAGTCATATCCACCAGCCATCACTTCCGCTTGGAATGAAC
rhesus         G-CCCTCAGGAATTCCCCTCAAGTCACATCCAACAGCCATCAGTTCCGTTTGGAATGAAC
mouse          GCCCCTCAGGAATTCCCCGAAAGTCATGTCCAGCAGCCATCCGTTCCTCTCGGGATGAAC
rat            GCCCTCAGGAATTCCCCCCCAAGTCATGTCCAGCAGCCTTCCGTTCCTCTCGGAATGAAC
               * **   .*.*:* **** .****** .**** *****:**. ****  * **.******
                             MEF                                Creb3L2
hErn1          CACGGCCAAGTCACTCACCAAAAATTATAAGGAGGAAACCATTTCGGCCACTTGTGTCAC
chimp          CACGGCCAAGTCACTCACCAAAAATTATAAGGAGGAAACCATTTCGGCCACTTGTGTCAC
rhesus         CACGGCCAAGTCACTCACCAAAAATTATAAGGAGGAAACCAATTCGGCCACTTGTGTCAC
mouse          AACAGCCAACTCATTCACCAAAAATTATAAG---GAAATCATGCGGGCCTCTCGTGTCAC
rat            CACAGCCAACTCATTCACCAAAAATTATAAG---GAAATCATGCAGGCTTCTTGTGTCAC
               .**.***** *** *****************    **** **:    *** :** *******
                    Creb312
hErn1          GTGCGTTGATGTGACACGTGTGGATGTG--------AAGTTTGAGATGGTCTGCGTCACA
chimp          GTGCGTTGATGTGACACGTGTGGATGTG--------AAGTTTGAGATGGTCTGCGTCACA
rhesus         GTGCGTTGATGTGACACGTGTGGATGTG--------AAGTTTGAGATGGCCTGCGTCACA
mouse          ATGTGTTTATGTGACCTTTGTAGATGTGGCCTGGTGAAGCTTGAGAGGACATGTGTCACA
rat            ACGTGTTTACGTGACCTTCGTAGATGTGGCCGGGTGAAGCTTGAGAGGACATGTGTCACA
               . * *** * *****.   **.******        *** ****** *.  .** ******
```

---

**Ern2:**

```
                                   Creb312
hErn2          GTGTCATCTGGGTCTCCCCTGACTCAGGGTCCACCACAAACCAGG
Mouse          GTGTCATCTGAGTCTCCCCTGACTCAGGGTCCACTACAAACCAGG
Chimp          GTGTCATCTGGGTCTCCCCTGACTCAGGGTCCACCACAAACCAGG
Rhesus         GTGTCATCTGGGTCTCCCCTGACTCAGGGTCCACCACAAACCAGG
Rat            GTGTCATCTGAGTCTCCCCTGACTCAGGGTCCACCACAAACCAGG
               **********.*********************** **********
```

---

**Xbp1:**

```
hXbp1          ------GCGCGCCGCAGCCGCCCAGCGCCCAGCCTCGCCGCGCCCGGCCTTTCTACGGTC
Chimp          ------GCGCGCCGCAGCCGCCCAGCGCCCAGCCTCGCCGCGCCCGGCCTTTCTACGGTC
```

```
baboon          CGCGCGCCGCGCCGCCGCCGCCCAGCGGCCAGCCCAGACGCTCCCGGCCTTTCTACGGTC
mouse           ----------GCCGCCACCGCCCCGTGGCCT--CCTGCCGCGCCCAGCCTTTCTAGGGTC
rat             ----------GCCGCCACCGCCCCGTGGCCG--CCTGCCGCGCCCAGCCTTTCTAGGGTC
                          *****..******.* * **   *  *.*** ***.********* ****
                     ERSE I                         ERSE II-like
hXbp1           GTGGCCCTCCGCGATTGGCCAGCGTCGCGTGACGCACGGCCGAGCTCGGCGTCCATTGGT
Chimp           GTGGCCCTCCGCGATTGGCCAGCGTCGCGTGACGCACGGCCGAGCTCGGCGTCCATTGGT
baboon          GTGGCCCTCCGCGATTAGCCAGCGTCGCGTGACGCACGGCCGAGCTCGGCGTCCATTGGC
mouse           GTGGCCGTCCGCGATTGGCCGGTGCTCCGTGACGCGCGGGTATATTCATCACTTATTGGT
rat             GTGGCCGTCTGCGATTGGCCTGTGCTGCGTGACGCGCGGCTGTATTCATCGTTTATTGGT
                ****** ** ******.*** * *   ********.***  .:. **. *.   *****

hXbp1           CCGGCCTGCCCGGCG--GGGCGTTTCAGGACCGTGGCTATGGAGTCCGGCGTGGCAGC
Chimp           CCGGCCTGCCCGGCG--GGGCGTTTCAGGACCGTGGCTATGGAGTCCGGCGTGGCAGC
baboon          CCGGCCTGCCCGGCG--GGGCGTTTCGGGGGCGTGACTATGGAGTCCGGAGTGGCAGC
mouse           CCTGTAGTCCCGGGGGCGGGCTCAACTCGGGCGGGGTGAG--TGTCCAGCGTGGCTGC
rat             CCTGCAGGCCCGGGGGCGGGCTCAACGCGGGCGGGGTGAG--TGTCCAGCGTGGCTGC
                ** * .  ***** *  ****  ::*  *. ** *.  *   :****.*.*****:**

---
hXbp1intron      TCTGAGCTGCTACTCTGTTTTTCAGTTTCCTAGGAAGAGAAGGA----ACATACCACACT
chimp            TCTGAGCTGCTACTCTGTTTTTCAGTTTCCTAGGAAGAGAAGGA----ACATACCACACT
baboon           TCTGAGCTGCTACTCTGTTTTTCAGTTTCCTG---AGAAAAGGA----ACATACCACACT
mouse            TCTGCGCTGCTACTCTGTTTTTCAGTTTCCTTGGGG----AGGAGACAGGGAACATAACT
rat              TCTGTGCTGCTACTCTGTTTTTCAGTTTCCTTGAGGGAGGGGGAGACAAAGAACATATCT
                 **** **************************    .    .***    . .:**.:.:**
                 Creb3l2                                      E-box
hXbp1intron      GAGTCATATCAAACCTTATTTATGTCTTTATTTGAAAACTTTACCAGCTGGTGCTTTCAT
chimp            GAGTCATATCAAACCTTATTTATGTCTTTATTTGAAAACTTTACCAGCTGGTGCTTTCAT
baboon           GAGTCATTTCAAACCTTATTTATGTCTTTATTTGAAAACTTGACCAGCTGGTGCTTTCAT
mouse            GAGTCATCTCCAGCTTTCATGTGTCTT-CATTTGTAGACGTGGCCAGCTGGTGCTTTCGT
rat              GAGTCATCTCCAGCTTTCCTGTGCCTTCCATTTGTAAACGTGGCCAGCTGGTGCTTTCGT
                 ******* **.*.* **. * :    *  *****:*.** * .***************.*
```

***Creb3l2:***

```
hCreb3L2            AGCTTCTGCTTCCACTCACACCTAAAAGAGTAAAACAGCCTGAAAATAAATCTTAAGCCT
chimpCreb3L2        AGCTTCTGCTTCCACTCACACCTAAAAGAGTAAAACAGCCTGAAAATAAATCTTAAGCCT
rheususCREB3L2      AGCTTCTGCTTCCCCTCACACCTAAAGGAGTAAAACAGCCTGAAAATAAATCTTAAGCCT
MouseCREB3L2        GGCCTCAGTGTCCCCTTGCACCTAAGG-GAATACAGAGCCTGAAAACAAATCTTAAGCCT
ratCreb3L2          GCCCTGCGCTTCCCCTTACACCCGAGG-AAATACACATCCTAAAAATAAATCTTTGGTCT
                    . * *  *  ***.** .**** .*.. ..::*.* * ***.**** *******:.* **
                    Creb3l2
hCreb3L2            GAGTCAATGTTCCCTGCAACTCCCCTTCTCTGGCCTTTTCTGAGATTCTATTGG-CAAAG
chimpCreb3L2        GAATCAATGTTCCCTGCAACTCCCCCTCTCTGGCCTTTTCTGAGATTCTATTGG-CAAAG
rheususCREB3L2      GAGTCAATGTTCCCTGCAACTCCCCCTCTCTGGCCTTTTCTGAGACTCCTTTTGGCAAAG
MouseCREB3L2        GAGTCAGTCTTCCCTGCAAATCCCTCTCCTCGGTCTTTTCTGGGATTCTT---GGCAAAG
ratCreb3L2          GAGTCAGTCCTCCCTGAGAACCCCTCTCCTTGGCCTTTTCTGGGACTCCT---GGCAAAG
                    **.***.*  ******..*. ***  **   ** ********.** ** :   * *****
                                        CREB3L2  E-box(Mist1)
hCreb3L2            AATGGCTTCCCTTGAGGATGGCTGAGCTAACAGCTGGGCAGTACTGGGTGATCTCTGCCC
chimpCreb3L2        AATGGCTTCCCTTGAGGATGGCTGAGCTAACAGCTGGGCAGTACTGGGTGATTTCTGCCC
rheususCREB3L2      AATGGGTTCCCTTGAGGATCGTTGAGCTAACAGCTGGGCAGTACTGGATGATTTCCGCCC
MouseCREB3L2        AATTTGCTTCCCTGAGGACAGCTGAGCTAACAGCTGGACAGTAGTGGCTGATTTCCATAA
ratCreb3L2          AATTTGCTTCCCTTCGGACAGCTGAGCTAACAGCTGGACAGTACTGACTGATTTCCACAA
                    ***    * ** * .***  * ***************.***** **. **** ** . ..
                           E-box for Mist1                   GATA (GATA2)
hCreb3L2            TTTCAACAGCTGGTCTCTGAGACAAACAACCAGACAGGC-TTTATCAGAAAAAA------
chimpCreb3L2        TTTCAACAGCTGGTCTCTGAGACAAACAACCAGACAGGC-TTTATCAGAAAAAAA-----
rheususCREB3L2      TTTCAACAGCTGGTCTCTGAGACAAACAACCAGACAGGC-TTTATCAGAAAAAA------
MouseCREB3L2        TTCCCACAGCTGGCCTTAGAGACAAACAGCCGGAAAGGCGTTTATCAACAAAACTAAACC
ratCreb3L2          TTCCCACAGCTGGCCTTAGAGACAAACAGCGGGAAAGGCGTTTATCAACAAAACAAAACC
                    ** *.******** ** :**********.* .**.**** *******  ****
                     Creb3l2
hCreb3L2            -TGACTCAACTTGTGGAGGCCAAGCATAGATGGAGAGCAG-TATCAGACTTAAAACTACT
chimpCreb3L2        -TGACTCAACTTGTGGAGGCCAAGCATAGATGGAGAGCAGTTATCAGACTTAAAACTACT
rheususCREB3L2      -TGACTCAACTTGTGGAGACCAAGCATAGATGGAGAGCAG-TATCAGACTTAAAACTACA
MouseCREB3L2        ATGACTCAACTTTTACAGGTCAAGTGTGGATGGAGA------------CTTGAAAGCCCC
ratCreb3L2          GTGACTCAACTTTTACAGGCCAAGTGTGGACAGAGA------------CCTAAAAGCCCT
                     *********** *. **. **** .*.** .****            * *.***  .*
```

#### ***Creb3l1:***

```
                           CEBP                E-box for Mist1
hCreb3l1       GGGGCTTGGCAAACCCTTATCGCCTGAGAAGCAGCTGCTGGAGAAATACCTTAGCGGCTG
Chimp          GGGGCTTGGCAAACCCTTATCGCCTGAGAAGCAGCTGCTGGAGAAATACCTTAGCGGCTG
Rhesus         GGGGCTTGGCAAACCCTTATCGCCTGAGAAGCAGCTGCTGGAGAAATACCTTAGCGGCTG
Mouse          GGGGGTTGGCAAACTCTTATCGCCTGCGGAGCAGCTGCTGGAGAAATACCGTGGCCGCTG
Rat            GGGGCTTGGCAAATCCTTATCGCCTGCGGAGCAGCTGCTGGAGAAATACCGTGGCCGCTG
               **** ********  ***********.*.******************** *.** ****
               Creb3L2
hCreb3l1       ACTCACCGGCTCCCTCTGCTCA---CAGCCCCCTC
Chimp          ACTCACCGGCTCCCTCTGCTCA---CAGCCCCCTC
Rhesus         ACTCACCGGCTCCCTCTGTTCA---CAGCCCCCTC
Mouse          ACTCATGG-CTCTTCCAGCTCTGTGCAACCCCCTC
Rat            ACTCACGG-CTCCTTCAGCTCTGCGTAACCCCCTC
               *****   * ***    *:* **:     *.*******
```

```
                                                             ETS for Spdef
hCreb3l1       CCATGCAGCCCAATTTAGGGAGCTGCTAGAGAACAGAGGGCCCCGTCTGCACCCCGGAAG
rhesus         CCATGCGGCCCATTTTAGGGAGCTGCAAGAGAACAAAGGGCCCCGTCTGCACCCCGGAAG
rat            CCCTGCAACCCATTCTAGGGAACCACAGAGAACAAAGGACCCCCATCTGCACCCCGGAAG
mouse          CCCTGCAACCCATTCTAGAGAAGCACAGAGAACAAAGGGCCCCCACCTGCACCCCGGAAG
               **.***..****:* ***.**.   .*:....*..*..*. ****. **************
                       ETS for Spdef
hCreb3l1       TCAGTCTAATCTCCAGGATGAATATGTTTTCATTTCTCAGAAGTTCCTGTTAGAGTGTGC
rhesus         TCAGTTTAATCTCCAGGATGAATATGTTTTCATTTCTCAGAAGTTCCTGTTAGAGTGTGC
rat            TCAGTCTGATCTCCAGGATGAATATGTTTTTATTCCCCACAAGTTCCTGCTCAAGTGTGC
mouse          TCAGTCTGATCTCCAGGATGAATGTGTTTTCATTCCCCACAAGTTCCTGCTCAAGTGTGC
               ***** *.***************.****** *** * ** ********* *..*******
                       IRF
hCreb3l1       CAAAGCATTTTTCCCTTTACAACCCAGGCAAACAAGACAAGTTTACTTTGGGTGACCTTA
rhesus         CAAAGCATTTTTCCCTTTACAACCCAGGCAAACAAGACAGGTTTACTTTGGGTGACCTTA
rat            CAAAACG-TTTTTCCTTTACGATGCAAGCAAACAAGGCAGGTTTACTTTGGGTGACCTTC
mouse          CAAAACGTTTTTTCCTTTACGAGGCAGGCAAACAAGGCAGGTTTACTTTGGGTGACCTTC
               ****.*. **** *******.*   **.*********.**.********************.
```

#### ***Mbtps1:***

```
                  TCFL5/TFE3          FOXO          TEAD          Xbp1/AHR/HIF
hMBTPS1          CCAGTCTCGCGAGAGTTGGGAGTAAACAGCCCC-GAATGGAGTGCCCAGGCGTGTTCGCC
gorilaMBTPS1     CCAGTCTCGCGAGAGTTGGGAGTAAACAGCCCC-GAATGGAGTGCCCAGGCGTGTTCGCC
chimpMBTPS1      CCAGTCTCGCGAGAGTTGGGAGTAAACAGCCCC-GAATGGAGTGCCCAGGCGTGTTCGCC
ratMBTPS1        GTGGTCTCGCGAGATTTGCGAGTAAACATCCCCCGAATGGATACCCGAGGCGTGTTCGCG
rhesusMBTPS1     CCAGTCTCGCGAGAGTTGGGAGTAAACAGCCCC-GAATGGAGTGCCCAGGCGTGTTCGCT
mouseMBTPS1      GTGGTCTCGCGAGACTTGCGAGTAAACATCCCCCGAATGGAGACCCGAGGCGTGTTCGCG
                    .*********** *** ********* **** ******* : ** ************

hMBTPS1          GCGGAGGCGCCGTTATCCCGGGCC
gorilaMBTPS1     GCGGAGGCGCCGTTATCCCGGGCC
chimpMBTPS1      GCGGAGGCGCCGTTATCCCGGGCC
ratMBTPS1        GCGGAGCCGCCGTTTTCCCGGGTC
rhesusMBTPS1     GCGGAGGCGCCGTTATCCCGGGCC
mouseMBTPS1      GCGGAGCCGCCGTTCTCCCGGGTC
                 ****** ******* ******* *
```

#### ***Mbtps2:***

```
                     ETS
hMBTPS2          TTCTCTTCCCCTGCCTGG-CGCCTC-CCGGGCTGG------------------ACGGTGC
Gorilla          TTCTCTTCCCCTGCCTGG-CGCCTC-CCGGGCTGG------------------ACGGTGC
rhesus           TTCTCTTCCCCTGCCTCG-CGCCTC-CCGGGCTGG------------------ACGGTGC
mouse            TTCTCTTCCCCTTCGGAGTCTCCGCGCCGGGCCGGGAGCCGCTCCGAGCGCCCGCCGCGT
rat              TTCTCTTCCCCGGCCGAGTTAACTCGGCGGGCTGCAACCCGCTCCGT----CCTTGGTGT
                 ***********  *   *    .* *  ***** *                      * *
                    XBP1/Creb3l2   AHR site
hMBTPS2          CCACGTGCGACAGTCCGCGTGGTGGATCGGCCCGGTGGAGTAAATAAAACATT----GCC
Gorilla          CCACGTGCGACAGTCCGCGTGGTGGATCGGCCCGGTGGAGTAAATAAAACATT----GCC
rhesus           CCACGTGCGACAGTCCGCGTGGTGGATCGGCCCGGTGGAGTAAATAAAGCATT----GCC
mouse            CTGCGTGCGCTCTCCCGCGTGGCGCATCCGTCCGACCGAGTAAAGAAAGTAGGGCAGGCA
```

```
rat          CTACGTGCACTCTACCGCGTGGCGCATCCGTCCGGCGGAGTAAATAAAGTAGG----GCA
             * .*****.. .  ******** * *** * ***.  ******* ***. *      **.
                  KLF
hMBTPS2      GGCCGCGGGAGGGTTGTGGCGGCCTGCGCGGCTGCCACTGT---------------
Gorilla      GGCCGCGGGAGGGGTGTGGCGGCCTGCGCGGCTGACACTGTTGCAGTGCGGGGCAG
rhesus       GGCCGCGGGAGGGGTGTGACGGCCTGCGCGGCTGCCACCGT---------------
mouse        GGCTGCGGGAGGGGAGGGGTG----TGGCGGCCACCGCATTGCGGGCGCCGCC---
rat          GGCTGCGGGAGGGGTGTGGCGGCCCGTGCCACCGCCGCATTGCGGGCACC------
             *** ********* :* *. *      ** .* ..*.*  *
```

---

```
                                Creb3l2
hMbtps2      -----------CCCATTTCCATGAGTCATGGATGTTGTTTTTAATCATGTTCTTATTAAT
rhesus       -----------CCCATTTCCATGAGTCATGGATGTTGTTTTTAATCGTGTTCTTATTAAT
chimp        -----------CCCATTTCCATGAGTCATGGATGTTGTTTTTAATCATGTTCTTATTAAT
mouse        ATACTTGTAGTTTCTACTCCATGAGTTATGAGTGTTGCTTTTAATAATGTTCTTATTCAT
rat          ATACTTGTAGTTTCTGTCCCATGAGATATGAGTGTTGTTTTTAATAATGTTCTTATTTAG
                         *:    *******: ***..***** *******..********** *

hMbtps2      GGAGTAGACTAAGGATTAGTAACAGATTGG-----GTTTTTTTTTCTTGTCTCTTTGTGT
rhesus       GGAGTAGACTAAGGATTAGTAACAGATTGGGTTTTTTTTTTTTTTCTTGTCTCTTTGTGG
chimp        GGAGTAGACTAAGGATTAGTAACAGATTGGG----TTTTTTTTTTCTTGTCTCTTTGTGT
mouse        AGGACATTTAAGG--TTAGGAACAGATTATT----TTAAAAAGAAGGAAATCTGATCTC
rat          AGGACATTCAAATGGTTAGGAACAGATTTTT----TTCTTTTTTTTTTAATCTTTATTA
             .*.. * : :*.   **** ********        * ::::: :    :.*** :
```

---

#### *Golgi-resident Proteins*

#### ***Golga2:***

```
                                                   Creb3l2
hGolga2intron2   AAGATACGAGTTACCCAAGGCTACACCATGAGTCAGTGGCACAGCTGGCACTAGAGCTTT
chimp            AAGACACGAGTTACCCAAGGCTACACCATGAGTCAGTGGCACAGCTGGCACTAGAGCTTT
rhesus           AAGACATGAGTTACCCAAGGCTACACCATGAGTCAGTGGCGCAGCTGGCACTAGAGCTTT
mouse            -----CTGAGTTACCTGGGGCCACAGCATGAGTCAGGAGCATTGCTGGGACCAGAGCACA
rat              -----CTGAGTTACCTGGGACCACACCATGAGTCAGGAGCATGGCTGGGACCAGAGCTCA
                      . ******** ..*.* *** ********** .**.  ***** ** *****: :

hGolga2intron2   GCTGGGCACACATGCACAC
chimp            GCTGGGCACACATGCACAC
rhesus           GCTGGGCACACATGCAAAC
mouse            GCTCTACACAGACCCAAAC
rat              GCTCTACACAAACCCAAGC
                 ***  .**** *  **..*
```

**---**

```
                                      Creb3l2     TEAD
hGolga2Intronic   CCG----AGGGGGATTGTGACGTCACTACATTCC
Chimp             CCG----AGGGGGATTGTGACGTCACTACATTCC
Rhesus            CCG----AGGGGGC-TGTGACGTCACTACATTCC
Mouse             TCG---AGGGGGGACTGTGACGTAACTACATTCC
Rat               TCGGGGGGGGGGGACTGTGACGTAACTACATTCC
                   **    .*****. ********.**********
```

---

#### ***Golgb1:***

```
                         ERSE II     Xbp1
hGolgb1promoter   CCCCCCTCCCAATTCAAGCCACGTCAGCTCGGGAAGCCCGTACGCGACACCTTCCACCGC
chimp             CCCCCCTCCCAATTCAAGCCACGTCAGCTCGGGAAGCCCGTACGCGACACCTTCCACCGC
rhesus            CCCCCCTCCCAATTCAAGCCACGTCAGCTCGGGAACCCCGTACGCAACACCTTCCACCGC
rat               CCCCCCGCCCAATCTAAGCCACGTCAGCTCGGGTAACCCAAGCGCAGCGCCA-CCACCGA
mouse             CCCCCCGCCCAATCGAAGCCACGTCAGCTCGG-TAACCCAAGCACGGCGCCA-CCGTCGA
                  ****** ******  ***************** :* ***.:.*.*..*.**: **. **.
                                                         ETS
hGolgb1promoter   CGCGCCCCGCCAGGGACTCAGCCACTAGCAGTTGGGACGGAAGATGGGATTAAGACACCG
chimp             CGCGCCCCGCCAGGGACTCAGCCACTAGCAGTTGGGACGGAAGATGGGATTAAGACACCG
rhesus            CGCGCCCCACCAGGGACTCAGCCACTAGCAGTTGGGACGGAAGATGGGATTAAGACACCG
rat               CGCGCCCC---TGGGAGTCAGCCACTCGCTG-TGAAACGGAAGGCTGGATTAAGGCGTCG
mouse             CGCGCCCCT---GGGAGTCAGCCACTCGCCG-TGAGACGGAAGGCTGAATCAAGGCGTCG
```

```
                     ********    **** *********.** * **..*******.  *.** ***.*. **

hGolgb1promoter     ACCAATCAGGACTATCGGCAGGACGTTACCAGGGAGACCAGGCACACCTTGTTTGTTACC
chimp               ACCAATCAGGACTATCGGCAGGACGTTACCAGGGAGACCAGGCACACCTTGTTTGTTACC
rhesus              ACCAATCAGGACTATCGGCAGTGGGTTACCAGGGAGACCAGGCACGCCTTGTTTGTTACC
rat                 ACCTATCGGAGCTACGCACAGGGCGTTACCATGGAGATCAGGCACGCCTTGTTTGTAACT
mouse               ACCTATCGGAGCGGTGCTCAGGGCGTTGCTATGGGAACCAGGCACGCCTTGTTTGTAACT
                    ***:***.*..*  .     ***  . ***.* * **..* *******.**********:**
                                                                 ETS
hGolgb1promoter     AACCTATCAACAATGACGGAAACCAATCAACGACAGAAGATGCTCCGCTTCCTCAAAGAA
chimp               AACCTATCAACAATGACGGAAACCAATCAACGACAGAAGATGCTCCGCTTCCTCGAAGAA
rhesus              AACCTATCAACAATGACGGAAACCAATCAACGACAGAAGATGCCCCGCTTCCTCAAAGAA
rat                 AACTGCGTCACAATGACAGACACCAACCAATGCCCTAAAGCCTTCCACTTCCTCAAAGAC
mouse               AACTGCGTTACACTGACAGAAACCAACCAATGCCCTAAAGCCTTCCACTTCCTCAAAGAC
                    ***   .   ***.****.**.*****  *** *.*. **..    **.*******.****.
                              ERSE II-like
hGolgb1promoter     ACAGCCAATGGACCCTAAAGAGTGAATGAGGGGAGTGATTAGACGCGAGTAGGTGCCCCG
chimp               ACAGCCAATGGACCCTAAAGAGTGAATGAGGGGAGTGATTAGACGCGAGTAGGTGCCCCG
rhesus              ACAGCCAATGGACCCTAAAGAGTGAATGAGGGGCGTGATTAGACGCGAGTAGGTGCCCCG
rat                 GGAACCAATGGATTCTAAGAAGGACACGCAGGACGGAATCATATGGTGGGGTGTGACCCG
mouse               GGAGCCAATGGATTCTAAGAAGGACGTGCCGGGTGGAATCACATGGTGGGATGTGACCCG
                    . *.********  ****..**  ... *. **. *  .** * * *   .*  . ***.****
```

#### *Golph3:*

```
                          ETS
hGolph3promoter     ------CTTCCGGCGCCTCCCGGCCCAGCTCTCCCGCGCCGACTCTGCCACATCCTCCGG
chimp               ------CTTCCGGCGCCTCCCGGCCCAGCTCTCCCGCGCCGACTCTGCCACATCCTCCGG
baboon              CCGCCGCTTCCGGCGTCTCTCGGCCCAGTTCTCCCGCGCCGACTCTGCCACATCCTCCGG
mouse               ------CTTCCG--GCACCCCGGCCCCGC---CCCGCGCCGGCGCTGCCACATCCTCCGG
rat                 ------CTTCCG--GTACCCCGGCCCCGC---CCCGCGCTGGCGCTGCCACATCCTCCGG
                          ******  *  . * ******.*    ******* *.* ****************
                    Creb3l2               NFYA                KLF
hGolph3promoter     TGACGTCAGCCGGGGCCGCCATATTGGAAAGG--CG---CCGCCGCCGCCTCCGCCTTGG
chimp               TGACGTCAGCCGGGGCCGCCATATTGGAAAGG--CG---CCGCCGCCGCCTCCGCCTTGG
baboon              TGACGTCAGCCGGGGCCGCCATATTGGAAAGG--CG---CCGCCGCCGCCTCCGTTTCGG
mouse               TGACGTCAGCGCGCGCCGCCATATTGGGAAGCGCCACCGCCGCCGCCGCCTCCGCTCGGG
rat                 TGACGTCAACGCGCGCCGCCATATTGGGAAGTGCCA---CCGCCGCCGCCTCCGCTCGGG
                    ********.*  * *************.***    *.   ***************    **

hGolph3promoter     AGCTCGGGGTGTTTCGGGGACTGCGGCCACAGGCAGGAAGGCGCTCCTCTCCTGCCCCGC
chimp               AGCTCGGGGTGTTTCGGGGGCTGCGGCCACAGGCAGGAAGGCGCTCCTCTCCTGCCCCGC
baboon              AGCTCGGGCTGTTTCGGGTGCCGCGACCGCAGGCAGGAAAGCGCCCCTCTCCTGCTCCGC
mouse               CTCCTGGGCTGTGTGGGGGGGCCACAGCCGCGGGCTG-AAGGCTCCCTTGTG-TACCCCAT
rat                 CTCCTGGGCTGTGTGGGG--CCACAGCCGCGGGCTG-AAGGCTCCCTTGTG-TATCCCAT
                    . *   *** *** * ***   *  .*..**.*.***:* **.** * * * *  *.   **.

hGolph3promoter     CGACGCCCGGCCAGCCCGCTTCGCCCTGACCTGTTTCCTCATGACTGCCCCCGGCCCTGC
chimp               CGACGCCCGGCCAGCCCGCTTCGCCCTGACCTGTTTCCTCATGACTGCCCCCGGCCCCGC
baboon              CGACGCCCGGCCAGCCCGCTTCGCCCTGACCTGTCTCCTCATGACTGCCCCCGGCCCTGC
mouse               CGACGCCCGGCCCGCCGGCTTCGCCCAGCTCCGGCTCCTCATGACCGCCTTCGGCCCCGC
rat                 CGACGCCTGGCGCGCCCGCTTCGCTCAGACCCGGCTCCTCATGACTGCCTTCGGACCCGC
                    ******* ***  .*** ******* *:*.  * *  ********** ***  ***.** **
                                                  SRF      ETS
hGolph3promoter     TGCCGACGGACGTCGCCCCGGCGTCCGGATTTAACACGGAA--ACCCGGATCGGAGGCCG
chimp               TGCCGACGGACCTCGCCCCGGCGTCCGGATTTAACACGGAA--ACCCGGGTCGGAGGCCG
baboon              TGCCTACGGACGTCGCCCCGGCGTCCGGATTTAACACGGAA--ACCCGGGTCGGAGGCCG
mouse               AGCCCACGGACATCGCCCCAGCTCCAAGAATTAACACGGAAGCTCCCGGGGCCGGGGCCG
rat                 AGCCCACGGACATCGCCCCAGCTCCGAGAATTAACACGGAAGCTCCTGGGGCCGGGACCG
                    :*** ****** *******.**  *  .**:***********   :** **. * *.*.***

hGolph3promoter     CGCGGGGAGGAGGAGGGCGACCCGGTCGGTCCTGCGACCCTCTCGGCCCGGCTCGGCGCC
chimp               CGCGGGGAGGAGGAGGGCGACCCGGTCGGTCCCGCGACCCTCTCGGCCCGGCTCGGCGCC
baboon              CGCGGGGAGGAGGAGGGCGAGCCGGTCGGTCCCGCGTCCCTCTCGGCCCGGCTCGGCGCC
mouse               T-----------GAGGGAGAGCCGGCCGGTCCCGCGTCCCTGTCGGTCCGGCGCGGCTCT
rat                 T-----------GAGGGAGAGCCGGCCGGTCCCGCGTCCCTGTCGGTCTGGCGCGGCTCT
                                *****.** **** ****** ***:**** **** * *** **** *

hGolph3promoter     TCGGCGGGAGCCATGACCTCGCTGACCCAGCGCAGCTCCGGCC
chimp               TCGGGGGGAGCCATGACCTCGCTGACCCAGCGCAGCTCCGGCC
baboon              TCGGCGGGAGCCATGACCTCGCTGACCCAGCGCAGCTCGGGCC
mouse               TGTGCGGGAGCCATGACCTCGCTGACCCAGCGGAGCTCGGGCC
rat                 TCGGCTGGAGCCATGACCTCGCTGACCCAGCGGAGCTCGGGCC
                    *  *  ************************* ***** ****
```

#### ***Lineage-specific and State-specific genes in HECs and GCs***

##### *Ltbr:*

```
                         KLF                                     Xbp1/Creb3l2
hLTBR           CGAAGAAGGGAGGAGGCCGG-TTCCGGCCCCGCGGCCCTCACGTGCTTTCCCGGCCG-CC
chimp           CGAAGAAGGGAGGAGGCCGG-TTCCGGCCCCGCGGCCCTCACGTGCTTTCCCGGCCG-CC
rhesus          CGAAGAAGGGAGGAGGCCGG-TTCCGGCCCCGCGGCCCTCACGTGCTTTCCCGGCCG-CC
mouse           CGAAGAAAGGTGGAGGCCGGGTTCCGGGCCTGCAGCTCTCACGTGCTTTCCCGGCCACCC
rat             CGAAGAAAGGTGGAGACCGGGTTCCGGGCCTGCAGCTCTCACGTGCTTTCCCGGCCACCC
                *******.**:****.**** ****** ** **.** ******************. **
                 KLF                            ETS
hLTBR           CCTCCCGCCCCGCATCGAGGCAGACAAGCCTGTTCCTCTT----CCCTGGGCTGCGATTG
chimp           CCTCCCGCCCCGCATCGAGGCAGACAAGCCTGTTCCTCTT----CCCTTGGCTGCGATTG
rhesus          CCTCCCGCCCCGCATCGAGGCAGACAAGCCTGTTCCTCTT----CCCTTGGCTGCGATTG
mouse           CCTCCCGCCCTGCGTCGAGGCGGCCAAGCCTGTTCCTCTTCCCCCCCCCGTCGCGATTG
rat             CCTCCCGCCCTGCGTCGAGGCGGCCAAGCCTGTTCCTCT-----TCCCCGTCTCGATTG
                ********** **.*******.*.***************      **    *    ******
```

##### *Tnfrsf11a:*

```
                                  GATA
hTNFRSF11aintron        CTTGGCTGATAAGGCCTTTGCACTCAGATTAAGAGCCCTGGCCCACAAAACGTGCGGTGG
chimp                   CTTGGCTGATAAGGCCTTTGCACTCAGATTAAGAGCCCTGGCCCACAAAACGTGCGGTGG
rat                     CATGGATGATAAGGTCTTTGCCTTGAGACTAAAAGCCCTG-GCCAGGAGACTTGCAGTGG
rhesus                  CTTGGCTGATAAGGCCTTTGCACTCAGATTAAGAGCCCTGGCCCACAAAATATGCGATGG
mouse                   CTTGGATGATAAG-TCTTTGCCTTCCAACTGAAGGCCCTG--GAATGAGACATGCAATGG
                        *:***.*******   ******. *  ..* *.*..******    .*  .*.*   ***..***
                                                   Creb3L2
hTNFRSF11aintron        AACTTGCCCAAAGAGGTCGTGACAGCTGAGTCACAGGGAATAGCACTGAAACTGCGATCC
chimp                   AACTTGCCCAAAGAGGTCGTGACAGCTGAGTCACAGGGAATAGCACTGAAACTGCGATCC
rat                     AACTTGCCCAGAGG--TCATGACAGCTGAGTCACAGAGCGCAGCATTGAAACTGAGATCT
rhesus                  AACTTGCCCAAAGAGGTCGTGACAGCTGAGTCACAGGGAATAGCACTGAAACTGCAATCC
mouse                   AACTTGCCCAAAGAGGTCACGACAGCTGAGTCACAGAGAACAGCATTGAAACTGAGATTC
                        **********.**.   **. ****************.*.. **** ********..**
                                                                  CREB3l2
hTNFRSF11aintron        TGGGGCATGGAGAATGTCTATGAGGGGTG-AGCCAGGGTGCTGAGTCATGCCTCTCCGCC
chimp                   TGGGGCATGGAGAATGTCTATGAGGGGTG-AGCCAGGGTGCTGAGTCATGCCTCTCCGCC
rat                     GAG----AGGAGAGTGCCCACTGGGGCTGGAGCCAAGGTGCTGAGTCATGCCACAAC---
rhesus                  TGGGGCATGGAGAATGTCTATGAGGGGTG-AGCCAGGGCGCTGAGTCATGTCTCTCAGCC
mouse                   TGAG---AGGAGAGTGCCCACTGGGAGTGGAGCCAAGGTGCTGAGTCATG-CCCTGGGCT
                         ..     :*****.** * *   .**. ** *****.** *********** * *:

HTNFRSF11aintron        TGGCACTGACTTGCTGAGTGTCAGAGCTTCCAGAA
chimp                   TGGCACTGACTTGCTGAGTGTCAGAGCTTCCAGAA
rat                     ----------TGGCTCTGTGTCAAAGCTTCTAGAA
rhesus                  TGGCATTGACTTGCTGAGTGTCAGAGCTTCCAGAA
mouse                   CTGCATC-ACTTGCTCTGTGTCAAAGCTTCTAAAA
                                  * *** :******.****** *.**
```

##### *Relb:*

```
                                                Sp1         Xbp1/Creb3l2   NF-kB
hRelB           CCAGCCTCCCTTTCCCGGCCGGCCCCGCCCCCGCGATCACGTGACGCGTGGGGGTTTTCC
chimp           CCAGCCTCCCTTTCCCGGCCGGCCCCGCCCCCGCGATCACGTGACGCGTGGGGGTTTTCC
gorilla         CCAGCCTCCCTTTCCCGGCCGGCCCCGCCCCCGCGATCACGTGACGCGTGGGGGTTTTCC
mouse           CAAGCCTTCGCTTCCCGGCCGACCCCGCCCCCGCAGTCACGTGAGCAGCGGGGGTTTTCC
rat             CAAGCCTTCCCTTCCCGGCCGGCCCCGCCCCCGCAGTCACGTGAGCAGCGGGGGTTTTCC
                *.***** *  **********.************..********   .* ***********
                              CREB3l2                NFYA/B     Xbp1 (ERSEII-like)
hRelB           CGTTCCCCAGG-CGTGACGTCACGGCGGGGTGCAGACCAATGGGCGCGCAGGCCGGCGGC
chimp           CGTTCCCCAGG-CGTGACGTCACGGCGGGGTGCAGACCAATGGGCGCGCAGGCCGGCGGC
gorilla         CGTTCCCCAGG-CGTGACGTCACGGCGGGGTGCAGACCAATGGGCGCGCAGGCCGGCAGC
mouse           CATTCCCTTTGACGTGACGTCACGGAGGGGTACGAGCCAATGGGTGCGCAGGCCGGCGGC
rat.            CATTCCCTTTGCCGTGACGTCACGGAGGGGTGTGAGCCAATGGGCGCGCAGGCCGGCGGC
                *.***** : * *************.*****.  ...******** ************.**


                     NF-kB            Sp1         KLF
hRelB           GCCGGGGAATTCCGCCGCCCCGCCCCCGCCCGCCGCCCGCCCGGCCCGGCCCCGCGCCCC
chimp           GCCGGGGAATTCCGCCGCCCCGCCCCCGCCCGCCGCCCGCCCGGCCCGGCCCCGCGCCCC
gorilla         GCCGGGGAATTCCGCCGCCCCGCCCCCGCCCGCCGCCCGCCCGGCCCGGCCCCGCGCCCC
mouse           GCCGGGGAATTCCGCCGCCCCGCCCCCGCCCGCCGCCCGCTCGGCCCG-----GCGCCCC
rat             GCCGGGGAATTCCGCCGCCCCGCCCCCGCCCGCCGCCCGCTCGGCCCG-----GCGCCCC
                ***************************************** *******     *******
```

##### *Nfkb2:*

```
                                                                 NFYA/B
hNFKB2         GCGCCCGAGTCGCTCCGGGTTGGCTGCGCCAGTCCAG-AGTTAAACTTTCAGCCAATGAA
rheusus        GCGCCCGAGTCGCTCCGGGTTGGCTGCGCCAGTCCAG-AGTTAAACTTTCAGCCAATGAA
chimp          GCGCCCGAGTCGCTCCGGGTTGGCTGCGCCAGTCCAG-AGTTAAACTTTCAGCCAATGAA
mouse          GCGCCTGCGTTGCTCCGGGTTGGCTACAAGAGTCCTGGAGTTAAACTTTCAACCAATAAA
rat            GCGCCTTCGTCGCTCCGGGTTG-CTACAAGAGTCCTGGAGTTAAACTTTCAGCCAATAAA
               *****   .** *********** **.*..  *****:* *************.*****.**
                              Creb3L2  Creb3L2     NFKB
hNFKB2         AAAGGGCGCGAGGCGTGACGCACGGAAACGTCATGGGAATTCCCCCCTCCGGGGGGCCGA
rheusus        AAAGGGCGCGAGGCGTGACGCACGGAAACGTCATGGGAATTCCCCCCTCCGGGGGGCCGA
chimp          AAAGGGCGCGAGGCGTGACGCACGGAAACGTCATGGGAATTCCCCCCTCCGGGGGGCCGA
mouse          AAAGGGCATGGGGAGTGACGCACGGGATCGTCACGGGAATTCCCCCCCTCGGGGGGCCGA
rat            AAAGGGCATGGGGAGTGACGCACAGGAACGTCACGGGAATTCCCCCCCTCGGGGGGCCGA
               *******. *.**.*********.*.*:***** *************  ***********
                   NF-kB
hNFKB2         GAAGGGGCTTTCCCGGCCCTGAGCCCTGCTGGCAGGCGAGGTGTCGCGACCGGTCCCAGG
rheusus        GAAGGGGCTTTCCCGGCCCCAAGCCCTGCTGGCCGGCGAGGTGTCGCGACCGGTCCCAGG
chimp          GAAGGGGCTTTCCCGGCCCTGAGCCCTGCTGGCCGGCGAGGTGTCGCGACCGGTCCCAGG
mouse          -AAGGGGCTTTCCCGGCTG-GAGCCCTTTGG-CTGGAGAGGAGCAACGACCGGTCCGGGG
rat            -AAGGGGCTTTCCCGGCTG-GAGCCCTACTG-CTGGAGAGGAGCAACGACCGGTCCAGGG
                ***************   .******   *  * **.****:*  ..**********  .**
```

#### *Bhlha15 (Mist1):*

```
                                                                Xbp1/Creb3l2
hbhlha15       CCCGCCCCGAGGGCTCATTTGCATGCGTCCG-CGGCCCCTCCCGGGGCGGCTAAAGCGAC
chimpbhlha15   TCCGCCCCGAGGGCTCATTTGCATGCTTCCG-CGGCCCCGCCCGGGGCGGCTAAAGCGAC
mouseMist1     CCCTCCCAGCGAACTCATTTACATGCTTCCCGCAGCCCCGCCCGTGGTGGCTAAAGCTAC
rat            CCCGCCCAGCGAGCTCATTTACATGCTTCCCGCAGCCCCGCCTGTGGTGGCTAAAGCGAC
                ** ***.*.*..*******.***** ***  *.***** ** * ** ********* **
                                                       NF-kB
hbhlha15       GTGTCCTTGTCCCCCGTGGGCAGCCTCGGCGCGTGTGGCCGCGGATCCCCAGGTAACCCG
chimpbhlha15   GTGTCCTTGTCCCCCGTGGGCAGCCTCGGCGCGTGTGGCCGCGGATCCCCAGGTAACCCG
mouseMist1     GTGTCCTTGTCCCCCGTGGGCAGCCTCTGCGCGTACGGCCTCGAATCCCCAGGTAAACCA
rat            GTGTCCTTGTCCCCCGTGGGCAGCCTCTGCGCGTACGGCCTCGGATCCCCAGGTAAACCA
               *************************** ******. **** **.************.**.

hbhlha15       GGAG--CTGGGAACTCGGGTGGGGCCGCGGGGAGGGGACCCAGGGG-CAGGACC
chimpbhlha15   GGAG--CTGGGAACTCGGGTGGGGCCGCGGGGAGGGGACCCAGGGG-CAGGACC
mouseMist1     AGGGGTTGGGGAAACTATATAGGACTGCTGGGAGGGAACCCCAGAAGCTTGATC
rat            AGGGGTTGGGGGAACTGGATGGGACTGCTGGGAAGGGACCCAGAAG-CGGGACC
               .*.*    ***.*.  . .*.**.* ** ****.**.****..... *  ** *
```

#### *Bcl3:*

```
                          E-Box           NF-kB               Creb3l2
hBcl3          AAACACTTCTGCCTCAGCTGCCTGCTGGGGGAAATCCCTTCCCGCAGAACTTGACCGCAA
chimpBcl3      AAACACTTCTGCCTCAGCTGCCTGCTGGGGGAAATCCCTTCCCGCAGAACTTGACCGCAA
baboonBcl3     AAACACTTCTGCCTCAGCTGCCTGCTGGGGGAAATCCCTTCCCGCAGAACTTGACCGCAA
mousebcl3      AAACACTTCTGCCTCAGCTGCCGCCTCGGGGAAATCCCTTCCCGCACAACTTGACCGCAG
ratBcl3        AAACACTTCTGCCTCAGCTGCCGCCTCGGGGAAATCCCTTCCCGCACAACTTGACCGCAG
               **********************  ** ******************* ************.
                  E-box           Xbp1/Creb3l2
hBcl3          CCCAGCTGCTCTGCCTCCCCCACGTCAGCACCCGTCACTCACTCGCAGCCTCCCCCCTCC
chimpBcl3      CCCAGCTGCTCTGCCTCCCCCACGTCAGCACCCGTCACTCACTCGCAGCCTCCCCCCTCC
baboonBcl3     CCCAGCTGCTCTGCCTCCCCCACGTCAGCACCCGTCACTCACTCGCAGCCTCCCCCCTCC
mousebcl3      CCCAGCTGCTCAGCCTCCCCCACGTCAGCATCCTCCACTC-CTCACAGCCTG-CCCCTCC
ratBcl3        CCCAGCTGCTCAGCCTCCCCCACGTCAGCATCCTCCACTC-CTCGCAGCCTG-CCCCTCC
               ***********:***************** **  ***** ***.******  *******
                                              ETS                ETS
hBcl3          TGGCGCTTCCTCCAACCTTAACCCCTTCCGCTGGGGCTGAGACTTTACCGGAACGGGCAG
chimpBcl3      TGGCGCTTCCTCCAACCTTAACCCCTTCCGCTGGGGCTGAGACTTTACCGGAACGGGCAG
baboonBcl3     TGGCGCTTCCTCCAACCTTAACCCCTTCCGCTGGGGCTGAGACTTTACCGGAACGGGCAG
mousebcl3      CGGTGCTCCCTCCAACCTTAACCCCTTCCGCTGGGGCT-------TACCGGAATGGGCAG
ratBcl3        CGATGCTCCCTCCGACCTTAACCCCTTCCGCTGGGGCT-------TACCGGAATGGGCAG
                *. *** *****.************************       ******** ******
```

#### *Nod1:*

```
                                                                 NFAT
hNod1          AGCACAGAATTGAGACTTTCTCTTTAAGATCATCACAAGGCTGAGAGAAAACTGGAAAGA
chimp          AGCACAGAATTGAGACTTTCTCTTTAAGATCATCACAAGGCTGAGAGAAAACTGGAAAGA
rhesus         AGCACAGAATTGAGACTTTCTCTTTAAGATCATCACAAGGCTGAGAGAAAACTGGAAAGA
mouse          AGCCCAGAGTGGAAACTTTCTCTTGAAATG-CTCACAAGTGTGAGAGAACA-GGGAAAGG
```

```
rat               AGCCCAGGGTGGAAACTTTCTCCTGAAAACGATCACAAGTCTGAGAGAACA-GGGAAAGG
                  ***.***..* **.******** * **.:  .*******  ********.*  ******.
                                    Creb3L2                           CEBP
hNod1             AGTCACTTCTCACTTTGTGATTCAGCATTTTGCAATGCCATGCTCCATTCTTTAGAAAGC
chimp             AGTCACTTCTCACTTTGTGATTCAGCATTTTGCAATGCCATGCTCCATTCTTTAGAAAGC
rhesus            AGTCGCTTCTCACTTCATGATTTAGCGTTTTTCAACGTCATGCTCCATTCTTTAGAAAGC
mouse             CCGCGGTTCTTCCCGGGTGAGTCAGCGTG---------------------TTTAGAAGGC
rat               CCGTGTTTCTCCCTGGGTGAGTCAGCATGT--------------------TTTAGAAGGC
                  .   . **** .*   .*** * ***.*                      *******.**
                                  CEBP
hNod1             TGGG---AAGGGGTGTTGCACAGCTTCCTTCATCCCTCCTTTCCCGACCTTTGCCCGCCT
chimp             TGGG---AAGGGGTGTTGCACAACTTCCTTCATCCCTCCTTTCCCAACCTTTGCCTGCCT
rhesus            TGGG---AAGGGGTATTGCACAACTTCCTTCATCCCTCCTTTCCCGACCTTTGCCCGCCT
mouse             TGGGGAAAGG--AGGTTGAGCAACTGCCTTCATTCCACTTGACCCAACTCTT----ATCA
rat               TGGGGAAAAAGGAGGTTGAGCAACTTCCTTCGTTCCTCTTGACCCAACTCTT----ATCA
                  ****   *..  . .***..**.** *****.* **:* * :***.**  **     . *:
```

#### *Nod2:*

```
                                E-box          E-box     ETS
hNOD2             GTGTGGTAAACTTGCTCAGTTGGGCATAGCAGTTGTTAGGAAACCTGAGGCTGGTAACAC
chimp             GTGTGGTAAACTTGCTCAGTTGGGCATAGCAGTTGTTAGGAAACCTGAGGCTGGTAACAC
rhesus            GTGTGGTAAACTTGCTCAGTTGGGCATAGCAGTTGTTAGGAAACCTGAGGCTGGTAACAC
mouse             GTATAG---TTTTGCTCAGTTGGGGATAACAGTTGTTTGGAAACCTCAGGCTGGTAACAC
rat               GCGTAG---CTTTGCTCAGTTGGGGATAACAGTTGTTGGGAAACCTCAGGCTGGTAACAC
                  *  .*.*      ************* ***.******** ******** *************
                             E-box       Creb312
humanNOD2         CAGCTGTAATACCAGCTGTCCGTCTGACTCATGCAACTGTTAAAGTTGATAGGGCTGAGG
chimp             CAGCTGTAATACCAGCTGTCCGTCTGACTCATGCAACTGTTAAAGTTGATAGGGCTGAGG
rhesus            CAGCTGTAATACCAGCTGTCCGTCTGACTCATGCAACTGTTAAAGTTGATAGGGCTGAGG
mouse             CAACTGTGATGCCAGCTGTCCCTGTGACTCATGCAGATGCTAAAGCCAATGGGACTGAGG
rat               CAACTGTGATGCCAGCTGTCCGTGTGACTCATGCAGATGCTAAAGCCCATGGGGCTGAGG
                  **.****.**.********** * ***********..** *****   **.**.******
```

#### *ATG16L1:*

```
hATG16L1          GTCTGGTTCTGCTGTACGAGAACGCAGTAGTGGTTAGATATGCTGCAGAGCTATTCATAT
chimp             GTCTGGTTCTGCTGTACGAGAATGCAGTAGTGGTTAGATATGCTGCAGAGCTATTCATAT
rhesus            CTCTGGTTCTGCTGTAGGAGAATGTAGTAGCAGTTAGATATGCTGCAGAGCTATTCATAT
mouse             TTCTGGTTCTGCCACACGAGAGTGCAGTGTTGGTGAGACACCCTGTGGAGCTATAAATAT
rat               TTCTGGTTCTGCCACAGGAGAGTGCAGTGTTGGTGAGACACCCTGTGGAGCTATAAATAT
                   ***********  . * ****. * ***.   .** *** *   *** .*******:.****
                      Creb312                                   IRF
hATG16L1          CTCGTGAGTCATTGGAACAGTGCGTTTTGAAATGGTGTCCCTTCACTTTCTGTTTAG--G
chimp             CTCGTGAGTCATTGGAACAGTGCGTTTTGAAATGGTGTCCCTTCACTTTCTGTTTAG--G
rhesus            CTTGTGAATCATTGGAACAGTGCGTTTTCAAATGGTGTCCCTTCACTTTCCGTTTAG--G
mouse             CTCATGACTCATTTGAACAGCCCGTTTTGAAATGGTGTCCCTTCATTTTCTGTTTAGGAA
rat               CTCATGACTCATTTAAGCAGCCCGTTTTGAAATGGTGTCCCTTCATTTTCTGTTTAG--G
                  ** .*** ***** .*.***  ****** **************** **** ******   .
```

## *IL13ra:*

```
                                                Creb312    Creb312
hIL13ra1          GATGCCTGAGTGGATGTAAGGGAAGGG--ACAGATGACTCATATATGACTCTTGTGATTT
chimp             GATGCCTGAGTGGATGTAAGGGAAGGG--ACAGATGACTCATATATGACTCTTGTGATTT
rhesus            GATGCCTGAGTGGATGTAAGGGAAGGAG-ATAGATGACTCATATATGACTCTTGTGATTT
mouse             AGTTGATGACTGCATGTGAAGAAAAAAAGACAGATGACTCACATATGACTCTTATGGTTT
rat               GATGACTGCATGCGAAGGAAAAAAAGG---CAGATGACTCACATATGACTCTTACGATTT
                  ..*  .**. ** .:. .*...**...     ********** ***********. *.***
                          Stat
chimp             CCTTAACTTTTCCCGAAATGTGCATGCCTCTGCTAGCTGCTGTGATTGGACAGGCGAGAG
rhesus            CCTTAACTTTTCCCGAAATGTGCATGCCTCTGCCAGCTGCTGTGATTGGACAGGCGAGAG
hIL13ra1          CCTTAACTTTTCCCAAAATGTGCATGCCTCTGCTAGCTGCTGTGATTGGACAGGCGAGAG
mouse             CCTTCACTTTTCCCGAAATCTACGTGCCTCTGCCAACTCCTGAGATTGAACAAGAGAAAG
rat               CTTTCACTTTTCCCGAAATCTACATGCCTCTGCCAACTCCTGAGATTGAACAAGAGAAAG
                  * **.*********.**** *.*.********* *.** ***:*****.***.*.**.**
```

#### Nr2f2 (COUP-TFII)

```
                                             SP1/KLF
hNR2f2Promoter    TAGGACCGGGCTGCGCCCGCGGCCGCCATGGCGGGGCCGCCGCAGTCCGG
chimp             TAGGACCGGGCTGCGCCCGCGGCCGCCATGGCGGGGCCGCCGCAGTCCGG
orangutan         TAGGACCGGGCTGCGCCCGCGGCCGCCATGGCGGGGCCGCCGCAGTCCGG
```

```
makedmolerat      TAGGACCAGGCTGCGCCCGCGGTCGCCATGGCGGGGCCGCCGCGCTCCGG
rat               TAGGACCGGGCTGTTCCCTCTGCCGCCGTGGCGGGGCCGCTGCGCTCCGG
mouse             TAGGACCGGGCTGCTCCCGCTGCCGCCATGGCGGGGCCGCTGCGCTCCGG
                  *******.*****  *** * * ****.*********** **. *****
                  NFYA/B  ERSE II    XBP1
humanNR2f2        CCAATGACGGCGAGGGGGCGGCGCGCGCGCGCCTGGCCAGGCCAACCCCG
chimp             CCAATGACGGCGAGGGGGCGGCGCGCGCGCGCCTGGCCAGGCCAACCCCG
orangutan         CCAATGACGGCGAGGGGGCGGCGCGCGCGCGCCTGGCCAGGCCAACCCCG
makedmolerat      CCAATGACGGCGAGGGGGCGGCGCGCGCGCGCCTGGCCAGGCCAACCCCG
rat               CCAATGACGACGAGGGGGCGGCGCGCGCGCGCCGGGCCAGGCCAACCCCG
mouse.            CCAATGACGACGAGGGGGCGGCGCGCGCGCGCCGGGCCAGGCCAACCCCG
                  *********.*********************** ****************
                           SRF                    XBP1/Creb3l2
humanNR2f2        GCTGCTGCCTTATAAGGCGCGCCGTCGCCATGGCAACGTGCGCTAAGTTG
chimp             GCTGCTGCCTTATAAGGCGCGCCGTCGCCATGGCAACGTGCGCTAAGTTG
orangutan         GCTGCTGCCTTATAAGGCGCGCCGTCGCCATGGCAACGTGCGCTAAGTTG
makedmolerat      GCTGCTGCCTTATAAGGCGCGCCGTCGCCATGGCAACGTGCGCTAAGTTG
rat               GCGGCTGCCTTATAAGGCGCGCCGTCGCCATGGCAACGTGCGCTAAGTTG
mouse.            GCGGCTGCCTTATAAGGCGCGCCGTCGCCATGGCAACGTGCGCTAAGTTG
                  ** ***********************************************
                                      SRF            Creb3l2-like
humanNR2f2        CAGCAGTCGTGTCAAAGTTCACTATATAGAGAGCTCAGTGAGCTGATCGC
chimp             CAGCAGTCGTGTCAAAGTTCACTATATAGAGAGCTCAGTGAGCTGATCGC
orangutan         CAGCAGTCGTGTCAAAGTTCACTATATAGAGAGCTCAGTGAGCTGATCGC
makedmolerat      CAGCAGTCGTGTCAAAGTTCACTATATAGAGAGCTCAGTGAGCTGATCGC
rat.              CAGCAGTCGTGTCAAAGTTCACTATATAGAGAGCTCAGTGAGCTGATCGC
mouse             CAGCAGTCGTGTCAAAGTTCACTATATAGAGAGCTCAGTGAGCTGATCGC
                  **************************************************
                                              TATA-box  TEAD
humanNR2f2        GGAGAAGCCACTTCTGCCAGCCCCGGCGCCTATAAATCGCATTCCCTCCC
chimp             GGAGAAGCCACTTCTGCCAGCCCCGGCGCCTATAAATCGCATTCCCTCCC
orangutan         GGAGAAGCCACTTCTGCCAGCCCCGGCGCCTATAAATCGCATTCCCTCCC
makedmolerat      GGAGAAGCCACTTCTGCCAGCCCCGGCGCCTATAAATCGCATTCCCTCCC
rat               GGAGAAGCCACTTCTGCCAGCCCCGGCGCCTATAAATCGCATTCCCTCCC
mouse             GGAGAAGCCACTTCTGCCAGCCCCGGCGCCTATAAATCGCATTCCCTCCC
                  **************************************************
```

---

***Spdef:***

```
                     ETS               Srebp-binding site
hSpdef            CAAGGAAGGAAAGGTCAGGGTGGGGTGGGGTGGGCAGAGACCAAGGCGGGGCTGGG---G
chimp             CAAGGAAGAAAAGGTCAGGGTGGGGTGGGGTGGGCAGAGGCCAAGGCGGGGCTGGG---G
gorilla           CAAGGAAGGAAAGGTCAGGGTGGGGTGGGGTGGGCAGAGACCAAGGCGGGGCTGGG---G
rhesus            CAAGGAAGGAAAGGTCAGGGTGGGGTGGGGTGGGCAGAGACCGAGGCGGGGCTGGG---G
mouse             CAAGGAAGAATAGGTTTGGGTG-GGTGGGGTGGACAGGGGTCGGGGTTGGGTCTGGATTG
rat               CAAGGAAGAATAGGTTTGGGTG-GGTGGGGTGGACAGGGGTCAGGGTGGGTCTGGATTAA
                  ********.*:**** :***** **********.***.*. *..**  **    *.   .

hSpdef            ATGTGGGGGACAGAGGGCAGACCTG-GAATCATTTCAAGTCCTTCTGTGCAAA-------
chimp             ATGTGGGGGACAGAGGGCAGACCTG-GAATCATTTCAAGTCCTTCTGTGCAAA-------
gorilla           ATGTGGGGGACAGAGGGCAGACCTG-GAATTATTTCAAGTCCTTCTGTGCAAA-------
rhesus            ATGTGGGGGACAGAGGGCAGACCTG-GAATCATGTCAAGTCCTTCTGTGCAAA-------
mouse             ATTCAAGTCCTTTTGTGCAAATATGTGTGTGTGTGTGTGTGTGTGTGTGCACG-------
rat               TTTCAAGTCCTTCTGTGCAACCGTGTGTGTGTGTGTGTGTGTGTGTGTGTGTGTGTGTGT
                  :*   ..*   . : :* ***..   ** *:.* :     .:**    * ****  . .

hSpdef            ----------------------CACTCGTGTGTGCTGTGGGCCCACATTATTTACTCAGGC
chimp             ----------------------CACTCGTGTGTGCTGTGGGCCCACATTATTTACTCAGGC
gorilla           ----------------------CACTCGTGTGTGCTGTGGGCCCACATTATTTACTCAGGC
rhesus            ----------------------CACTCATGTGTGCTGTGGGCTCACATTATTGACTCAGGC
mouse             ----------------------CGCACGCACACTCCACGGGACCTTGGTATTTACCAAGGC
rat               GTGTGCGTGTGTGTGTGTGTGCGCGCGCGCGCACCACAGGCTCTTGCTATTTACCGAGGC
                                        *.* *. .  .   *  . .**. *: . **** **  ****
                                                            KLF
hSpdef            AGTGCCAACAGCTCGCTCTGCCCCTGGGCCCCACCCTGCCTGGTGCCCATGCCCTACCCG
chimp             AGTGCCAACAGCTCGCTCTGCCCCTGGGCCCCACCCTGCCTGGTGCCCATGCCCTACCCG
gorilla           AGTGCCAACAGCTCGCTCTGCCCCTGGGCCCCACCCTGCCTGGTGCCCATGCCCTACCCG
rhesus            AGTGCCAACAGCTCGCTCTGCCCCTGGGCCCCACCCTGGCCGGTGCCCATGCCCTACCCG
mouse             AGCTTCCATAGCCTGTTCT-------GTCCCCACCCTGGCCTATGCCCACACCCTATACC
rat               TGTGCCCGTAGCCTGCTCTGCCCGA-GTCCCCACCCTGGCTTGTGCCCACACCCTGTGCC
                  :*    *.. ***  * ***        * ********** *   .****** .****.  *
                               KLF
hSpdef            CTAG---GTGGCTCCACCCCTGGGGACCCCCCCCCG--CAGCCCCCCACCCCTCCTTCAT
```

```
chimp           CTAG---GTTGCTCCACCCCTGGGGACCCCACCCG---CAATAACAAAAAAAAATTTTAT
gorilla         CTAG---GTGGCTCCACCCCTGGGGACCCCCCCCTCCGCAGCCCCCCATCCCTCCTTCAT
rhesus          CTAG---GTGGCTCCACCCCTGGGGACTCCCCCCTG--CAGCCCCCCGCCCCTCCTTCAT
mouse           CCAGGTGGTGGCCCCACCCAAGGAGAGTCTTCCCCC----AGTTTCCATCCCTCCTCCTC
rat             CCAGGTGGTGGCCCCACCCGAGGAGAG-CCTCCTCC----AGTCTCCATCCCCCCTCC--
                * **   ** ** ****** :**.**  *  **       .    ... ... . *
                                             Spdef                Xbp1/Creb3l2
hSpdef          CACCCCCTGCCTGCCTCCTGCCCCGTCCACATCCCCAGAGCCCCCATGCCTGCCACGTTA
chimp           CTCCCCCATTTTTTTTTCTATCTGACCCACATCCCCAGAGCCCCCATGCCTGCCACGTTA
gorilla         CACCCCCTGCCTGCCTCCTGCCCCGTCCACATCCCCAGAGCCCCCATGCCTGCCACGTTA
rhesus          CACCCCCTGCCTGCCTCCCGCCCCGTCCACATCCCCAGAGCCCCCATGCCTGCCACGTTA
mouse           CACCTCACAATGGCCTCCTGCCCCGTCCACATCCCCAATGGCCCCATGCCTGCCACGTTA
rat             -ATCCCACAATGGCCTCCTGCCCCGTCCACATCCCCAGCGGCCCCATG-CTGCCACGTTA
                 : * *.        * * . *  . ***********. * ******* ***********

hSpdef          GGACAGTCTCTGCCAACACCCTGGGCGCCATGCCAGGGAGACAGAAGCTTTTCCAAGAAG
chimp           GGACAGTCTCTGCCAACACCCTGGGCGCCATGCCAGG-----------------------
gorilla         GGACAGTCTCTGCCAACACCCTGGGCGCCATGCCAGGGAGACAGAAGCTTTTCCAAGAAG
rhesus          GGACAGTCTGTGCCAACACCCTGGGCGCCATGCCAGGGAGACAGAAACTTTTCCAAGAAG
mouse           GGACAGTCTCTGCCAGCTCCCTGGGCCCCATGCCCAGCCTCTAGCTTTTCCAAGAGGCAG
rat             GGACAGTCTCTGCCAGCTCCCTGGGCCCTGTGCCCAGCCGCTAGCTTTTCCAAGAG----
                ********* *****.*:******** * .****..*
```
