## Supplementary material for "Coordinated Transcriptional Networks Program Organelle Expansion and Metabolic Flows for High Endothelial Morphology and Function": https://drive.google.com/drive/folders/16LrJoVh-ugeEoXfNzdhAxGWC4Km_MaxT?usp=sharing

**Data S3:**
Additional Phylogenetic Analysis of Key Genes Controlling Specialized Glycosylation in stomach resident plasma cells and pituitary gland.

IgGs have invariant N-linked glycosylation at 297N – carrying Fucose (by **FUT8**) and 2-6-linked sialic acid (**St6gal1**).

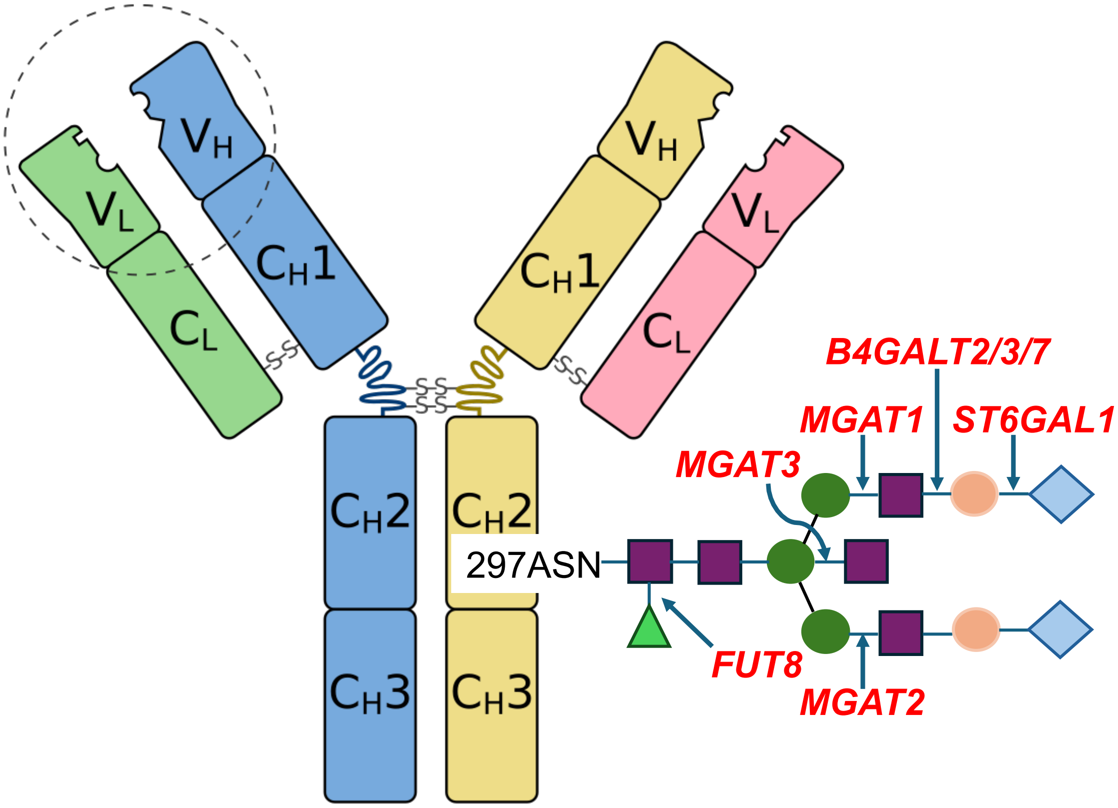


**Fut8**

```
                                                                    ERSE-II
hFUT8           CAAGGCTACAGGGAAGAGTTTGGAACGGGAAGCTCATCTTCCGGCCCTCTGATTGGCCGG
chimp           CAAGGCTACAGGGAAGAGTTTGGAACGGGAAGCTCATCTTCCGGCCCTCTGATTGGCCGG
rhesus          CAAGGCTACAGGGAAGAGTTTGGAACGGGAAGCTCATCTTCCGGCCCTCTGATTGGCCGG
mouse           CCAGGCTACAGGGAAGAGTTCGGCACGGGAAGCTCATCTTCCGGCCCTCTGATTGGCCAG
rat             CCAGGCTACAGGGAAGAGTTCTGCACGGGAAGCTCATCTTCCAGCACTCTGATTGGCCAG
                *.******************  *.******************.**.************.*

                        Xbp1      ERSEII-like   NFYA/B                  Xbp1
hFUT8           CTCGCACTCCACTCACGCGGCGCGCAGCTCTGATTGGCCTCGGCGGCACCCCT-CGTCCC
chimp           CTCGCACTCCACTCACGCGGCGCGCGGCTCTGATTGGCCCCGGCGGCACCCCT-CGTCCC
rhesus          CTCGCACTCCACTCACGCGGCGCGCGGCTCTGATTGGCCTCGGCGGCACCCCT-CGGCCC
mouse           TTCGCACTCCACTCACTGGTTTTCCCGCTCTGATTGGCCCCGGCAGCACCCCTTCGGCCC
rat             TTCTCACTCCACTCACTGGTTTTCCCGCTCTGATTGGCCCCGGTGGCACCCG----GCCG
                 ** ************  *     * ************* *** .******      ** 

hFUT8           GCGACTACTTTGTGTGCTGGGGCGGCGCGCTCCGGTCCTCCCGCTCAGCTGGCGGTCTGG
chimp           GCGACTACTTTGTGTGCTGGGGCGGCGCGCTCAGGTCCTCCCGCTCAGCTGGCGGTCTGG
rhesus          GCGACTACTTTGTGTGCTGGGGCGGCGCGCTCAGGTCCTCCCGCTCAGCTGGCGGTCTGG
mouse           GCGACTACTTTGTGTGCCCTGGCTGCGCGGCTGGGTCCTCCCGCCCAGCTCGCCCTCCCG
rat             GCGACTACTTTGTGTGCCCTGGCGGCGCGGCTGGGTCCTCCCGCTTAGCTCGCCCTCTAG
                *****************   *** *****    ***********  **** **  **  *
```

**St6gal1**

```
                                   Xbp1/Creb3l2
hSt6gal1        CCTGCGGATCCACAACAAACCCACGTGCGCACAGCCCCGGGGAACACGGATTTCTCCACG
chimp           CCTGCGGATCCACAACAAACCCACGTGCGCGCAGCCCCGGGGAACACGGATTTCTCCACG
rhesus          CCTGCGGATCCACAACAAACCCACGTGCGCGCAGCCCCGGGGAACACCGATTTCTCCACC
mouse           CCTGCGCACCCATAACAAACCCACGTGTGCGGGGACGCAGG---------AACCTACACT
rat             CCTGCGCACCCATAACAAACCCACGTGTGCGGGGCCGCAGG---------AACCTACACC
                ****** * *** ************** **. .*.* *.**         :: **.***

hSt6gal1        GCACCGTCAGTATGGG-----------------------GTGGG------------GTGG
chimp           GCACCGTCAGTATTGG-----------GTGGGGTGGCGGGGGGGGCAGTGCGCCTTGCAG
rhesus          GCACCGTCAGTGTTTGGCTGGGGGGGTGGGGGGAGTGGGGAGGGGCGGTGCGCCTCGCAG
mouse           GCAGCGTCTGTATTGT-----------------------GTAAGGCGTTGA--CTCGCGG
rat             ACCGCGTCTGTATTGT-----------------------GTAAGGCGTTGC--CTTGCAG
                .*. ****:**.*                          * ..*            * .*

hSt6gal1        CGTGGGGG---CATTGCGCCTTGCAGAGTCTGGGTTTACGGATCGACAAATTGAGGCCCG
chimp           AGTCCGGGTTCCTTCAAATCCTGCCTT--CTGGGTTTACGGATCGACAAATTGAGGCCCG
rhesus          AGTCCGGGTTCCTTCAAATCCTGCCTT--CTGGGTTTACGGATCGACAAATTGGGGCCCG
mouse           AGTGAGGGTCTCTTTAGATCTTGCCCA--CTTTGGTAACAGAGCAGGAAAACAAGCCCGG
rat             AGTGTGGGTCCCCTTAGATCTTGCTCA--CCGGGGTTACAGATCAGGAAACTAAGTCTGG
                .**  ***   * * . . * ***  :  *   * *:**.** *.. ***  ..* *  *
                                                     Xbp1/Creb3l2
hSt6gal1        CGGCAGCGGAGGG--GGCACAGGCACCATAACAAACGCACGTGCGCCCTGCCCGGAGAGC
chimp           CGGCAGCGGAGGG--GGCACAGGCACCATAACAAACGCACGTGCGCCCTGCCCGGAGAGC
rhesus          CGGCAGCGGAGGG--GGCACAGGCACCATAACAAACGCACGTGCGCCCTGCCCAGAGATC
mouse           TGGAAGGAGAGGGGGAGCGCAGGCACCACAACAAACGCACGTGTGCCCCG----------
rat             TGGAGGGAGAAGG--AGTGCAGGCACCACAACAAACGCACGTGTGCCCCGCCAGGG---A
                 **..* .**.**  .* .********* ************** **** *

hSt6gal1        TGGACGCACTGCATACCCTTAGGGGCCGCCCTGCGGTGCCGGGCCATTGTCTCAGCCTCG
chimp           TGGACGCACTGCATACCCTTAGGGGCCGCCCTGCGGTGCCGGGCCATTGTCTCAGCCTCG
rhesus          TGGACGCACCGCAAACCCTTAGGGGCTGCCCTACGGTGCCGGGCCATTGTCTCAGCCTAG
mouse           -------------------GCTGAACCTCTTTTGG---TCAGGTTATTGTTCCAGCCAAA
rat             TCGGGGCGCAGCCATCCCTGCTGAACCTCCTTTGG---GTAGGTTATTGTTCCA------
                                    . *..*  *  *  *     .**  *****  **
```

In the pituitary gland including lactotrophs, corticotrophs and gonadotrophs, several peptide hormones—including lutropin—undergo specialized glycosylation steps that generate distinct oligosaccharide structures critical for controlling their circulating half-lives. Two enzymes are central to this process: the β-1,4-N-acetylgalactosaminyltransferase *B4GALNT3/4* and the sulfotransferase *CHST8*, each of which contributes specific modifications that fine-tune hormone stability and systemic bioavailability**.**

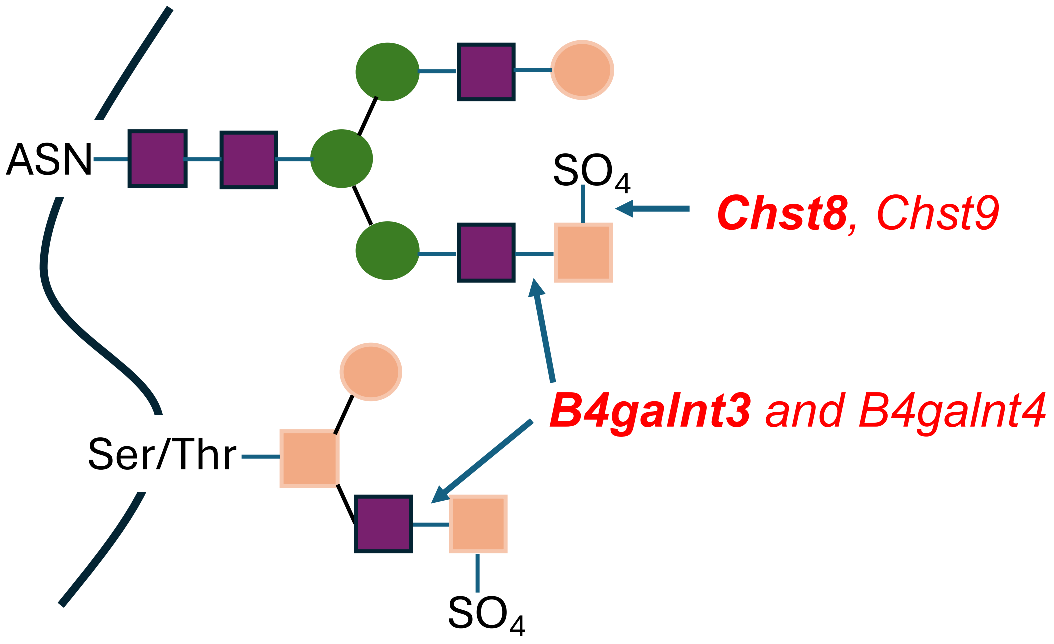


**B4galnt3**

```
                      TEAD
hB4galnt3       AACACTGAATGGGAAGAGAGGGAGGCTTTCTAGGGGATTTAAATGATACTGAAACCCGGA
rhesus          AACACCGAATGGAGACAGAGGGAGGCTTTCGGGGGGATTCAGATGATACTGAAACCCGGA
chimp           AACACTGAATGGGAAGAGAGGGAGGCTTTCTAGGGGATTTAAATGATACTGAAACCCGGA
mB4galnt3       AACATGGAATGGTGAATGGAGCAG-CCTCCCGGGGGATTTAGATGGTACGGAAACCCAGA
                ****  ****** .* :*..* ** * * * .******* *.***.*** *******.**

hB4galnt3       TACGGAATAGGAGAAAAAATACCAACTTGCTCTCAGGGTGTCTTTACTCTCCCCAGTGCA
rhesus          TATGGAACAGGACAAAAAACAGCATCTTGCTCTCAGGGTGTCTTTATTCTCCCGAGTGCT
chimp           TACGGAATAGGAGAAAAAATACCAACTTGCTCTCAGGGTGTCTTTACTCTCCCCAGTGCA
mB4galnt3       TACAGAATTAGAGAAGAAACAACAGCCT-TGCTCAGGGACTCTTCGCTCTCCCAAATGTT
                ** .*** :.** **.*** * ** * *   *******: **** . ****** *.** :
                Nkx2-1   Creb312
hB4galnt3       GATTTCATTGACCTAAATCAGGTCTCTGCTCCTATTTGCTGGTACCTAGTTCAAGAGAGT
rhesus          GATTTCATTGACCTAAATCAGGTCTCCGCTCCTGTATGCTGGTACCTAGTTTAAGAGAGT
chimp           GATTTCATTGACCTAAATCAGGTCTCTGCTCCTATTTGCTGGTACCTAGTTCAAGAGAGT
mB4galnt3       GATTTCATTGACCTAAATCAAGCCTCTGCTCCCATGTGCGGGGACCTATTTTAAGAGAGT
                ********************.* *** ***** .* *** ** ***** ** ********
                                   ETS
hB4galnt3       CTTATAAATACACAAAGCCGGAAATGCCTG-----------GGTAGGGGGGTGGCAAGAG
rhesus          CCTATAAATACACAAAGCCGGAAATGCCTG-----------GGTAGCGGGGTGGCAGGAG
chimp           CTTATAAATACACAAAGCCGGAAATGCCTG-----------GGTAGGGGGGTGGCAAGAG
mB4galnt3       CATAAATATACAAACT--CGGAAATGCCTGGTGGTGGTGGTGGTGGGGTGGTGGCAAGAG
                * **:*:*****.*.:  ************           ***.* * *******.***
```

**B4galnt4**

```
                                                     SP1
hB4galnt4         --CG--CCCGCTGTTCTCCGCTCTGCGATTTGGGGGGAGGGGGCGCGGTTCGGCTCGGCC
chimp             --CG--CCCGCTGTTCTCCGCTCTGCGATTTGGGGGGAGGGGGCGCGGTTCGGCTCGGCC
greenmonkey       --CG--CCCGCTGTTCCCTGCCCTGCGATTTGTGAGGAGGGGGCGCGGTTCGGCTCGGCC
mouse             TGCTGTCCTGCTCCTCCCAGCTCAGCTATATTGGGGGAGGGGTCGCGGTTCTGTTCGG-C
rat               TGCTGTCCTGTTCCAACCAGCTCCGCTATATTGGGGGAGGGGTCGCGACTCTGTTAGG-C
                    *   ** * *  :. * ** * ** **:*  *.******* ****. ** * *.** *
                             Creb3l2
hB4galnt4         CGGACGCTCGATGACGTCGGCAGTGACGCGCGCGACCCTCCCAGAGAGGGTCCCTG----
chimp             CGGACGCTCGATGACGTCGGCAGTGACGCGCGCGACCCTCCCAGAGAGGGTCCCTG----
greenmonkey       CGGACGCTCGATGACGTCGGCAGTGACGCGCGCGACCCTCTCTGGGAGGGTCCCTG----
mouse             GGGACGCCCAGTGACGTCAGCAGTAACGCCAGGGACCCTCG--GGGCTCATCCCTACCTT
rat               GGGACGCCCAATGACGTCAGCAGTAACGCCAGGGACCCTCG--GGACTCATCCCTAACCT
                   ****** *..*******.*****.**** .* *******   *...  .*****.

hB4galnt4         ------ACTCTGGGATCTGGCTTGGATCCGGTAGTGAGGGG-CTGACGGTGGGCTCCTGG
chimp             ------ACTCTGGGATCTGGCCTGGATCCGGTAGTGAGGGG-CTGACGGTGGGCTCCTGC
greenmonkey       ------GCTCTGGGATCCGGCCTGGATCTGGTAGTGAGCGG-CTGACGG-GGGCTCCCGG
mouse             TTCCG-ACCCTGGAATCTGGCTTAGATCCGGTGGTGATGGAGTTAACGGTGGGCCCTTGG
rat               TTTTGGACCCTGGAATCTGTCTTAGATCCGGTGGTGATGGAATTAACGGTGGGCCCTTGG
                        .* ****.*** * * *.**** ***.****  *.  *.**** **** *  *
```

---

**Chst8**

```
                                              Creb3l2
hChst8            CAGAGCTGGGGAGCAGGGAAGCAGGTCTGTGAGTCACCCCATTTCCCGTTGTGCCCATGC
chimp             CAGAGCTGGGGAGCAGGGAAGCAGGTCTGTGAGTCACCCCATTTCCCGTTGTGTCCATGC
rhesus            CAGAGCTGGGGAGCAGGGAAGCAGGTCTGTGAGTCACCCTGTTTCCCGTTGTGCCCACAC
mouse             CAGAGCTGGGGTGGGGGAGAGCAGGACTGTGAGTCACCCTACTTCCTGGTGTACCCATGC
rat               TAGAGCTGGGGTGGGGGAGAGCAGGCCTGTGAGTCACCCTACTTCCTGGTGTACCCATGC
                   **********:* .**..****** ************* . **** * ***. *** .*
                                 Creb3l2-like                        ER-response
hChst8            TGCTGGCTGCAGGCCCTGAGCTGAGTGGCCACCGAGCTGCAGGCAACAGCCCAGGTCGCT
chimp             TGCTGGCTGCAGGCCCTGAGCTGAGTGGCCACCGAGCTGCAGGCAACAGCCCAGGTCGCT
rhesus            TGCTGGCTGCAGGCCCTGAGCTGAGTGGCCGCCGAGCTGCAGGCAACAGCCCAGGTCGCT
mouse             TGCTGGCTGTGGGCCCTGAGCTGAGCGGCTGCCGAGCTGCAGGCAACAGCCCAGGTCGCT
rat               TGCTGGCTGCAGGCCCTGAGCTGAGAGGCTGCCGAGCTGCAGGCAACAGCCCAGGTCGCT
                  ********* .************** *** .*****************************

hChst8            GTCACCTTGTCCTCACTGT--CTGGGCCTGAGCCAGCTCCTCCCTGACTGGGCCAGACTC
chimp             GTCACCTTGTCCTCACTGT--CTGGGCCTGAGCCAGCTCCTCCCTGACTGGGCCAGACTC
rhesus            GTCACCTTGTCCTCACTGC--CTGGGCCTGAGCCAGCTCCT-CCTGGCTGGGCCAGACTC
mouse             GTCACCTTGTCCTCACTCTGCCTGGGCCAGAGCCAGTTGCTTCCAAGCTCTGACAGCCTT
rat               GTCACCTTGTCCTCACTCTGCCTGGGCCAGAGCCAGCTGCTTCCAAGCTCGGCCAGCCTT
                  *****************    *******:******* * ** **:..**  *.***.**
```
